# Genomics of Neotropical biodiversity indicators: two butterfly radiations with rampant chromosomal rearrangements and hybridisation

**DOI:** 10.1101/2024.07.07.602206

**Authors:** Eva SM van der Heijden, Karin Näsvall, Fernando A. Seixas, Carlos Eduardo Beserra Nobre, Artur Campos D Maia, Patricio Salazar-Carrión, Jonah M Walker, Daiane Szczerbowski, Stefan Schulz, Ian A Warren, Kimberly Gabriela Gavilanes Córdova, María José Sánchez-Carvajal, Franz Chandi, Alex P Arias-Cruz, Nicol Rueda-M, Camilo Salazar, Kanchon K Dasmahapatra, Stephen H Montgomery, Melanie McClure, Dominic E Absolon, Thomas C Mathers, Camilla A Santos, Shane McCarthy, Jonathan MD Wood, Gerardo Lamas, Caroline Bacquet, André Victor Lucci Freitas, Keith R. Willmott, Chris D Jiggins, Marianne Elias, Joana I Meier

**Author notes:** **Corresponding author**: Joana Isabel Meier. **Author contributions:** PSC, JIM, ESMH, KGGC, MJSC, DS, FC, APAC, CB, NRM, CS, KKD, NN, SHM, MM, KRW, AVLF, CDJ and ME collected samples. GL, KRW and ME oversaw the butterfly identification and taxonomic revision. ESMH, JIM, JMW and IAW performed lab work. ESMH mapped and filtered the sequencing data, and ran the phylogenetic and hybridisation analyses. KN did the chromosomal rearrangement analysis. CEBN, ACDM, DS, SS and AVLF analysed the androconial semiochemicals. FAS ran the MSCi analysis. DEA, TCM, CAS, SM, KN and JW assembled and curated the new genomes. JIM designed the study with contributions from ESMH, ME and CDJ. ESMH, JIM and KN wrote the manuscript with contributions from all authors.

## Abstract

A major question in evolutionary biology is what drives the diversification of lineages. Rapid, recent radiations are ideal systems for addressing how new species arise because they still show key morphological and ecological adaptations associated with speciation. While most studied recent radiations have evolved in an insular environment, less research has been carried out on continental radiations with complex species interactions. *Melinaea* and *Mechanitis* butterflies (Nymphalidae: Ithomiini) have rapidly radiated in the Neotropics. They are classical models for Amazonian biogeography and colour pattern mimicry and have been proposed as biodiversity indicators. We generated reference genomes for five species of each genus, and whole-genome resequencing data of most species and subspecies covering a wide geographic range to assess phylogeographic relationships, patterns of hybridisation and chromosomal rearrangements. Our data help resolve the classification of these taxonomically challenging butterflies and reveal very high diversification rates. We find rampant evidence of historical hybridisation and putative hybrid species in both radiations, which may have facilitated their rapid diversification. Moreover, dozens of chromosomal fusions and fissions were identified between congeneric species, and even some within species. We conclude that interactions between geography, hybridisation and chromosomal rearrangements have contributed to these two rapid radiations in the highly diverse Neotropical region. We suggest that rapid radiations may be spurred by repeated periods of geographic isolation during Pleistocene climate oscillations, combined with lineage-specific rapid accumulation of incompatibilities during allopatric phases, followed by secondary contact with some gene exchange.

**Significance Statement:** Understanding factors contributing to rapid speciation is a key aim of evolutionary biology. Here we focus on two rapid radiations of Neotropical butterflies. Our genomic data with broad taxonomic and geographic coverage reveal rampant hybridisation and chromosomal rearrangements, each likely contributing to the high diversification rates. Our study highlights the use of genomic data to resolve taxonomically challenging species groups and elucidate drivers of diversification in rapid radiations. We show that for biodiversity hotspots with recent radiations, barcoding is insufficient to characterise species richness due to gene flow and recent speciation. The taxonomic implications of both introgression and karyotype diversity for species delimitation are important to consider during monitoring and management of biodiversity in these vulnerable habitats.

## Introduction

Rapid radiations, where a lineage diversifies into many different species over a short time period, are ideal systems for studying how new species evolve (1, 2). They can be driven both by non-adaptive processes, such as the accumulation of differences during periods of allopatry leading to incompatibilities upon secondary contact, and by adaptive processes such as adaptation to different ecological niches or sexual selection for different traits and preferences (3, 4). Sympatric radiations require some degree of niche differentiation among the species for stable coexistence and sufficient reproductive isolation such that incipient lineages do not merge (2).

While most lineages do not readily radiate even in the face of ecological opportunity, some lineages are particularly prone to rapid radiations, and do so repeatedly. We are only starting to understand the factors explaining these lineage-specific differences (5). Most knowledge stems from well-studied radiations that evolved in insular environments with little competition with other species and a relatively simple and geographically limited environment (e.g. Darwin’s finches on the Galapagos Islands, cichlid fishes in lakes, *Anolis* lizards on the Caribbean islands or Hawaiian silverswords) (2, 6). However, many rapid radiations evolved on the much more complex continents and much less is known about drivers leading to their diversification (7–9). Reduced competition in insular environments allows niche specialisation without being immediately outcompeted when a lineage is not yet well-adapted to its environment. However, in large continental areas such as the hyperdiverse Neotropics, competition is much stronger limiting ecological speciation. On the other hand, large and complex environments on continents may provide more opportunity for allopatric divergence than a small island. The speed of accumulating incompatibilities in allopatry may thus be more important in rapid radiations on continents than insular environments.

Here, we study the drivers of diversification in two rapid continental radiations of the Neotropical butterfly tribe Ithomiini. Ithomiine butterflies (Nymphalidae: Danainae, ca. 400 species in 42 genera) are found across Central and South America (10, 11). They constitute a substantial part of the butterfly species assemblage and are regarded as good indicators of spatial patterns of biodiversity in the Neotropics, the most biodiverse areas in the world (10, 12, 13). Sequestration of pyrrolizidine alkaloids from Asteraceae and Boraginaceae plants render most Ithomiini unpalatable (14–18), and their colour patterns advertise this unpalatability to predators. They form Müllerian mimicry rings, where locally co-occurring species have converged in colour pattern, and thus share the cost of educating predators (10, 19, 20). We focus on two ithomiine genera, *Melinaea* and *Mechanitis*, which have diversified exceptionally fast with most species younger than a million years (11, 21). Hitherto, the study of these radiations has been hampered by taxonomic challenges. *Melinaea* and *Mechanitis* are among the most taxonomically difficult of Ithomiini, as Fox noted (1967) (22): ‘these insects [are] so thoroughly confusing and so thoroughly confused by my predecessors’. The species do not differ in genital or other morphological characteristics and show substantial intraspecific wing pattern variation and mimicry between taxa. Barcoding does not reliably distinguish species either (23, 24). As prior studies have only used few or no genetic markers or did not have broad geographic coverage, the taxonomy is still partially unresolved, despite many taxonomic revisions (22, 24–26, e.g. 27–29).

While the exact causes of their rapid radiations are unknown, different contributing factors have been proposed. Ecological adaptation may be relevant as species show differences in microhabitats, host plants, mimicry rings, and altitude, but on the other hand many species share habitats and host plants, and the majority of species occurs in the lowlands (25, 30–33). Moreover, where species co-exist, they converge in colour patterns and thus assortative mating may rely more strongly on chemical cues. Ithomiine species differ in male-specific androconial compounds (chemical compounds secreted from specialised wing scales where the fore- and hindwing overlap), which likely act as pheromones (Trigo et al. 1996; Schulz et al. 2004; Blow et al. 2023).

Allopatric accumulation of differences could also have played a role in the rapid diversification of ithomiini, as this could have occurred in different rainforest refugia during climatic oscillations e.g. in the Pleistocene (26, 34, but see 35, 36) or on opposite sides of geographic barriers such as the Andes. Both climatic refugia and the Andes have been proposed as “speciation pumps” in the Neotropics (37), also for Ithomiini (38), where periods of allopatry followed by secondary contact create favourable circumstances for speciation.

Another factor that might contribute to the diversification of ithomiine butterflies is hybridisation. Phylogenetic studies using a limited number of markers have revealed mito-nuclear discordances and paraphyletic taxa in Ithomiini (23, 36). This could be due to limited geographic or genetic resolution, incomplete lineage sorting in the rapidly speciating lineages, or introgressive hybridisation. While gene flow between sister lineages can homogenise their gene pools, opposing speciation, recent studies have shown that sometimes introgressive hybridisation from more distant relatives can facilitate rapid diversification by enriching the genetic diversity with novel, potentially adaptive variants or contributing to the origin of new hybrid species (39–43). Admixture has been shown to kickstart adaptive radiation (e.g. 44, 45), facilitate parallel adaptation (e.g. 42, 46), and novel adaptations (e.g. 47), but the role in ithomiini diversification is hitherto unknown.

Ithomiini butterflies have an unusually high diversity in chromosome number (48), which could also contribute to their rapid diversification. Offspring from parents with different karyotypes may suffer reduced fitness, due to mismatch in pairing of homologous chromosomes that results in aneuploidy, meiotic failure or hybrid sterility (49, 50). Furthermore, chromosomal rearrangements might facilitate divergence in the face of gene flow by accumulation of incompatibilities in low recombining regions (51) or if they link together co-adapted variants (52, 53). In *Melinaea* and *Mechanitis* butterflies, chromosome counts range from 13 to 30 (48), and chromosomal rearrangements likely contribute to reproductive isolation, as a cross between two closely-related *Melinaea* species with different karyotypes resulted in nearly sterile hybrids (54). However, pervasive intraspecific variation in chromosome counts (48, 54) indicates that not all rearrangements reduce fitness and their role in speciation thus remains an open question.

Here, we use five reference genomes of each genus and whole-genome resequencing data of almost all species and many subspecies, to resolve taxonomic uncertainties and explore whether geography, introgressive hybridisation or chromosomal rearrangements may have played a role in the rapid diversification of *Mechanitis* and *Melinaea* butterflies.

## Results

### Taxonomic revision

In this manuscript, we adhere to the ‘genotypic cluster’ species concept, which defines a species as ‘a morphologically or genetically distinguishable group of individuals that has few or no intermediates when in contact with other such clusters’ (55). Through whole-genome resequencing of 135 *Mechanitis* and 109 *Melinaea* individuals from across South and Central America, with additional individuals from the outgroup genera *Forbestra*, *Eutresis* and *Olyras*, we shed light on the phylogenomic relationships and taxonomy (Fig. 1). Our results, laid out in the following sections (see also Text S1), confirm species versus subspecies status for most known taxa in agreement with (29), support two recent species reclassifications (*Mel. tarapotensis* (54), *Mel. mothone* (56)) and reveal three additional taxa that need to be elevated to species level (*Mel. maeonis* **stat rest** (Hewitson 1869), *Mec. nesaea* **stat rest** (Hübner 1820), *Mec. macrinus* **stat rest** (Hewitson 1860), Fig. 1; Text S1). The placement of *Mel. menophilus mediatrix* (French Guiana) has been uncertain (56); our dataset places it as the most divergent subspecies of *Mel. menophilus*. A revised, annotated taxonomic list for these two genera can be found in Text S2.

**Figure 1.**
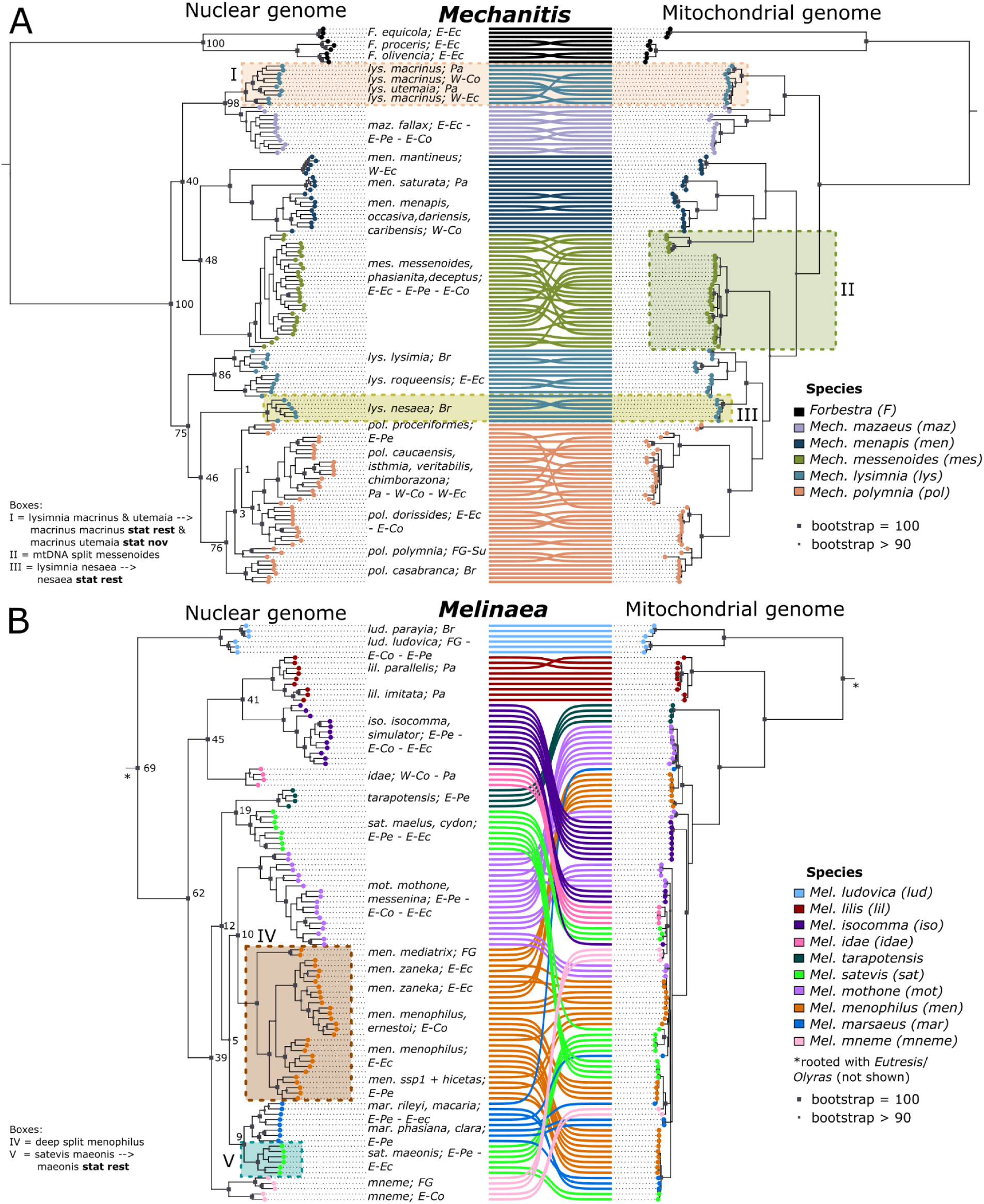
Rampant cytonuclear discordance in Mechanitis and Melinaea and a need for taxonomic revision. Co-phyloplot showing the nuclear and mitochondrial phylogenies of 135 Mechanitis and 109 Melinaea individuals. Nuclear phylogeny (left) based on 537,500 SNPs for Mechanitis **(A)** and 784,526 SNPs for Melinaea **(B)** and full mitochondrial genome phylogeny (right). The coloured circles and connecting lines indicate the currently classified species (25, 29). Br=Brazil; FG=French Guiana; E-Co=eastern Colombia; W-Co=western Colombia; E-Ec=eastern Ecuador; W-Ec=western Ecuador; E-Pe=eastern Peru; Pa=Panama, Su=Suriname. The coloured boxes highlight key findings. The node labels show concordance factors, indicating the percentage of trees produced for windows across the genome that contain that clade.

### Mitonuclear discordance

In both *Mechanitis* and *Melinaea*, we find rampant mitonuclear discordance, *i.e.* mismatches between mitochondrial and nuclear phylogenies (IQtree2; maximum likelihood) (Fig. 1). For instance, *Mec. nesaea* is sister to *Mec. polymnia* in the nuclear phylogeny, but to *Mec. lysimnia* in the mitochondrial phylogeny (Fig. 1A - box III; Fig. S1-2). This result could either be indicative of incomplete lineage sorting (ILS) or admixture (more details below). *Mec. messenoides* harbours two divergent mitochondrial haplotypes, consistent with barcoding results (23, 25) (Fig. 1A - box II), one clustering with the *polymnia*-*lysimnia* clade, and the other one with its nuclear sister species, *Mec. menapis*. The mitochondrially divergent *Mec. messenoides* individuals do not form separate clades in the nuclear phylogeny, nor do they differ in collection location or subspecies (Table S1; Fig. S1-2).

While the *Melinaea* species are clearly differentiated in the nuclear phylogeny, the mitochondrial phylogeny shows almost no variation among species, confirming previous barcoding results (56) (Fig. 1B; Fig. S3-4). *Mel. ludovica*, *Mel. tarapotensis* and *Mel. lilis* are the only species that form monophyletic mitochondrial clades that are no part of the shallow clade without species differentiation. Notably, *Mel. ludovica* is the only species that is also an outgroup to the other *Melinaea* species in the nuclear genome, whereas *Mel. lilis* and *Mel. tarapotensis* are sister to *Mel. isocomma* and *Mel. satevis*, respectively.

### Calibrated phylogenetic tree

We approximated the divergence times in the two genera using a Bayesian MCMC method for inferring trees (BEAST2), with one individual representing each lineage to produce a phylogeny calibrated with divergence times from (11) (Fig. 2). The seven *Mechanitis* species are estimated to have diversified within the past 1.36 million years, resulting in a speciation rate of 1.431 speciation events per lineage/My (assuming a pure birth model with a constant speciation rate). The ten species of the core *Melinaea* clade (excluding the most divergent species, *Mel. ludovica*) have diversified in the past 1.41 million years (Fig. 2B), giving a speciation rate of at least 1.633. Of the four potential *Melinaea* species missing in our analysis (*Mel. ethra*, *mnasias, mnemopsis* and *scylax*), two likely form part of the core clade (11, 57), which would increase the speciation rate to 1.672.

**Figure 2.**
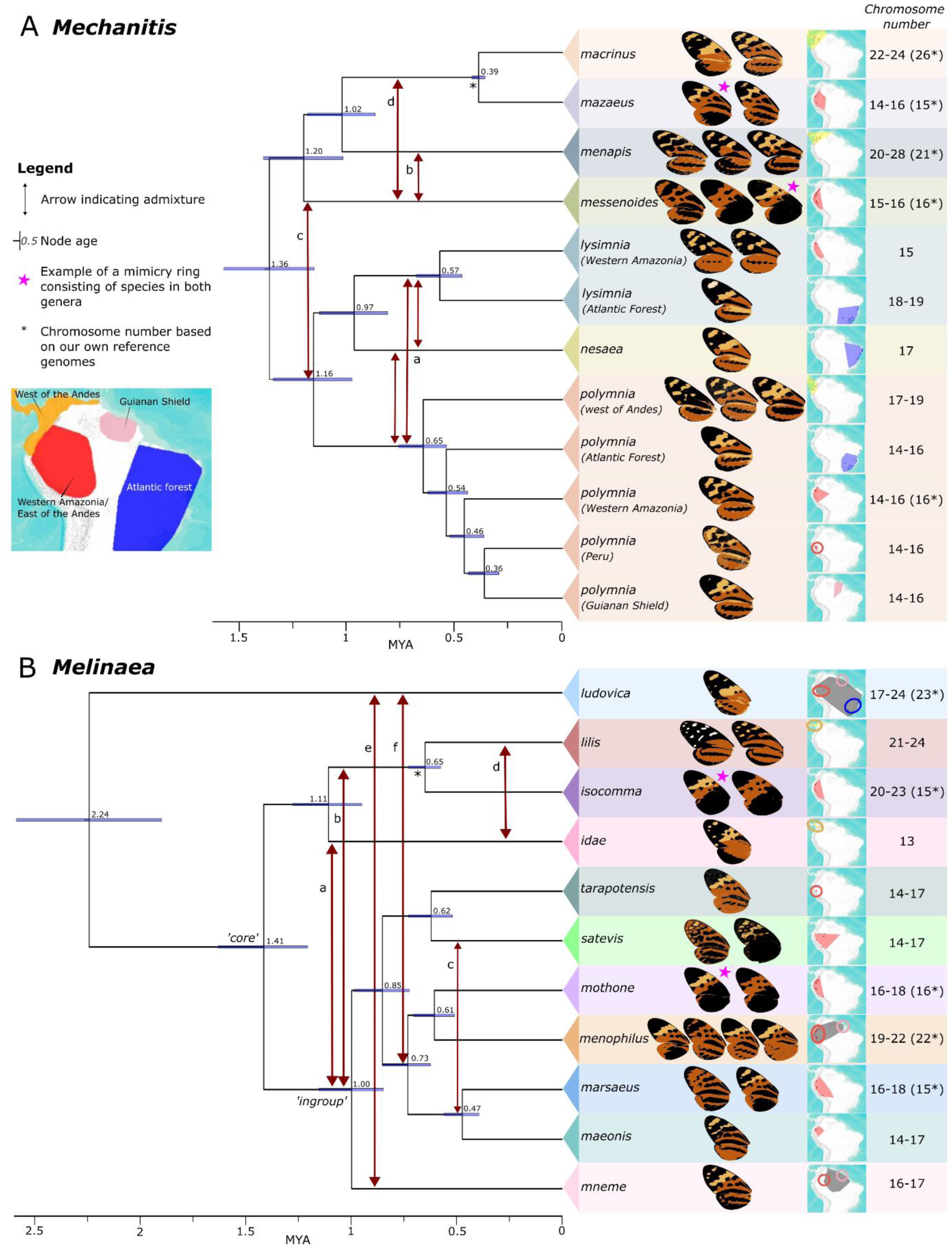
Calibrated phylogeny of Mechanitis and Melinaea butterflies with evidence of introgression and biogeographic patterns. Time-calibrated BEAST2 phylogenies of Mechanitis **(A)** and Melinaea **(B)** with the newly proposed species classification and secondary calibrations from (11) (asterisk at node indicates which node was used to calibrate). One individual was included at the species level, or subspecies-level if they were very divergent. The regions in the overview map are based on the combined distribution of our subspecies and samples from (10) with region names adapted from (11). Arrows between clades indicate potential hybridisation events (based on AIM, Fbranch and BPP). The node labels indicate the age as obtained by the calibration. ‘Core’ in the Melinaea phylogeny indicates the core clade of fast diverging Melinaea, and ‘ingroup’ is a clade referred to in the text. For each clade, cartoon wings based on representative colour patterns are shown, and as an example, pink stars designate taxa part of the same mimicry ring. The distribution of our individuals is indicated by coloured dots and coloured rings on a distribution map based on the subspecies distribution from (10)); map from USGS, Esri, TANA, DeLorme and NPS; see Fig. S6-7 for larger maps). The chromosome numbers in the right column are based on (48) or our reference genomes (asterisk).

However, note that gene flow could affect our estimates, as suggested by discordance between the BEAST2 and IQtree2 topologies (Fig. 2 vs Fig. 1 (note low concordance factors in Fig. 1)): *Mec. messenoides* is not sister to *Mec. menapis* anymore, and *Mec. nesaea* has shifted to be sister to *Mec. lysimnia* instead of *Mec. polymnia*. The relationships among lineages within *Mec. polymnia* also changed. While ILS could partially explain this, introgression could also cause it (see details below). Gene flow between sister lineages will make their apparent divergence times shorter than the initial split time, and gene flow with non-sister lineages will make them appear longer.

### Phylogeographic patterns

Through combining our sampling locations with those compiled by (10) adjusted according to our taxonomic revision (Fig. 2), we assessed the biogeographic distribution of the species. We find four main biogeographic regions (Fig. 2; Fig. S6-7), with most species restricted to one of them. Some sister-species are separated by the Andes: *Mec. messenoides* and *mazaeus* (Western Amazonia) versus their respective sisters *Mec. menapis* and *macrinus* (West of the Andes). However, note that the placement of *Mec. messenoides* differs in the BEAST2-topology and is affected by hybridisation (see next section). Within *Mec. polymnia*, Ecuadorian and Colombian individuals from opposite sides of the Andes are highly divergent, indicating little gene flow, though our study includes one putative hybrid (Fig. S8). This pattern of species separation by the Andes was not previously as apparent due to species misclassification.

Most individuals in our *Melinaea* dataset are from Western Amazonia (east of the Andes) and we find that many sister species are sympatric. Only two species in our dataset occur west of the Andes (*Mel. lilis* and *idae*) and they form a clade with a third species from east of the Andes (*Mel. isocomma*). Our dataset lacks a potential third species occurring west of the Andes, *Mel. scylax* (10), which may represent a subspecies of *Mel. lilis* (26, 57). Other potential species missing in our dataset are *Mel. mnemopsis* from Western Amazonia, *Mel. ethra* from the Atlantic forest and *Mel. mnasias* from Western Amazonia and the Atlantic Forest (10, 57).

### Signatures of rampant introgression throughout both genera

Phylogenetic discordance between phylogenies constructed for the nuclear and mitochondrial genome (Fig. 1), for different genomic regions inferred with concordance factors (Fig. 1), and with IQtree2 and BEAST2 (Fig. 1 vs Fig. 2) is indicative of a history of introgression and/or ILS in both genera. We assessed hybridisation throughout both radiations with windowed species tree inference with BPP (Fig. S5), excess allele sharing between non-sister taxa estimated from Fbranch (Fig. S9) and joint inference of species tree with gene flow using the approximate isolation with migration (AIM) model (Fig. S10). The resulting phylogenies and introgression histories are summarised in Fig. 2 (Text S3), but we stress that this is only one of multiple possible scenarios consistent with our observed patterns of excess allele sharing and gene tree discordance.

In *Mechanitis*, two lineages show uncertain placements. As mentioned, the placement of *Mec. messenoides* and *nesaea* varies depending on the methodology (IQtree2 and BEAST2) and genomic regions (Fig. 1A – box II and III). Contrary to the nuclear phylogeny inferred with BEAST2, but in line with the mitochondrial phylogeny, the species phylogeny inferred allowing for gene flow (AIM) places *messenoides* closer to the *polymnia-nesaea-lysimnia* clade rather than with *menapis-mazaeus-macrinus*. AIM indicates admixture between these clades (Fig S10A – arrows B-D), and Fbranch also confirms excess allele sharing between *messenoides* and *polymnia-nesaea-lysimnia* (Fig. S9A - #1). According to the BPP-analysis, *Mec. messenoides* groups with menapis-mazaeus-macrinus in ∼51% of the genome, and with *polymnia-lysimnia-nesaea* in ∼28% of the genome (sometimes still with *Mec. menapis*) (Fig. 5B). The branching order within the *polymnia-lysimnia-nesaea* clade is also highly variable across the genome: *polymnia* and nesaea are sister in 46.3% of the genome grouping, while 31.7% of trees group *polymnia* with *lysimnia*, and 14.7% group *nesaea* with *lysimnia* (Fig. 5C) (see also Fbranch - Fig. S9A #3). In short, these results are consistent with *Mec. messenoides* being introgressed (Fig 2A - arrows a-d) and gene flow between *polymnia-lysimnia-nesaea*.

In *Melinaea*, IQtree2 and BEAST2 produce the same topology (Fig. 1 vs Fig. 2), with *Mel. idae* sister to *Mel. lilis* and *Mel. isocomma*. However, the AIM analysis groups *Mel. idae* with the ingroup clade, although showing extensive gene flow from the lineage of *Mel. lilis* and *Mel. isocomma* (Fig. 2B - arrow a; Fig S10B – arrow A,D; see also Fbranch (Fig. S9B - #1)). To resolve the position of *Mel. idae*, as well as putative introgression between deeper branches of the phylogeny, we ran BPP focusing on the relationships between the *lilis-isocomma-idae* clade and representatives of the ingroup clade (*mneme*, *marsaeus* and *mothone*) (Fig. 5E-F). In the majority of the genome (43.4%) *lilis, idae* and *isocomma* group together (like IQTree2 and BEAST2), while in 34.5% of the genome *idae* groups with *mothone-marsaeus-mneme* (like AIM). Notably, the Z chromosome supports almost exclusively the latter relationship. In 4.3% of the genome, *isocomma* clusters with *mothone-marsaeus-mneme*. *Mel. lilis* is almost as often sister to *idae* (30.3%) as to *isocomma* (38.3%), and less commonly an outgroup to both (4.1%), indicating more recent shared ancestry between *lilis* and *idae*, which is confirmed by Fbranch excess allele sharing (Fig. S10B; Fig S9B - #2). *Mel. mothone, marsaeus* and *mneme* also vary in their respective relationships (Fig. 5E).

We further investigated the timing of divergence and introgression for the three species showing the strongest signals of introgression (*Mec. messenoides*, *Mec. nesaea* and *Mel. idae*), using a multispecies coalescent-with-introgression (MSCi) approach. In all three cases, introgression is estimated to be old (200-425 kya) (Fig. 3A-B scenario 1; Fig. 3C). For *Mec. messenoides* and *Mel. idae,* different replicate runs produce different outcomes reflecting the uncertainty in the placement of these two species in the phylogeny. Notably, some replicates indicate the origin of the introgressed lineages closely coincides with the split time from both parents, suggestive of a hybrid origin (Fig. 3A-B scenario 2; Fig. S11; Table S2).

**Figure 3.**
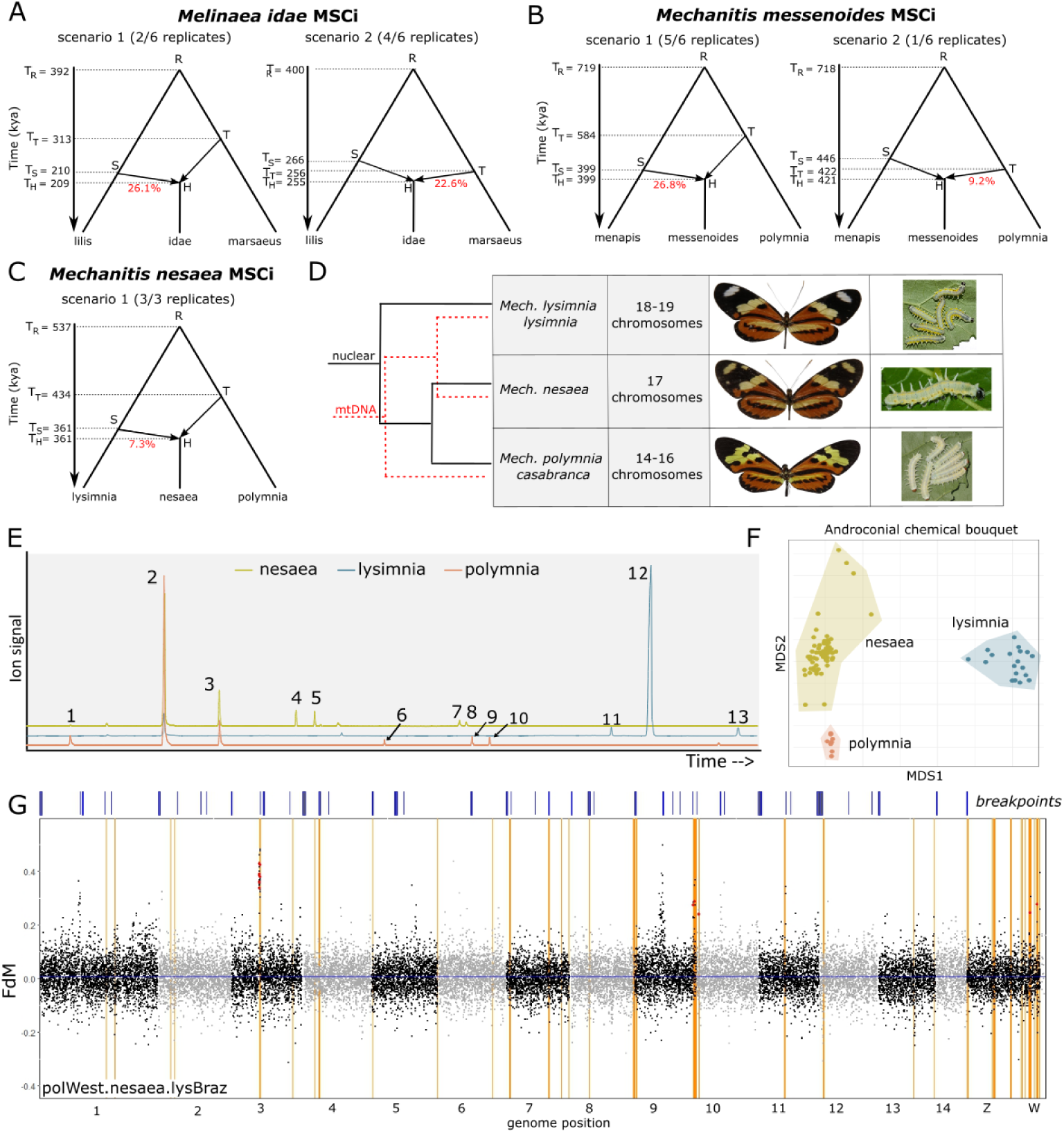
Three ancestrally introgressed species, with a focus on Mechanitis nesaea. A multispecies coalescent-with-introgression model explored the relation and timing of introgression relative to the divergence times, in **A)** Mel. lilis, Mel. idae and Mel. marsaeus; **B)** Mec. menapis, Mec. messenoides, and Mec. polymnia; and **C)** the Brazilian Mec. lysimnia lysimnia, Mec. nesaea and Mec. polymnia casabranca. **D)** A closer look into the restored species Mec. nesaea: phylogenetic relationships, chromosome numbers (48), a photo of a representative adult and an early fifth instar larva (123, 124). The photo of M. lysimnia lysimia is a courtesy of Augusto Rosa. **E)** Overlaid chromatograms of androconial extracts of representative individuals of Mec. nesaea (yellow line), Mec. l. lysimnia (blue) and Mec. polymnia casabranca (orange). Peaks: (1) 4-Hydroxy-3,5,5- trimethylcyclohex-2-enone, (2) Hydroxydanaidal, (3) Methyl hydroxydanaidoate, (4) Methyl farnesoate isomer, (5) Methyl (E,E)-farnesoate, (6) m/z 57, 43, 55, 56, 85, (7) Octadecatrienoic acid (cf.), (8) Octadecanoic acid, (9) Ethyl linolenate, (10) (E)-Phytyl acetate (11) Hexacosene, (12) Heptacosene, (13) Nonacosene (not all compounds of Table S3 are found in these three individuals). **F)** NMDS shows the androconial chemical bouquet of Mec. nesaea is clearly distinct from both putative parental lineages, most similar to Mec. polymnia. **G)** A genome scan of f_dM_ across the genome (in 20kb windows) reveals that strong signatures of introgression (f_dM_) between Mec. nesaea and Mec. lysimnia (P1=allopatric polymnia,P2=nesaea,P3=lysimnia,P4=Forbestra) overlaps with regions of high differentiation (F_ST_) between Mec. nesaea and its sister species Mec. polymnia (orange vertical lines - high F_ST_, red dots - high f_dM_ and high F_ST_). Chromosomal breakpoints between Mec. polymnia and the four other reference genomes are shown with blue bars on top.

### A focus on *Mec. nesaea*

As we propose to re-elevate *Mec. nesaea* to a species from its previous classification as a subspecies of *Mec. lysimnia*, we investigated the relationships and reproductive isolation between *Mec. nesaea*, *Mec. lysimnia* and *Mec. polymnia* in more detail. We resequenced 15 *Mec. nesaea*, 19 *Mec. l. lysimnia* and 9 *Mec. p. casabranca*, of which 24 were sampled from sympatry, and assessed the androconial compounds that might act as pheromones, potentially contributing to assortative mating. Even though *Mec. lysimnia* and *Mec. polymnia* have been observed to interbreed in nature (58) and putative hybrids have been observed (Fig. S12), we find relatively high F_ST_ throughout the genome between all three species and no evidence of ongoing gene flow based on ADMIXTURE analyses (Fig. S8; S13). Furthermore, they have been shown to differ in number of chromosomes (Fig. 3D) (48), and we find their androconial bouquets to be clearly distinct (Fig. 3E-F, Text S4; Table S3-4). All these lines of evidence suggest strong reproductive isolation between *Mec. nesaea* and both *Mec. polymnia* and *Mec. lysimnia.* Genome scans for introgression (f_dM_) between *Mec. lysimnia* and *Mec. nesaea* compared to allopatric *Mec. polymnia* show that this introgression is restricted to few genomic regions (Fig. 3G; Fig. S14-S15). The MSCi model for *Mec. nesaea* suggests *Mec. nesaea* diverged from *Mec. polymnia* prior to *Mec. lysimnia* introgression. However, *lysimnia*-introgression could have contributed key genetic variation to *Mec. nesaea* and strengthened reproductive isolation to *Mec. polymnia*. Consistent with this hypothesis, we find that introgression peaks coincide with peaks of elevated D_xy_ and F_ST_ between *Mec. nesaea* and *Mec. polymnia* and mostly show no evidence of excess allele sharing between *Mec. nesaea* and sympatric *Mec. polymnia* compared to allopatric *Mec. polymnia* (Fig. 3G; Fig. S13-S15), but the low levels of ongoing gene flow make these measures poor predictors of reproductive isolation barriers.

### Chromosomal rearrangements

To study chromosomal rearrangements, we generated haplotype-resolved genomes of five *Melinaea* and five *Mechanitis* species using PacBio and HiC-data (Fig. S16), resulting in high quality genomes with >96% BUSCO completeness and >94% of the genome assembled to chromosomes (Table S5-S6). The inferred chromosome numbers match the chromosome counts from karyotyping (48). The *Mechanitis* genomes are substantially shorter (291-320 Mb) than the *Melinaea* genomes (496-661 Mb, Table S5).

Whereas most Lepidoptera have conserved karyotypes with 31 chromosomes (59), our synteny analysis using BUSCO genes show that *Mechanitis* and *Melinaea* genomes are highly rearranged compared to the ancestral karyotype (Fig. S17-S19), corroborating earlier findings from two *Melinaea* genomes (60). The median length of conserved syntenic blocks is 32-36 genes, compared to 168 genes in the outgroup *Danaus plexippus* (Table S7; Fig. S19). The canonical Z has not undergone any fissions and has retained the longest conserved syntenic blocks (225-227 genes; Fig. 4A; Table S7; Fig. S19) in our genomes. However, both genera share a fusion of Z with parts of ancestral autosome 10 and *Mechanitis* has a further fusion with parts of ancestral autosome 6 (Table S8; Fig. S18). In addition, a fusion-fission polymorphism involving the Z and another autosome was observed between the Zs of the male *Mel. isocomma* (Table S8; Fig. S16). A W-chromosome was identified in all females as a segment of varying size depleted of BUSCO genes with moderate sequence similarity within genera, but none between (Fig. 4A). We found W-autosome fusions in four species (Table S8). *Mec. macrinus* and *mazaeus* show complex fusions between the W and multiple autosomes that are partially shared. Four species have two Z chromosomes, as by definition, the homologue of the chromosome fused to the W becomes a Z chromosome. Similarly, three species have two or three W chromosomes. Surprisingly, seven individuals were heterozygous for one or more simple autosomal rearrangement and in *Mel. ludovica* we detected a complex chain rearrangement involving four autosomes in one haplotype and three autosomes in the other haplotype (Table S8; Fig. S16). None of the simple autosomal polymorphisms are shared among taxa.

**Figure 4.**
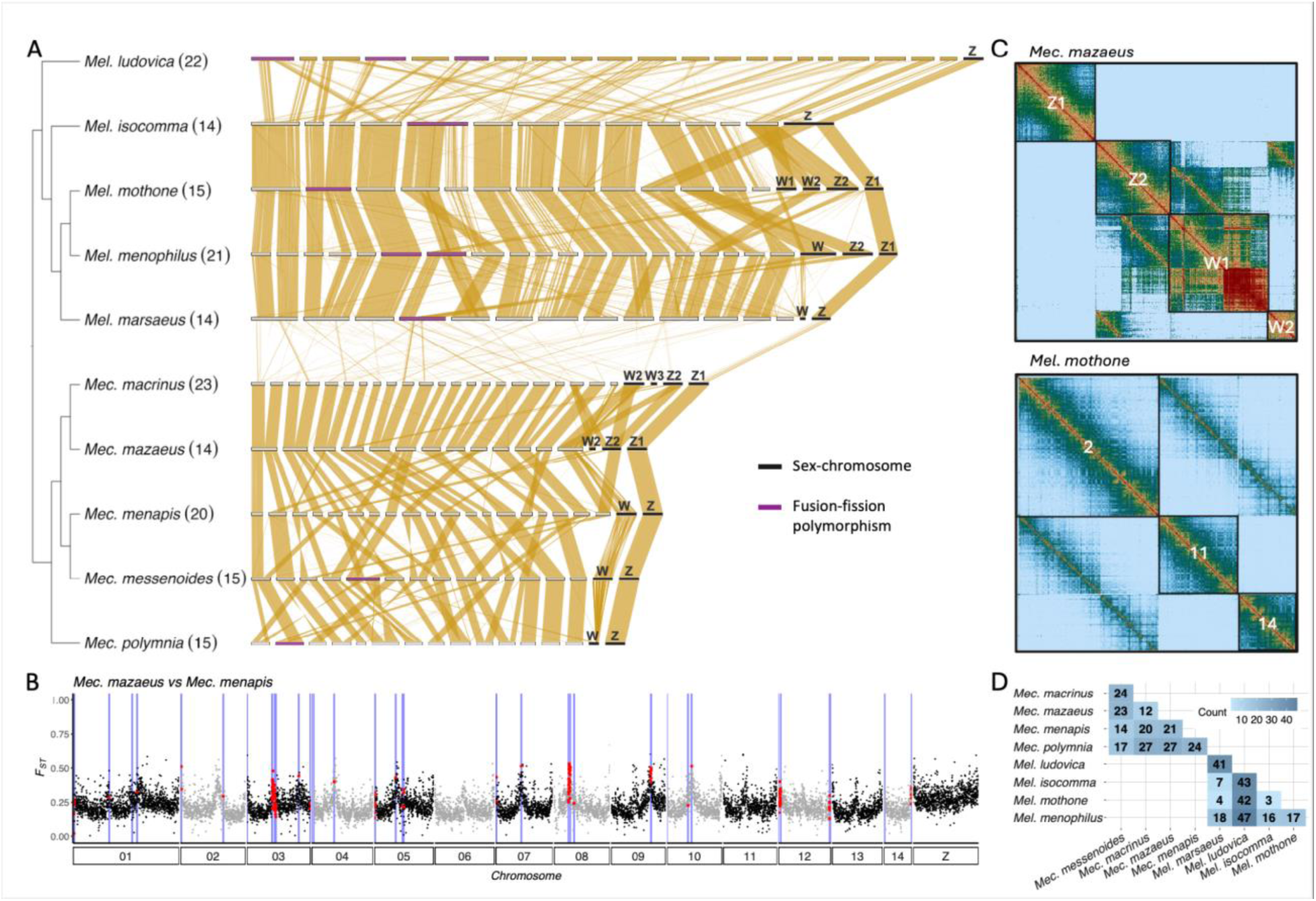
Chromosomal rearrangements. **A)** Synteny between Melinaea and Mechanitis genomes based on whole genome alignments. Horizontal bars represent individual chromosomes, with sex-chromsomes (black bar) and chromosomes involved in within-species polymorphic fusion-fissions (purple bar) highlighted. The cladogram is based on Fig. 2 and shows haploid chromosome numbers in parentheses. **B)** Example of differentiation (F_ST_) and breakpoints (blue vertical lines) between Mec. mazaeus and Mec. menapis along the genome. The red dots indicate windows coinciding with breakpoints. **C)** Examples of HiC-contact maps. Top panel: Sex-chromosomes in Mec. mazaeus. Lower panel: Autosomal fission-fusion heterozygote in Mel. mothone. **D)** Matrix displaying number of fusion-fission rearrangements between species in each genus.

While chromosome spreads had revealed species differences in chromosome counts (48), we here show that these are not due to a few fissions or fusions, but complex rearrangements. Closely related sister species show 3-47 chromosomal rearrangements (Fig. 4), which could contribute to reproductive isolation. To assess if these chromosomal rearrangements confer reproductive isolation barriers, we mapped the location of the chromosomal breakpoints between all species pairs to test for an association between breakpoints and reduced gene flow. We found increased *F_ST_* in all *Mechanitis* comparisons, and reduced diversity (π) in breakpoint regions in seven of the ten comparisons (Fig. 4B, Fig. S20, Table S9). In principle, elevated *F_ST_* could be caused by increased background selection as recombination tends to be reduced at chromosome ends and levels of ongoing gene flow are low (61, 62). However, we also find elevated absolute divergence (D_XY_) in windows coinciding with breakpoints in four of the ten comparisons, which indicates that background selection alone cannot explain the pattern (63). In *Melinaea*, we did not observe significantly elevated F_ST_ in the breakpoint regions, but we detected an increase in D_XY_ especially in the comparisons involving the more distantly related *Mel. ludovica* (Table S9).

## Discussion

Our results confirm that the two ithomiine genera *Melinaea* and *Mechanitis* represent fast and recent radiations. They diverged in the past 1-2 million years, and speciated much faster than the other well-studied Neotropical butterfly radiation of *Heliconius* (respectively 1.633 and 1.431 versus 0.324 speciation events per lineage/My (46 species in 11.8 My)) (64). Kawahara et al. (21) previously found a significant rate shift towards high diversification rates in ithomiine butterflies (clade L; 0.23 speciation events per lineage/My) and we show that among Ithomiini, *Mechanitis* and *Melinaea* have an even higher speciation rate, consistent with previous results (11). Our results shed light on potential drivers of their rapid diversification, detailed below.

Given the high number of ancient introgression events across both genera, hybridisation may have sped up speciation by boosting genetic variation as seen in other systems (e.g. 42). We identified three species showing ancient introgression that might have a hybrid origin. Further research on which genomic regions are more or less likely to introgress could inform us as to what extent introgression contributed to reproductive isolation between those taxa and their parental lineages. For instance, similar to *Heliconius* butterflies, introgression might have contributed to both mimicry ring switches and colour-based assortative mating, which generate reproductive isolation, thereby driving speciation (65).

Despite ancient hybridisation, we find little evidence for ongoing gene flow between the species, suggesting strong reproductive isolation. Some of this reproductive isolation is likely attributed to the exceptionally high rates of chromosomal rearrangements across both genera, as complex chromosomal rearrangements are expected to constitute barriers to gene flow (66, 67). While Lepidoptera chromosomes are holocentric and may be able to tolerate simple fusions and fissions during meiosis, most Lepidoptera have retained the same highly conserved karyotypes of 31 chromosomes (59, 68). The massive chromosomal rearrangements we found in ithomiini are thus unusual, but there are other lineages that also show high rates of fissions and fusions (e.g. *Leptidea* (69); and *Erebia* (70)) and there is a slight association with variation in chromosome number and speciation rates (71).

Even though chromosomal rearrangements are thought to limit gene flow, many of our genomes showed heterozygosity for chromosomal rearrangements. The heterozygote fission-fusions are unambiguous in the HiC-data, matching the intraspecific variation in chromosome counts documented previously (48) and cytogenetical evidence for pervasive polymorphism in *Mel. satevis cydon* (54). This may be akin to *Leptidea sinapis*, where, despite multiple fusion/fission polymorphisms segregating within populations (69, 72), crosses between chromosomal extremes show low survival of F2 hybrids (73). That complex chromosomal rearrangements involving multiple chromosomes likely constitute stronger barriers to gene flow compared to simple fusions and fissions has also been found in other taxa such as shrews (74) and wallabies (75) and is in line with findings of nearly sterile hybrids in a cross between two *Melinaea* sister species that differ in chromosome count (54). It is thus possible that simple fusions and fissions are not strongly selected against, but if too many sequential fusions accumulate in isolated populations, they cause reproductive isolation upon secondary contact.

Furthermore, we find many fusions between sex chromosomes and autosomes. Given the disproportionate role sex chromosomes are proposed to have in speciation (Haldane’s rule and/or large-Z effect) (76–78), they may play a strong role in reproductive isolation of ithomiini butterflies. We found two Z-autosome fusions shared among all congenerics, but no fission in the canonical Z, consistent with conserved Z chromosomes in other Lepidoptera lineages (68). The heterozygote Z-autosome fusion in *Mel. isocomma* potentially represents an ongoing fixation of a novel Z-autosome fusion, which might rapidly rise in frequency if there are advantageous loci involved (79). The simple W-autosome fusion observed in *Mel. menophilus* resemble those described in some *Heliconius* butterflies (80). The complex fusion-fissions with multiple Ws and Zs in *Mec. macrinus* and *Mec. mazaeus* are similar to *Leptidea* butterflies (81) and may constitute particularly strong barriers to gene flow (82).

Since the divergence times of *Mechanitis* and *Melinaea* species match Pleistocene climatic fluctuations (2.58 Ma-11.7 ka), and many sister species are separated by the Andes or restricted to different geographic regions corresponding to postulated Pleistocene refugia (26, 34, 37), we believe that our findings are consistent with the speciation pump hypothesis. This hypothesis postulates that periods of allopatry, followed by secondary sympatry, can facilitate diversification (37), as proposed in e.g. fishes (45, 83) and Andean plants (84). The high rates of chromosomal rearrangements in *Mechanitis* and *Melinaea* might accelerate the accumulation of chromosomal differences in isolated populations. On coming together in secondary sympatry, these rearrangements likely cause low hybrid fitness, which in turn might facilitate adaptation to different niches due to low levels of gene flow. Selection against interbreeding might also lead to reinforcement of reproductive isolation via assortative mating, e.g. based on pheromones (18, 85).

Our data suggests that hybridisation-derived genetic variation and high rates of chromosomal rearrangements both may have played key roles in the fast diversification of *Melinaea* and *Mechanitis*, and potentially contributed to adaptation and assortative mating facilitating species coexistence. Notably, while an important role of hybridisation in diversification has been found in both insular environments (e.g. silverswords (44), or cichlids (45)) and continental radiations (e.g. this study, *Heliconius* (86), or *Rhagoletis* flies (87)), rapid radiations with high rates of chromosomal rearrangements seem to be restricted to continents (e.g. this study, shrews (74), or wallabies (75)). Due to more opportunities for geographical isolation on large land masses than in insular environments, factors such as high rates of chromosomal rearrangements that speed up the allopatric accumulation of incompatibilities may thus be particularly important in continental radiations.

Our study not only sheds light on drivers of continental radiations, but also largely resolves the taxonomy of important biodiversity indicators. Hitherto, the study of these radiations has been hampered by taxonomic challenges, whereas our combination of whole-genome resequencing with vast taxonomic and geographic coverage, genome assemblies and androconial chemical analysis allowed us to resolve taxonomic issues. Our study confirms that DNA barcoding can be misleading, massively underestimating species richness, and should only be used with care to assess biodiversity. The taxonomic implications of both introgression and karyotype diversity for species delimitation and designation of conservation units are important to consider during monitoring and management of biodiversity in these vulnerable habitats.

## Materials and methods

### Collecting butterfly specimens

157 specimens of *Mechanitis*, 9 specimens of *Forbestra*, 109 specimens of *Melinaea,* 1 *Eutresis* and 1 *Olyras* were collected over the years 2000 to 2023 across Central and South America (Table S1). Adult butterflies were caught with a net, and their bodies were subsequently preserved in ethanol, DMSO or flash-frozen and stored at -70°C. Moreover, a few legs from dried museum specimens were used (Florida Natural History Museum; Natural History Museum London). Wings were photographed and stored in envelopes. The resulting dataset covers almost all species of *Melinaea* and *Mechanitis* from a wide geographical range. In addition, 65 individuals of *Mec. lysimnia nesaea* (status restored to *Mec. nesaea* in this paper; hereafter called *Mec. nesaea*), 19 *Mec. lysimnia lysimnia* and 8 *Mec. polymnia casabranca* were collected for the androconial chemical analysis (Table S3).

### DNA extractions & whole genome resequencing

DNA extractions were done with either the Qiagen MagAttract High Molecular Weight kit (Qiagen ID 67563), or the Qiagen QiaAmp DNA mini kit (51304), or a PureLink digestion and lysis step followed by a magnetic bead DNA extraction (88). The dried museum specimens were extracted using a Lysis-C buffer and a MinElute DNA extraction kit (protocol adapted from (89); Qiagen ID 28006). Library preparations were performed using homemade TN5-transposase-mediated tagmentation (protocol adapted from (90)), or following the manufacturer’s guidelines with the Illumina DNA PCR-free library prep kit and sequenced (150 bp paired-end) on Illumina NovaSeq 6000 or NovaSeq X machines at Novogene or the Wellcome Sanger Institute.

### Reference genomes

Haplotype-level chromosomally resolved reference genomes were assembled for five species of *Mechanitis* (*Mec. messenoides, menapis, mazaeus, macrinus* and *polymnia*) and *Melinaea* (*Mel. ludovica, marsaeus, mothone, isocomma* and *menophilus*). Note that earlier versions of two *Melinaea* genomes were published previously (60). In short, we combined 12-57x PacBio HiFi sequencing and 33-197x Illumina sequencing of HiC libraries (haploid coverages, Table S5) and assembled the genomes according to the Tree of Life pipelines (https://github.com/sanger-tol/genomeassembly) (Text S5).

### Whole genome mapping

To prepare the whole genome data for analysis, read quality was checked with FastQC (v0.11.9) (91). Sequences below 50 bp were discarded and adapters and PolyG-tails were trimmed with FastP (v0.23.2) (92), before they were aligned to *Melinaea marsaeus* (60) or *Mechanitis messenoides* using BWA-mem (v.0.7.17) (93). Picard was used to remove PCR duplicates (v3.0.0) (94). Samtools (v1.17) (95) and GATK3 HaplotypeCaller (v3.8.1.0) (96, 97) were used for variant calling, with a minimum base quality score of 20.

VCFtools (v0.1.16) (98) was used for filtering. Based on the distribution of sequencing depth (mean *Melinaea*: 7; *Mechanitis*: 15), all sites with a mean depth below 3 (*Melinaea*) or 5 (*Mechanitis*), and above 15 (*Melinaea*) or 30 (*Mechanitis*) were removed. Insertions, deletions, sites with >50% missing data, as well as genotypes with a depth below 2 (*Melinaea*) or 3 (*Mechanitis*) were removed. The mitochondrial DNA was filtered separately, with a maximum depth of 1700 (*Melinaea*) or 1200 (*Mechanitis*).

### Phylogenetic analyses

For each genus, we ran a Principal Component Analysis (PCA) using Plink (V1.9) to explore population structure (99) (Fig. S21). Next, we inferred a phylogenetic tree based on a filtered subset of the whole genome sequence data (also including monomorphic sites, thinned to 1 in 500 sites, with a minimum genotype quality of 10). Our filtered VCF-files were converted to phylip with a custom script which calls heterozygous sites as ‘ambiguous’ (equal likelihood for both alleles) to generate one sequence per individual (vcf2phylip.py, http://www.github.com/joanam/scripts), and subsequently, IQtree2 (v2.1.2) (100) produced phylogenetic trees with ultrafast bootstrap approximation (-B 1000; UFBoost) (101) and the GTR-model.

We inferred separate phylogenies for mitochondrial and nuclear DNA. The nuclear trees are based on 537,500 SNPs for *Mechanitis* and 784,526 SNPs for *Melinaea*. The mitochondrial phylogenies are based on the full mitochondrial genome (not thinned) including 11,818 bp for *Mechanitis* and 11,815 bp for *Melinaea.* For *Mechanitis*, we included a maximum of six individuals of the same subspecies and country in the phylogenetic analyses, thus excluding several Brazilian *Mec. polymnia*, *Mec. nesaea* and *Mec. lysimnia*. These individuals were included in hybridisation-analyses. For *Melinaea*, all individuals were used.

In addition, a phylogenetic tree calibrated with divergence times from (11) was produced using BEAST2 (102) (following https://beast2-dev.github.io/beast-docs/beast2/DivergenceDating/DivergenceDatingTutorial.html) for the same dataset as the nuclear phylogenetic tree, but thinned further to 1 in 5000 sites and with only one individual per lineage. Divergence times between *Mec. mazaeus* and *Mec. macrinus* (0.39 MYA) and *Mel. isocomma* and *Mel. lilis* (0.65 MY) were used for calibration. We used the HKY gamma-4 site model with a strict molecular clock and the Calibrated Yule model. The model was run for 15.000.000 chains, stored every 5000 trees.

Phylogenies were visualised using the packages ‘ape’ (v5.7-1) and ‘phytools’ (v2.1-1) in R (Paradis and Schliep 2019; Revell 2024) and FigTree (v1.1.4) (http://tree.bio.ed.ac.uk/software/figtree/). To calculate a constant speciation rate (*λ*) we assumed a pure birth model, with n as the number of species and T the time from root to tip: *n* = *e*^*λ*∗*T*^ which gives *λ* = *ln*(*n*)/*T*.

### Distribution maps

The coordinates of our individuals and the individuals from (10) were plotted using the libraries ‘sf’, ‘tmap’, ‘tmaptools’ and ‘mapview’ in R (103–105). We classified the subspecies of (10) into species following our taxonomic revision. Our individuals were plotted as dots on top of the overall distribution combining data with (10). The basemap ‘Esri.WorldTerrain’ was used, provided through ‘Leaflet’ (ESRI, ARCGIS; https://github.com/leaflet-extras/leaflet-providers); Fig. S6-7 has maps including the attribution.

### Window trees

IQtree2 was used to produce phylogenetic trees in windows across the genome (one 20kb window every 200kb) and to calculate gene concordance factors between these window-trees and the whole-genome phylogenetic tree (106).

### DSuite analysis

We explored excess allele sharing between species or divergent subspecies using DSuite (107). We filtered our genomic dataset to be 1 in 100 sites, only including variable sites. Dtrios calculated D and f4-ratio statistics for all trios and Fbranch then summarised them as f-branch statistics using a species tree based on the nuclear phylogeny of Fig. 2 (Table S1). The results of this analysis were visualised with a python script provided by DSuite.

### AIM with StarBEAST2

We followed the Approximate Isolation with Migration (AIM) in StarBEAST2 tutorial to obtain a phylogenetic tree with admixture arrows (102, 108). For each genus, we picked one individual per species (highest depth), and randomly extracted 150 (*Mechanitis*) or 200 (*Melinaea*) 800-1000bp windows across the genome. We used the HKY gamma-4 site model with a strict molecular clock and the Yule model. The migration rate was set to 25 with an initial value of 0.1, and we ran the model for 100.000.000 chains, storing every 5000 trees. We updated the parameters according to the suggestions in the output of the first run and re-ran the model to improve the results.

### Species-tree inference

Phylogenetic relationships across the genome between species of *Melinaea* and *Mechanitis* were inferred using the multispecies coalescent (MSC) approach implemented in BPP v.4.6.2 (109), while accounting for incomplete lineage sorting. For each species, only the individual with the highest coverage was included. To minimise the effect of linked selection, loci were required to be at least 2 kb from annotated exons. Because the analysis assumes no intra-locus recombination and independence between loci, we selected loci of 300 bp and at least 2 kb from neighbouring loci. Sequence alignments were produced for all loci, masking repetitive elements annotated in the reference genome using RepeatMasker v4.1.5 (http://repeatmasker.org/). For each locus, individuals with more than 80% missing genotype calls were excluded from the alignment and only loci with all individuals passing filters were considered. Furthermore, sites with missing genotype calls were removed and loci with less than 30 bp passing filters were excluded. Heterozygous sites were assigned IUPAC codes. Loci were grouped into blocks of 100 loci. Species-tree estimation was then performed in BPP v.4.6.2 using the A01 analysis (species-tree inference assuming no gene flow). Inverse gamma priors (invGs) were applied both to the root age (τ0) and to effective population sizes (θ) – invG(3, 0.06) and invG(3, 0.04), respectively. Parameters were scaled assuming a mutation rate of 2.9 × 10−9 substitutions per site per generation and 4 generations per year. The MCMC was run for 1,000,000 iterations after 50,000 iterations of burn-in, sampling every 10 iterations.

### Demographic modelling

For the three highly introgressed species *Mel. idae, Mec. messenoides* and *Mec. nesaea, we ran* a multispecies-coalescent-with-introgression (MSCi) model implemented in BPP v.4.6. 2 (109) to better estimate their position in the phylogeny and divergence timing while accounting for admixture. For *Mel. idae* we considered *Mel. lilis* and *Mel. marsaeus* as sister/donor species, while for *Mec. messenoides*, *Mec. menapis* and *Mec. polymnia* (West) were chosen. For *Mec. nesaea*, we used one individual of each of the Brazilian populations (*Mec. nesaea, Mec. lysimnia lysimnia*, *Mec. polymnia casabranca*). Loci were selected randomly from autosomes, and required to be at least 2 kb from annotated exons and 10 kb from the nearest locus, and a maximum size of 2 kb. For each locus, individuals with more than 50% missing data and sites containing missing genotypes or overlapping annotated repetitive elements were removed. Only loci at least 800 bp long after filters and without missing individuals were considered. Heterozygous sites were assigned IUPAC codes. Demographic parameters were estimated using a fixed species tree with introgression events (Fig. S11). An inverse gamma prior (invG) was applied for all population size parameters θ (α=3; β=0.04) and root age parameter τ (α=3; β=0.06). A beta prior was applied to the introgression probability (ϕ) (α=1; β=1). Three replicate MCMC runs of 1,000,000 iterations, after a burn-in period of 50,000 iterations, sampling every 50 iterations were performed for each dataset. Divergence time was calculated based on 4 generations per year, and a mutation rate of 2.90E-09 as in *Heliconius* (43) (T=τ/mutation rate/generations/million years).

### Genome scans and ADMIXTURE

F_ST_, D_XY_ and pi (π) were calculated in all species pairs in 20kb windows (including monomorphic sites, with a minimum of 10,000 sites per window) across the genome with the scripts from Simon Martin (https://github.com/simonhmartin/genomics_general). We also ran ADMIXTURE in subsets of *Mechanitis* and *Melinaea* to see clustering between individuals with k ranging from 2 to 9 (110). For this dataset, we used the same filter settings as for the Fbranch analysis (see above).

### Introgression (f_dM_) scans

To examine gene flow across the genome of *Mec. nesaea*, we computed f_dM_ with the script by Simon Martin (ABBABABAwindows.py, https://github.com/simonhmartin/genomics_general) (111) with various populations for P1,2,3,4. If for example P1=*Mec. polymnia casabranca* (Brazil), P2=*Mec. nesaea*, P3=*Mec. lysimnia* Brazil, and P4=*Forbestra*, high f_dM_ values indicate introgression between *Mec. lysimnia* and *Mec. nesaea*. We also tested other outgroups (*Mec. messenoides*) and with five single *Mec. nesaea* individuals, to see if the signal was consistent in different individuals (Fig. S14).

### Androconial Chemistry

We obtained samples of the androconial secretions from 92 males (Text S5). To exclude non-androconial compounds from our analyses, for each population the same extraction procedure was adopted with wings of conspecific females and two non-androconial wing areas of males. Eight solvent negative controls were also taken for each sampling event.

The peak areas of each chromatogram were integrated with MSD ChemStation E.02.01.1177 (Agilent Technologies, USA) to obtain the total ion current signals. A series of linear alkanes (C7–C40) was used to determine the linear retention indices (RI) of each compound (112). Compounds were identified by comparing their mass spectra and retention indices with those of reference samples available from personal and commercial mass spectral libraries (FFNSC 2, MassFinder 4, NIST17, MACE v.5.1 (113), and Wiley Registry™ 9th ed.). The peaks exclusive to the androconial samples were used to determine the relative percentages of each compound per sample.

Dihydropyrrolizines, such as hydroxydanaidal or methyl hydroxydanaidoate, are typically accompanied by smaller peaks formed by degradation during GC/MS analyses and/or storage (114). These degradation peaks were excluded from the generated ion chromatograms and statistical analysis. To avoid non-evident contaminants, we only considered compounds present in more than three individuals for any given taxon.

The data were plotted in R (115) using the vegan package (https://doi.org/10.32614/CRAN.package.vegan) for nonmetric multidimensional scaling with ‘monoMDS’, specifying a global model, square root transformation and Wisconsin double standardisation (autotransform=TRUE).

### Chromosome rearrangements

Synteny and breakpoint analysis was performed using single copy orthologous genes identified with BUSCO version 5.7.1 with the lineage database lepidoptera_odb10 and otherwise default options (116). To compare large scale rearrangements in the *Mechanitis* and *Melinaea* versus the butterfly ancestral linkage groups (Merian elements), we used two outgroup genomes, *Melitaea cinxia* (117) and *Danaus plexippus* (118). The output from BUSCO was filtered with a custom script to contain only complete genes located in chromosome-sized scaffolds and excluding W. Only single copy genes were included with the exception of genes on the neo-Z2, where we also included genes classified as duplicated. The sex linking of Z2 appears to be recent and most of the genes had high similarity to the genes on the corresponding W’s and were therefore classified as duplicated by BUSCO. Genes that were actually duplicated (occurring more than once or present on other chromosomes than W) on the Z2 were removed. We determined syntenic blocks, excluding single gene translocations, by comparing the position of the BUSCO-genes in each genome against the Merian elements (*Melitaea cinxia*), and visualised the syntenic relationship with the R-package gggenomes (119). Minimum number of fusions were determined by the number of different Merian elements located in every query chromosome and fissions were determined as the number of query chromosomes containing parts of each Merian element for each species. The breakpoints between all species within *Mechanitis* and *Melinaea* was determined by the same principle as above using an all against all approach for each genome to compare the BUSCO-gene positions. Synteny analysis of the sex chromosomes and between haplotypes were performed with whole genome alignment using minimap2/2.27 with default settings and -x asm10 (1% sequence divergence) (120) and visualised after removing short alignments (<100kb for multispecies alignment, <500kb for haplotype alignment) using a modified version of the R-package Farre-lab/syntenyPlotteR (121).

To investigate the association between chromosomal rearrangements and species divergence we mapped the location of the breakpoints between each comparison to the reference genomes *Mec. messenoides* and *Mel. marsaeus*. Divergence and diversity was estimated in 20 kb windows along the genome (detailed above). To determine whether the observed statistics in the breakpoint regions were different from the random expectation of regions of the same size and number across the genome, we used permTest with the randomise function option ‘randomizeRegions’ and evaluate function option ‘meanInRegions’ for 50,000 permutations, as implemented in the R-package regioneR (122). The analyses above were performed in R v4.4.0 (115). For conservative interpretation multiple correction was performed with Bonferroni adjustment (ɑ/n), where ɑ = 0.05 and n = 10 for the comparisons within each genus, resulting in values considered significantly different if their p-value from the randomisation test was less than 0.005.

## Supporting information

All supplemental figures, tables and text

## Acknowledgements

We thank Ismael Aldas and Raúl Aldaz for assistance with catching butterflies in Ecuador, Mario Tuanama and Ronald Mori-Pezo for help with rearing and collecting butterflies in Peru, Augusto Rosa for sampling support in São Paulo, Erika de Castro for sampling in Brazil, Mathieu Joron for sampling in French Guiana and John R. MacDonald for sampling in Panama. We thank the Peruvian, Ecuadorian and Brazilian authorities as well as the Museo de Historia Natural in Peru and the Museo Ecuatoriano de Ciencias Naturales in Ecuador for their support. Many thanks to Dr Blanca Huertas and Robyn Crowther from the Natural History Museum in London (NHMUK) for providing butterfly legs of *Mel. isocomma* and *Mel. mothone* and to Dr Petra Korlevic for assistance with extracting DNA from museum specimens. We are grateful to the editor and reviewers for their helpful comments that have greatly improved our manuscript.

## Data, materials and software availability

All code and tables underlying the figures are found in our GitHub repository: https://github.com/rapidspeciation/mechanitis_melinaea.

