## Supplementary material for "Genomics of Neotropical biodiversity indicators: two butterfly radiations with rampant chromosomal rearrangements and hybridisation": All supplemental figures, tables and text

PNAS

**This PDF file includes:**

Supporting text S1 to S5
Figures S1 to S21
Tables S1 to S9
SI References

**Supporting Information Text**

**Supporting Text 1 – Taxonomic revision**

For decades, taxonomists have acknowledged that species in *Melinaea* and *Mechanitis* are notoriously difficult to classify (1–3). For instance, (3) recognized only five species of *Mechanitis*, whereas (4) split them into 16 species in Colombia alone. In order to study diversification and speciation, we need to know how populations relate to each other, and what the species and subspecies are.

In *Mechanitis*, our data suggests two clades should be reclassified. The nuclear genome phylogeny of *Mechanitis* shows that the subspecies *Mec. lysimnia macrinus* (Panama and western Colombia) and *Mec. lysimnia utemaia* (Panama) form a clade that is phylogenetically distinct from the rest of *Mec. lysimnia* (Fig. 1A - box I; Fig. S1). We recognize them as a distinct species named *Mec. macrinus* **status restored** (Hewitson 1860). Furthermore, the Brazilian subspecies *Mec. lysimnia nesaea* does not cluster with the remaining South American *Mec. lysimnia* in the nuclear phylogeny (Fig. 1A - box III; Fig. S1), but is sister to *Mec. polymnia*. *Mec. lysimnia nesaea* is sympatric with *Mec. polymnia casabranca* and we therefore treat it as a distinct species, *Mechanitis nesaea* **status restored** (Hübner, 1820).

Two species show deeply divergent lineages that could potentially merit species classification, but in the absence of sequenced specimens from zones of sympatry, and given their monophyly, we currently refrain from a reclassification. There is deep divergence between Brazilian *Mec. lysimnia lysimnia* and the east Ecuadorian *Mec. lysimnia roqueensis* (Fig. 1; Fig. S1) and between the Panamanian *Mec. menapis saturata*, *Mec. menapis mantineus* from Ecuador, and a third clade with several west Colombian subspecies (*Mec. menapis occasiva* and *Mec. menapis menapis*). The overall placement of *Mechanitis menapis* in our phylogenetic tree does not match the placement of *Mechanitis menapis* in (5), where *Mech. menapis* is the most basal group. In our phylogenetic tree, it is the sister-species of *Mech. messenoides*.

Within *Mechanitis polymnia*, most individuals cluster by region (Fig. 1 and 2). There are three mitochondrial clades, which don't match the nuclear tree completely. The three *Mechanitis messenoides* subspecies (Peruvian *phasianita*, Colombian and Ecuadorian *messenoides* and *deceptus*) form a mixed monophyletic clade together; they do not cluster by country or colour pattern. This is also seen for *Mechanitis mazaeus.*

In *Melinaea*, one clade has to be reclassified. The *Mel. satevis* subspecies are not monophyletic (Fig. 1B - box V; Fig. S3). *Mel. satevis maeonis* clusters with *Mel. marsaeus*, and the other *Mel. satevis* subspecies (*Mel. s. maelus*, *Mel. s. cydon, Mel. s. lamasi*) cluster with the former *satevis* subspecies that was recently reclassified as its own species, *Mel. tarapotensis* (6). We propose to restore *Mel. satevis maeonis* as a distinct species *Melinaea maeonis* **status restored** (Hewitson 1869).

*Mel. menophilus* forms four clades (Fig. 1 - box IV; Fig. S3), the first consisting of the French Guianan *Mel. men. mediatrix,* the second consisting of the Colombian *Mel. men. ernestoi/menophilus* and Ecuadorian *Mel. men. zaneka*, a third with the Ecuadorian *Mel. men. menophilus* and another containing the Peruvian *Mel. men. 'ssp1' and Mel. men. hicetas.* However, since these subspecies do not occur sympatrically, we cannot assess the levels of reproductive isolation, and we retain them all as subspecies.

**Supporting Text 2 – Annotated synonymic list of *Mechanitis* and *Melinaea***

Genus ***Mechanitis*** Fabricius

***Mechanitis*** Fabricius, 1807: 284. Type-species, *Papilio polymnia* Linnaeus (Scudder, 1875: 212).

*Nerëis* [sic] Hübner, [1806]a: [1]. Type-species, *Papilio polymnia* Linnaeus (mon.). Invalid (ICZN, Opinion 97).

*Nerëis* [sic] Hübner, [1806]b: pls. [1-2], [5-8], [12], [15]. Type-species, *Papilio polymnia* Linnaeus (Scudder, 1875: 227). Junior homonym of *Nereis* Linnaeus, 1758.

*Hymenitis* [Illiger], 1807: 1180. Type-species, *Papilio polymnia* Linnaeus (Hemming, 1934: 28). Suppressed (ICZN, Opinion 985).

*Epimetes* Billberg, 1820: 77. Type-species, *Papilio polymnia* Linnaeus (Hemming, 1933: 199). Junior objective synonym.

***lysimnia elisa*** (Guérin), [1844]: 472 (*Heliconia élisa* [sic]). TL: Bolivia, [Santa Cruz, Santa Cruz de la Sierra]. D: Bolivia, NW Argentina, S Brazil. Lectotype ♂, MNHN (Fox, 1967: 123). **reinst. stat.** [raised to species by Le Crom & Winhard (2024: 28) on the basis of "sus *habitus* diferentes", i.e. their different wing patterns, the molecular data confirm that SE Brazil and west Amazonian populations form a clade that we believe is best treated as a species]

*elisa connectens* Talbot, 1928: 412, pl. 14, fig. 8. TL: Brazil, Mato Grosso, Rio Serragem, 1500'. Holotype ♂, NHMUK.

***lysimnia lysimnia*** (Fabricius), 1793: 161 (*Papilio*). TL: Unknown. D: SE Brazil, Paraguay, Uruguay, NE Argentina. Syntypes (♀), CCWM? Fig.- Jones, 2: pl. 8 (see Vane-Wright, 2021).

*lysimene* (Godart), 1819: 218 (*Heliconia*). *Nomen novum*.

*aurea* (Moreira), 1881: 2, pl. 1, figs. A-E (*Heliconia*). TL: "Brazil". Lectotype ♀, MNRJ (lost) (Lamas, MS).

*lysimnia* ab. *albescens* Haensch, 1905: 148. TL: Brazil, [Minas Gerais, Fazenda] Monte Cristo. “Lectotype” ♂, MfN (Lamas, 1994: 273). Unavailable name.

*castalia* (Larrañaga), 1923: 417 (*Papilio*). TL: "Uruguay". Syntypes, lost. Junior primary homonym of *Papilio castalia* Fabricius, 1793.

***lysimnia menecles*** Hewitson, 1860: [30], pl. [15], fig. 13 (*Mechanitis m.*). TL: "Amazon". D: E Colombia, E Ecuador, E Perú, N Bolivia, W Brazil. Syntypes (♀), NHMUK. **reinst. stat.** [treated as subspecies of *M. elisa* by Le Crom & Winhard (2024)]

*elisa acreana* d'Almeida, 1950: 394, figs. 2-3. TL: Brazil, Acre, Xapuri. Holotype ♂, DZUP.

*elisa roqueensis* Bryk, 1953: 30. TL: Perú, [San Martín], Roque]. Holotype ♂, RMS. **n. syn.** [treated as a valid subspecies in Lamas (2004)]

***lysimnia ocona*** H. Druce, 1876: 208 (*Mechanitis o.*). TL: Perú, [Cuzco], Valle del Río Santa Ana, Huiro. D: E Perú. Holotype ♂, NHMUK. **reinst. stat.** [treated as subspecies of *M. elisa* by Le Crom & Winhard (2024)]

*vilcanota* Röber, 1904: 105. TL: Perú, Cuzco. Lectotype ♀, SMTD (Lamas, 1992: 176).

***lysimnia tapajona*** Freitas & Mota, *in* Mota, Rosa, Vasconcellos, Willmott & Freitas, 2022: 48, figs. 1-3. TL: Brazil, Pará, Novo Progresso, Alvorada da Amazônia, Côrrego rio Quico/Arco-íris, -7.281835°, -55.311290°. D: Brazil. Holotype ♂, ZUEC. [relationships with respect to *M. nesaea* **rev. stat.** and *M. l. lysimnia* need clarifying]

***macrinus bipuncta*** Forbes, 1948: 19, fig. 7 (*Mechanitis mazaeus b.*). TL: Venezuela, [Bolívar], upper Caroní River, Surukum Basin. D: S Venezuela, Guyana, N Brazil. Holotype ♀, CUIC. **rev. stat.** [treated as a subspecies of *M. lysimnia* in Lamas (2004)]

***macrinus labotas*** Distant, 1876: xii (*Mechanitis l.*). TL: "Costa Rica". D: Pacific Costa Rica, Pacific Panamá. Syntypes (♀), NHMUK. **rev. stat.** [treated as a subspecies of *M. lysimnia* in Lamas (2004)]

***macrinus limnaea*** Forbes, 1930: 317 (*Mechanitis l.*). TL: French Guiana, [Guyane], St. Laurent. D: Guyana, Suriname, French Guiana, N Brazil. Holotype ♂, CUIC. **rev. stat.** [treated as a subspecies of *M. lysimnia* in Lamas (2004)]

*mantineus* var. Forbes, 1924: 146, 156.

*mantineus* f. *forbesi* Bryk, 1937: 491, 641. TL: French Guiana, [Guyane, St. Jean]. Holotype ♀, USNM.

***macrinus macrinus*** Hewitson, 1860: [29], pl. [15], fig. 11. TL: "New Granada". D: E Panamá, Colombia, W Ecuador, NW Perú. Syntypes (♂), NHMUK. [treated as a species by Le Crom & Winhard (2024)]

*numerianus* C. Felder & R. Felder, 1865: 368, pl. 45, fig. 9. TL: Colombia, "Bogotá" (error). Syntypes (♀), NHMUK.

*macrinus blissi* Fox, 1942: 26, figs. 7-8. TL: Panamá, [Canal Zone], Fort Kobbe. Holotype ♂, RPM. [*m. macrinus* x *m. utemaia*].

*macrinus pacificus* Le Crom & Winhard, 2024: 5, 29, pl. 29. TL: [Colombia], Antioquia, Turbo, Río León, Urabá. 50m. Holotype ♂, IAVH. **n. syn.** [described by Le Crom & Winhard (2024) based on supposedly reduced yellow DFW postdiscal markings in comparison with typical *M. macrinus macrinus*, but the ST of *macrinus* (Warren et al., 2024) shows even more reduced markings than the HT of *pacificus*]

*aurantiacus* Winhard & Le Crom, *in* Le Crom & Winhard, 2024: 5, 29, pl. 30. TL: [Colombia], Valle [del Cauca], El Salto, 9km W Cisneros, 300m. Holotype ♂, ZSM. **n. syn.** [described by Le Crom & Winhard (2024) as a species in comparison with *M. macrinus macrinus* based on complete orange DHW submarginal band and black at the base of DFW cell Cu2-Cu1; the extent of DHW orange is variable throughout the range of *M. m. macrinus*, and a black spot at the base of DFW cell Cu2-Cu1 occurs occasionally in specimens in Panama and western Ecuador; in the absence of other morphological or genetic evidence, we consider the type specimen to represent a form of *M. m. macrinus*]

*aurantiacus magdalenae* Le Crom & Winhard, 2024: 5, 30, pl. 30. TL: [Colombia], Huila, Garzón, Vereda El Espinal, Reserva Privada Taky-Huaylla, 1000m. Holotype ♂, IAVH. **n. syn.** [described by Le Crom & Winhard (2024) as a subspecies in comparison with '*M. aurantiacus*', the wing pattern is typical of *M. macrinus macrinus* specimens with more extensive DHW orange]

***macrinus solaria*** Forbes, 1948: 15, fig. 5 (*Mechanitis polymnia s*.). TL: Venezuela, Sucre, El Chorro. D: NE Venezuela, Trinidad. Holotype ♂, CUIC. **rev. stat.** [treated as a subspecies of *M. lysimnia* in Lamas (2004)]

***macrinus utemaia*** Reakirt, 1866: 241 (*Mechanitis u.*). TL: "Honduras". D: S México to W Panamá, N Colombia, N Venezuela. Holotype ♀, FMNH. **rev. stat.** [treated as a species distinct from *M. lysimnia* by Le Crom & Winhard (2024) on the basis of the "*habitus* diferentes" (different appearance), the molecular data in the present study suggest this taxon is best treated as a subspecies of *M. macrinus*]

*doryssus* f. *utemaia* ab. *extrema* Hoffmann, 1940: 636, fig. 1. TL: México, Chiapas, Mapastepec. “Holotype” ♂, AMNH. Unavailable name.

*utemaia guajira* Le Crom & Winhard, 2024: 6, 33, pl. 34. TL: [Colombia], Guajira, Río Palomino, 50m. Holotype ♂, ZSM. **n. syn.** [described by Le Crom & Winhard (2024) based on supposedly more reddish DFW yellow markings and broader DFW yellowish subapical marking, in comparison with *M. m. utemaia*; these supposed characters lie within the typical range of variation of *M. m. utemaia*]

***mazaeus beebei*** Forbes, 1948: 18, fig. 6. TL: Venezuela, Monagas, Caripito. D: E Venezuela, Guyana. Holotype ♂, CUIC.

***mazaeus lanei*** Fox, 1967: 83, figs. 63, 69B, 71, 80A-B (*Mechanitis l.*). TL: Brazil, Pará, Igarapé Açú. D: Brazil (E Pará). Holotype ♂, NHMUK.

***mazaeus mazaeus*** Hewitson, 1860: [28], pl. [14], fig. 8. TL: "Amazon". D: E Colombia, E Ecuador, E Perú, W Brazil. Lectotype ♀, NHMUK (Lamas, 1994: 284).

*fallax* Butler, 1873: 154. TL: Colombia, "Bogotá" (error). Lectotype ♀, NHMUK (Lamas, 1999: 133). **rev. stat.** [treated as a species by Le Crom & Winhard (2024) on basis of its sympatry with *M. mazaeus*, but without discussion of diagnostic characters, the LT of *fallax* (see Warren et al. 2024) is a common phenotype found throughout the western Amazon]

*polymnia* var. *plagifera* Staudinger, 1885: 62. TL: Perú, [Loreto], Yurimaguas. Lectotype ♂, MfN (Lamas, MS).

*mazaeus* ab. *lucifera* Haensch, 1905: 146, pl. 4, fig. 1. TL: Perú, [Loreto], Yurimaguas. “Lectotype” ♀, MfN (Lamas, 1994: 284). Unavailable name.

*mazaeus* ab. *phasianita* Haensch, 1905: 146. TL: [Perú, Loreto], Yurimaguas. “Lectotype” ♂, MfN (Lamas, 1994: 290). Unavailable name.

*mazaeus* ab. *nigroapicalis* Haensch, 1905: 146. TL: Brazil, [Amazonas], São Paulo de Olivença. “Lectotype” ♂, MfN (Lamas, 1994: 287). Unavailable name.

*visenda elevata* Riley, 1919: 182. TL: Brazil, [Amazonas], Rio Purús, Aliança, Canutama; Tefé; Fonteboa. Syntypes (♂♀), NHMUK. **rev. stat.** [treated as a subspecies of '*M. fallax*' by Le Crom & Winhard (2024), the ST of *elevata* (see Warren et al. 2024) is a common phenotype found throughout the western Amazon]

*lorigae* Fernández, v.1928: 209, fig. 6. TL: Perú, [Loreto], Iquitos. Lectotype ♀. MNCN (Lamas, 1999: 133, fig. 6).

*egaensis septentrionalis* Apolinar, xi.1928: 180. TL: Colombia, Boyacá, Garagoa, Valle de Tensa. Holotype, lost (Lamas, 1979: 67).

*mazaeus williamsi* Fox, 1941: 6, pl. 2, fig. 16. TL: Perú, [San Martín], Achinamiza. Holotype ♂, RPM.

*fallax pothetoides* d'Almeida, 1951: 13, pl. 5, fig. 2. TL: Brazil, [Rondônia], Rio Verde. Holotype ♀, MNRJ (lost). **n. syn.** [Lamas (2004) treated *pothetoides* as a valid subspecies, but the wing pattern of the HT occurs commonly throughout the western Amazon, so we here treat it as a synonym of *M. mazaeus mazaeus*]

*foxi* d'Almeida, 1951: 14, pl. 1, figs. 2-3. TL: Brazil, Acre, alto Rio Juruá. Holotype ♂, MNRJ (lost).

*messenoides dissimilis* Le Crom & Winhard, 2024: 6, 35, pl. 38. TL: [Colombia], Vaupés, Taraira, Estación Caparú. Holotype ♂, IAVH. **n. syn.** [the protruding edge of the DFW yellow postdiscal band in cell M2-M1, relatively broad forewings, and white VHW marginal spots, suggest that this is a form of *M. mazaeus mazaeus* (see Hill et al. 2012)]

*deceptus dicolor* Le Crom & Winhard, 2024: 6, 36, pl. 39. TL: [Colombia], Guaviare, San José del Guaviare. Holotype ♂, IAVH. **n. syn.** [the protruding edge of the DFW orange postdiscal band in cell M2-M1, relatively broad forewings, and white VHW marginal spots, suggest that this is a form of *M. mazaeus mazaeus* (see Hill et al. 2012)]

*mazaeus metaensis* Le Crom & Winhard, 2024: 6, 36, pl. 39. TL: [Colombia, Meta], Monterredondo, 1600m [error ?]. Holotype ♂, ICNB. n**. syn.** [described by Le Crom & Winhard (2024) from two specimens, based on supposedly less extensive black markings on the DFW in comparison with typical *M. m. mazaeus*; these differences are well within the range of wing pattern variation observed in *M. mazaeus* (see Hill et al. 2012)]

***mazaeus neukircheni*** Neild, 2008: 109, 234, figs. (*Mechanitis lysimnia n.*) TL: Venezuela, Táchira, Parque Nacional El Tamá, Río Frío, 600m. D: NE Colombia, SW Venezuela. Holotype ♂, MIZA. **rev. stat.** [Neild (2008) described this taxon as a subspecies of *M. lysimnia*, due to its similarity to *M. macrinus utemaia* (then a subspecies of *M. lysimnia*), but discussed the possibility that it might represent *M. mazaeus*, whereas Le Crom & Winhard (2024) treated it as subspecies of '*M. dorissides*' (=*M. polymnia*); our molecular data support Neild's (2008) supposition of a close relationship between *M. macrinus utemaia* and *M. mazaeus mazaeus*, and we here treat *neukircheni* as a subspecies of *M. mazaeus* because of its geographic proximity to *M. mazaeus mazaeus*]

***mazaeus pannifera*** Butler, 1877: 150 (*Mechanitis p.*) TL: Brazil, [Amazonas], Ega (= Tefé); [Pará], Óbidos. D: ?E Colombia, SE Venezuela, Guyana, Suriname, French Guiana, N Brazil. Syntypes (♂), NHMUK. **reinst. stat.** [treated as a species by Le Crom & Winhard (2024) on basis of its yellow rather than orange DFW subapical band, a highly variable character (in color, size, presence or absence) as supported by lack of correlation of DNA sequences with these forms; *pannifera* represents a relatively stable phenotype in the northern Amazon and Guianas, albeit one that also occurs infrequently in the western Amazon, and we retain it as a distinct subspecies for the present]

***mazaeus visenda*** Butler, 1877: 150 (*Mechanitis v.*). TL: Brazil, [Pará, Rio] Tapajós, [Trovador]. D: Brazil (Pará). Syntypes (♂), NHMUK.

***menapis caribensis*** Fox, 1967: 114, figs. 69H, 104, 106. TL: Venezuela, Distrito Federal, Puerto La Cruz. D: N Colombia, N Venezuela, ?Curaçao, Trinidad. Holotype ♂, CMNH.

***menapis dariensis*** Brown, 1977: 187, fig. 80. TL: Panamá, Darién, near El Real, base of Cerro Pirre, near sea level. D: E Panamá, NW Colombia. Holotype ♂, AMNH.

*macrinus extremus* Le Crom, *in* Le Crom & Winhard, 2024: 5, 29, pl. 30. TL: [Colombia], Antioquia, San Jerónimo, 700m. Holotype ♂, IAVH. **n. syn.**

*doryssus subobscurus* Winhard & Le Crom, *in* Le Crom & Winhard, 2024: 5, 30, pl. 31. TL: [Colombia, Risaralda], 22km W Pueblo Rico, 700m. Holotype ♂, ZSM. **n. syn.**

***menapis doryssus*** Bates, 1864: 33 (*Mechanitis d.*). TL: Guatemala, [Alta Verapaz, San Gerónimo]. D: S México to W Panamá. Lectotype ♂, NHMUK (Lamas, 1988: 49, fig. 1).

*doryssus* f. *saturata* ab. *escalantei* Hoffmann, 1940: 636, fig. 2. TL: México, Guerrero, Acahuizotla. “Holotype” ♀, AMNH. Unavailable name.

*saturata* Godman, *in* Godman & Salvin, 1901: 642 (*Mechanitis s*.). TL: Costa Rica, [Cartago], Caché. Lectotype ♀, NHMUK (Lamas, 1988: 49, fig. 2). **n. syn.** [Lamas (1988) clarified the taxonomic identities of *doryssus* and *saturata*, and treated them as valid subspecies of *M. menapis*, a classification followed by Lamas (2004). Earlier authors, e.g. Fox (1967), regarded all individuals from Mexico to Panamá as representing the same taxon, although treating *doryssus* as a subspecies of *M. polymnia*; we see insufficient geographic variation to retain the name]

***menapis mantineus*** Hewitson, 1869: 12 (*Mechanitis mantineus*). TL: Ecuador, [Cañar, Río] Angas. D: SW Colombia, W Ecuador, ?NW Perú. Syntypes (♂), NHMUK. **reinst. stat.** [treated as species by Le Crom & Winhard (2024) based on its distinctive wing pattern, our molecular data show that this taxon groups with other *M. menapis* taxa, albeit with some distance between the clades, and we tentatively treat it as a subspecies, for the present]

***menapis menapis*** Hewitson, [1856]: [17], pl. [9], fig. 1. TL: "New Granada". D: Colombia. Syntypes (♀), NHMUK.

*franis* Reakirt, 1868: 90. TL: Colombia, [Cundinamarca], "Insagasugá" (= Fusagasugá). Syntypes (♀), FMNH. **reinst. syn.** [treated as species by Le Crom & Winhard (2024) based on a narrower black DHW postdiscal band and orange DFW tornal marking, both of these phenotypes occur close together in the same clade in our data (vouchers Sal_5683 and Sal_4723, for example), with variation in both characters among other specimens; we therefore restore the name as a synonym of *M. menapis menapis*]

*menapis* var. *peruana* Hopffer, 1879: 419. TL: ["Bolivia"] (error). Lectotype ♂, MfN (Lamas, MS).

*menapis anchicayaensis* Le Crom & Winhard, 2024: 5, 31, pl. 33. TL: [Colombia, Valle del Cauca], Anchicayá, 400-500m. Holotype ♂, MPUJ. **n. syn.** [the holotype is within the range of typical variation of *M. menapis menapis*, and we conclude that all four type specimens, bearing the same label data in the collection of the Universidad Javeriana in Bogotá, Colombia (Le Crom & Winhard, 2024), are mislabeled]

***menapis nariniensis*** Le Crom & Winhard, 2024: 5, 30, pl. 31. (*M. doryssus nariniensis*). TL: [Colombia], Nariño, Ricaurte, La Planada, 1600m. D: SW Colombia, ?NW Ecuador. Holotype ♂, IAVH. **rev. stat.** [described as a subspecies of '*M. doryssus*' [*M. menapis doryssus*] by Le Crom & Winhard (2024), this phenotype resembles specimens known from northwestern Ecuador which DNA barcodes indicate to belong to *M. menapis*, and we thus place it as a subspecies of that species]

***menapis occasiva*** Fox, 1967: 117, figs. 104, 108. TL: Colombia, Cauca, Munchique, [2000m]. D: W Colombia (Cauca). Holotype ♂, AMNH. **reinst. stat.** [treated as a subspecies of *Mechanitis franis* by Le Crom & Winhard (2024), see discussion under *franis*]

*aurantiacus tenebrosus* Winhard, *in* Le Crom & Winhard, 2024: 5, 29, pl. 30. TL: [Colombia], Valle [del Cauca], Quebrada El Pital, 10km N La Paila, 800m. Holotype ♂, ZSM. **n. syn.** [described from a single specimen from Valle del Cauca (Le Crom & Winhard, 2024), the status of this name is speculative; the reduced yellow DFW postdiscal band is consistent with *M. macrinus macrinus* (although some specimens of that taxon do show a broken band, albeit not as reduced as in the HT of *tenebrosus*), while the reduced DHW orange band is consistent with *M. menapis occasiva*. Without DNA sequence data the issue is unlikely to be resolved, but given the large wing pattern variation in all *Mechanitis* species from western Colombia, and the fact that this subspecies would otherwise be based on a single specimen, we believe there is insufficient evidence to regard it as valid, and we treat it for the present as a synonym of *M. menapis occasiva*]

*franis salazari* Le Crom, *in* Le Crom & Winhard, 2024: 5, 31, pl. 32. TL: [Colombia], Caldas, Anserma, 1000m. Holotype ♂, IAVH. **n. syn.** [described by Le Crom in Le Crom & Winhard (2024) based on the thin yellow line joining the postdiscal and discal markings on the DFW, and more extensive black markings, in comparison with *M. m. occasiva*; both are variable characters within series of specimens from the same locality, e.g. Valle del Cauca, Cali, Cañas Gordas (specimens in MGCL), and we therefore treat this name as a synonym of *M. m. occasiva*]

*franis morales* Winhard, *in* Le Crom & Winhard, 2024: 5, 31, pl. 32. TL: [Colombia], Cauca, Vereda Los Cafés, cerca de Morales, 1550m. Holotype ♂, ZSM. **n. syn.** [described by Winhard in Le Crom & Winhard (2024) based on the narrower black bar at the end of the FW discal cell and supposedly broader orange DHW postdiscal band;

both are variable characters within series of specimens from the same locality, e.g. Valle del Cauca, Cali, Cañas Gordas (specimens in MGCL), and we therefore treat this name as a synonym of *M. m. occasiva*]

*menapis flavus* Winhard & Le Crom, *in* Le Crom & Winhard, 2024: 5, 32, pl. 33. TL: [Colombia], Valle [del Cauca], Calcedonia, 1150m. Holotype ♂, IAVH. **n. syn.** [described by Le Crom & Winhard (2024) based on the more extensive yellow markings in the middle of the DFW, which is variation seen within series of specimens from the same locality, e.g. Valle del Cauca, Cali, Cañas Gordas (specimens in MGCL), and we therefore treat this name as a synonym of *M. m. occasiva*]

*menapis submonte* Le Crom & Winhard, 2024: 5, 32, pl. 33. TL: [Colombia], Meta, Sierra La Macarena, Río Guayabero. Holotype ♂, ICNB. **n. syn.** [the HT is indistinguishable from specimens of *M. menapis* from the Cauca valley (Valle del Cauca), while the figured female (Le Crom & Winhard, 2024, plate 33), is typical of specimens from the Magdalena valley (*M. menapis menapis*). The putative type locality (Meta, Sierra La Macarena, Río Guayabero) is in the Amazon basin, but it, and all of the type specimens with dates, were collected prior to 1960, and the absence of modern records from the Amazon (including in online resources such as iNaturalist), and the fact that there are no clear characters separating this putative taxon from others, suggests the possibility of mislabeling]

***messenoides*** C. Felder & R. Felder, 1865: 356. TL: "New Granada". D: E Colombia, E Ecuador, E Perú, Bolivia. Holotype ♀, NHMUK. *Nomen novum* for *Mechanitis menophilus* Hewitson, [1856] (in part).

*menophilus* Hewitson, [1856]: [17] (in part), pl. [9], fig. 2 (*nec* 3). Misidentification.

*deceptus* Butler, 1873: 154. TL: Ecuador, [Azuay], Cuenca. Syntypes (♂), NHMUK. **n. syn.** [treated as a species by Le Crom & Winhard (2024) due to the orange rather than yellow distal half of the FW median band, Hill et al. (2012) showed that these phenotypes are not distinguishable genetically; although an entirely orange form is somewhat more common in Ecuador than it is in Colombia, both forms are frequent throughout the distribution of this species, so we treat *deceptus* as a synonym of *messenoides*]

*mazaeus simplex* Bryk, 1953: 29. TL: Perú, [San Martín], Roque. Holotype ♀, RMS.

*mazaeus holmgreni* Bryk, 1953: 29. TL: Perú, [Puno], Chaquimayo. Holotype ♂, RMS. **n. syn.** [the HT of *ballucatus* has the distal half of the FW median band tinged yellow, but as discussed under *deceptus*, despite likely geographic variation in the frequency of yellow vs. orange forms, we do not think that they are sufficiently geographically stable to merit recognition as distinct subspecies, so we treat *holmgreni* as a synonym of *messenoides*]

*messenoides ballucatus* Fox, 1967: 110, figs. 98, 102. TL: Bolivia, [Santa Cruz], Río Yapacani. Holotype ♂, CMNH.

*messenoides calidus* Le Crom & Winhard, 2024: 6, 35, pl. 38. TL: [Colombia], Guaviare, San José del Guaviare. Holotype ♂, IAVH. **n. syn.** [described by Le Crom & Winhard (2024) based on the presence of an orange discal band and apex on the DHW, which is an infrequent variation seen within populations of typical *M. messenoides* (e.g., Hill et al. 2012, Fig. 1B)]

*pannifera australis* Le Crom, *in* Le Crom & Winhard, 2024: 6, 36, pl. 40. TL: [Colombia], Putumayo, Puerto Leguízamo, 400m. Holotype ♀, IAVH. **n. syn.** [the HT female has the FW band without a strong distal projection in cell M2-M1, and no white VHW marginal spots, suggesting it is a form of *M. messenoides* (e.g., Hill et al. 2012, Fig. 1B)]

***nesaea*** Hübner, [1820]: pl. [2], figs. 1-4. TL: Unknown. D: NE Brazil. Syntypes (♂♀), lost? **rev. stat.** [as discussed in the text, our molecular data support recognition of this taxon as a distinct species]

*lysimnia* ab. *sulphurescens* Haensch, 1905: 148. TL: Brazil, Bahia. "Lectotype" ♀, MfN (Lamas, 1994: 293). Unavailable name.

***polymnia angustifascia*** Talbot, 1928: 411, pl. 14, fig. 7; pl. 16, figs. 4, 7. TL: Brazil, Mato Grosso, Rio Serragem, 1500'. D: SE Perú, Bolivia, S Brazil. Holotype ♂, NHMUK.

*travassosi* d'Almeida, 1951: 11, pl. 2, figs. 1-2. TL: Brazil, Acre, [alto Rio Juruá]. Holotype ♂, DZUP.

***polymnia apicenotata*** Zikán, 1941: 15 (*Mechanitis p.* r. *a.*). TL: Brazil, [Amazonas], Rio Negro, São Gabriel. D: E Colombia, S Venezuela, NW Brazil. Lectotype ♂, IOC (d'Almeida, 1956: 3, fig. 2).

*polymnia sancti-gabrielis* [sic] Bryk, 1953: 29. TL: Brazil, [Amazonas], São Gabriel. Lectotype ♂, RMS (Lamas, MS).

***polymnia bolivarensis*** Fox, 1967: 77, figs. 71, 74 (*Mechanitis isthmia b.*). TL: Venezuela, Bolívar, Río Suapure. D: E Venezuela, Guyana. Holotype ♂, AMNH.

***polymnia casabranca*** Haensch, 1905: 145, fig. 3 (*Mechanitis c.*). TL: Brazil, [Minas Gerais, Fazenda] Monte Cristo. D: E Brazil, NE Argentina. Lectotype ♂, MfN (Lamas, 1994: 278).

***polymnia caucaensis*** Haensch, 1909: 124 (*Mechanitis p.* f. *caucaënsis* [sic]). TL: Colombia, Valle del Cauca. D: E Panama (?), W Colombia. Lectotype ♂, MfN (Lamas, 1994: 278). **reinst. stat.** [treated as subspecies of *Mechanitis chimborazona* by Le Crom & Winhard (2024)]

*isthmia lapaila* Winhard & Le Crom, *in* Le Crom & Winhard, 2024: 6, 34, pl. 37. TL: [Colombia], Valle del Cauca, Quebrada Pital, 10km N La Paila, 800m. Holotype ♂, ZSM. **n. syn.** [described by Le Crom & Winhard (2024) from four specimens from Valle del Cauca, in a transition zone between *M. p. caucaensis* and *M. p. veritabilis* (treated as distinct species by Le Crom & Winhard (2024)), the status of this name is not clear, but we feel there is insufficient evidence to regard it as representing a distinct phenotype inhabiting a clearly defined geographic region]

*utemaia luteola* Winhard & Le Crom, *in* Le Crom & Winhard, 2024: 6, 32, pl. 34. TL: [Colombia], Valle del Cauca, Quebrada Las Cañas, 10km N La Paila, 900m. Holotype ♂, ZSM. **n. syn.** [described by Le Crom & Winhard (2024) based on yellowish DHW discal band, broader black postdiscal band and margin, and more yellowish markings in cell Cu2-Cu1, in comparison with *M. macrinus utemaia*; the HT is from the same locality as *M. isthmia lapaila*, and it seems most parsimonious to assume they represent the same species; without DNA sequence data and better information about nearby populations and geographic variation, the status of this name will remain unclear]

***polymnia chimborazona*** Bates, 1864: 33, footnote (*Mechanitis c.*). TL: Ecuador, [Chimborazo], western foot of Chimborazo, 2-4000'. D: SW Colombia, W Ecuador, NW Perú. Syntypes (♂♀), NHMUK. **reinst. stat.** [treated as species by Le Crom & Winhard (2024), the molecular data are consistent with this and other Transandean taxa being treated as subspecies of *M. polymnia*]

*werneri* Hering, *in* Hering & Hopp, 1925: 188. TL: Colombia, [Cauca], Río [San Juan de] Micay, El Correo. Holotype ♀, MfN. **n. syn.** [Le Crom & Winhard (2024) applied the name *werneri* to a population of *M. polymnia* from the western slopes of the northern Cordillera Occidental in Colombia, which differs from *M. p. chimborazona* in having a complete black discal band on the VHW in the female; this may well represent a distinct subspecies, but the HT of *werneri* has a reduced black VHW postdiscal band typical of *M. p. chimborazona*, and we thus synonymize it with that subspecies]

***polymnia dorissides*** Staudinger, [1884]: pl. 28, fig. [8] (*Mechanitis d.*); 1885: 62 (*Mechanitis doryssides* [sic]). TL: Perú, [Loreto], Yurimaguas. D: E Colombia, E Ecuador, NE Perú. Lectotype ♂, MfN (Fox, 1967: 81). **reinst. stat.** [raised to species by Le Crom & Winhard (2024), the molecular data show that this taxon forms a clade with Guianan *M. polymnia* and we thus treat it as a subspecies]

*aurantiacus orientalis* Le Crom, *in* Le Crom & Winhard, 2024: 5, 30, pl. 31. TL: [Colombia], Caquetá, San Vicente del Caguán, Vereda Las Vegas, Pato Bajo, Finca Napuraña. Holotype ♂, ICNB. **n. syn.** [the DFW of the HT has the yellow postdiscal band broken, reduced orange in cell Cu2-Cu1, and an extensive yellow-orange subapical band, but these are all characters seen within west Amazonian populations of *M. polymnia*, from which the HT otherwise does not differ]

*dorissides vaupesensis* Le Crom & Winhard, 2024: 6, 33, pl. 35. TL: [Colombia], Vaupés, Río Apaporis, Comunidad de Jirijirimo. Holotype ♀, ICBN. **n. syn.** [described by Le Crom & Winhard (2024) based on a supposedly broader yellow DFW postdiscal band, the HT female is a typical specimen of *M. p. dorissides*, whereas the only other type specimen, the male figured under this name on p. 33 and pl. 35 is *M. mazaeus mazaeus*].

*isthmia remolinensis* Le Crom & Winhard, 2024: 6, 35, pl. 37. TL: [Colombia], Meta, Remolinos. Holotype ♂, IAVH. **n. syn.** [without explanation, Le Crom & Winhard (2024) described this name as a subspecies of *Mechanitis isthmia*, noting that the yellow DFW postdiscal band was broken in the middle (as in the HT of *aurantiaca orientalis* Le Crom, from the same region); this is an uncommon variant within typical *M. p. dorissides* that, based on images in iNaturalist, is not stable even within the putative range of *remolinensis*] *isthmia tacana* Le Crom & Winhard, 2024: 6, 35, pl. 38. TL: [Colombia], Amazonas, Tacana. Holotype ♂, UCaldas. **n. syn.** [Le Crom & Winhard (2024) described this putative subspecies as differing from *isthmia remolinensis* in having reduced black discal markings on the DFW, a variable character throughout the distribution of *M. p. dorissides*, from which the HT otherwise does not differ]

***polymnia egaensis*** Bates, 1862: 531, pl. 56, fig. 7a (*Mechanitis p*. var. *e*.). TL: Brazil, [Amazonas], Ega (= Tefé). D: W Brazil. Lectotype ♀, NHMUK (Fox, 1967: 106).

*obscura* Butler, 1877: 149. TL: Brazil, [Amazonas], Ega (= Tefé); Rio Juruá, Pupunha. Syntypes (♂♀), NHMUK. *Nomen novum* for *Mechanitis egaensis* var. 1 Bates, 1862.

*egaensis contracta* Riley, 1919: 182. TL: Brazil, [Amazonas], Rio Purús, Aliança, Canutama. Syntypes (♂♀), NHMUK.

*egaensis obumbrata* d'Almeida, 1951: 10, pl. 1, fig. 1. TL: Brazil, Acre, alto Rio Juruá. Holotype ♀, MNRJ (lost).

***polymnia eurydice*** Haensch, 1905: 147, fig. 4 (*Mechanitis e*.). TL: Perú, [Cuzco], Illapani. D: E Perú. Lectotype ♂, MfN (Lamas, 1994: 280). **reinst. stat.** [treated as a subspecies of *Mechanitis dorissides* by Le Crom & Winhard (2024)]

*lycidice* f. *argentea* Prüffer, 1922: 5, pl. 2, fig. 3. TL: Perú, [Ayacucho], Monte Rico. Syntypes (♂), lost.

***polymnia isthmia*** Bates, 1863: 247, pl. 29, fig. 1 (*Mechanitis i*.). TL: Panamá, [Panamá], Lion Hill. D: Honduras, E Nicaragua, Costa Rica, W Panamá. Syntypes (♂♀), NHMUK. **reinst. stat.** [treated as a species by Le Crom & Winhard (2024), the molecular data show that this taxon forms a clade with other *M. polymnia*, including the nominate subspecies, and we believe it is best treated as a subspecies]

*isthmica* Bates, 1863: pl. 29, fig. 1. Incorrect original spelling.

*californica* Reakirt, [1866]: 223. TL: "USA, California, Los Angeles" (error). Syntypes (2♂), FMNH.

*ovata* Distant, 1876: xi. TL: "Costa Rica". Syntypes (♀), NHMUK.

***polymnia kayei*** Fox, 1967: 78, figs. 71, 75 (*Mechanitis isthmia k*.). TL: Trinidad, Mt. Michael. D: Trinidad. Holotype ♂, CMNH.

***polymnia lycidice*** Bates, 1864: 33 (*Mechanitis l*.). TL: Guatemala, [Baja Verapaz], Chuacús. D: S México, Guatemala, Belize, El Salvador, W Honduras, W Nicaragua. Lectotype ♂, NHMUK (Lamas, 1994: 275).

*doryssa* Boisduval, 1870: 31. TL: "Honduras"; "México". Syntypes (♂♀), NHMUK. Junior primary homonym of *Mechanitis doryssus* Bates, 1864.

*lycidice* f. *arcana* Haensch, 1909: 126. TL: Honduras, [Cortés, San Pedro Sula]. Lectotype ♀, MfN (Lamas, 1994: 275).

***polymnia mauensis*** Forbes, 1948: 15, fig. 4. TL: Brazil, [Pará], Santarém. D: Brazil (Tapajós). Holotype ♂, CUIC.

***polymnia polymnia*** (Linnaeus), 1758: 466 (*Papilio*). TL: "Surinam". D: Guyana, Suriname, French Guiana, N Brazil. Lectotype ♀, UUZM (Honey & Scoble, 2001: 370).

*mopsus* (Linnaeus), 1758: 487 (*Papilio*). TL: "Indiis". Syntypes, lost?

*plagigera* Butler, 1877: 150. TL: Brazil, [Pará], Prainha. Syntypes (♂), NHMUK.

*polymnia atrata* Brévignon, 2007: 539, pl. 1, figs. 1-4. TL: French Guiana, Saint-Laurent-du-Maroni. Holotype ♀, coll. Brévignon. **n. syn.** [the extent of black on the HW is locally variable in this and many co-mimetic taxa from the Guianan region]

***polymnia proceriformis*** Bryk, 1953: 30 (*Mechanitis doryssus p*.). TL: Perú, [San Martín], Roque. D: E Perú (Huallaga). Holotype ♂, RMS.

***polymnia veritabilis*** Butler, 1873: 155 (*Mechanitis v*.). TL: Colombia, "Bogotá" (error); "Venezuela". D: E Panamá, N & W Colombia, N Venezuela. Syntypes (♂♀), NHMUK. **reinst. stat.** [treated as a subspecies of *Mechanitis isthmia* by Le Crom & Winhard (2024)]

*utemaia fluminea* Le Crom & Winhard, 2024: 6, 33, pl. 34. TL: [Colombia], Santander, Socorro, Potrero. Holotype ♂, ICNB. **n. syn.** [described by Le Crom & Winhard (2024) in comparison with *Mechanitis 'lysimnia' utemaia*, the HT is a typical specimen of *M. p. veritabilis*]

*isthmia occidentalis* Winhard & Le Crom, *in* Le Crom & Winhard, 2024: 6, 34, pl. 36. TL: [Colombia], Valle del Cauca, Río Dagua, El Salto, W of Cisneros, 300m. Holotype ♂, ZSM. **n. syn.** [described by Le Crom & Winhard (2024) from two specimens from Valle del Cauca and Risaralda, they noted the more yellowish discal cell patch in comparison with typical *M. p. veritabilis*; given the few specimens examined and lack of information about neighboring populations, this might represent a distinct subspecies on the western slopes of the Cordillera Occidental, or transitional forms between *M. p. veritabilis* and *M. p. isthmia*; for the moment, we feel there is insufficient evidence to regard this as a distinct phenotype inhabiting a clearly defined geographic region]

*Nomina nuda*

*anah* Doubleday, 1847: 130 ("Guyana")
*balea* Doubleday, 1847: 130 ("Honduras")
*baucis* Doubleday, [1845]: 148 ("Venezuela")
*polymnia* var. *cruda* Fassl, 1914: 43
*didyma* Doubleday, 1847: 130 ("Brazil")
*polymnia* var. *donna* Fassl, 1914: 43
*doryssa* Doubleday, [1845]: 55 ("Colombia")
*edocla* Doubleday, 1847: 130
*francesca* Doubleday, 1847: 130 ("Brazil")
*hispulla* Doubleday, 1847: 130 ("Brazil")
*lysidice* Doubleday, [1845]: 55 ("Colombia")
*macrinellus* Herrich-Schäffer, 1865: 175
*neria* Herrich-Schäffer, 1865: 175
*rythiria* Herrich-Schäffer, 1865: 175
*zarina* Doubleday, 1847: 130 ("Bolivia")
*zillah* Doubleday, 1847: 130 ("Honduras")

Genus ***Melinaea*** Hübner

***Melinaea*** Hübner, 1816: 11. Type-species, *Papilio egina* Cramer (Scudder, 1875).
*Melinaea* Bates, 1862: 549. Type-species, *Papilio egina* Cramer (Hemming, 1964: 118). Junior homonym and objective synonym.
*Czakia* Kremky, 1925: 192. Type-species, *Melinaea mneme* var. *mediatrix* Weymer (orig. des.).

***ethra*** (Godart), 1819: 221 (*Heliconia*). TL: "Brazil". D: E Brazil. Syntypes (♀), MNHN.
*phasis* Herrich-Schäffer, 1865: 175. *Nomen nudum*.
*phasis* C. Felder & R. Felder, 1865: 354. TL: Brazil, Bahia. Lectotype ♂, NHMUK (Lamas, MS).

***idae idae*** (C. Felder & R. Felder), 1862: 414 (in part) (*Mechanitis*). TL: Colombia, "Bogotá, 7200'" (error). D: E Panamá, NW Colombia. Lectotype ♂, NHMUK (Lamas, MS).
***idae vespertina*** Fox, 1945: 1, fig. 21. TL: Ecuador, Los Ríos, Playas de Juan Montalvo, 30m. D: SW Colombia, W Ecuador. Holotype ♂, AMNH.
*idae* (C. Felder & R. Felder), 1862: 414 (in part) (*Mechanitis*). Misidentification.

***isocomma isocomma*** Forbes, 1948: 6, pl. 1, fig. 2. TL: Colombia, [Villavicencio], upper Río Negro, 800m. D: E Colombia. Holotype ♂, CUIC.
***isocomma simulator*** Fox, 1960: 162, pl. 5, fig. 69 (*Melinaea comma s.*). TL: Ecuador, [Pastaza], Puyo, 1000m. D: E Ecuador, E Perú. Holotype ♂, CMNH.

***lilis dodona*** Hopffer, 1874: 344 (*Melinaea d.*). TL: "Bolivia" (error). D: W Colombia. Holotype ♀, MfN.
*lilis imitata* f. *erica* Bargmann, 1929: 141. TL: Colombia, [Valle del Cauca], Río Dagua. “Syntypes” (♂♀), NHMUK. Unavailable name.
***lilis ezra*** Fox, 1939: 74. TL: Colombia, Magdalena, Minca. D: N Colombia, NW Venezuela. Holotype ♂, CMNH.

***lilis flavicans*** Hoffmann, 1924: 70, fig. 3 (*Melinaea imitata f.*). TL: México, "Tabasco, Villa Hermosa" (error) [= Colima, Colima]. D: W México. Holotype ♀, AMNH.

***lilis imitata*** Bates, 1864: 55 (*Melinaea i.*). TL: Guatemala, [Alta Verapaz, Río Polochic valley]. D: E México to Atlantic Costa Rica. Syntypes (♂), NHMUK.

*tachypetis* Herrich-Schäffer, 1865: 175. *Nomen nudum*.

*tachypetis* C. Felder & R. Felder, 1865: 355. TL: "México". Lectotype ♂, NHMW (Lamas, MS).

***lilis kayei*** Brown, 1977: 167, fig. 24 (*Melinaea ethra k.*). TL: Guyana, [Mazaruni-Potaro], Quonga. D: Guyana. Holotype ♂, NHMUK.

***lilis lateapicalis*** A. Hall, 1935: 227. TL: Venezuela, Mérida. D: N Colombia, SW Venezuela. Syntypes (2♂), NHMUK.

***lilis lilis*** (Doubleday), 1847: 130, pl. 17, fig. 4 (*Mechanitis*). TL: "Venezuela". D: N Venezuela. Holotype ♂, NHMUK.

***lilis limitata*** A. Hall, 1935: 227. TL: "Venezuela". D: NE Venezuela. Syntypes (2♂, 1♀), NHMUK. [raised to subspecies by Neild, 2008: 98]

***lilis messatis*** (Hewitson), [1856]: [18], pl. [9], fig. 4 (*Mechanitis m.*). TL: "New Granada". D: E Panamá, N Colombia. Neotype ♂, NHMUK (Fox, 1965: 78), **reinst. stat.** [treated as a species by Le Crom & Winhard (2024) on the basis of sympatry with *M. lilis dodona* in west Colombia; genomic data from two similar white (*M. lilis parallelis*) and yellow (*M. lilis imitata*) taxa that are sympatric in Panama grouped specimens by phenotype, although they were not differentiated in the mitochondrial DNA alone. Nevertheless, the split in the genomic tree is very shallow, more study is needed to determine the status of these phenotypes, and for the present we believe they are best treated as representing a single species]

***lilis parallelis*** Butler, 1873: 155 (*Melinaea p.*). TL: Panamá, [Panamá, Lion Hill]. D: SE Costa Rica, Central Panamá. Holotype ♀, NHMUK.

***lilis sola*** Kaye, 1925: 413 (*Melinaea mneme s.*). TL: Trinidad, [St. Patrick, Silverstream River]. D: Trinidad. Syntypes (♀), MGCL.

***ludovica ludovica*** (Cramer), 1780: 17, 250, pl. 297, fig. E (*Papilio*). TL: "Surinam". D: S Venezuela, S Guyana, S Surinam, French Guiana, N Brazil, E Colombia, E Ecuador, NE Perú, E Bolivia. Syntypes (♂), NHMUK?

*egina* (Cramer), 1777: 144, 148, pl. 191, fig. D (*Papilio*). TL: "Surinam". Syntypes (♂), NHMUK? Junior homonym of *Papilio egina* Cramer, 1775.

***ludovica manuelito*** Tessmann, 1928: 119, pl. 5, fig. 22 (*Melinaea m.*). TL: Perú, [Pasco], Pozuzo valley, 750m; between San Nicolás and Puchalina, Pichis road, ca. 1000m. D: E Perú. Lectotype ♂, MfN (Lamas, MS).

***ludovica paraiya*** Reakirt, 1866: 242 (*Melinaea p.*). TL: Brazil, Rio de Janeiro; [Santa Catarina], Ilha de Santa Catarina. D: E Brazil. Syntypes (♂), FMNH.

***maeonis maeonis*** Hewitson, 1869: 11. TL: Ecuador, [Pastaza], Sarayacu. D: E Colombia, E Ecuador, N Perú. Syntypes (♂), NHMUK, **rev. stat.**

*strigilis* Weymer, 1891: 280. TL: Unknown. Holotype ♂, MfN.

*madeira aequatoriensis* d'Almeida, 1951: 23, pl. 5, fig. 1. TL: "Ecuador". Holotype ♂, MNRJ (lost).

***maeonis zamora*** Haensch, 1909: 123 (*Melinaea m.* f. *z.*). TL: "Perú, Ucayali" (error) [= Venezuela, Zamora (= Barinas)]. D: SW Venezuela. Lectotype ♂, MfN (Lamas, 1994: 296), **rev. stat.**

*maeonis borealis* Hall, 1935: 226. TL: Venezuela, [Mérida], Mucuchachí. Syntypes, NHMUK (3♂, 1♀).

***marsaeus clara*** Rosenberg & Talbot, 1914: 672 (*Melinaea orestes c.*). TL: Perú, [Puno], Yahuarmayo, 1200'. D: SE Perú, NW Bolivia, SW Brazil. Holotype ♂, NHMUK. Fig.- Jordan (1923: pl. 4, fig. 9).

*clara juruaënsis* [sic] d'Almeida, 1943: 165, pl., fig. [1]. TL: Brazil, [Amazonas], Rio Juruá. Holotype ♂, MZUSP. [*m. clara* x *m. pothete*].

***marsaeus macaria*** Godman & Salvin, 1898: 107 (*Melinaea macaria*). TL: Colombia, [Meta], San Martín, Llanos of Río Meta. D: E Colombia, E Ecuador, NE Perú. Syntypes (5♂), NHMUK, **reinst. stat.** [treated as a species by Le Crom & Winhard (2024) on the basis of wing pattern differences in comparison with "*M. marsaeus*"; although there are no molecular data for the nominate subspecies, the genomic data for four other taxa, *macaria*, *rileyi*, *phasiana*, and *clara*, support these as being conspecific, despite their wing pattern differences, so for the present we retain the previously existing taxonomy until further evidence suggests it should be modified]

*macaria casanare* Winhard, 2021: 478, fig. 1. Tl: Colombia, Casanare, Vereda Cunamá, 650m. Holotype ♀, ZSM, **n. syn.** [Winhard (2021) stated that this taxon differed from *M. marsaeus macaria* in having reduced black in the wings, but this character is variable throughout the range of the species, and we regard the two listed type specimens from a single locality as insufficient evidence to conclude that they represent a consistent phenotype within a larger geographic area; the name was not mentioned by Le Crom & Winhard (2024)]

***marsaeus marsaeus*** (Hewitson), 1860: [28], pl. [14], fig. 10 (*Mechanitis*). TL: [Brazil], Amazonas, [Ega (= Tefé)]. D: W Brazil. Lectotype ♂, NHMUK (Lamas, 1994: 286).

*maelus* var. (Hewitson), 1860: [27] (in part), pl. [14], fig. 6 (*nec* 9) (*Mechanitis*).

*manga* Haensch, 1909: 123. TL: [Brazil], Amazonas, [Ega (= Tefé)]. Holotype ♀, NHMUK. *Nomen novum* for *Mechanitis maelus* var. Hewitson, 1860. [*m. marsaeus* x *m. phasiana*].

***marsaeus phasiana*** Butler, 1870: 489 (*Melinaea p.*). TL: "Peruvian Amazons", [Loreto, Nauta]. D: E Perú, W Brazil. Syntypes (♀), NHMUK.

*lucifer* var. *divisa* Staudinger, 1885: 71. TL: Perú, [Loreto], Yurimaguas. Lectotype ♂, MfN (Lamas, MS). [*m. phasiana* x *m. rileyi*].

*acreana* d'Almeida, 1951: 22, pl. 4, fig. 2. TL: Brazil, Acre, alto Rio Juruá. Holotype ♂, MNRJ (lost). [*m. phasiana* x *m. rileyi* x *m. marsaeus*]

***marsaeus pothete*** d'Almeida, 1945: 12, pl. 2, figs. 1-3. TL: Brazil, [Rondônia], Rio Verde. D: SW Brazil. Holotype ♂, MNRJ (lost).

***marsaeus rileyi*** Fox, 1942: 3. TL: Perú, [Amazonas], upper Río Marañón, [Pongo de Manseriche]. D: SE Ecuador, SE Colombia, NE Perú, W Brazil. Holotype ♂, AMNH.

*macaria rocafuerte* Winhard, 2021: 478, figs. 4-5. TL: Ecuador, Orellana, Laguna Jatuncocha, 200m. Holotype ♀, ZSM, **n. syn.** [the female holotype has the FW yellow band tinged orange, like the type of *rileyi*, and given that *rileyi* was not mentioned in the original description, we see no reason to regard it as anything other than a specimen of *M. m. rileyi*]

***menophilus cocana*** Haensch, 1903: 164. TL: Ecuador, [Napo], Coca. D: SE Colombia, E Ecuador, NE Perú. Lectotype ♂, MfN (Lamas, 1994: 278).

***menophilus ernestoi*** Brown, 1977: 180, figs. 62-63. TL: Colombia, Caquetá, 40km above Florencia, 1000m. D: SE Colombia. Holotype ♂, MNRJ (lost), **reinst. stat.** [treated as a subspecies of *M.* "*zaneca*" [= *zaneka*] by Le Crom & Winhard (2024), see discussion under that taxon]

***menophilus hicetas*** Godman & Salvin, 1879: 150 (*Melinaea h.*). TL: Perú, [Loreto], Iquitos. D: E Colomibia, E Perú. Lectotype ♂, NHMUK (Lamas, 1994: 285), **reinst. stat.** [Le Crom & Winhard (2024) treated this taxon as a species based on sympatry with *M. menophilus menophilus*, but genomic data of both phenotypes suggest that they are conspecific, and that the yellow FW apical spots that differentiate them represent variation that is partly local, partly geographic]

*hicetas* ab. *flavosignata* Staudinger, 1885: 71. TL: Perú, [Loreto], Pebas; Yurimaguas. “Lectotype” ♂, MfN (Lamas, MS). Unavailable name.

*egesta* Godman & Salvin, 1898: 107. TL: "Interior of Colombia"; Perú, [Loreto], Yurimaguas. Syntypes (4♂), NHMUK. [*m. hicetas* x *m*. ssp. n.]

*magnifica* Haensch, 1905: 145. TL: "Perú". Holotype ♂, MfN.

*hicetas godmani* Winhard & Le Crom, *in* Le Crom & Winhard, 2024: 5, 23, pl. 22. TL: [Colombia], Casanare, Vereda Cunamá, near Planbrisas, N of Aguazul, 650m. Holotype ♂, ZSM, **n. syn.** [Le Crom & Winhard (2024) stated that the name *egesta* was invalid, representing a hybrid between subspecies, and, (presumably) assuming that the name otherwise would apply to an east Colombian taxon, proposed the name *godmani* based on specimens from Casanare, Cundinamarca and Meta; however, we know of no characters that might consistently differentiate *godmani* from *hicetas*, and none were described by Winhard & Le Crom (2024)] *hicetas staudingeri* Winhard & Le Crom, *in* Le Crom & Winhard, 2024: 5, 24, pl. 22. TL: [Colombia], Meta, Humadea, N of San Martín, 500m. Holotype ♂, ZSM, **n. syn.** [the reduced yellow FW apical spots described Le Crom & Winhard (2024) as differentiating this taxon from *M.* "*hicetas godmani*" are variable within populations]

***menophilus maenius*** (Hewitson), 1860: [27], pl. [14], fig. 7 (*Mechanitis maenius*). TL: [Brazil], Amazonas, [Tefé (= Ega)]. D: W Brazil. Syntypes (♂), NHMUK.

***menophilus mediatrix*** Weymer, 1891: 282 (*Melinaea mneme* var. *mediatrix*). TL: French Guiana, Cayenne. D: S Venezuela, Guyana, Surinam, French Guiana, N Brazil. Lectotype ♂, MfN (de Lesse, 1970: 855).

*mediatrix* ab. *anina* Haensch, 1909: 123. TL: "Guyana". "Lectotype" ♂, MfN (Lamas, 1994: 275). Unavailable name.

***menophilus menophilus*** (Hewitson), [1856]: [17] (in part), pl. [9], fig. 3 (*nec* 2) (*Mechanitis*). TL: "New Granada". D: E Colombia. Syntypes (♂), NHMUK.

*ishka* Butler, 1870: 489. TL: Colombia, "Bogotá" (error). Syntypes (♀), NHMUK.

*menophilus membrosa* Forbes, 1927: 33. TL: Unknown. Syntypes, CUIC?

***menophilus orestes*** Salvin, 1871: 412 (*Melinaea o.*). TL: Perú, [Pasco], Pozuzo. D: E Perú, Bolivia, W Brazil. Syntypes (♂), NHMUK.

*chincha* Druce, 1876: 211, pl. 17, fig. 3. TL: Perú, [Pasco], Pozuzo. Syntypes (♂), NHMUK.

***menophilus zaneka*** Butler, 1870: 490 (*Melinaea z.*). TL: Ecuador, [Napo], Archidona. D: E Colombia, E Ecuador, N Perú. Lectotype ♂, NHMUK (Lamas, 1994: 280), **reinst. stat.** [treated as a species by Le Crom & Winhard (2024), there is some support for this hypothesis from the genomic data, which group this Andean foothill taxon with *M. m. ernestoi*, separate from lowland *M. menophilus* taxa; nevertheless, given the absence of sympatry between these phenotypes in well-sampled areas of possible overlap, we feel that they are better treated as conspecific; Le Crom & Winhard (2024) stated that *M. m. zaneka* and *M. m. menophilus* are sympatric in Colombia, but the elevational ranges they provided (1400-2000m and 200-1300m, respectively) do not support this claim. Furthermore, the type specimens of the names *maculosa* and *discurrens* (see Warren et al. (2024) under *M. m. zaneka*) may represent hybrids between *M. m. zaneka* and *M. m. menophilus*]

*dora* Strecker, 1876: 120. TL: Ecuador, "Esmeraldas" (error). Holotype ♂, FMNH.

*zaneka* ab. *maculosa* Haensch, 1903: 163. TL: Ecuador, [Tungurahua], Santa Inés. "Lectotype" ♂, MfN (Lamas, 1994: 285). Unavailable name.

*zaneka* ab. *discurrens* Haensch, 1909: 122. TL: Ecuador, [Tungurahua, Santa Inés]. "Lectotype" ♀, MfN (Lamas, 1994: 280). Unavailable name.

***mnasias abitagua*** Brown, 1977: 164, fig. 11. TL: Ecuador, [Napo], km 43 Puyo-Napo road, 600m. D: E Ecuador. Holotype ♀, MNRJ (lost).

***mnasias comma*** Forbes, 1927: 34, pl. 2, fig. [1]; pl. 3, fig. [13] (*Melinaea c.*). TL: Perú, [Junín], Chanchamayo. D: E Perú. Holotype ♂, CUIC.

***mnasias eratosthenes*** A. Hall, 1935: 226, pl. 6, fig. 5 (*Melinaea e.*). TL: "French Guiana". D: SE Venezuela, French Guiana, N Brazil. Holotype ♂, NHMUK.

***mnasias lucifer*** Bates, 1862: 551 (*Melinaea l.*). TL: Brazil, [Amazonas], São Paulo [de Olivença]. D: NE Perú, W Brazil. Syntypes (♂♀), NHMUK, **reinst. stat.** [treated as a subspecies of *Melinaea* (*marsaeus*) *macaria* by Le Crom & Winhard (2024: 23) on the basis of similarity in wing pattern, the figured putative Colombian specimen of *lucifer* differs from the type of *Melinaea lucifer* (see Warren et al. 2024) in the shape of the FW postdiscal band and in lacking orange FW marginal spots, and actually represents a specimen of *M. marsaeus rileyi*]

*hicetas eryx* d'Almeida, 1951: 24, pl. 3, fig. 3. TL: Brazil, Acre, alto Rio Juruá. Holotype ♂, MNRJ (lost). [*m. lucifer* x *m. romualdo*].

***mnasias lutzi*** Fox, 1942: 1, fig. 2 (*Melinaea lucifer l.*). TL: Perú, [Amazonas], upper Río Marañón, [Quebrada Huachinza]. D: N Perú. Holotype ♂, AMNH.

***mnasias mnasias*** (Hewitson), [1856]: [18], pl. [9], fig. 5 (*Mechanitis*). TL: [Brazil, Pará, Pará (= Belém)]. D: Brazil (Pará). Syntypes (♂), NHMUK.

***mnasias neblinae*** Brown, 1977: 164, fig. 12. TL: Brazil, Amazonas, Pico Neblina, 1500m. D: S Venezuela, NW Brazil. Holotype ♂, MIZA.

***mnasias romualdo*** Fox, 1965: 80 (*Melinaea comma r.*). TL: Perú, [Puno], "Carabaya, Río Huacamayo, La Unión, 2000'" [= Yahuarmayo, 1200']. D: SE Perú. Holotype ♂, NHMUK.
***mnasias rondonia*** Brown, 1977: 166, fig. 13. TL: Brazil, Rondônia, near Riozinho, 300m. D: SW Brazil. Holotype ♂, MNRJ (lost).
***mnasias tecta*** Haensch, 1909: 124, pl. 33d, fig. [4] (as *mnasias*) (*Melinaea thera* f. *tecta*). TL: "Guyana". D: Guyana. Holotype ♂, MfN.
***mnasias thera*** C. Felder & R. Felder, 1865: 354 (*Melinaea t.*). TL: Unknown. D: SE Brazil. Holotype ♂, NHMUK.
*thera* Herrich-Schäffer, 1865: 175. *Nomen nudum*.

***mneme mauensis*** Weymer, 1891: 283 (*Melinaea mneme* var. *mauensis*). TL: Brazil, [Amazonas], Ega (= Tefé). D: S Venezuela, E Colombia, NE Perú, Brazil. Holotype ♂, MfN.
***mneme mneme*** (Linnaeus), 1763: 20 (*Papilio*). TL: "China" (error). D: ?SE Venezuela, Guyana, Surinam, French Guiana, N Brazil. Lectotype ♂, LSL (Kaye, 1925: xxii).

***mnemopsis*** Berg, 1897: 234. TL: "Perú" (error?). D: ?SE Perú, Bolivia. Holotype ♂, MBR.
*boliviana* Weymer, 1907: 2, pl. 1, fig. 2. TL: "Bolivia". Holotype ♂, MfN.

***mothone messenina*** C. Felder & R. Felder, 1865: 356, pl. 45, fig. 11 (*Melinaea messenina*). TL: Colombia, "Bogotá" (error). D: E Colombia, W Venezuela. Syntypes (♂), NHMUK.
***mothone mothone*** (Hewitson), 1860: [30], pl. [15], fig. 14 (*Mechanitis*). TL: [“Ecuador”]. D: SE Colombia, E Ecuador, NE Perú. Neotype ♂, NHMUK (Fox, 1965: 79).
*cydippe* Salvin, 1871: 412. TL: Ecuador, [Morona-Santiago], Gualaquiza; Perú, [Pasco], Pozuzo. Syntypes (♂), NHMUK.

***satevis aurantia*** Forbes, 1942: 27 (*Melinaea mneme a.*). TL: Venezuela, Monagas, Caripito. D: NE Venezuela, Trinidad. Syntypes (7♂), CUIC (3♂), MIZA (1♂), USNM (2♂), CMNH (1♂).
***satevis crameri*** Godman & Salvin, 1898: 107 (*Melinaea c.*). TL: Guyana, [Mazaruni-Potaro, Mt. Roraima]. D: E Venezuela, N Guyana, N Surinam. Syntypes (5♂), NHMUK.
*mneme* (Cramer), 1777: 142, 149, pl. 190, fig. C (*Papilio*). Misidentification.
*crameri* ab. *incisa* Kaye, 1925: xxiii. TL: Guyana, [Mazaruni-Potaro], Essequibo River. “Syntypes” (♂), MGCL. Unavailable name.
***satevis cydon*** Godman & Salvin, 1879: 151 (*Melinaea c.*). TL: Brazil, [Amazonas], Tabatinga; Perú, [Loreto], Pebas. D: SE Colombia, E Perú, W Brazil. Syntypes (♂), NHMUK.
***satevis flavomacula*** Weymer, 1894: 322 (*Melinaea pardalis* var. *f.*). TL: [Brazil], "obern Amazonenstrom". D: Brazil. Lectotype ♂, MfN (Lamas, 1994: 285).
*madeira* Kaye, 1904: 4. *Nomen nudum*.
*pardalis madeira* Moulton, 20.i.1909: 597, 604, pl. 32, fig. 2. TL: [Brazil], "Amazons". Holotype ♀, OXF.
*madeira* Haensch, 26.v.1909: 123, pl. 33b, fig. [3]. TL: [Brazil, Amazonas], Manicoré. Lectotype ♂, MfN (Lamas, 1994: 285). Junior primary homonym of *Melinaea pardalis madeira* Moulton, 1909.
***satevis lamasi*** Brown, 1977: 171, figs. 31-32 (*Melinaea ethra l.*). TL: Perú, Madre de Dios, Iberia, 200m. D: SE Perú, SW Brazil. Holotype ♂, MUSM.
***satevis maelus*** (Hewitson), 1860: [27] (in part), pl. [14], fig. 9 (*nec* 6) (*Mechanitis m.*). TL: [Brazil], Amazonas, [Ega (= Tefé)]. D: E Colombia, NE Perú, W Brazil. Syntypes (♂♀), NHMUK.
*pardalis* (C. Felder & R. Felder), iii.1862: 78 (*Mechanitis*). *Nomen nudum*.
*pardalis* Bates, 13.xi.1862: 552. *Nomen novum*.
*hicetas brunnea* Riley, 1919: 181. TL: Brazil, [Amazonas], upper Rio Purús, Boca do Acre. Holotype ♀, NHMUK. [*s. maelus* x *s. lamasi*]
*hicetas purusana* Riley, 1919: 181. TL: Brazil, [Amazonas], Rio Purús, Aliança, Canutama, Sebastopol, Lábrea; [Rondônia], Abunã. Syntypes (♂♀), NHMUK. [*s. maelus* x *s. flavomacula*]

*madeira* var. *purusana* Aurivillius, 1929: 155. TL: Brazil, [Amazonas], Rio Purús, Hiutanaã. Lectotype ♂, RMS (Lamas, MS). Junior primary homonym of *Melinaea hicetas purusana* Riley, 1919. [*s. maelus* x *s. flavomacula*].

*maelus romani* Bryk, 1937: 641. *Nomen novum* for *Melinaea madeira* var. *purusana* Aurivillius, 1929, *nec M. hicetas purusana* Riley, 1919.

***satevis satevis*** (Doubleday), 1847: 130, pl. 17, fig. 3 (*Mechanitis*). TL: "Bolivia". D: ?SE Perú, Bolivia. Holotype ♀, NHMUK.

***scylax*** Salvin, 1871: 412. TL: Panamá, Chiriquí, Bugaba, [800-1000']. D: Pacific Costa Rica, Pacific W Panamá, NW Colombia. Lectotype ♂, NHMUK (Lamas, MS).

*ribbei* Staudinger, v.1875: 97. TL: Panamá, Chiriquí. Lectotype ♂, MfN (Lamas, MS).

*ribbei* Weymer, vii.1875: 379, pl. 2, fig. 4. TL: Panamá, Chiriquí. Lectotype ♂, MfN (Lamas, MS). Junior primary homonym of *Melinaea ribbei* Staudinger, 1875.

***tarapotensis*** Haensch, 1909: 122 (*Melinaea menophilus* f. *t.*). TL: [Perú, San Martín], Tarapoto. D: NE Perú. Lectotype ♀, MfN (Lamas, 1994: 294).

Interspecific hybrids

*agricola* Hall, 1935: 227, pl. 6, fig. 6. TL: Brazil, [Amazonas], Ega (= Tefé). Holotype ♂, NHMUK. [*ludovica ludovica* x *mneme mauensis*]

*mayi* d'Almeida, 1951: 20, pl. 4, fig. 1. TL: Brazil, Acre, alto Rio Juruá. Holotype ♂, MNRJ (lost). [*marsaeus marsaeus* x *mneme mauensis*].

*Nomina nuda*

*eginella* Herrich-Schäffer, 1865: 175.

*vilis* Herrich-Schäffer, 1865: 175.

**Collection codens**

AMNH: American Museum of Natural History, New York, NY, USA
CCWM: Chau Chak Wing Museum, University of Sydney, Sydney, Australia.
CMNH: Carnegie Museum of Natural History, Pittsburgh, USA
CUIC: Cornell University, Ithaca, USA
DZUP: Departamento de Zoologia, Universidade Federal do Paraná, Curitiba, Brazil
FMNH: Field Museum of Natural History, Chicago, IL, USA
IAVH: Instituto Alexander von Humboldt, Bogotá, Colombia
ICNB: Instituto de Ciencias Naturales, Universidad Nacional, Museo de Historia Natural, Bogota, Colombia
IOC: Instituto Oswaldo Cruzo, Rio de Janeiro, Brazil
LSL: Linnaean Collection, Linnaean Society of London, London, United Kingdom
MBR: Museo Argentina de Ciencias Naturales "Bernardino Rivadavia", Buenos Aires, Argentina
MfN: Museum für Naturkunde, Leibniz-Institut für Evolutions- und Biodiversitätsforschung an der Humboldt Universität, Berlin, Germany
MGCL: McGuire Center for Lepidoptera and Biodiversity, Florida Museum of Natural History, Gainesville, USA
MIZA: Museo del Instituto de Zoología Agrícola, Universidad Central de Venezuela, Maracay, Venezuela
MNCN: Museo Nacional de Ciencias Naturales, Madrid, Spain
MNHN: Muséum National d'Histoire Naturelle, Paris, France
MNRJ: Museu Nacional Rio de Janeiro, Rio de Janeiro, Brazil
MPUJ: Museo de Historia Natural de la Pontificia Universidad Javeriana, Bogotá, Colombia
MUSM: Museo de Historia Natural, Universidad Nacional Mayor de San Marcos, Lima, Peru
NHMUK: Natural History Museum, London, UK
OXF: Oxford University Museum, Oxford, UK
RMS: Rijksmuseum, Amsterdam, Netherlands

RPM: Reading Public Museum, Philadelphia, USA
SMTD: Staatliches Museum für Tierkunde, Dresden, Germany
UCaldas: Museo de Historia Natural, Universidad de Caldas, Manizales, Colombia
USNM: National Museum of Natural History, Smithsonian Institution, Washington, DC, USA
UUZM: Uppsala University, Uppsala, Sweden
ZSM: Zoologische Staatssammlung München, Munich, Germany
ZUEC: Museu de Zoologia da Universidade Estadual de Campinas 'Adão José Cardoso', Campinas, Brazil

**Supporting Text 3 – Hybridisation analysis**

The poorly supported nodes and discordance between phylogenies constructed with different methods (Fig. 1 vs Fig. 2), from nuclear versus mitochondrial genomes (Fig. 1) or different genomic regions (Fig. S5) could be due to either ILS or hybridisation. To test for hybridisation, we assessed excess allele sharing between non-sister taxa with Fbranch (Fig. S9) and windowed phylogenetic tree discordance with BPP (Table S2; Fig. S5; Text S2). Moreover, we ran AIM which uses MCMC models to produce a phylogenetic tree that allows for gene flow (Fig. S10). The combined results of all hybridisation analyses are summarised with arrows in Fig. 2 (Text S2), but we would like to stress that this is only one of the multiple possible scenarios consistent with our observed patterns of excess allele sharing and gene tree discordance. Due to hybridisation, there is likely not 'one true species tree' that reflects the 'real' phylogeny of these genera (7, 8).

*Mechanitis*

In *Mechanitis*, various analyses show that there has been ancient hybridisation between *Mec. messenoides* and the ancestor of the *polymnia-lysimnia-nesaea*-clade*.* The IQtree2 nuclear phylogeny places *messenoides* as a sister-species of *menapis* (Fig. 1A), but BEAST2 places *messenoides* as the sister-species of *menapis*, *macrinus* and *mazaeus* (Fig. 2A). AIM places *messenoides* closer to the *polymnia-lysimnia-nesaea*-clade (Fig. S10A) with various admixture arrows between the *menapis-macrinus-mazaeus* clade and the *messenoides-polymnia-nesaea-lysimnia* clade (Fig S10A – arrows B-D). The IQtree2 mitochondrial phylogeny shows that *messenoides* has two mitochondrial haplotypes, one clustering with *menapis* and the other with *polymnia*, *lysimnia* and *nesaea* (Fig. 1A – box II). We performed an Fbranch-analysis (with the BEAST2-tree as the 'species'-tree) to assess allele sharing, and this confirms excess allele sharing between *Mec. messenoides* and *Mec. menapis*, as well as *Mec. messenoides* and the *polymnia*-clade (Fig S9A - #2-3). Moreover, in the PCA (Fig. S22), PC2 groups *Mec. messenoides* with the *polymnia-lysimnia-nesaea* clade. *Mec. menapis* clusters in between *Mec. mazaeus* and *Mec. macrinus* on one side, and *Mec. messenoides* on the other. *Mec. messenoides* co-occurs with *mazaeus*, *polymnia* and *lysimnia* in Western Amazonia, whereas *Mec. menapis* co-occurs with *macrinus* and *polymnia* West of the Andes.

There also appears to have been admixture within the *polymnia-lysimnia*-nesaea-clade. *Mec. nesaea* co-occurs with *polymnia* and *lysimnia* in the Atlantic Forest of Brazil, and both *polymnia* and *lysimnia* have subspecies across South America. *Mec. nesaea* is placed in a different position in the phylogenetic tree with the nuclear IQtree2 (Fig. 1A) and BEAST2 (Fig. 2A) (as sister to *polymnia* and *lysimnia*, respectively), and also the AIM phylogeny shows a different topology (Fig. S10A) (*polymnia* and *lysimnia* as closest sister-species). The mitochondrial IQtree2 phylogeny places *nesaea* as sister to *lysimnia* (Fig. 1A – box III). The Fbranch analysis confirms excess allele sharing between *Mec. nesaea, Mec. polymnia* and *Mec. lysimnia* (Fig S8A - #1*).* There is excess allele sharing between *nesaea* and Brazilian *polymnia*, as well as Brazilian *lysimnia* and Brazilian *polymnia* compared to the rest of *polymnia* (Fig. S9A - #1B, #1C). Interestingly, however, the Ecuadorian *lysimnia* share more alleles with *Mec. nesaea* than Brazilian *lysimnia* (Fig. S9A - #1A). The Ecuadorian *lysimnia* also share more alleles with all of the polymnia-lineages, compared to the Brazilian *lysimnia* (Fig. S9A - #1A).

The BPP-analysis, where we plotted phylogenies in windows across the genome, confirms both the jumping of *messenoides*, and the uncertainty within *polymnia-lysimnia-nesaea*. The three most common trees in the genome vary from each other by the relationship amongst *Mec. lysimnia, Mec. polymnia* and *Mec. nesaea* (Fig S5; accounting for ~41% of the genome). These three trees place *Mec. messenoides* and *Mec. menapis* as closest relatives, which is the topology we found with the IQtree2 analysis, and what the Fbranch analysis confirms with excess allele sharing between *messenoides* and *menapis* (Fig. S9A - #2). According to the BPP-analysis, *Mec. messenoides* clusters in the same clade as *Mec. menapis, mazaeus* and *macrinus* in ~51% of the genome (Fig S5B). However, in almost ~20% of the genome, *Mec. messenoides* is placed closer to or within the clade consisting of *Mec. polymnia*, *Mec. lysimnia* and *Mec. nesaea*, and not with *Mec. menapis*. The branching order within the *polymnia-lysimnia-nesaea* clade is highly variable across the genome*, with* 46.3% of the genome grouping *polymnia* with *nesaea*, 31.7% grouping *polymnia* with *lysimnia*, and 14.7% grouping *nesaea* with *lysimnia* (Fig. 5C).

In the Fbranch-analysis, another signal of excess allele sharing is found, namely between the *mazaeus-macrinus*-clade and the *polymnia-lysimnia-nesaea*-clade (Fig. S9A – #4). AIM also indicates some gene flow between these two clades (Fig. S10A – arrow E). *Mec. polymnia* and *nesaea* share more alleles with *Mec. mazaeus* compared to *Mec. macrinus*.

The relationships between the various *polymnia*-lineages shift around in the various analyses. With IQtree2, *polymnia* from west of the Andes is closest to the individuals from Western Amazonia, but with BEAST2 *polymnia-West* is the earliest diverging lineages in the *polymnia*-clade. Fbranch confirms this admixture (Fig. S9A - #5). We found a single individual in Colombia with equal ancestral contribution of lineages East and West of the Andes (Sal_4944 in Fig. S1; Fig. S8).

*Melinaea*

In *Melinaea*, we find hybridisation between various species and low concordance factors at all nodes, even at the base of the core clade (Fig. 1B). IQtree2 and BEAST produce the same topology, but AIM places *Mel. idae* with another clade and suggests gene flow between *Mel. idae* and the two different sister clades (Fig. 2B - arrow a; Fig. S10B – arrow A), which is confirmed by the Fbranch results (Fig. S9B - #1). We ran the BPP analysis for *Melinaea* in a subset of the species, as including individuals of all species created so many different trees across the genome that the results were difficult to interpret, likely due to the recent divergence and gene flow. To resolve the position of *Mel. idae*, as well as putative introgression between deeper branches of the phylogeny, we focused on the relationships between the *lilis-isocomma-idae* clade and representatives of the other clade (*mneme*, *marsaeus* and *mothone*; thus excluding *satevis*, *tarapotensis*, *maeonis* and *menophilus*) (Fig. 5E-F). The three most common trees in this analysis switch *Mel. idae* from a sister group to *lilis* (~15%; indicating admixture between *lilis* and *idae*, confirmed by Fbranch excess allele sharing; Fig. S5; Fig S8B - #4), to *idae* as the sister to *mneme-marsaeus-mothone* (like the AIM-results; ~12%), to *idae* as sister to both *lilis* and *isocomma* (like IQTree2 and BEAST2; ~10%).

In the majority of the genome (43.4%) *lilis, idae* and *isocomma* group together (like IQTree2 and BEAST2), while in 34.5% of the genome *idae* groups with *mothone-marsaeus-mneme* (like AIM). Notably, the Z chromosome supports almost exclusively the latter relationship. In 4.3% of the genome, *isocomma* clusters with *mothone-marsaeus-mneme*. *Lilis* and *isocomma* (38.3%) are almost as often sister-species compared to *lilis* and *idae* (30.3%), both a lot more than *idae* and *isocomma* (4.1%), indicating admixture between *lilis* and *idae*, which is confirmed by Fbranch excess allele sharing (Fig. S10B; Fig S9B - #2). *Mel. mothone, marsaeus* and *mneme* also vary in their respective relationships (Fig. 5E). In the PCA (Fig. S17B), *isocomma* and *idae* cluster together. *Mel. idae* and *Mel. lilis* are both found west of the Andes, whereas all other clades are found east of the Andes.

Moreover, AIM also places *Mel. satevis* closer to *marsaeus-maeonis* instead of *tarapotensis* (Fig. 2B - arrow c; Fig. S10B; Fig. S9B - #3), and Fbranch also indicates some excess allele sharing between *Mel. marsaeus/maeonis* and *Mel. isocomma* and *Mel. mothone* (Fig. S9B - #2,#7). Both AIM and Fbranch indicate admixture between *Mel. ludovica* and several clades in the ‘core’ *Melinaea* (Fig. 2B - arrows e-f; Fig. S10B – arrows E/F; Fig. S9B - #5-6). BPP always places *Mel. ludovica* as the outgroup in the phylogeny (Fig. S5). There is a slight excess of allele sharing between the *mothone/menophilus*-clade and the *lilis/isocomma/idae*-clade (Fig. S9B; #10). Overall, these results indicate that there has been rampant admixture between many *Melinaea* lineages, and that especially *Mel. idae* has been strongly introgressed.

**Supporting Text 4 – *Mechanitis neseae* – an ancestrally introgressed species**

*Mechanitis nesaea* occurs in the eastern tip of the Brazilian Atlantic Forest. *Mec. lysimnia lysimnia* occurs further south, with a small sympatric zone, and *Mec. polymnia casabranca* largely overlaps with both taxa (Fig. 2) (2, 9). *Mec. nesaea* is intermediate between *Mec. polymnia* and *Mec. lysimnia* in chromosome number (10), *lysimnia*-like in wing colour patterns and caterpillar phenotypes, although the head capsule colour of later caterpillar instars is sometimes more *polymnia*-like (11) (Fig. 3A). We find that in androconial compounds, *Mec. nesaea* is distinct from both putative parental species but closest to *Mec. polymnia* (Fig. 3B, Table S3-4). Some of the differences in androconial compounds might contribute to assortative mating, as has been suggested in other Ithomiini (12–14). Of the 26 androconial compounds we found, three show very high concentrations, two of which are derivatives of PAs: hydroxydanaidal (peak 2) and methyl hydroxydanaidoate (peak 3) (Fig. 3C). The content of these chemicals can vary greatly between individuals, because it depends on the access to PA-containing plants, but we find them to be consistently associated with *Mec. nesaea* and *Mec. p. casabranca* and present only in traces in *Mec. l. lysimnia* (across 92 individuals total from multiple locations, Table S3), consistent with previous findings of *Mec. l. lysimnia* (15). On the other hand, heptacosene (peak 12) was dominant in *Mec. l. lysimnia* while only present as traces in *Mec. nesaea* and absent in *Mec. p. casabranca* samples.

To study genomic signatures of differentiation and introgression and to examine whether introgression of *lysimnia* into *nesaea* contributed to reproductive isolation between *nesaea* and *polymnia*, we resequenced 15 *Mec. nesaea*, 19 *Mec. l. lysimnia* and 9 *Mec. p. casabranca*, of which 24 many were sampled from sympatry. We examined patterns of ADMIXTURE, $F_{ST}$, $D_{XY}$ and $f_{dM}$. ADMIXTURE and high $F_{ST}$ shows no evidence of recent gene flow between either *lysimnia* and *nesaea*, or *polymnia* and *nesaea* (Fig. S13), and introgression plots are consistent between various *nesaea*-individuals (Fig. S13), confirming that *lysimnia*-introgression is likely ancient or ancestral to *Mec. nesaea*. We find high peaks of differentiation that overlap with peaks of introgression between *lysimnia* and *nesaea* and also with chromosomal breakpoints (Fig. 3G; Fig. S13), but as we find overall low recent gene flow between *nesaea* and *polymnia*, it is difficult to interpret our $F_{ST}$ and $f_{dM}$ results. We conclude that more research is necessary, and future studies should test if high-differentiation peaks with signatures of *lysimnia*-introgression harbour reproductive isolation barriers such as genes contributing to chemoreceptor differences.

It is interesting to note that *Mec. lysiminia lysiminia* and *Mec. polymnia casabranca* have been observed to interbreed in nature (16) and putative hybrids have been found, some phenotypically resembling *Mec. nesaea* (Fig. S14). It might be the case that all putative hybrids are infertile F1-individuals, and that recent introgression is unlikely to spread.

**Supporting Text 5 – Methods**

*Androconial Chemistry*

To obtain samples of the androconial secretions, both wing androconia of each sampled male were excised and soaked into 1.5 mL bidistilled HPLC grade hexane. The extracts were filtered with silanized glass wool (Sigma-Aldrich, USA) to remove any debris, and then concentrated to approximately 25 µL under a gentle nitrogen flow. The samples were analysed on a gas chromatograph coupled to a mass spectrometer (GC-MS; Agilent 7890A™ gas chromatograph, Agilent 5975C Series MSD™ mass spectrometer) equipped with a non-polar DB-5ms column (Agilent J&W; 30 m × 0.25 mm i.d., 0.25 µm film thickness). A split/splitless inlet was fitted to an Agilent Thermal Separation Probe (TSP). For the analyses, 1 µL aliquots were injected into quartz micro vials which were then inserted in the TSP vial holder with the inlet set to split mode (1:1) and the GC injector temperature set to 250 °C. The GC oven temperature started at 60 °C for 2 min and then increased at a rate of 10 $°C \cdot min^{-1}$ to 280 °C. The final temperature was maintained for 12 min. Helium carrier gas flow was maintained at a constant pressure of 7.3 psi. MS source and quadrupole temperatures were set at 230 °C and 150 °C, respectively. Mass spectra were taken at 70 eV (in EI mode) with a scanning speed of 1.0 $scan \cdot s^{-1}$ from m/z 35–450. To exclude non-androconial compounds from our analyses, the same extraction procedure was adopted to excise wing areas of conspecific females (n=2 / population) and non-androconial wing areas of males (n=2 / population). Solvent negative controls were also taken for each sampling event (n=8).

*Reference genomes*

High molecular weight DNA was extracted from half a thorax using the Qiagen MagAttract High Molecular Weight kit (Qiagen ID 67563), followed by shearing into an average fragment size between 12–20 kb in a Megaruptor 3 system. The sheared DNA was used for low-input PacBio library preparation according to the manufacturers' instructions (see Tree of Life protocol (17)). Pacific Biosciences HiFi circular consensus libraries were constructed and sequencing was performed on PacBio Revio. Next, we used the head for HiC library prep using Arima v2 kit following the manufacturer's protocol, followed by 150 bp paired-end Illumina sequencing on a Novaseq 6000 performed by Scientific Operations at Wellcome Sanger Institute.

The genomes were assembled with Hifiasm version 0.19.5 (18), duplicates removed with purge_dups version 1.2.5 (19). The draft assembly was scaffolded with HiC-data using YaHS version 1.2a.2 (20), and both haplotypes were assembled separately. Contamination check and correction was performed with the TreeVal pipeline version 1.0.0 (21), thereafter the assemblies were manually curated with Higlass (22) and PretextView (https://github.com/sanger-tol/PretextView). Post-processing and evaluation of the genome assemblies were performed with three pipelines: sanger-tol/readmapping (23), sanger-tol/genomenote (24), and sanger-tol/blobtoolkit (25).

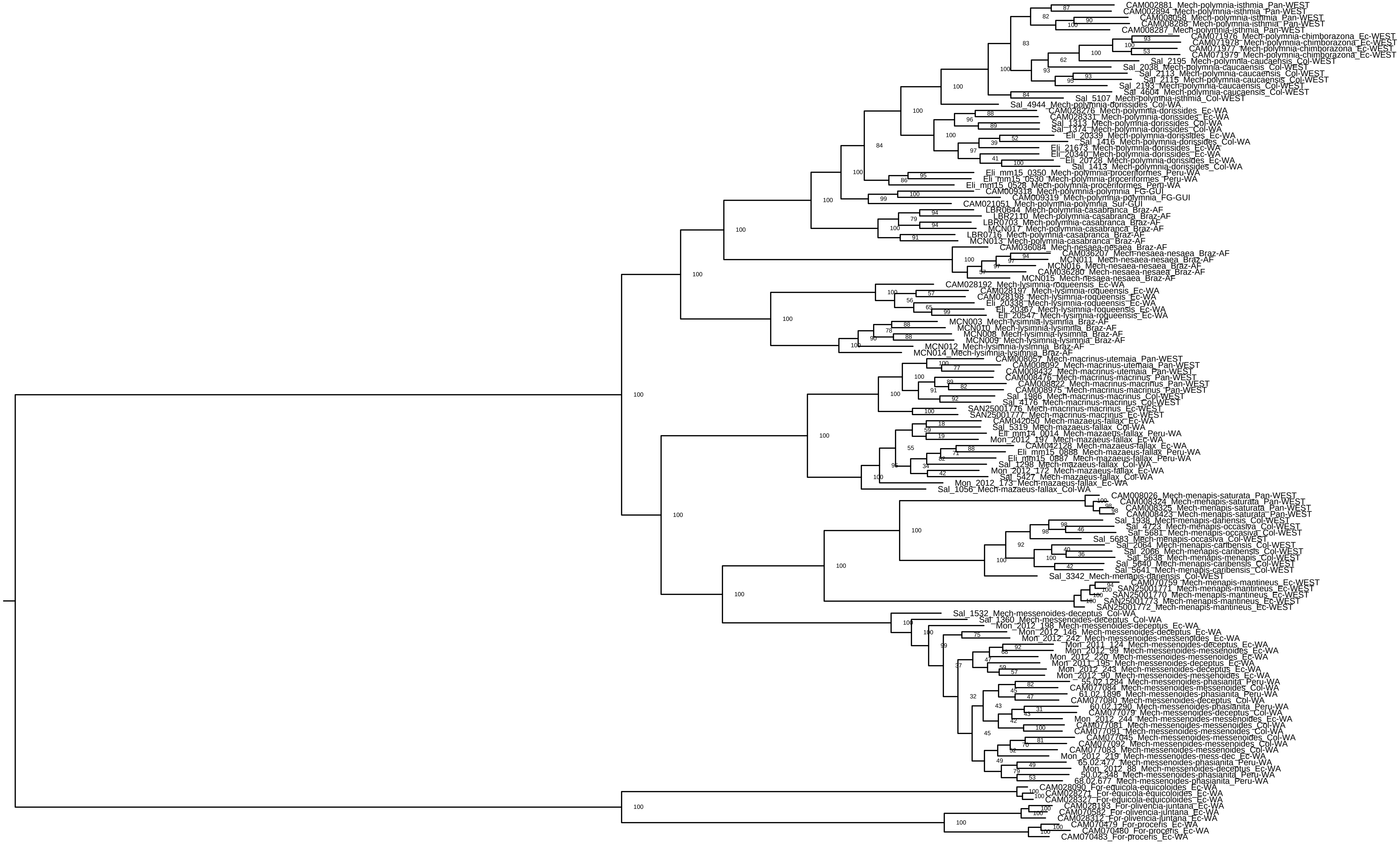
CAM002881_Mech-polymnia-isthmia_Pan-WEST
CAM002894_Mech-polymnia-isthmia_Pan-WEST
CAM008058_Mech-polymnia-isthmia_Pan-WEST
CAM008288_Mech-polymnia-isthmia_Pan-WEST
CAM008287_Mech-polymnia-isthmia_Pan-WEST
CAM071976_Mech-polymnia-chimborazona_Ec-WEST
CAM071978_Mech-polymnia-chimborazona_Ec-WEST
CAM071977_Mech-polymnia-chimborazona_Ec-WEST
CAM071979_Mech-polymnia-chimborazona_Ec-WEST
Sal_2195_Mech-polymnia-caucaensis_Col-WEST
Sal_2038_Mech-polymnia-caucaensis_Col-WEST
Sal_2113_Mech-polymnia-caucaensis_Col-WEST
Sal_2115_Mech-polymnia-caucaensis_Col-WEST
Sal_2193_Mech-polymnia-caucaensis_Col-WEST
Sal_4604_Mech-polymnia-caucaensis_Col-WEST
Sal_5107_Mech-polymnia-isthmia_Col-WEST
Sal_4944_Mech-polymnia-dorissides_Col-WA
CAM028276_Mech-polymnia-dorissides_Ec-WA
CAM028331_Mech-polymnia-dorissides_Ec-WA
Sal_1313_Mech-polymnia-dorissides_Col-WA
Sal_1374_Mech-polymnia-dorissides_Col-WA
Eli_20339_Mech-polymnia-dorissides_Ec-WA
Sal_1416_Mech-polymnia-dorissides_Col-WA
Eli_21673_Mech-polymnia-dorissides_Ec-WA
Eli_20340_Mech-polymnia-dorissides_Ec-WA
Eli_20728_Mech-polymnia-dorissides_Ec-WA
Sal_1413_Mech-polymnia-dorissides_Col-WA
Eli_mm15_0350_Mech-polymnia-proceriformes_Peru-WA
Eli_mm15_0530_Mech-polymnia-proceriformes_Peru-WA
Eli_mm15_0528_Mech-polymnia-proceriformes_Peru-WA
CAM009318_Mech-polymnia-polymnia_FG-GUI
CAM009319_Mech-polymnia-polymnia_FG-GUI
CAM021051_Mech-polymnia-polymnia_Sur-GUI
LBR0644_Mech-polymnia-casabranca_Braz-AF
LBR2110_Mech-polymnia-casabranca_Braz-AF
LBR0703_Mech-polymnia-casabranca_Braz-AF
MCN017_Mech-polymnia-casabranca_Braz-AF
LBR0716_Mech-polymnia-casabranca_Braz-AF
MCN013_Mech-polymnia-casabranca_Braz-AF
CAM036084_Mech-nesaea-nesaea_Braz-AF
CAM036207_Mech-nesaea-nesaea_Braz-AF
MCN011_Mech-nesaea-nesaea_Braz-AF
MCN016_Mech-nesaea-nesaea_Braz-AF
CAM036280_Mech-nesaea-nesaea_Braz-AF
MCN015_Mech-nesaea-nesaea_Braz-AF
CAM028192_Mech-lysimnia-roqueensis_Ec-WA
CAM028197_Mech-lysimnia-roqueensis_Ec-WA
CAM028198_Mech-lysimnia-roqueensis_Ec-WA
Eli_20338_Mech-lysimnia-roqueensis_Ec-WA
Eli_20367_Mech-lysimnia-roqueensis_Ec-WA
Eli_20547_Mech-lysimnia-roqueensis_Ec-WA
MCN003_Mech-lysimnia-lysimnia_Braz-AF
MCN010_Mech-lysimnia-lysimnia_Braz-AF
MCN008_Mech-lysimnia-lysimnia_Braz-AF
MCN009_Mech-lysimnia-lysimnia_Braz-AF
MCN012_Mech-lysimnia-lysimnia_Braz-AF
MCN014_Mech-lysimnia-lysimnia_Braz-AF
CAM008057_Mech-macrinus-utemaia_Pan-WEST
CAM008092_Mech-macrinus-utemaia_Pan-WEST
CAM008432_Mech-macrinus-utemaia_Pan-WEST
CAM008476_Mech-macrinus-macrinus_Pan-WEST
CAM008822_Mech-macrinus-macrinus_Pan-WEST
CAM008975_Mech-macrinus-macrinus_Pan-WEST
Sal_1986_Mech-macrinus-macrinus_Col-WEST
Sal_4176_Mech-macrinus-macrinus_Col-WEST
SAN25001776_Mech-macrinus-macrinus_Ec-WEST
SAN25001777_Mech-macrinus-macrinus_Ec-WEST
CAM042050_Mech-mazaeus-fallax_Ec-WA
Sal_5319_Mech-mazaeus-fallax_Col-WA
Eli_mm14_0014_Mech-mazaeus-fallax_Peru-WA
Mon_2012_197_Mech-mazaeus-fallax_Ec-WA
CAM042128_Mech-mazaeus-fallax_Ec-WA
Eli_mm15_0888_Mech-mazaeus-fallax_Peru-WA
Eli_mm15_0887_Mech-mazaeus-fallax_Peru-WA
Sal_1298_Mech-mazaeus-fallax_Col-WA
Mon_2012_172_Mech-mazaeus-fallax_Ec-WA
Sal_5427_Mech-mazaeus-fallax_Col-WA
Mon_2012_173_Mech-mazaeus-fallax_Ec-WA
Sal_1056_Mech-mazaeus-fallax_Col-WA
CAM008026_Mech-menapis-saturata_Pan-WEST
CAM008324_Mech-menapis-saturata_Pan-WEST
CAM008325_Mech-menapis-saturata_Pan-WEST
CAM008423_Mech-menapis-saturata_Pan-WEST
Sal_1938_Mech-menapis-dariensis_Col-WEST
Sal_4723_Mech-menapis-occasiva_Col-WEST
Sal_5681_Mech-menapis-occasiva_Col-WEST
Sal_5683_Mech-menapis-occasiva_Col-WEST
Sal_2064_Mech-menapis-caribensis_Col-WEST
Sal_2066_Mech-menapis-caribensis_Col-WEST
Sal_5638_Mech-menapis-menapis_Col-WEST
Sal_5640_Mech-menapis-caribensis_Col-WEST
Sal_5641_Mech-menapis-caribensis_Col-WEST
Sal_3342_Mech-menapis-dariensis_Col-WEST
CAM070759_Mech-menapis-mantineus_Ec-WEST
SAN25001771_Mech-menapis-mantineus_Ec-WEST
SAN25001770_Mech-menapis-mantineus_Ec-WEST
SAN25001773_Mech-menapis-mantineus_Ec-WEST
SAN25001772_Mech-menapis-mantineus_Ec-WEST
Sal_1532_Mech-messenoides-deceptus_Col-WA
Sal_1360_Mech-messenoides-deceptus_Col-WA
Mon_2012_198_Mech-messenoides-deceptus_Ec-WA
Mon_2012_146_Mech-messenoides-deceptus_Ec-WA
Mon_2012_242_Mech-messenoides-messenoides_Ec-WA
Mon_2011_124_Mech-messenoides-deceptus_Ec-WA
Mon_2012_99_Mech-messenoides-messenoides_Ec-WA
Mon_2012_220_Mech-messenoides-messenoides_Ec-WA
Mon_2011_195_Mech-messenoides-deceptus_Ec-WA
Mon_2012_243_Mech-messenoides-deceptus_Ec-WA
Mon_2012_90_Mech-messenoides-messenoides_Ec-WA
55.02.1284_Mech-messenoides-phasianita_Peru-WA
CAM077084_Mech-messenoides-messenoides_Col-WA
61.02.1896_Mech-messenoides-phasianita_Peru-WA
CAM077080_Mech-messenoides-deceptus_Col-WA
60.02.1290_Mech-messenoides-phasianita_Peru-WA
CAM077079_Mech-messenoides-deceptus_Col-WA
Mon_2012_244_Mech-messenoides-messenoides_Ec-WA
CAM077081_Mech-messenoides-messenoides_Col-WA
CAM077091_Mech-messenoides-messenoides_Col-WA
CAM077045_Mech-messenoides-messenoides_Col-WA
CAM077092_Mech-messenoides-messenoides_Col-WA
CAM077083_Mech-messenoides-messenoides_Col-WA
Mon_2012_219_Mech-messenoides-mess-dec_Ec-WA
65.02.477_Mech-messenoides-phasianita_Peru-WA
Mon_2012_88_Mech-messenoides-deceptus_Ec-WA
50.02.348_Mech-messenoides-phasianita_Peru-WA
68.02.677_Mech-messenoides-phasianita_Peru-WA
CAM028090_For-equicola-equicoloides_Ec-WA
CAM028271_For-equicola-equicoloides_Ec-WA
CAM028327_For-equicola-equicoloides_Ec-WA
CAM028193_For-olivencia-juntana_Ec-WA
CAM070582_For-olivencia-juntana_Ec-WA
CAM028312_For-olivencia-juntana_Ec-WA
CAM070479_For-procera_Ec-WA
CAM070480_For-procera_Ec-WA
CAM070483_For-procera_Ec-WA

**Fig. S1. Nuclear phylogeny for *Mechanitis***

*Each individual is labelled with 'ID_Species_Country-Region'. The countries are: Pan = Panama; Ec = Ecuador; Col = Colombia; Peru = Peru; FG = French Guiana; Braz = Brazil; Sur = Suriname. The regions are: WEST = west of the Andes; WA = Western Amazonia; GUI = Guianan Shield; AF = Atlantic Forest. Numbers at the node indicate bootstrap support.*

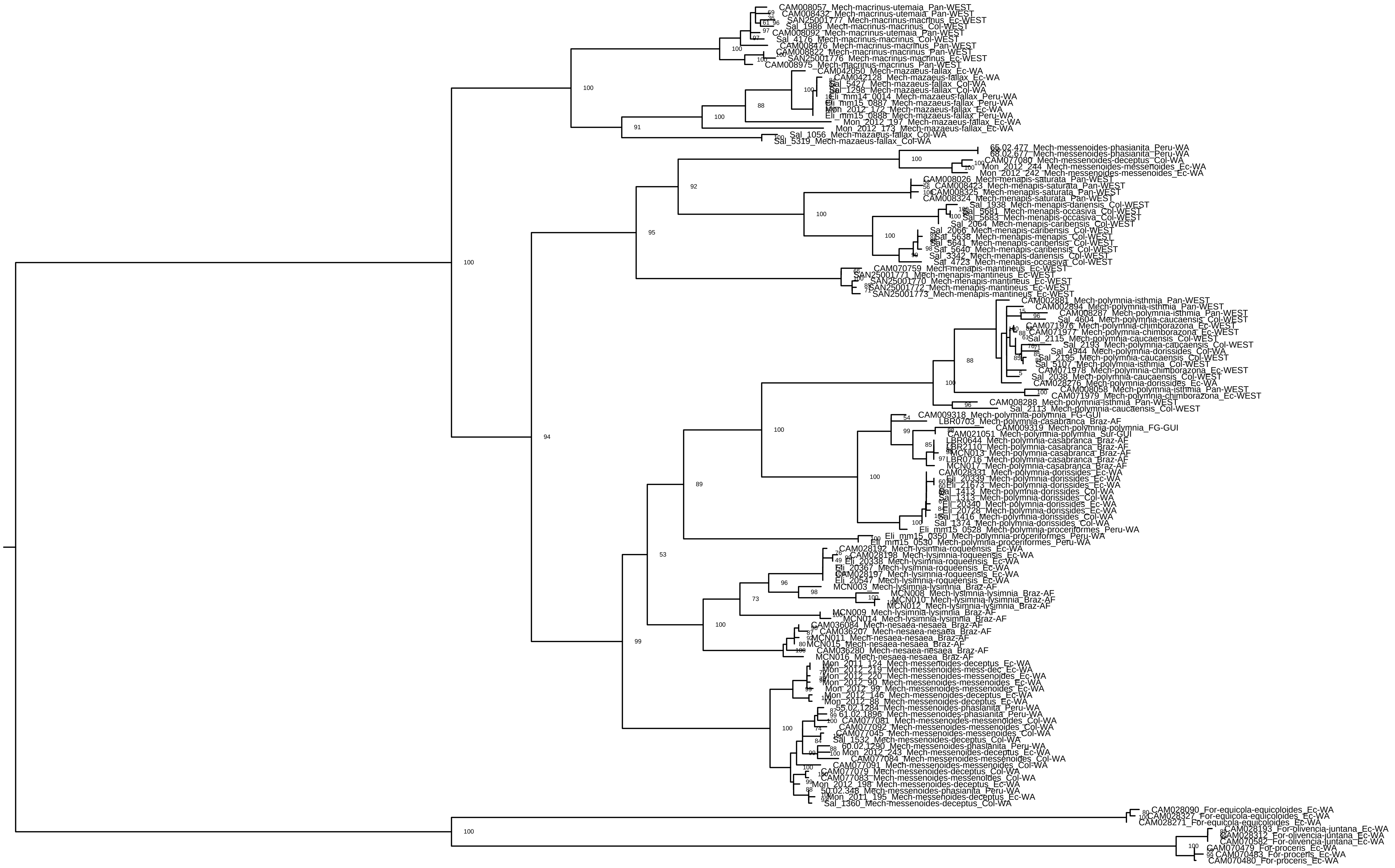
CAM008057_Mech-macrinus-utemaia_Pan-WEST
CAM008432_Mech-macrinus-utemaia_Pan-WEST
SAN25001777_Mech-macrinus-macrinus_Ec-WEST
Sal_1986_Mech-macrinus-macrinus_Col-WEST
CAM008092_Mech-macrinus-utemaia_Pan-WEST
Sal_4176_Mech-macrinus-macrinus_Col-WEST
CAM008476_Mech-macrinus-macrinus_Pan-WEST
CAM008822_Mech-macrinus-macrinus_Pan-WEST
SAN25001776_Mech-macrinus-macrinus_Ec-WEST
CAM008975_Mech-macrinus-macrinus_Pan-WEST
CAM042050_Mech-mazaeus-fallax_Ec-WA
CAM042128_Mech-mazaeus-fallax_Ec-WA
Sal_5427_Mech-mazaeus-fallax_Col-WA
Sal_1298_Mech-mazaeus-fallax_Col-WA
Eli_mm14_0014_Mech-mazaeus-fallax_Peru-WA
Eli_mm15_0887_Mech-mazaeus-fallax_Peru-WA
Mon_2012_172_Mech-mazaeus-fallax_Ec-WA
Eli_mm15_0888_Mech-mazaeus-fallax_Peru-WA
Mon_2012_197_Mech-mazaeus-fallax_Ec-WA
Mon_2012_173_Mech-mazaeus-fallax_Ec-WA
Sal_1056_Mech-mazaeus-fallax_Col-WA
Sal_5319_Mech-mazaeus-fallax_Col-WA
65.02.477_Mech-messenoides-phasianita_Peru-WA
68.02.677_Mech-messenoides-phasianita_Peru-WA
CAM077080_Mech-messenoides-deceptus_Col-WA
Mon_2012_244_Mech-messenoides-messenoides_Ec-WA
Mon_2012_242_Mech-messenoides-messenoides_Ec-WA
CAM008026_Mech-menapis-saturata_Pan-WEST
CAM008423_Mech-menapis-saturata_Pan-WEST
CAM008325_Mech-menapis-saturata_Pan-WEST
CAM008324_Mech-menapis-saturata_Pan-WEST
Sal_1938_Mech-menapis-dariensis_Col-WEST
Sal_5681_Mech-menapis-occasiva_Col-WEST
Sal_5683_Mech-menapis-occasiva_Col-WEST
Sal_2064_Mech-menapis-caribensis_Col-WEST
Sal_2066_Mech-menapis-caribensis_Col-WEST
Sal_5638_Mech-menapis-menapis_Col-WEST
Sal_5641_Mech-menapis-caribensis_Col-WEST
Sal_5640_Mech-menapis-caribensis_Col-WEST
Sal_3342_Mech-menapis-dariensis_Col-WEST
Sal_4723_Mech-menapis-occasiva_Col-WEST
CAM070759_Mech-menapis-mantineus_Ec-WEST
SAN25001771_Mech-menapis-mantineus_Ec-WEST
SAN25001770_Mech-menapis-mantineus_Ec-WEST
SAN25001772_Mech-menapis-mantineus_Ec-WEST
SAN25001773_Mech-menapis-mantineus_Ec-WEST
CAM002881_Mech-polymnia-isthmia_Pan-WEST
CAM002894_Mech-polymnia-isthmia_Pan-WEST
CAM008287_Mech-polymnia-isthmia_Pan-WEST
Sal_4604_Mech-polymnia-caucaensis_Col-WEST
CAM071976_Mech-polymnia-chimborazona_Ec-WEST
CAM071977_Mech-polymnia-chimborazona_Ec-WEST
Sal_2115_Mech-polymnia-caucaensis_Col-WEST
Sal_2193_Mech-polymnia-caucaensis_Col-WEST
Sal_4944_Mech-polymnia-dorissides_Col-WA
Sal_2195_Mech-polymnia-caucaensis_Col-WEST
Sal_5107_Mech-polymnia-isthmia_Col-WEST
CAM071978_Mech-polymnia-chimborazona_Ec-WEST
Sal_2038_Mech-polymnia-caucaensis_Col-WEST
CAM028276_Mech-polymnia-dorissides_Ec-WA
CAM008058_Mech-polymnia-isthmia_Pan-WEST
CAM071979_Mech-polymnia-chimborazona_Ec-WEST
CAM008288_Mech-polymnia-isthmia_Pan-WEST
Sal_2113_Mech-polymnia-caucaensis_Col-WEST
CAM009318_Mech-polymnia-polymnia_FG-GUI
LBR0703_Mech-polymnia-casabranca_Braz-AF
CAM009319_Mech-polymnia-polymnia_FG-GUI
CAM021051_Mech-polymnia-polymnia_Sur-GUI
LBR0644_Mech-polymnia-casabranca_Braz-AF
LBR2110_Mech-polymnia-casabranca_Braz-AF
MCN013_Mech-polymnia-casabranca_Braz-AF
LBR0716_Mech-polymnia-casabranca_Braz-AF
MCN017_Mech-polymnia-casabranca_Braz-AF
CAM028331_Mech-polymnia-dorissides_Ec-WA
Eli_20339_Mech-polymnia-dorissides_Ec-WA
Eli_21673_Mech-polymnia-dorissides_Ec-WA
Sal_1413_Mech-polymnia-dorissides_Col-WA
Sal_1313_Mech-polymnia-dorissides_Col-WA
Eli_20340_Mech-polymnia-dorissides_Ec-WA
Eli_20728_Mech-polymnia-dorissides_Ec-WA
Sal_1416_Mech-polymnia-dorissides_Col-WA
Sal_1374_Mech-polymnia-dorissides_Col-WA
Eli_mm15_0528_Mech-polymnia-proceriformes_Peru-WA
Eli_mm15_0350_Mech-polymnia-proceriformes_Peru-WA
Eli_mm15_0530_Mech-polymnia-proceriformes_Peru-WA
CAM028192_Mech-lysimnia-roqueensis_Ec-WA
CAM028198_Mech-lysimnia-roqueensis_Ec-WA
Eli_20338_Mech-lysimnia-roqueensis_Ec-WA
Eli_20367_Mech-lysimnia-roqueensis_Ec-WA
CAM028197_Mech-lysimnia-roqueensis_Ec-WA
Eli_20547_Mech-lysimnia-roqueensis_Ec-WA
MCN003_Mech-lysimnia-lysimnia_Braz-AF
MCN008_Mech-lysimnia-lysimnia_Braz-AF
MCN010_Mech-lysimnia-lysimnia_Braz-AF
MCN012_Mech-lysimnia-lysimnia_Braz-AF
MCN009_Mech-lysimnia-lysimnia_Braz-AF
MCN014_Mech-lysimnia-lysimnia_Braz-AF
CAM036084_Mech-nesaea-nesaea_Braz-AF
CAM036207_Mech-nesaea-nesaea_Braz-AF
MCN011_Mech-nesaea-nesaea_Braz-AF
MCN015_Mech-nesaea-nesaea_Braz-AF
CAM036280_Mech-nesaea-nesaea_Braz-AF
MCN016_Mech-nesaea-nesaea_Braz-AF
Mon_2011_124_Mech-messenoides-deceptus_Ec-WA
Mon_2012_219_Mech-messenoides-mess-dec_Ec-WA
Mon_2012_220_Mech-messenoides-messenoides_Ec-WA
Mon_2012_90_Mech-messenoides-messenoides_Ec-WA
Mon_2012_99_Mech-messenoides-messenoides_Ec-WA
Mon_2012_146_Mech-messenoides-deceptus_Ec-WA
Mon_2012_88_Mech-messenoides-deceptus_Ec-WA
55.02.1284_Mech-messenoides-phasianita_Peru-WA
61.02.1896_Mech-messenoides-phasianita_Peru-WA
CAM077081_Mech-messenoides-messenoides_Col-WA
CAM077092_Mech-messenoides-messenoides_Col-WA
CAM077045_Mech-messenoides-messenoides_Col-WA
Sal_1532_Mech-messenoides-deceptus_Col-WA
60.02.1290_Mech-messenoides-phasianita_Peru-WA
Mon_2012_243_Mech-messenoides-deceptus_Ec-WA
CAM077084_Mech-messenoides-messenoides_Col-WA
CAM077091_Mech-messenoides-messenoides_Col-WA
CAM077079_Mech-messenoides-deceptus_Col-WA
CAM077083_Mech-messenoides-messenoides_Col-WA
Mon_2012_198_Mech-messenoides-deceptus_Ec-WA
50.02.348_Mech-messenoides-phasianita_Peru-WA
Mon_2011_195_Mech-messenoides-deceptus_Ec-WA
Sal_1360_Mech-messenoides-deceptus_Col-WA
CAM028090_For-equicola-equicoloides_Ec-WA
CAM028327_For-equicola-equicoloides_Ec-WA
CAM028271_For-equicola-equicoloides_Ec-WA
CAM028193_For-olivencia-juntana_Ec-WA
CAM028312_For-olivencia-juntana_Ec-WA
CAM070582_For-olivencia-juntana_Ec-WA
CAM070479_For-proceris_Ec-WA
CAM070483_For-proceris_Ec-WA
CAM070480_For-proceris_Ec-WA

**Fig. S2. Mitochondrial phylogeny for *Mechanitis***

*Each individual is labelled with 'ID_Species_Country-Region'. The countries are: Pan = Panama; Ec = Ecuador; Col = Colombia; Peru = Peru; FG = French Guiana; Braz = Brazil; Sur = Suriname. The regions are: WEST = west of the Andes; WA = Western Amazonia; GUI = Guianan Shield; AF = Atlantic Forest. Numbers at the node indicate bootstrap support.*

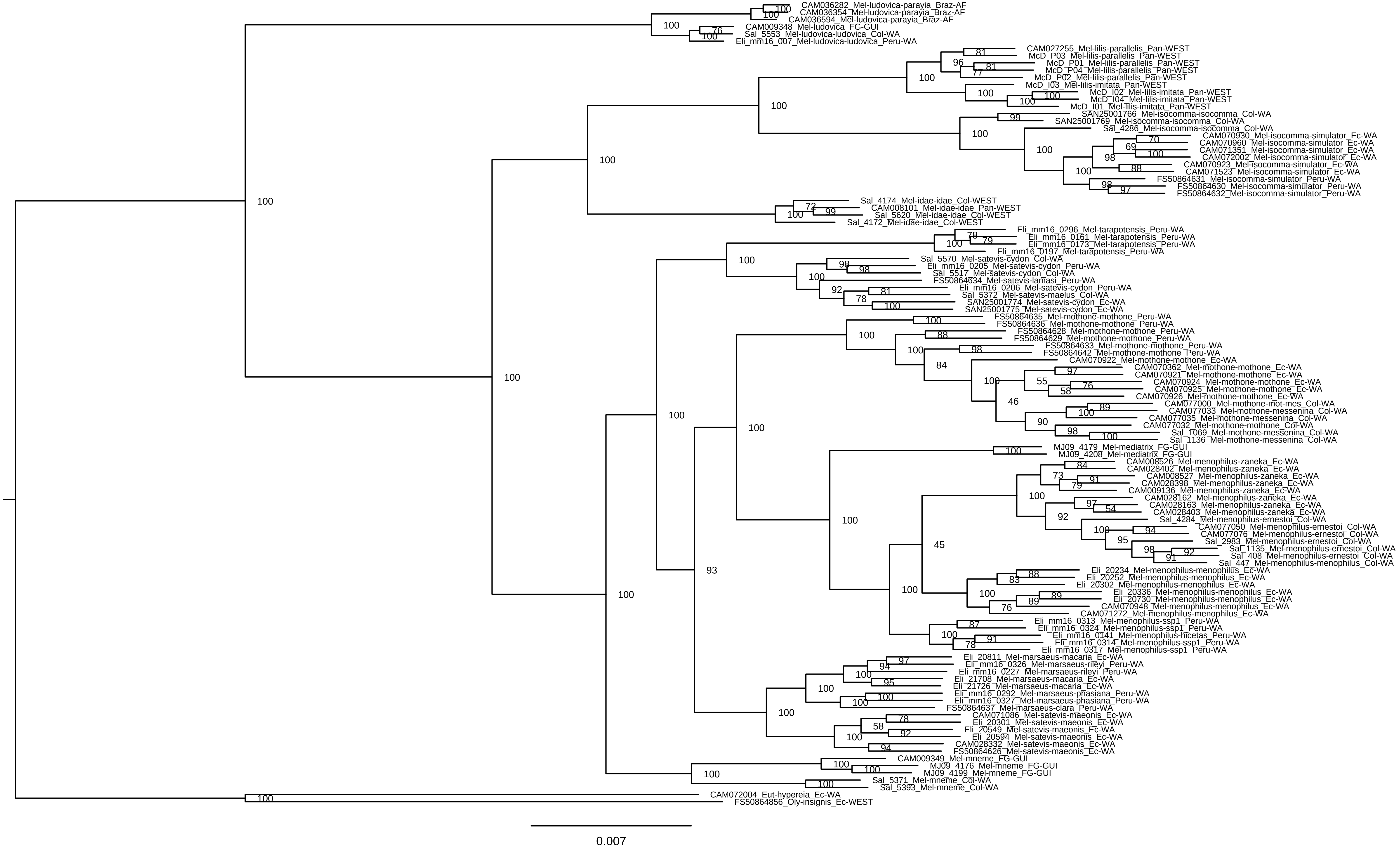
CAM036282_Mel-ludovica-parayia_Braz-AF
CAM036354_Mel-ludovica-parayia_Braz-AF
CAM036594_Mel-ludovica-parayia_Braz-AF
CAM009348_Mel-ludovica_FG-GUI
Sal_5553_Mel-ludovica-ludovica_Col-WA
Eli_mm16_007_Mel-ludovica-ludovica_Peru-WA
CAM027255_Mel-lilis-parallelis_Pan-WEST
McD_P03_Mel-lilis-parallelis_Pan-WEST
McD_P01_Mel-lilis-parallelis_Pan-WEST
McD_P04_Mel-lilis-parallelis_Pan-WEST
McD_P02_Mel-lilis-parallelis_Pan-WEST
McD_I03_Mel-lilis-imitata_Pan-WEST
McD_I02_Mel-lilis-imitata_Pan-WEST
McD_I04_Mel-lilis-imitata_Pan-WEST
McD_I01_Mel-lilis-imitata_Pan-WEST
SAN25001766_Mel-isocomma-isocomma_Col-WA
SAN25001769_Mel-isocomma-isocomma_Col-WA
Sal_4286_Mel-isocomma-isocomma_Col-WA
CAM070930_Mel-isocomma-simulator_Ec-WA
CAM070960_Mel-isocomma-simulator_Ec-WA
CAM071351_Mel-isocomma-simulator_Ec-WA
CAM072002_Mel-isocomma-simulator_Ec-WA
CAM070923_Mel-isocomma-simulator_Ec-WA
CAM071523_Mel-isocomma-simulator_Ec-WA
FS50864631_Mel-isocomma-simulator_Peru-WA
FS50864630_Mel-isocomma-simulator_Peru-WA
FS50864632_Mel-isocomma-simulator_Peru-WA
Sal_4174_Mel-idae-idae_Col-WEST
CAM008101_Mel-idae-idae_Pan-WEST
Sal_5620_Mel-idae-idae_Col-WEST
Sal_4172_Mel-idae-idae_Col-WEST
Eli_mm16_0296_Mel-tarapotensis_Peru-WA
Eli_mm16_0161_Mel-tarapotensis_Peru-WA
Eli_mm16_0173_Mel-tarapotensis_Peru-WA
Eli_mm16_0197_Mel-tarapotensis_Peru-WA
Sal_5570_Mel-satevis-cydon_Col-WA
Eli_mm16_0205_Mel-satevis-cydon_Peru-WA
Sal_5517_Mel-satevis-cydon_Col-WA
FS50864634_Mel-satevis-lamasi_Peru-WA
Eli_mm16_0206_Mel-satevis-cydon_Peru-WA
Sal_5372_Mel-satevis-maelus_Col-WA
SAN25001774_Mel-satevis-cydon_Ec-WA
SAN25001775_Mel-satevis-cydon_Ec-WA
FS50864635_Mel-mothone-mothone_Peru-WA
FS50864636_Mel-mothone-mothone_Peru-WA
FS50864628_Mel-mothone-mothone_Peru-WA
FS50864629_Mel-mothone-mothone_Peru-WA
FS50864633_Mel-mothone-mothone_Peru-WA
FS50864642_Mel-mothone-mothone_Peru-WA
CAM070922_Mel-mothone-mothone_Ec-WA
CAM070362_Mel-mothone-mothone_Ec-WA
CAM070921_Mel-mothone-mothone_Ec-WA
CAM070924_Mel-mothone-mothone_Ec-WA
CAM070925_Mel-mothone-mothone_Ec-WA
CAM070926_Mel-mothone-mothone_Ec-WA
CAM077000_Mel-mothone-mot-mes_Col-WA
CAM077033_Mel-mothone-messenina_Col-WA
CAM077035_Mel-mothone-messenina_Col-WA
CAM077032_Mel-mothone-mothone_Col-WA
Sal_1069_Mel-mothone-messenina_Col-WA
Sal_1136_Mel-mothone-messenina_Col-WA
MJ09_4179_Mel-mediatrix_FG-GUI
MJ09_4208_Mel-mediatrix_FG-GUI
CAM008526_Mel-menophilus-zaneka_Ec-WA
CAM028402_Mel-menophilus-zaneka_Ec-WA
CAM008527_Mel-menophilus-zaneka_Ec-WA
CAM028398_Mel-menophilus-zaneka_Ec-WA
CAM009136_Mel-menophilus-zaneka_Ec-WA
CAM028162_Mel-menophilus-zaneka_Ec-WA
CAM028163_Mel-menophilus-zaneka_Ec-WA
CAM028403_Mel-menophilus-zaneka_Ec-WA
Sal_4284_Mel-menophilus-ernestoi_Col-WA
CAM077050_Mel-menophilus-ernestoi_Col-WA
CAM077076_Mel-menophilus-ernestoi_Col-WA
Sal_2983_Mel-menophilus-ernestoi_Col-WA
Sal_1135_Mel-menophilus-ernestoi_Col-WA
Sal_408_Mel-menophilus-ernestoi_Col-WA
Sal_447_Mel-menophilus-ernestoi_Col-WA
Eli_20234_Mel-menophilus-menophilus_Ec-WA
Eli_20252_Mel-menophilus-menophilus_Ec-WA
Eli_20302_Mel-menophilus-menophilus_Ec-WA
Eli_20336_Mel-menophilus-menophilus_Ec-WA
Eli_20730_Mel-menophilus-menophilus_Ec-WA
CAM070948_Mel-menophilus-menophilus_Ec-WA
CAM071272_Mel-menophilus-menophilus_Ec-WA
Eli_mm16_0313_Mel-menophilus-ssp1_Peru-WA
Eli_mm16_0324_Mel-menophilus-ssp1_Peru-WA
Eli_mm16_0141_Mel-menophilus-hicetas_Peru-WA
Eli_mm16_0314_Mel-menophilus-ssp1_Peru-WA
Eli_mm16_0317_Mel-menophilus-ssp1_Peru-WA
Eli_20811_Mel-marsaeus-macaria_Ec-WA
Eli_mm16_0326_Mel-marsaeus-rileyi_Peru-WA
Eli_mm16_0227_Mel-marsaeus-rileyi_Peru-WA
Eli_21708_Mel-marsaeus-macaria_Ec-WA
Eli_21726_Mel-marsaeus-macaria_Ec-WA
Eli_mm16_0292_Mel-marsaeus-phasiana_Peru-WA
Eli_mm16_0327_Mel-marsaeus-phasiana_Peru-WA
FS50864637_Mel-marsaeus-clara_Peru-WA
CAM071086_Mel-satevis-maeonis_Ec-WA
Eli_20301_Mel-satevis-maeonis_Ec-WA
Eli_20549_Mel-satevis-maeonis_Ec-WA
Eli_20594_Mel-satevis-maeonis_Ec-WA
CAM028332_Mel-satevis-maeonis_Ec-WA
FS50864626_Mel-satevis-maeonis_Ec-WA
CAM009349_Mel-mneme_FG-GUI
MJ09_4176_Mel-mneme_FG-GUI
MJ09_4199_Mel-mneme_FG-GUI
Sal_5371_Mel-mneme_Col-WA
Sal_5393_Mel-mneme_Col-WA
CAM072004_Eut-hypereia_Ec-WA
FS50864856_Oly-insignis_Ec-WEST
0.007

**Fig. S3. Nuclear phylogeny for *Melinaea***

*Each individual is labelled with 'ID_Species_Country-Region'. The countries are: Pan = Panama; Ec = Ecuador; Col = Colombia; Peru = Peru; FG = French Guiana; Braz = Brazil; Sur = Suriname. The regions are: WEST = west of the Andes; WA = Western Amazonia; GUI = Guianan Shield; AF = Atlantic Forest. Numbers at the node indicate bootstrap support.*

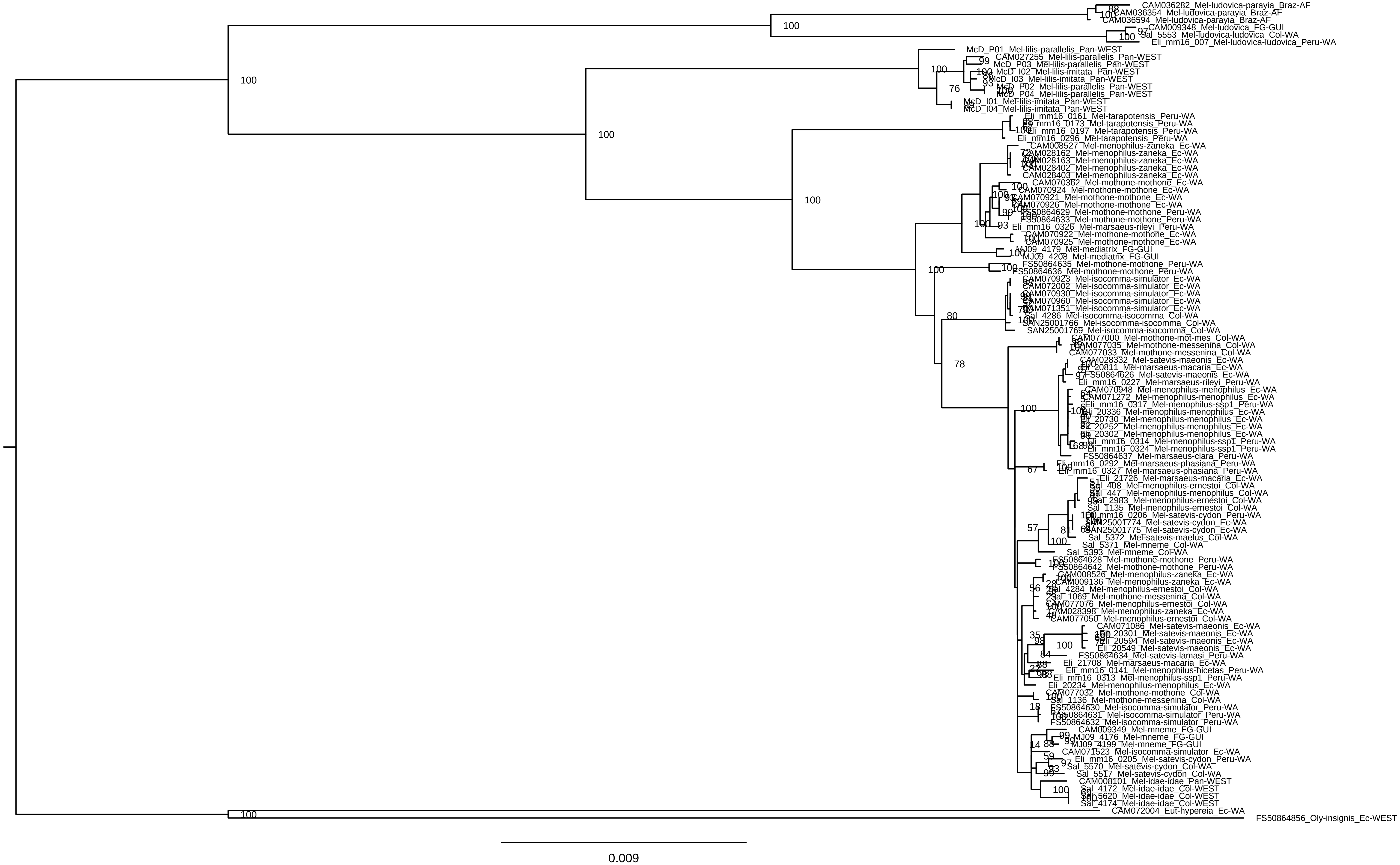

CAM036282_Mel-ludovica-parayia_Braz-AF
CAM036354_Mel-ludovica-parayia_Braz-AF
CAM036594_Mel-ludovica-parayia_Braz-AF
CAM009348_Mel-ludovica_FG-GUI
Sal_5553_Mel-ludovica-ludovica_Col-WA
Eli_mm16_007_Mel-ludovica-ludovica_Peru-WA
McD_P01_Mel-lilis-parallelis_Pan-WEST
CAM027255_Mel-lilis-parallelis_Pan-WEST
McD_P03_Mel-lilis-parallelis_Pan-WEST
McD_I02_Mel-lilis-imitata_Pan-WEST
McD_I03_Mel-lilis-imitata_Pan-WEST
McD_P02_Mel-lilis-parallelis_Pan-WEST
McD_P04_Mel-lilis-parallelis_Pan-WEST
McD_I01_Mel-lilis-imitata_Pan-WEST
McD_I04_Mel-lilis-imitata_Pan-WEST
Eli_mm16_0161_Mel-tarapotensis_Peru-WA
Eli_mm16_0173_Mel-tarapotensis_Peru-WA
Eli_mm16_0197_Mel-tarapotensis_Peru-WA
Eli_mm16_0296_Mel-tarapotensis_Peru-WA
CAM008527_Mel-menophilus-zaneka_Ec-WA
CAM028162_Mel-menophilus-zaneka_Ec-WA
CAM028163_Mel-menophilus-zaneka_Ec-WA
CAM028402_Mel-menophilus-zaneka_Ec-WA
CAM028403_Mel-menophilus-zaneka_Ec-WA
CAM070362_Mel-mothone-mothone_Ec-WA
CAM070924_Mel-mothone-mothone_Ec-WA
CAM070921_Mel-mothone-mothone_Ec-WA
CAM070926_Mel-mothone-mothone_Ec-WA
FS50864629_Mel-mothone-mothone_Peru-WA
FS50864633_Mel-mothone-mothone_Peru-WA
Eli_mm16_0326_Mel-marsaeus-rileyi_Peru-WA
CAM070922_Mel-mothone-mothone_Ec-WA
CAM070925_Mel-mothone-mothone_Ec-WA
MJ09_4179_Mel-mediatrix_FG-GUI
MJ09_4208_Mel-mediatrix_FG-GUI
FS50864635_Mel-mothone-mothone_Peru-WA
FS50864636_Mel-mothone-mothone_Peru-WA
CAM070923_Mel-isocomma-simulator_Ec-WA
CAM072002_Mel-isocomma-simulator_Ec-WA
CAM070930_Mel-isocomma-simulator_Ec-WA
CAM070960_Mel-isocomma-simulator_Ec-WA
CAM071351_Mel-isocomma-simulator_Ec-WA
Sal_4286_Mel-isocomma-isocomma_Col-WA
SAN25001766_Mel-isocomma-isocomma_Col-WA
SAN25001769_Mel-isocomma-isocomma_Col-WA
CAM077000_Mel-mothone-mot-mes_Col-WA
CAM077035_Mel-mothone-messenina_Col-WA
CAM077033_Mel-mothone-messenina_Col-WA
CAM028332_Mel-satevis-maeonis_Ec-WA
Eli_20811_Mel-marsaeus-macaria_Ec-WA
FS50864626_Mel-satevis-maeonis_Peru-WA
Eli_mm16_0227_Mel-marsaeus-rileyi_Peru-WA
CAM070948_Mel-menophilus-menophilus_Ec-WA
CAM071272_Mel-menophilus-menophilus_Ec-WA
Eli_mm16_0317_Mel-menophilus-ssp1_Peru-WA
Eli_20336_Mel-menophilus-menophilus_Ec-WA
Eli_20730_Mel-menophilus-menophilus_Ec-WA
Eli_20252_Mel-menophilus-menophilus_Ec-WA
Eli_20302_Mel-menophilus-menophilus_Ec-WA
Eli_mm16_0314_Mel-menophilus-ssp1_Peru-WA
Eli_mm16_0324_Mel-menophilus-ssp1_Peru-WA
FS50864637_Mel-marsaeus-clara_Peru-WA
Eli_mm16_0292_Mel-marsaeus-phasiana_Peru-WA
Eli_mm16_0327_Mel-marsaeus-phasiana_Peru-WA
Eli_21726_Mel-marsaeus-macaria_Ec-WA
Sal_408_Mel-menophilus-ernestoi_Col-WA
Sal_447_Mel-menophilus-menophilus_Col-WA
Sal_2983_Mel-menophilus-ernestoi_Col-WA
Sal_1135_Mel-menophilus-ernestoi_Col-WA
Eli_mm16_0206_Mel-satevis-cydon_Peru-WA
SAN25001774_Mel-satevis-cydon_Ec-WA
SAN25001775_Mel-satevis-cydon_Ec-WA
Sal_5372_Mel-satevis-maelus_Col-WA
Sal_5371_Mel-mneme_Col-WA
Sal_5393_Mel-mneme_Col-WA
FS50864628_Mel-mothone-mothone_Peru-WA
FS50864642_Mel-mothone-mothone_Peru-WA
CAM008526_Mel-menophilus-zaneka_Ec-WA
CAM009136_Mel-menophilus-zaneka_Ec-WA
Sal_4284_Mel-menophilus-ernestoi_Col-WA
Sal_1069_Mel-mothone-messenina_Col-WA
CAM077076_Mel-menophilus-ernestoi_Col-WA
CAM028398_Mel-menophilus-zaneka_Ec-WA
CAM077050_Mel-menophilus-ernestoi_Col-WA
CAM071086_Mel-satevis-maeonis_Ec-WA
Eli_20301_Mel-satevis-maeonis_Ec-WA
Eli_20594_Mel-satevis-maeonis_Ec-WA
Eli_20549_Mel-satevis-maeonis_Ec-WA
FS50864634_Mel-satevis-lamasi_Peru-WA
Eli_21708_Mel-marsaeus-macaria_Ec-WA
Eli_mm16_0141_Mel-menophilus-hicetas_Peru-WA
Eli_mm16_0313_Mel-menophilus-ssp1_Peru-WA
Eli_20234_Mel-menophilus-menophilus_Ec-WA
CAM077032_Mel-mothone-mothone_Col-WA
Sal_1136_Mel-mothone-messenina_Col-WA
FS50864630_Mel-isocomma-simulator_Peru-WA
FS50864631_Mel-isocomma-simulator_Peru-WA
FS50864632_Mel-isocomma-simulator_Peru-WA
CAM009349_Mel-mneme_FG-GUI
MJ09_4176_Mel-mneme_FG-GUI
MJ09_4199_Mel-mneme_FG-GUI
CAM071523_Mel-isocomma-simulator_Ec-WA
Eli_mm16_0205_Mel-satevis-cydon_Peru-WA
Sal_5570_Mel-satevis-cydon_Col-WA
Sal_5517_Mel-satevis-cydon_Col-WA
CAM008101_Mel-idae-idae_Pan-WEST
Sal_4172_Mel-idae-idae_Col-WEST
Sal_5620_Mel-idae-idae_Col-WEST
Sal_4174_Mel-idae-idae_Col-WEST
CAM072004_Eut-hypereia_Ec-WA
FS50864856_Oly-insignis_Ec-WEST
0.009

**Fig. S4. Mitochondrial phylogeny for *Melinaea***

*Each individual is labelled with ‘ID_Species_Country-Region’. The countries are: Pan = Panama; Ec = Ecuador; Col = Colombia; Peru = Peru; FG = French Guiana; Braz = Brazil; Sur = Suriname. The regions are: WEST = west of the Andes; WA = Western Amazonia; GUI = Guianan Shield; AF = Atlantic Forest. Numbers at the node indicate bootstrap support.*

A. *Mechanitis* – all species; each topology coloured differently

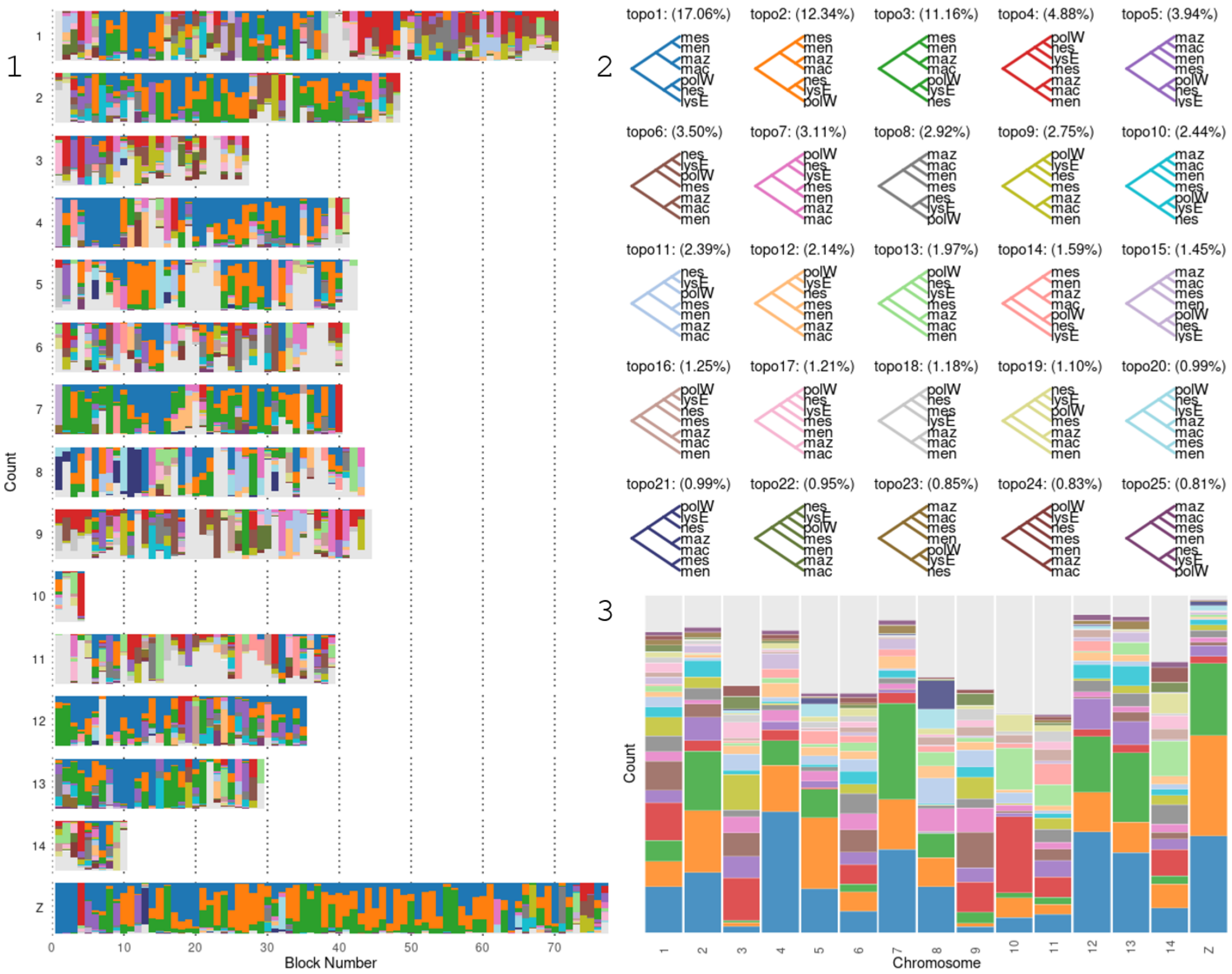

*B. Mechanitis* – all species; coloured by position of *Mec. messenoides*

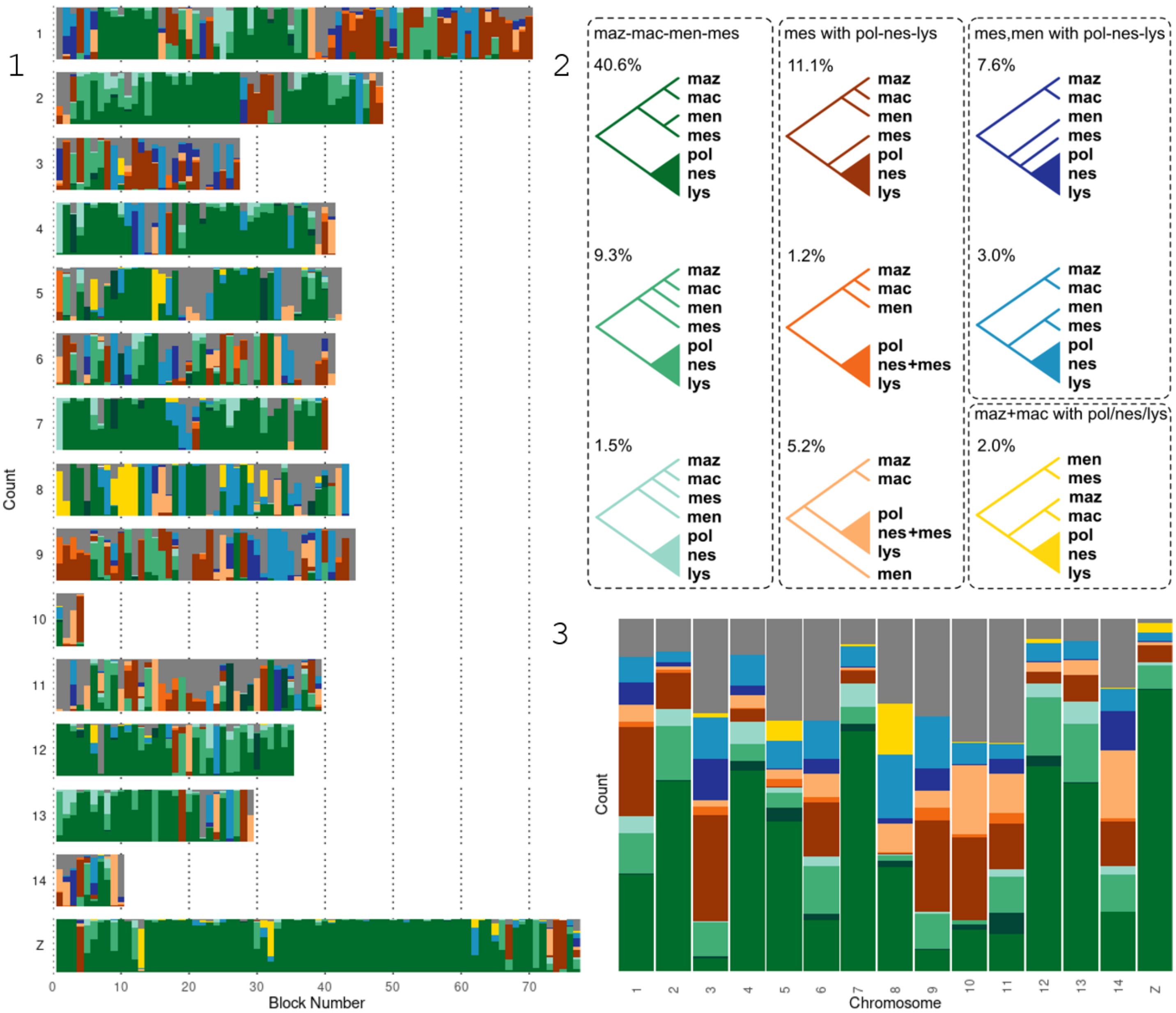

*C. Mechanitis* – all species; coloured by position of *Mec. nesaea*

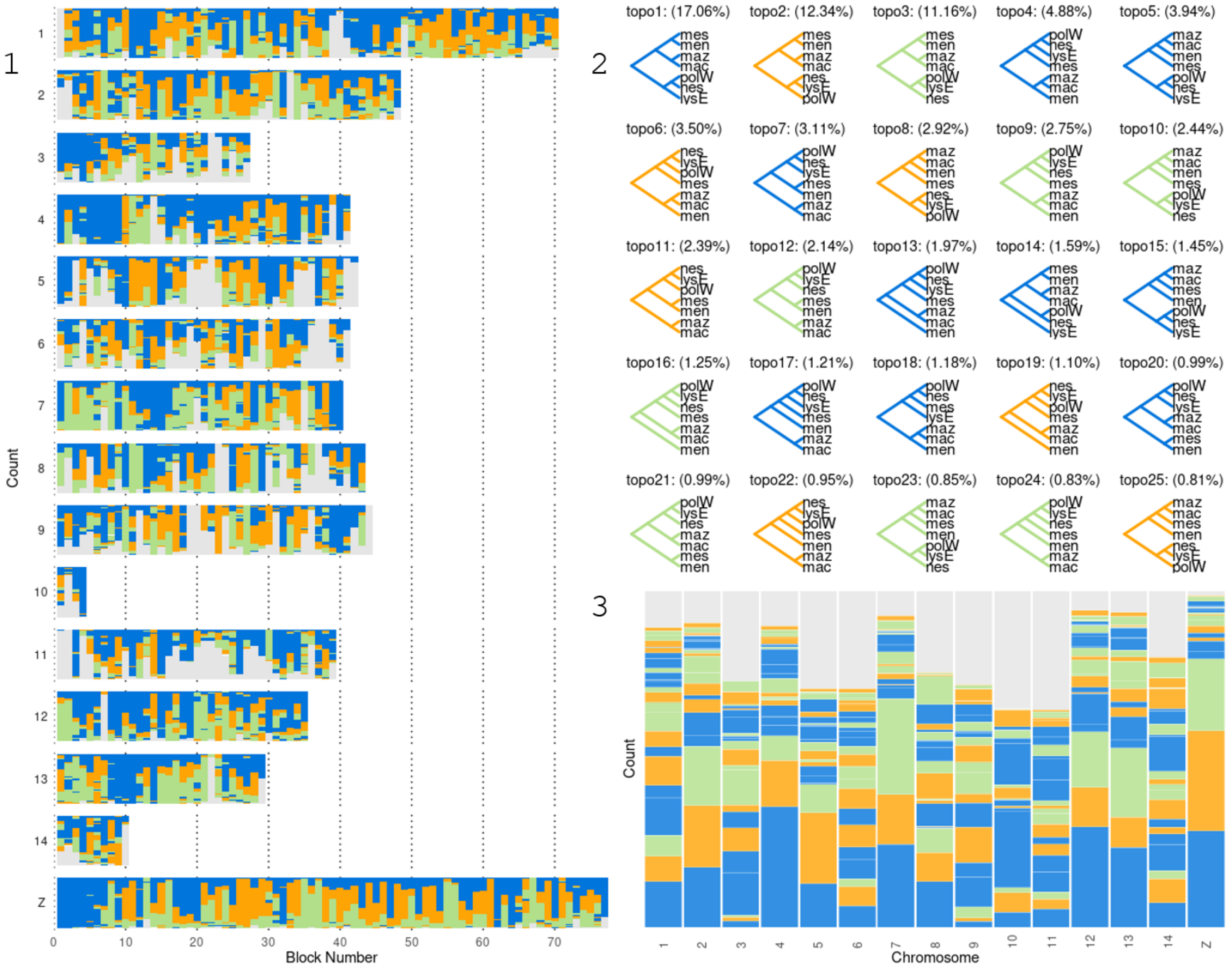

*D. Mechanitis – Mec. polymnia, Mec. nesaea, Mec. lysimnia*

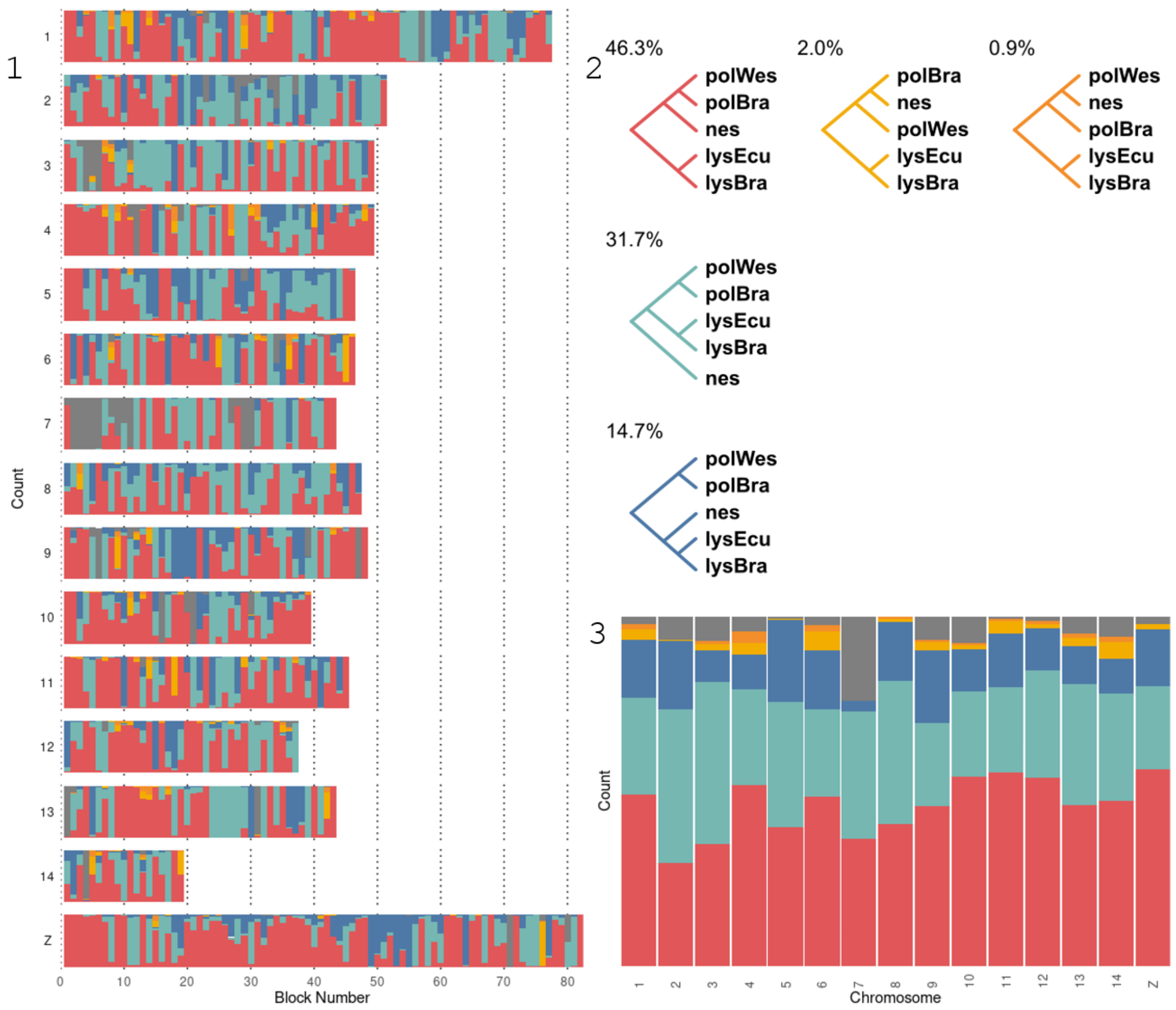

E. *Melinaea* – subset of species ; each topology coloured differently

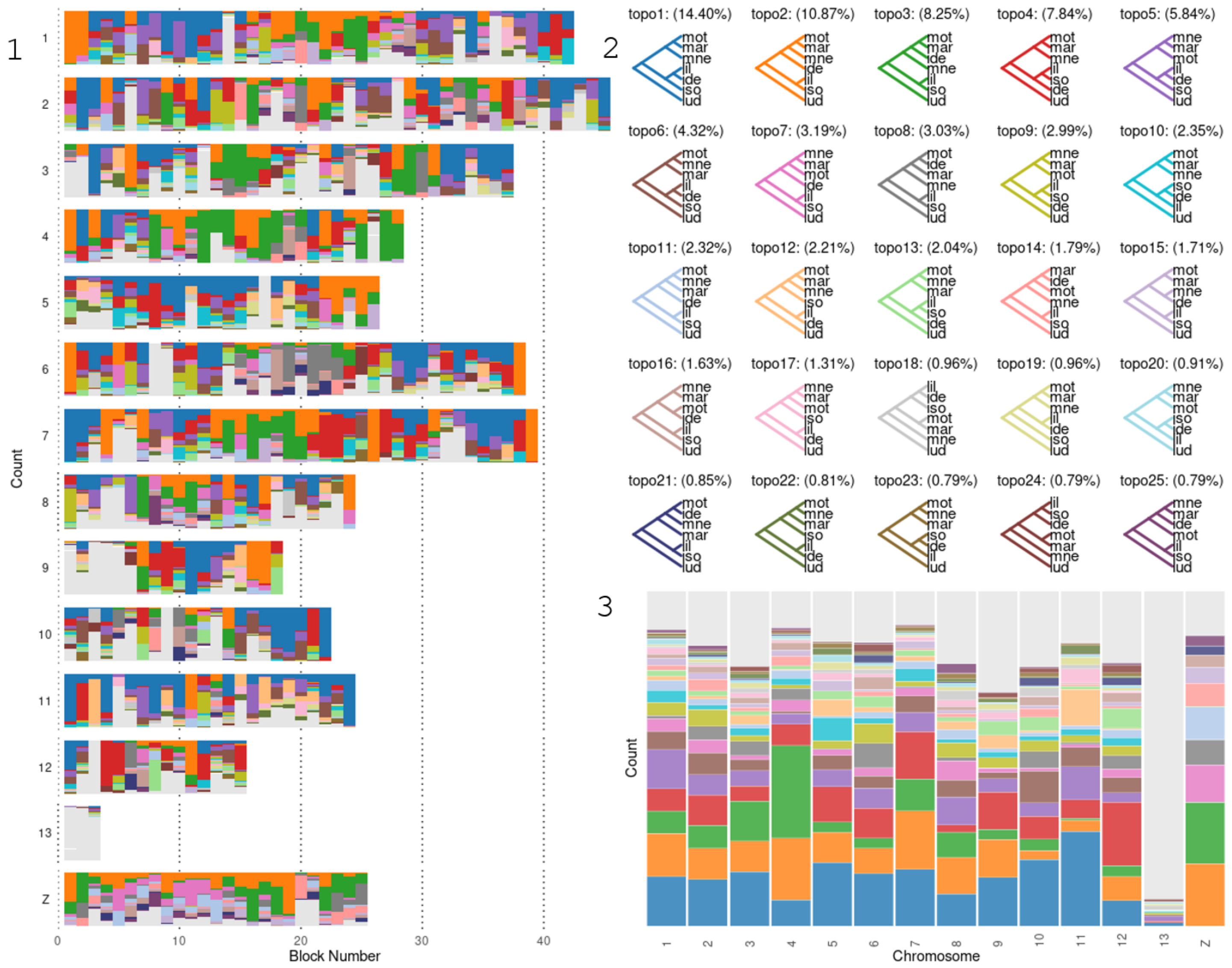

F. *Melinaea* – subset of species ; coloured by position of *Mel. idae*

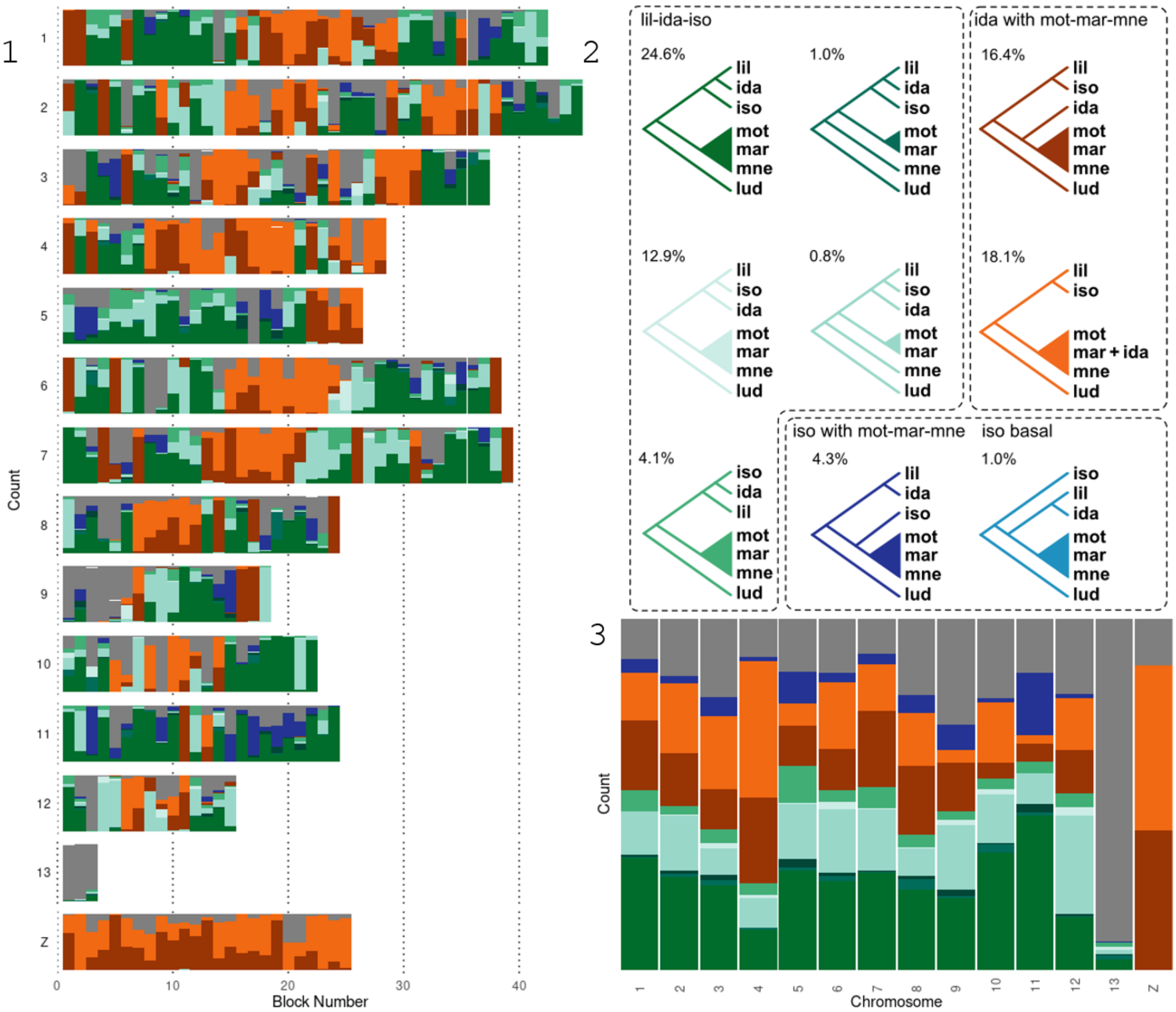


**Fig. S5. BPP analysis for *Mechanitis* and *Melinaea***

*BPP inferred tree topologies in windows across the genome. For each figure, part 1 shows one chromosome per row, with the windows across the chromosome coloured by the topologies for that window. Part 2 shows the various topologies, ordered by how often they occur or by an indicated grouping. The number above each topology indicates what percentage of the genome that topology occurs. Each column of part 3 represents a chromosome, coloured by how much each topology occurs in that chromosome. Grey colouration means that that window had a topology that occurred less than one of the top25 topologies. A, B and C show results for the analysis consisting of one individual of each* Mechanitis *species, with A coloured differently for each topology, B coloured according to the position of* Mec. messenoides*, and C coloured for the position of* Mec. nesaea*. mes = Mec. messenoides; men = Mec. menapis; maz = Mec. mazaeus; mac = Mec. macrinus; polW = Mec. polymnia (west); nes = Mec. nesaea; lysE = Mec. lysimnia (Ecuador). D is the result of an analysis including* Mec. polymnia *from Brazil (polBra) and west of the Andes (polWes),* Mec. nesaea *(Brazil; nes) and* Mec. lysimnia *from Brazil (lysBra) and Ecuador (lysEcu). E and F are the results for a subset of the* Melinaea *species, with E coloured differently for each topology, and F coloured by the placement of* Mel. idae*. mot = Mel. mothone; mar = Mel. marsaeus; mne = Mel. mneme; lil = Mel. lilis; ida = Mel. idae; iso = Mel. isocomma; lud = Mel. ludovica.*

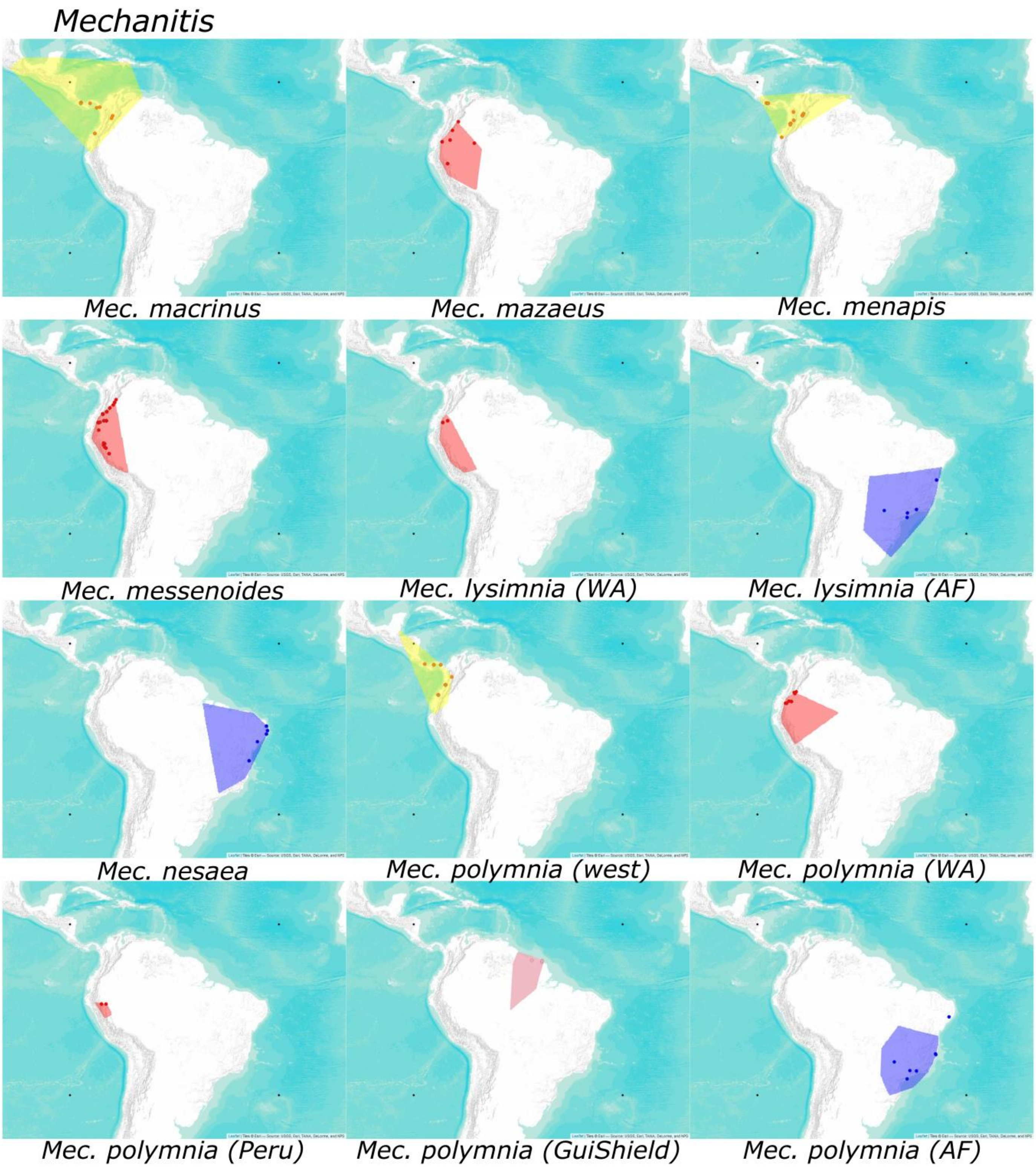


**Fig. S6. Distribution maps for *Mechanitis***

*Larger sizes of all the distribution maps in main figure 2 for Mechanitis. Concave shapes are based on distribution data from Doré et al (2022). Dots are our samples. Attribution for the maps is shown in the corner of each map (downloaded with 'Leaflet', map tiles from USGS, Esri, Tana, DeLorme, and NPS). The four black dots are an aid to crop the maps consistently for the main figure.*

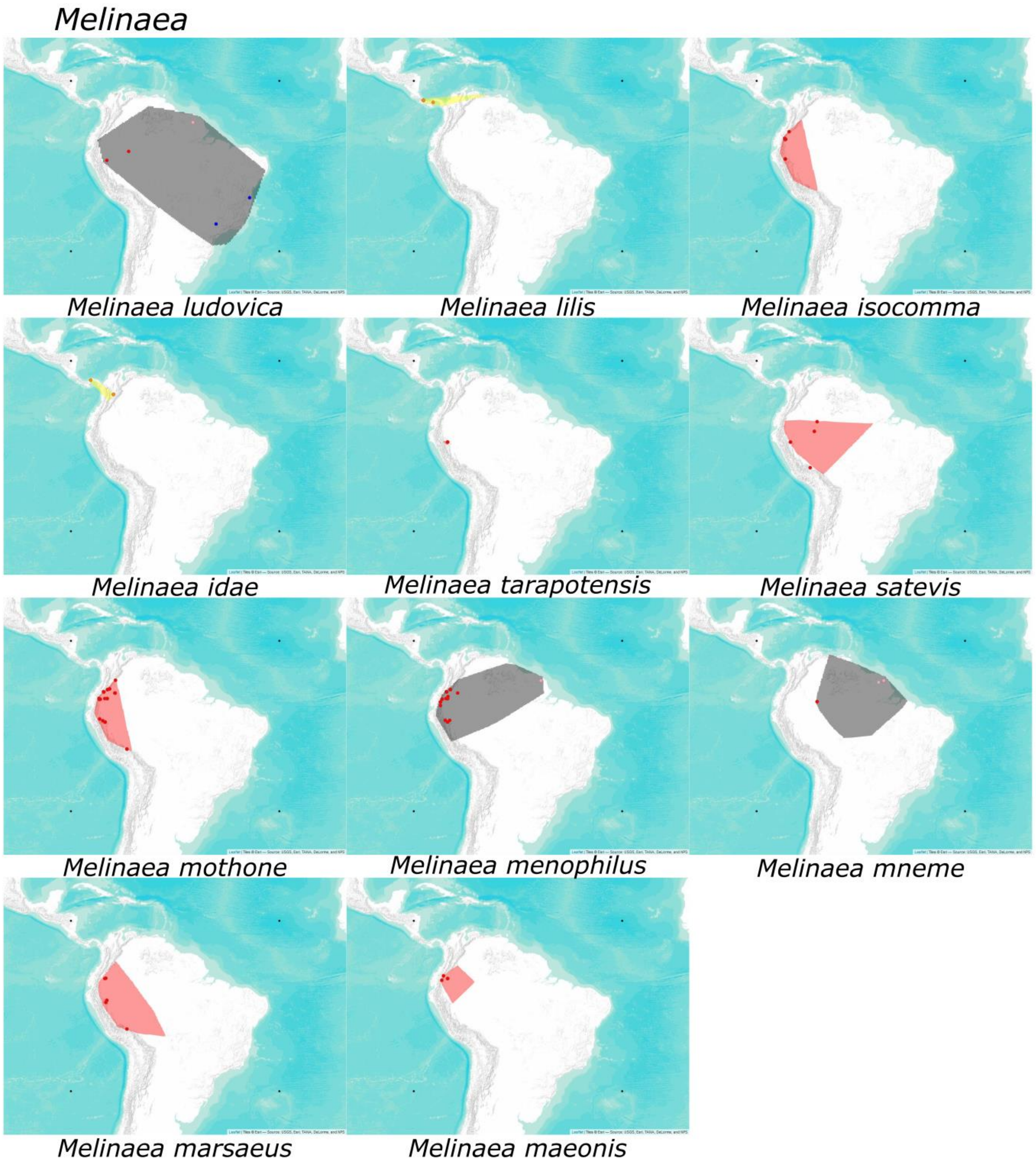


**Fig. S7. Distribution maps for *Melinaea***

*Larger sizes of all the distribution maps in main figure 2 for* Melinaea*. Concave shapes are based on distribution data from Doré et al (2022). Dots are our samples. Attribution for the maps is shown in the corner of each map (downloaded with 'Leaflet', map tiles from USGS, Esri, Tana, DeLorme, and NPS). The four black dots are an aid to crop the maps consistently for the main figure.*

A

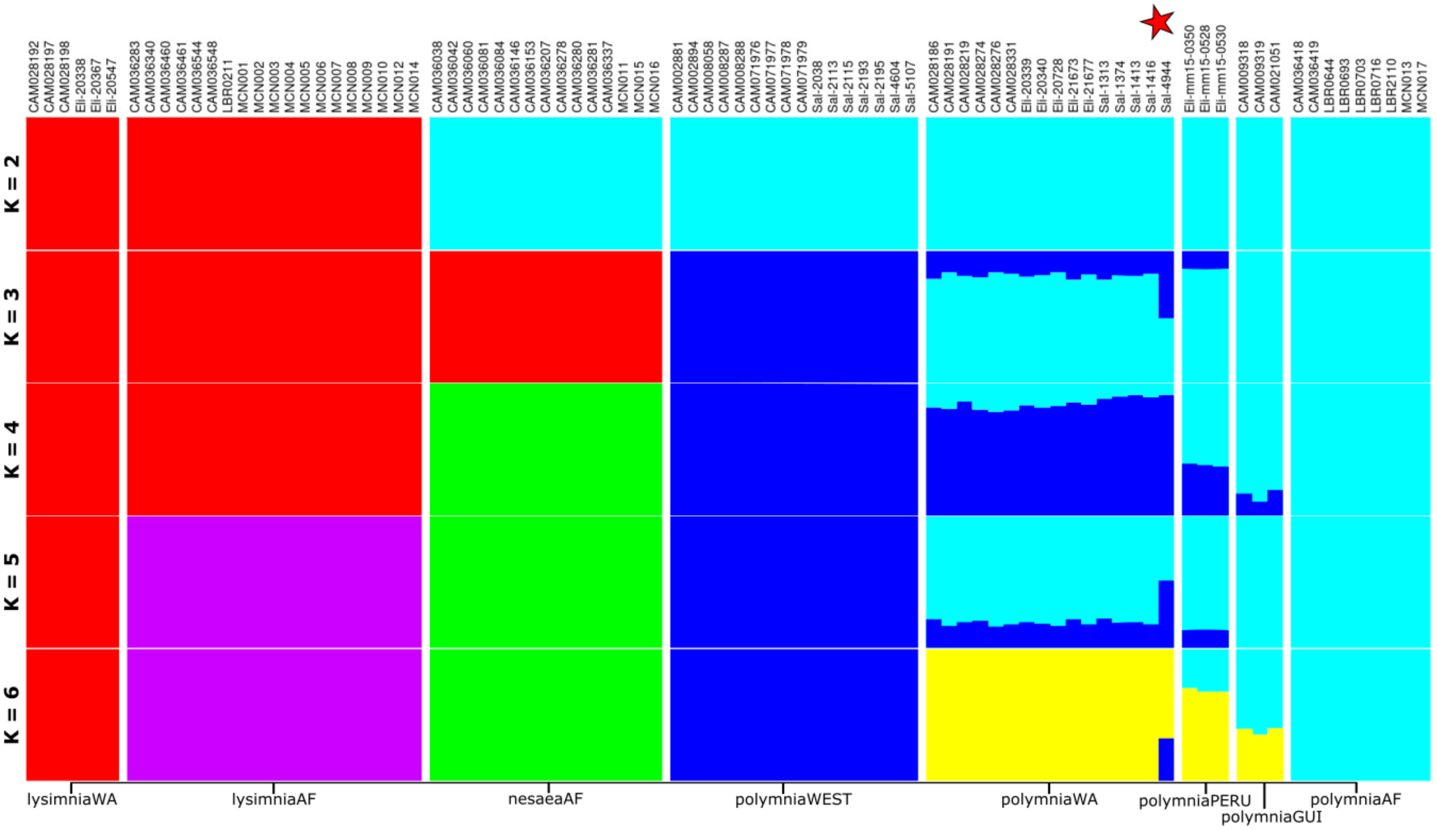
K = 2
K = 3
K = 4
K = 5
K = 6
lysimniaWA
lysimniaAF
nesaeaAF
polymniaWEST
polymniaWA
polymniaPERU
polymniaGUI
polymniaAF

B

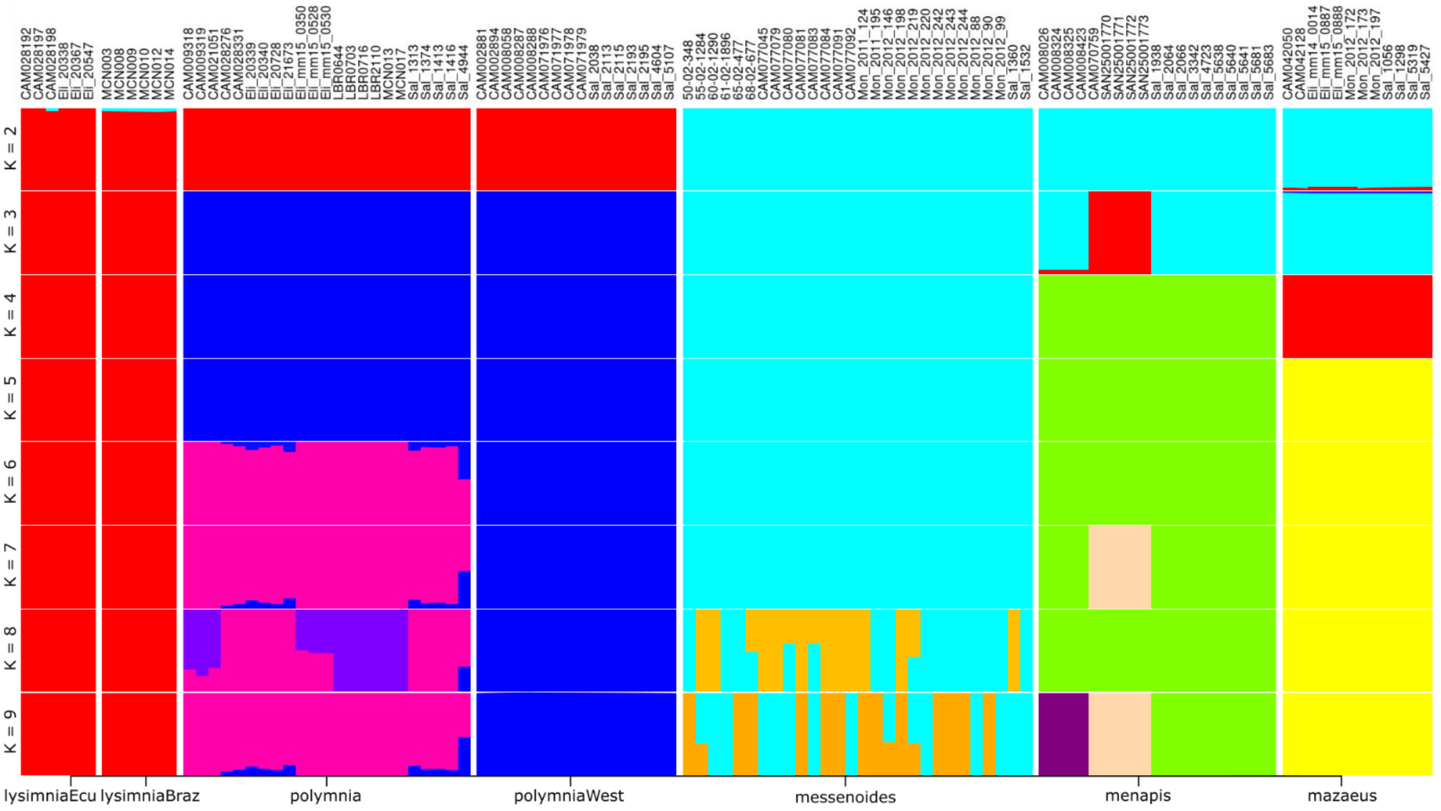
K = 2
K = 3
K = 4
K = 5
K = 6
K = 7
K = 8
K = 9
lysimniaEcu
lysimniaBraz
polymnia
polymniaWest
messenoides
menapis
mazaeus

C

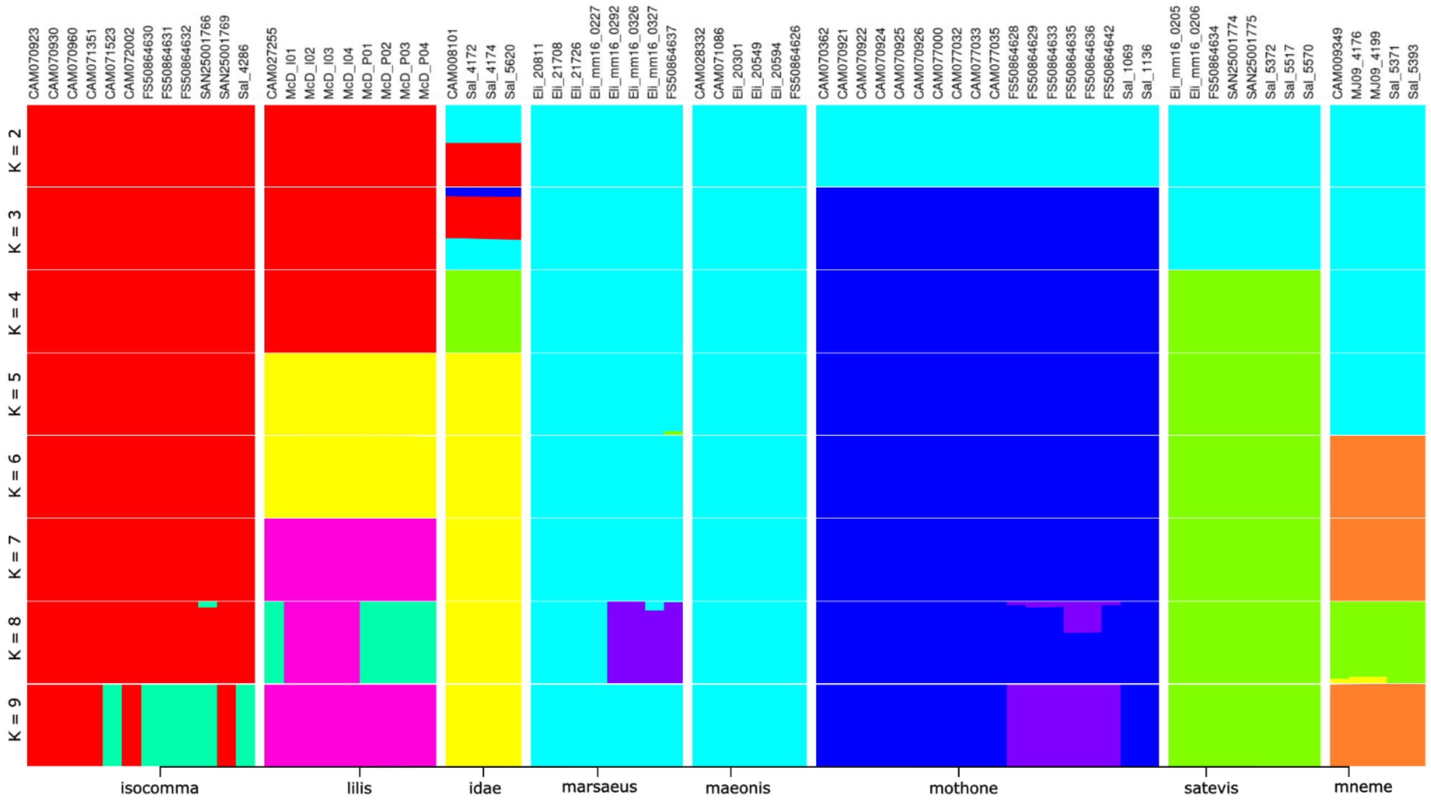


**Fig. S8. ADMIXTURE plots**

*A) ADMIXTURE in* Mec. polymnia*,* Mec. lysimnia *and* Mec. nesaea*. The red star indicates the Colombian polymnia individual (Sal_4944) caught in Western Amazonia, that appears to have equal ancestry with polymnia Western Amazonia and polymnia West of the Andes.*

*B) ADMIXTURE in* Mec. lysimnia, Mec. polymnia, Mec. messenoides, Mec. menapis, *and* Mec. mazaeus*.*

*C) ADMIXTURE in* Mel. isocomma, Mel. lilis, Mel. idae, Mel. marsaeus, Mel. maeonis, Mel. mothone, Mel. satevis and Mel. mneme.

*WA = Western Amazonia; AF = Atlantic Forest; West = west of the Andes; GUI = Guianan Shield; Ecu = Ecuador; Braz = Brazil (same region as Atlantic Forest)*

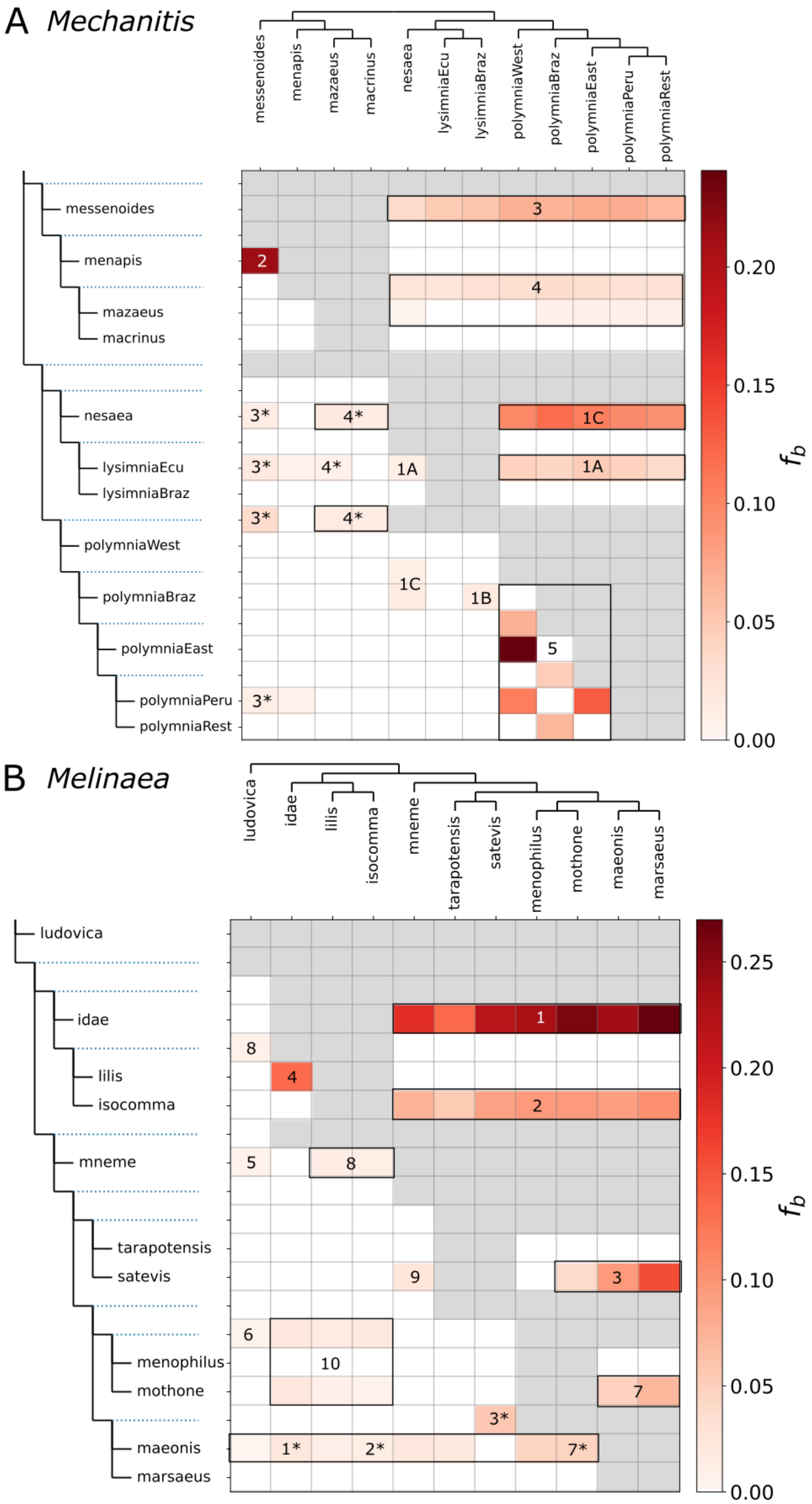


**Fig. S9. Fbranch DSuite results**

*Fbranch results for* Mechanitis *(A) and* Melinaea *(B). Stronger red colour indicates more excess allele sharing between the species shown on top and the species shown on the left as compared to the sister clade of the latter according to the phylogeny on the left. Note, if the direction of gene flow is from the species on top to the species on the left, close relatives of the species on top are also expected to show excess allele sharing. The numbers indicate which excess allele sharing pattern was used to draw the hybridisation arrows in Figure 2.*

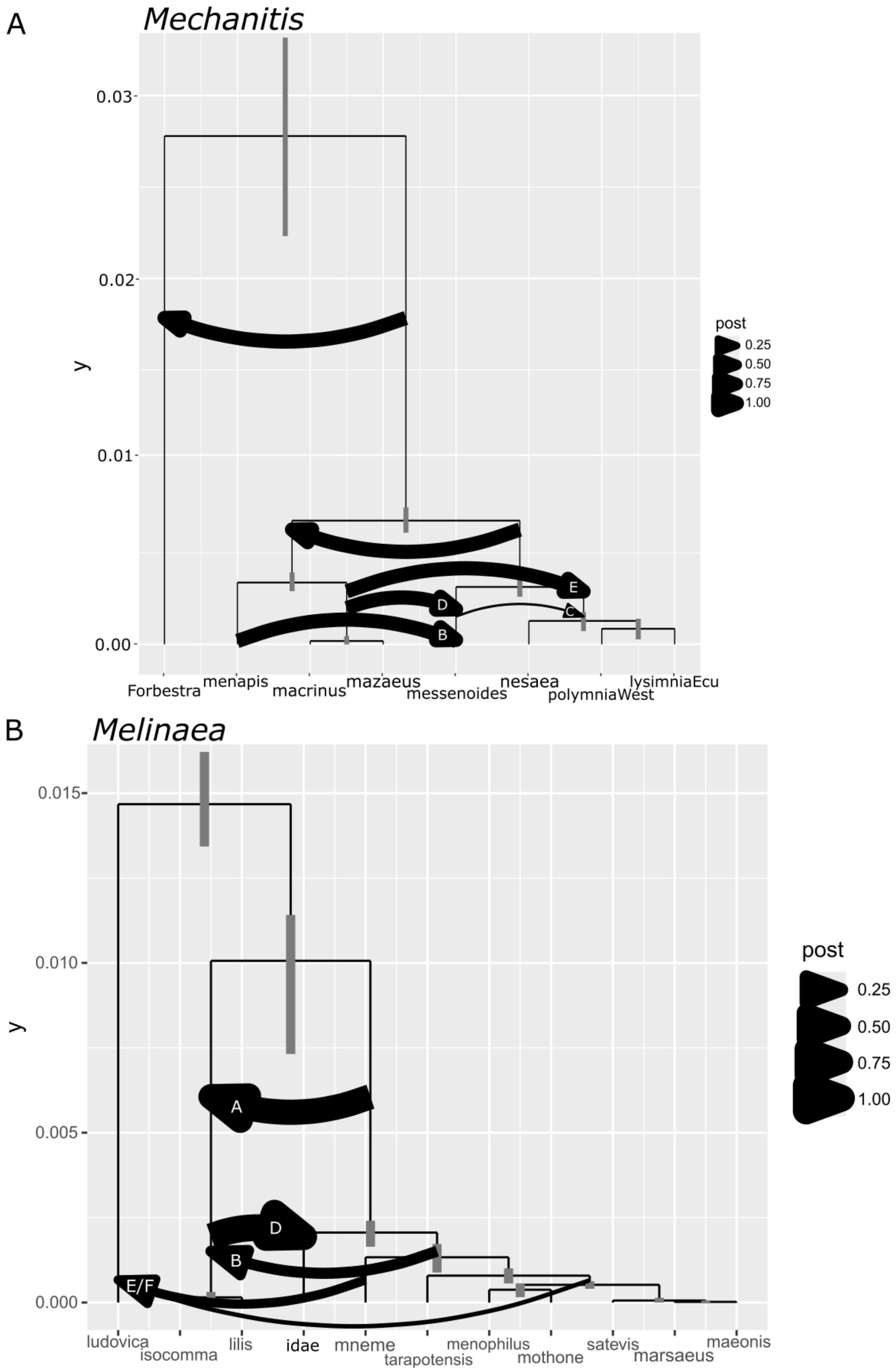


**Fig. S10. AIM (BEAST2) admixture trees**

*A tree with a set of admixture arrows that might explain the phylogenetic history of* Mechanitis *(A) and* Melinaea *(B), as produced by AIM. The letters in the arrows are used in the text to refer to different admixture events and they correspond to arrows in Figure 2.*

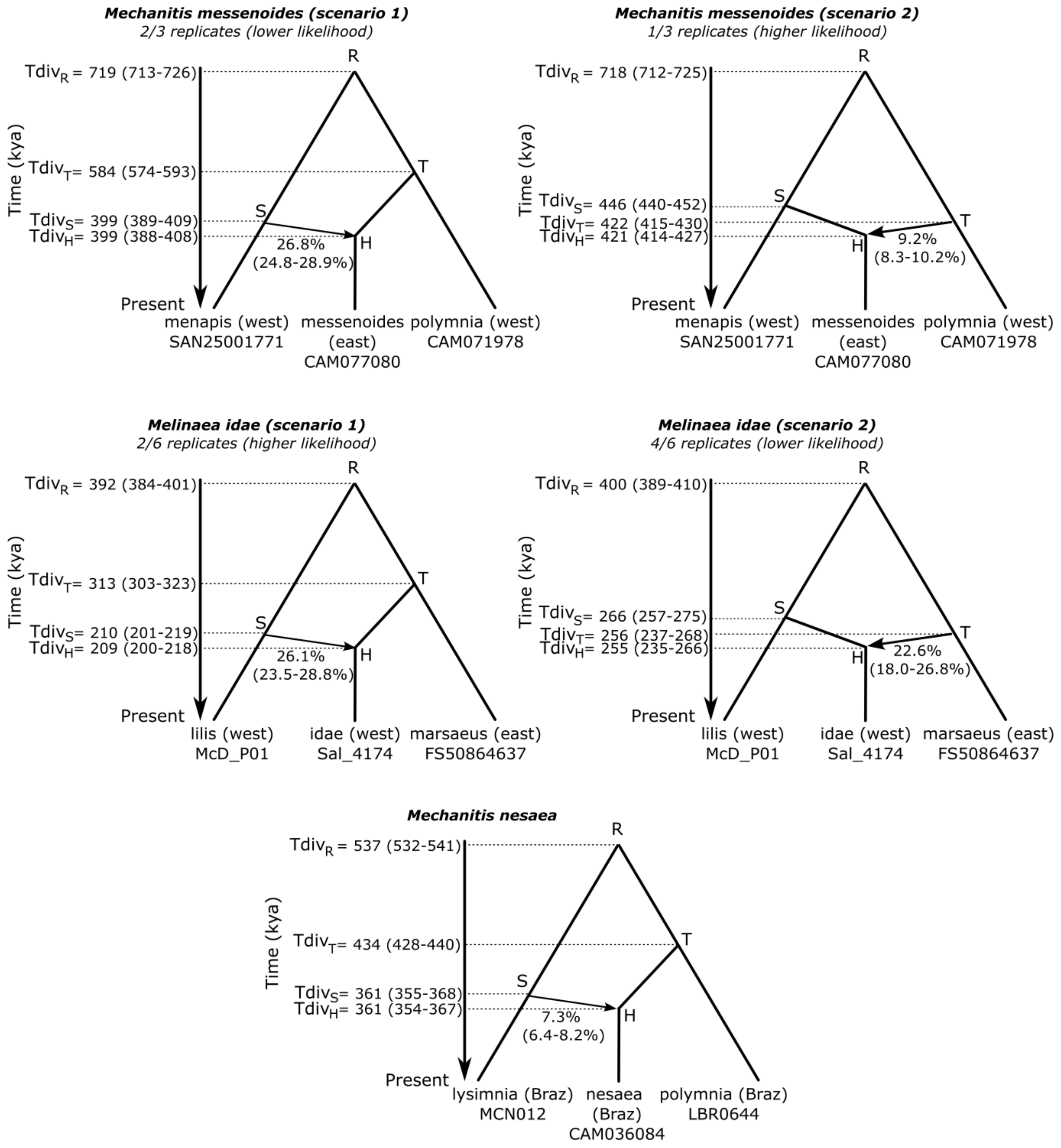


**Fig. S11. MSCi model results**

*The figure shows the multi-species coalescent model with introgression (MSCi) for each introgressed species of interest, with the obtained divergence times and the corrresponding confidence intervals. We assumed 4 generations per year, and a mutation rate of 2.9x10^9. For Mechanitis messenoides and Melinaea idae, different replicates converged on a different output, so we have provided both scenarios. The branching order is not pre-determined in the model; T can be younger or older than S.*

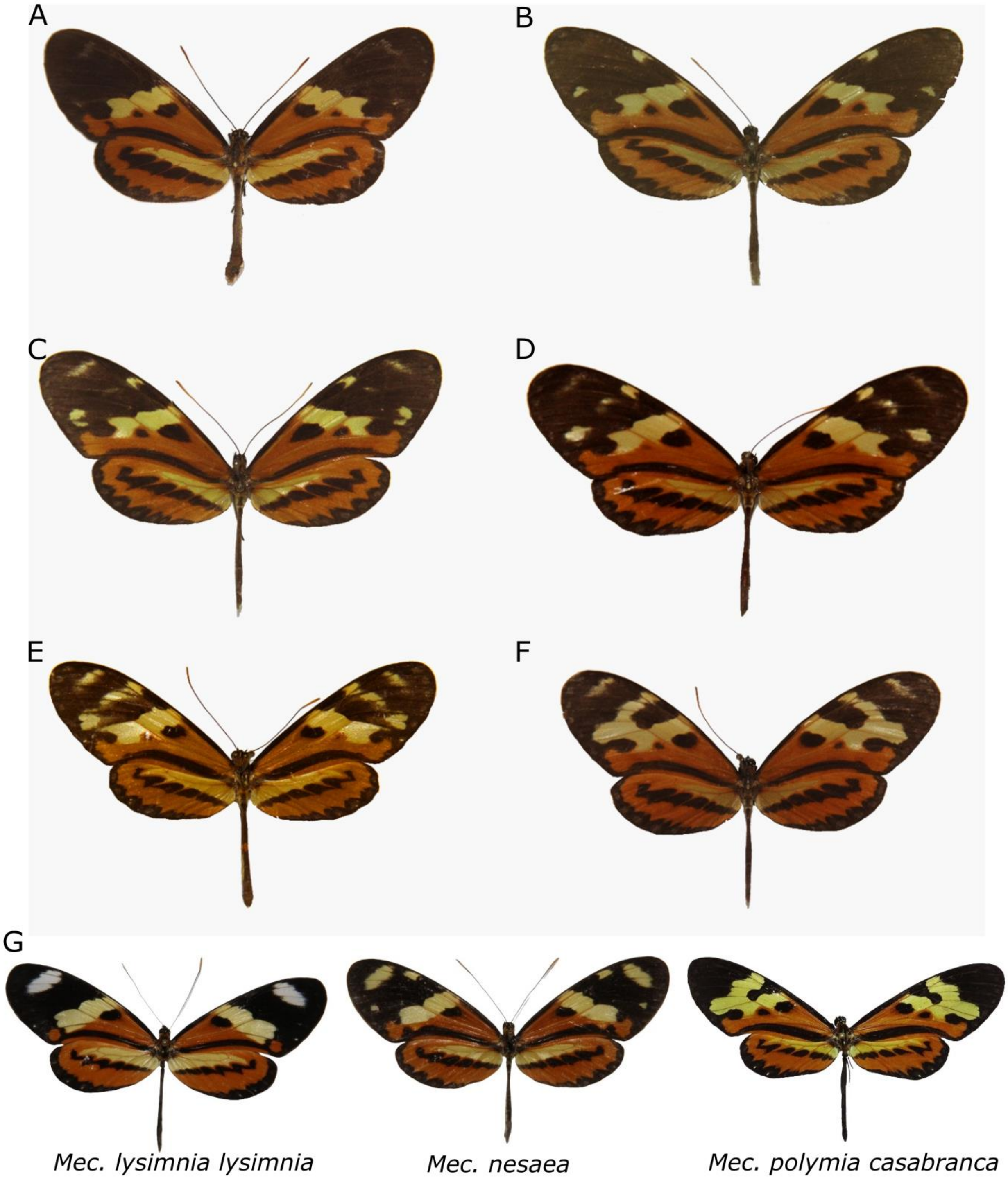


**Fig. S12. Hybrids between *Mec. polymnia* and *Mec. lysimnia* look like *Mec. nesaea***

*Hybrids between Mec. l. lysimnia and Mec. p. casabranca found in Brazil, some (e.g. C) look like the putative hybrid species Mec. nesaea. Photos coming from: A) Brazil, São Paulo, São Vicente, Morro do Japui, 25.v.1991 - ZUEC-AVLF. B) Brazil, Minas Gerais, Itueta, viii.1970 – ZUEC. C) Brazil, Minas Gerais, Itueta, v.1970 – ZUEC. D) Brazil, Espírito Santo, Baixo Guandú, vi.1970 – ZUEC. E) Brazil, Espírito Santo, Baixo Guandú, v.1970 – ZUEC. F) Brazil, Espírito Santo, Baixo Guandú, v.1970 – ZUEC. ZUEC = Zoological collection, Museu de Diversidade Biológica da Universidade Estadual de Campinas, Campinas, São Paulo, Brazil. ZUEC-AVLF = André V.L. Freitas Collection, Universidade Estadual de Campinas, Campinas, São Paulo, Brazil. G) Photos of Mec. lysimnia lysimnia, Mec. nesaea, and Mec. polymnia casabranca.*

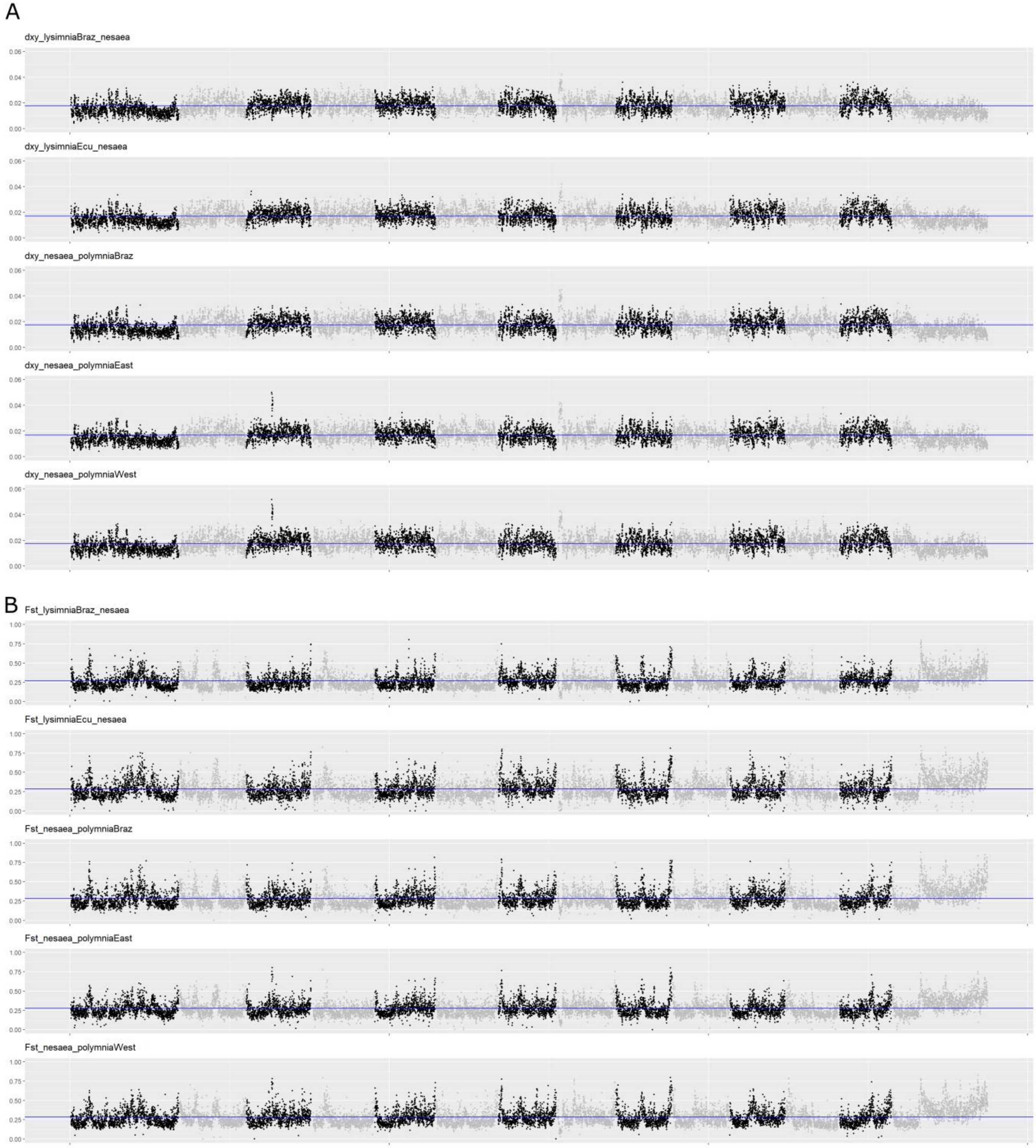


**Fig. S13. Divergence and diversity in *Mec. polymnia, Mec. lysimnia* and *Mec. nesaea***

*A) Dxy absolute divergence analysis. The title of each row indicates between which two populations the Dxy is calculated. High peaks indicate more divergence.*
*B) Fst relative differentiation analysis. The title of each row indicates between which two populations the Fst was calculated. High peaks indicate more differentiation.*
*Black versus grey dots indicate a new chromosome. Window-size was 20kb.*

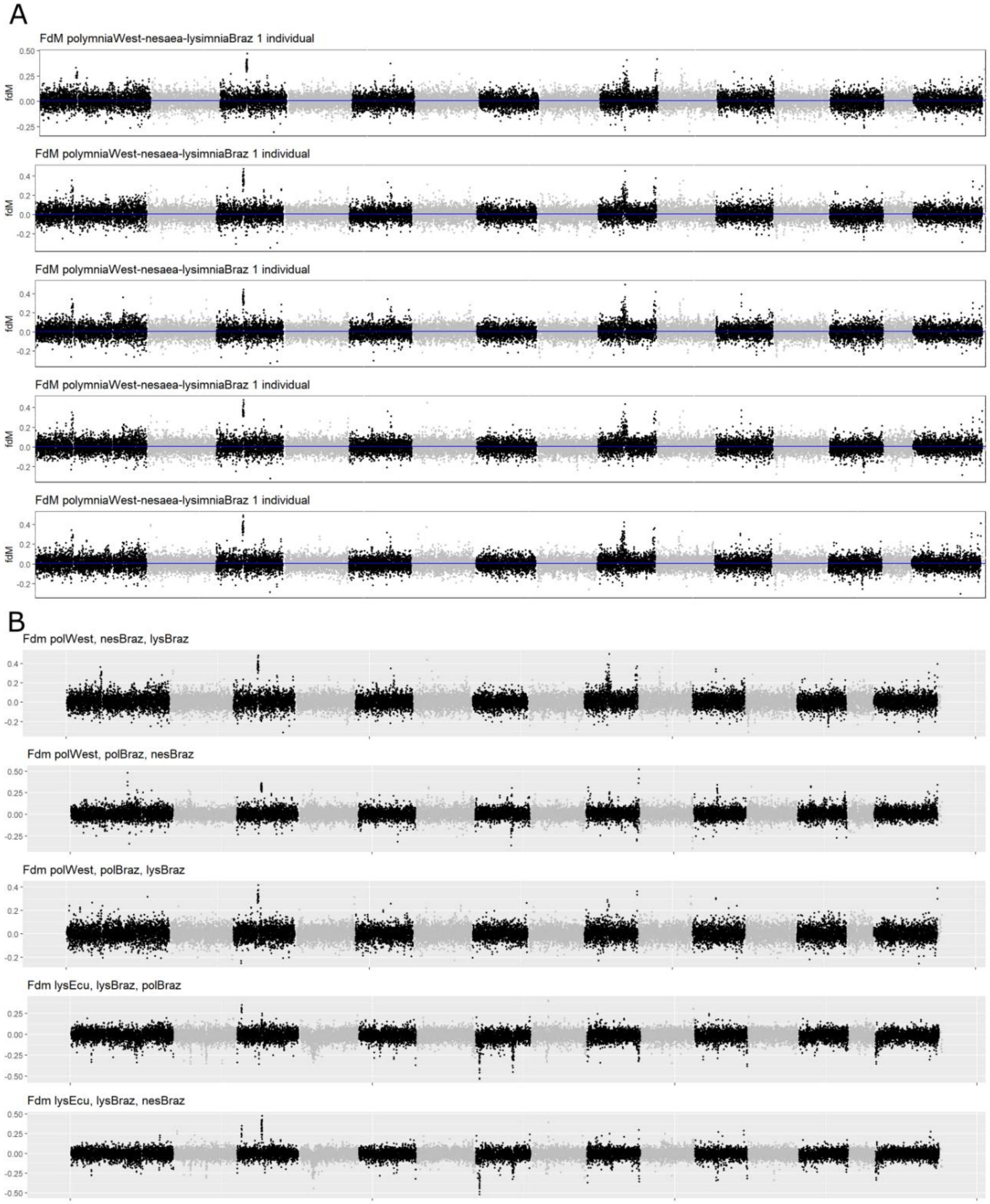


**Fig. S14. Introgression analysis in *Mec. polymnia, Mec. lysimnia* and *Mec. nesaea***

*A) fdM introgression analysis with P1 = Mec. polymnia (from West of the Andes), P2 = Mec. nesaea (1 individual from Brazil per row) and P3 = Mec. lysimnia (Brazil). P4 is the outgroup Forbestra. The same analysis was done for five different individuals of Mec. nesaea, to examine how consistent the results are between individuals. High fdM indicates high introgression between P2 and P3 (Mec. nesaea and Mec. lysimnia).*
*B) Various fdM introgression analyses to compare gene flow between different populations. The order of the names in the titel of each row gives P1, P2 and P3. Forbestra is the outgroup (P4) in each analyses. One can see that certain peaks of introgression are shared between populations.*
*Black versus grey dots indicate a new chromosome.*

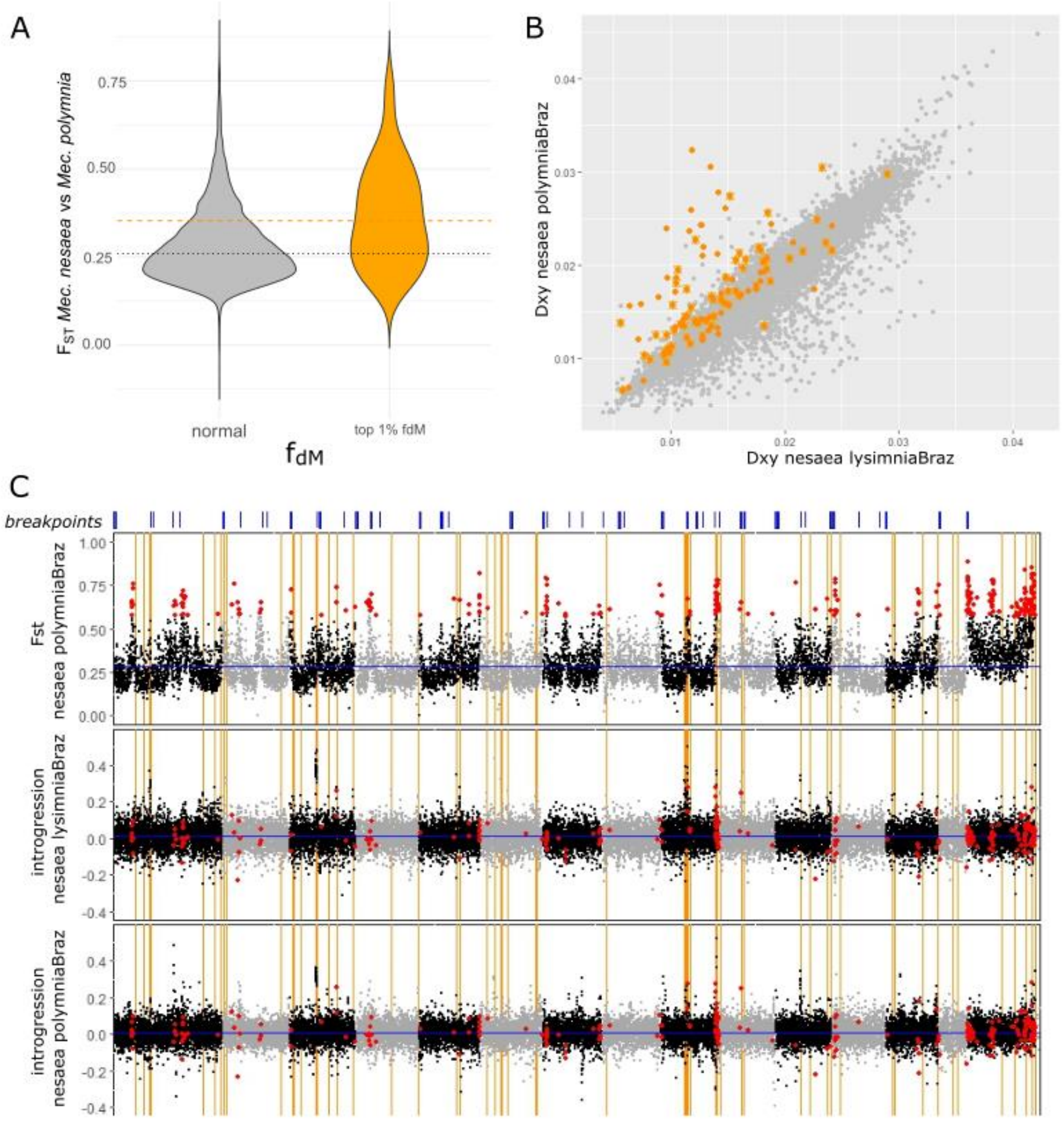


**Fig. S15.** ***Mec. nesaea*** **differentiation and introgression**

Mec. nesaea *shows increased differentiation from its sister species* Mec. polymnia *in regions with strong signatures of introgression with* Mec. lysimnia*. A)* $F_{ST}$ *between* Mec. polymnia *and* Mec. nesaea *computed in 20kb windows is higher in top 1%* $f_{dM}$ *windows (orange) than in the rest of the genome (grey, Kruskal-Wallis test: 39.395, df = 1, p-value < 3.5e-10). The dotted black horizontal line indicates the median value of Fst, and the dashed orange horizontal line indicates the median of the top 1%* $f_{dM}$ *windows. B)* $D_{XY}$ *values computed between* Mec. nesaea *and both close relatives with the top 1% fdM windows highlighted in orange. The orange stars are windows that also have low fdM values for* Mec. nesaea *and* Mec. lysimnia. *C) A scan across the genome (20kb windows) showing* $F_{ST}$ *(*Mec. nesaea *vs* Mec. polymnia *of Brazil) and* $f_{dM}$ *(P1=polymniaWest, P2=nesaea, P3=lysimniaBrazil, P4=Forbestra) and (P1=polymniaWest, P2=polymniaBrazil, P3=nesaea, P4=Forbestra). Blue lines at the top indicate potential chromosomal breakpoints (breakpoints from* Mec. polymnia *with other four reference species, as we don't have a reference genome for* Mec. nesaea*). Orange vertical lines indicate regions with strong signatures of introgression (top 0.5%). The red dots are the top 2% Fst windows.*

*Mec. macrinus*

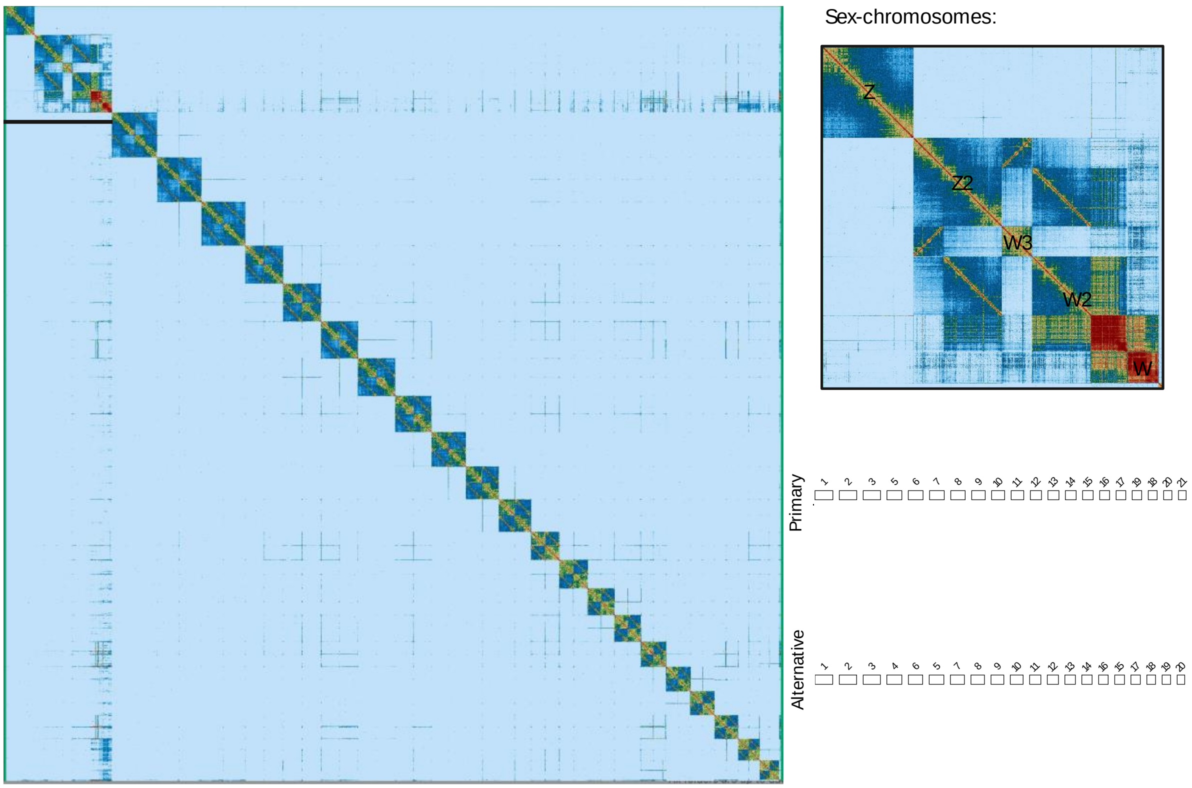


*Mec. mazaeus*

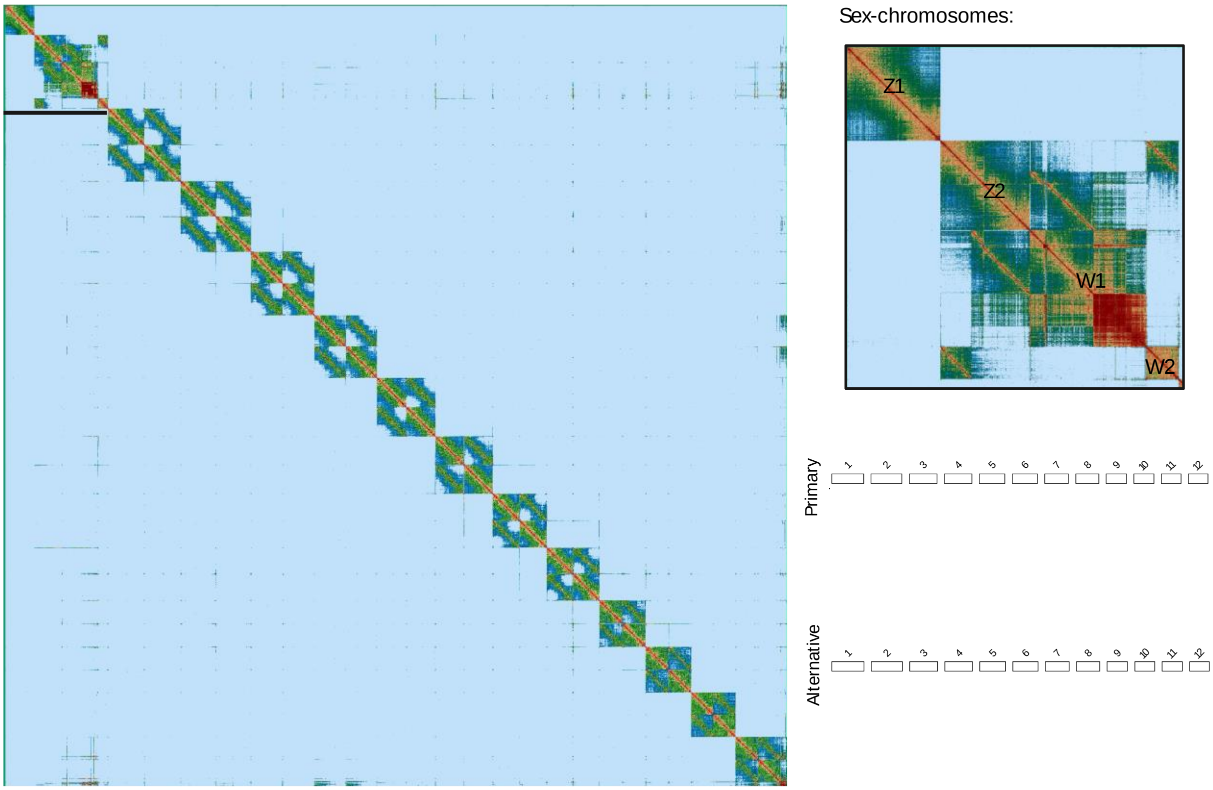

*Mec. menapis*

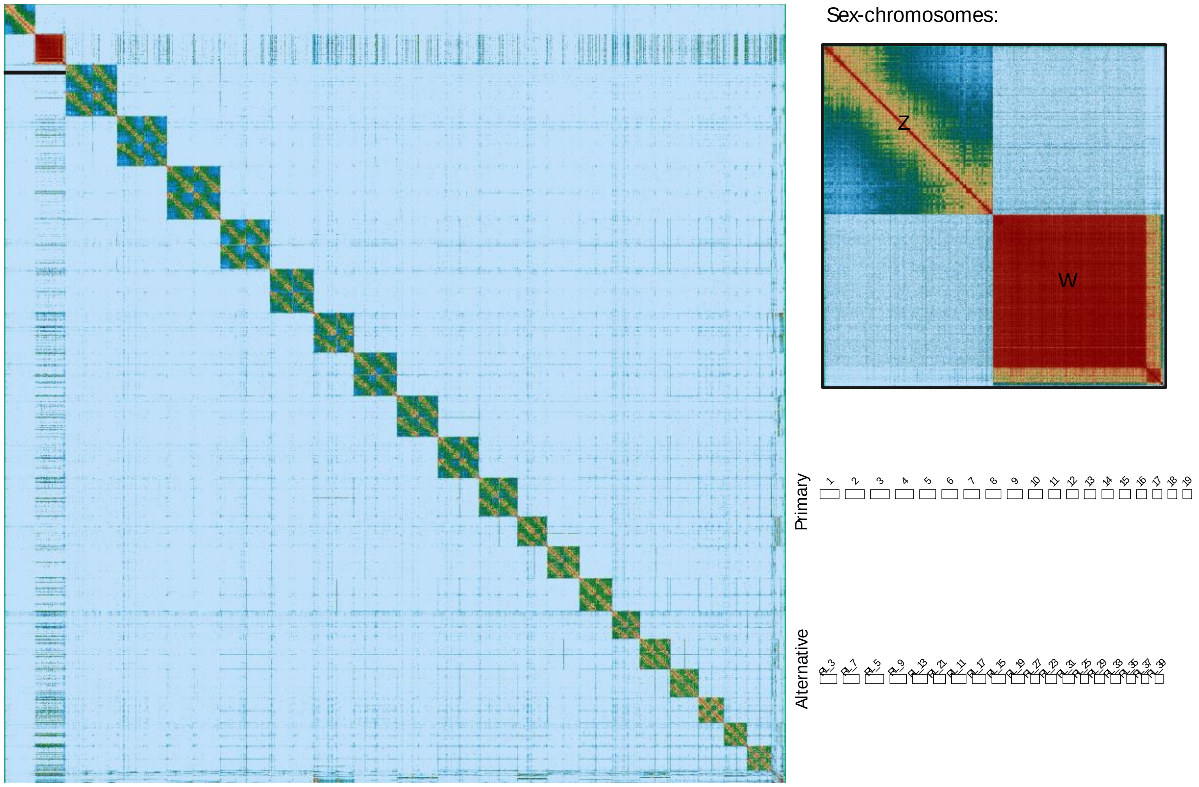


*Mec. messenoides*

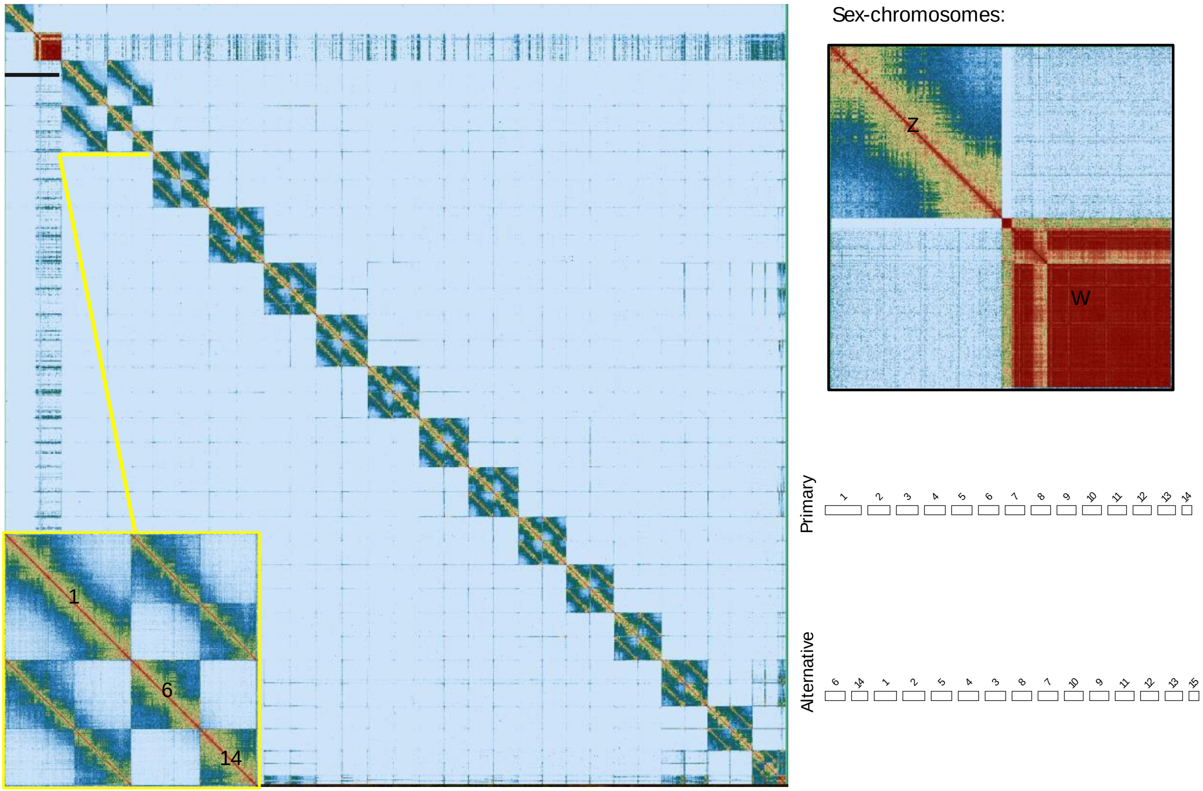

*Mec. polymnia*

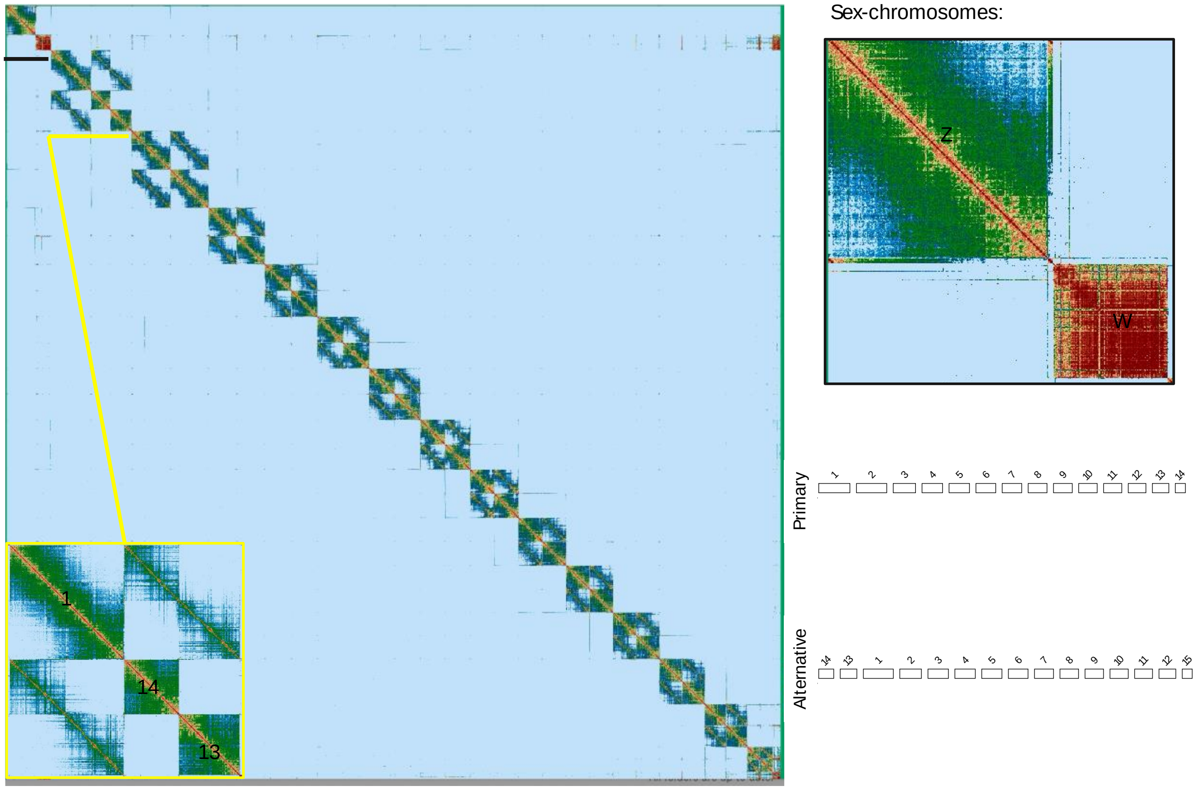


**Fig. S16a. Haplotype mapping**

*HiC contact maps of reads mapped simultaneously to the two haplotype phased assemblies in each* Mechanitis *genome, with sex-chromosomes (black line) and heterozygous rearrangements (yellow line) highlighted. Insets: Examples of autosomal heterozygous rearrangements. Right panel: Zoom in of sex-chromosomes. Bottom panel: Whole genome alignment between the two phased assemblies, excluding hemizygous chromosomes.*

*Mel. ludovica*

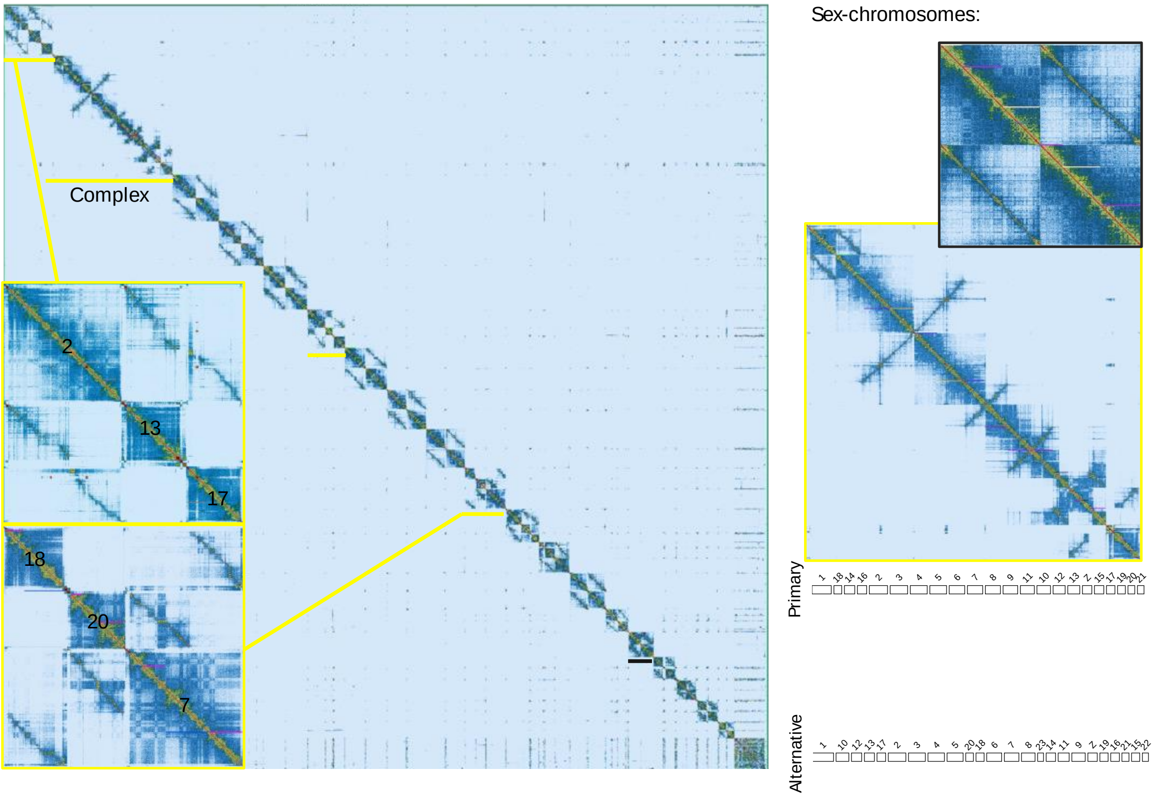


*Mel. isocomma*

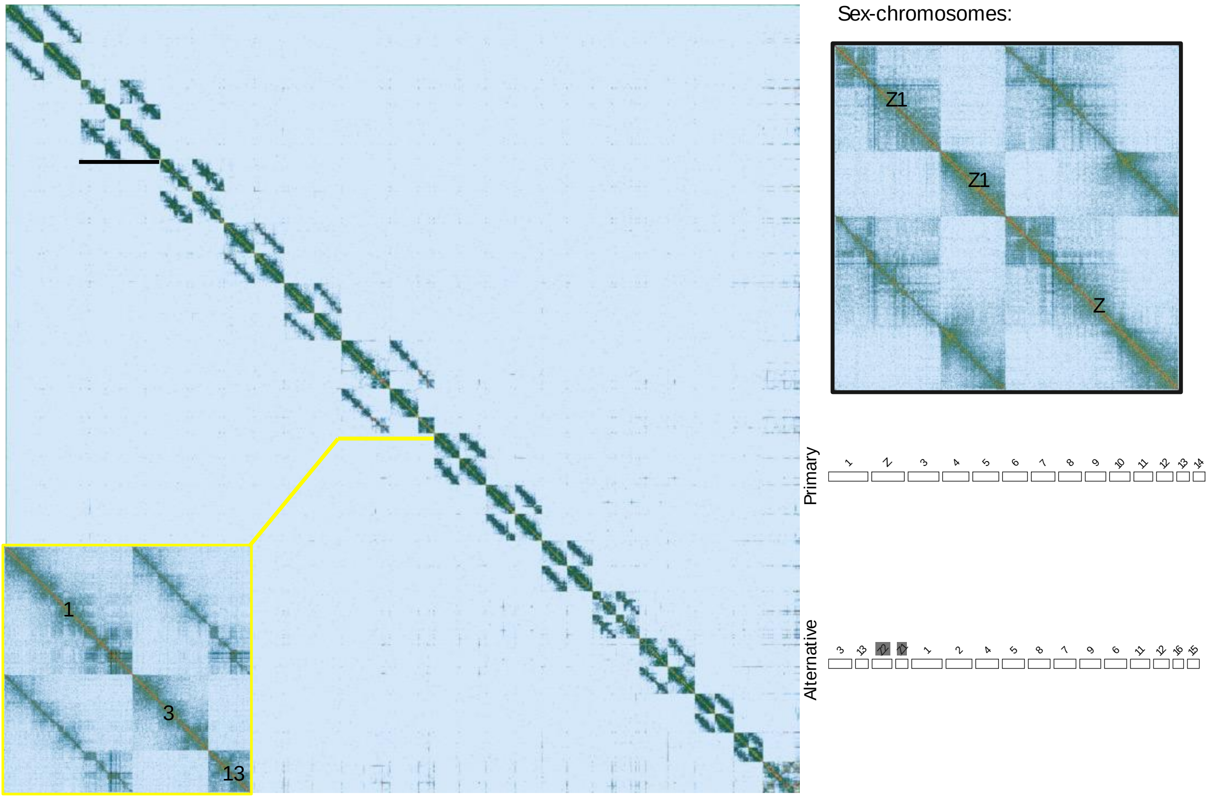

*Mel. mothone*

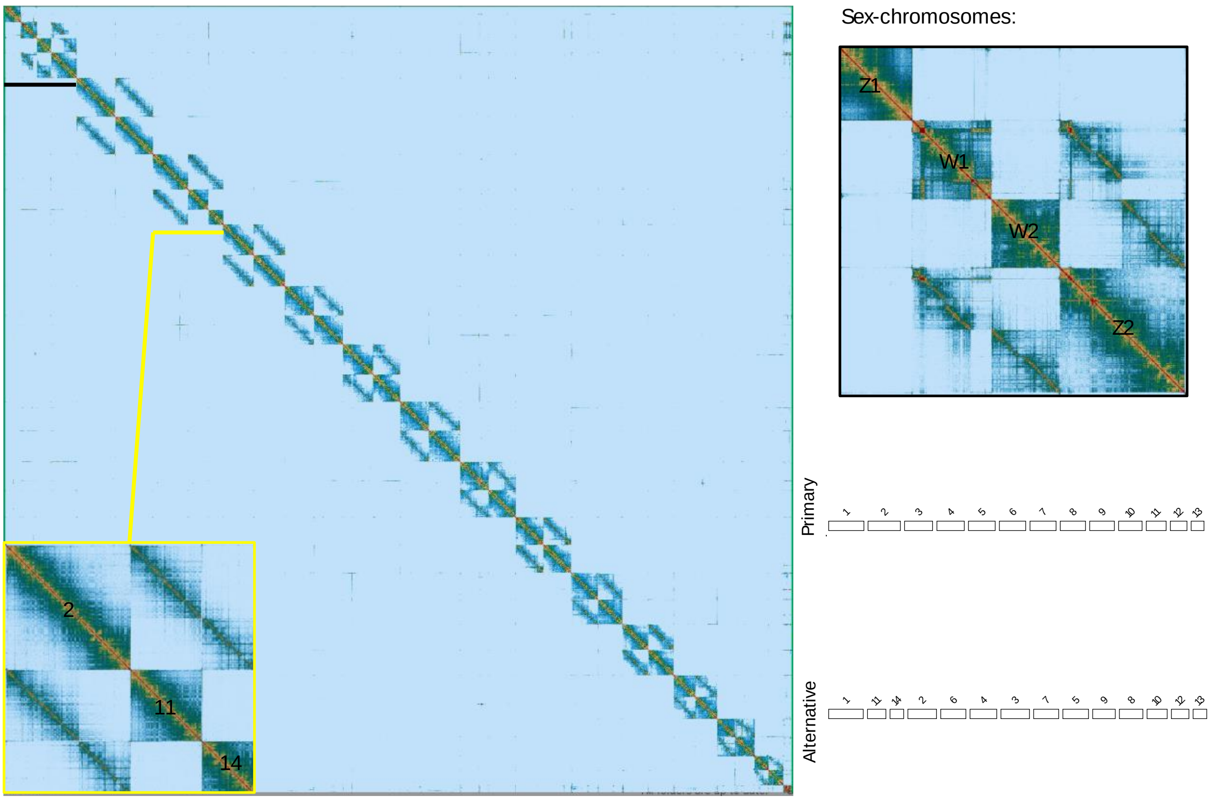


*Mel. menophilus*

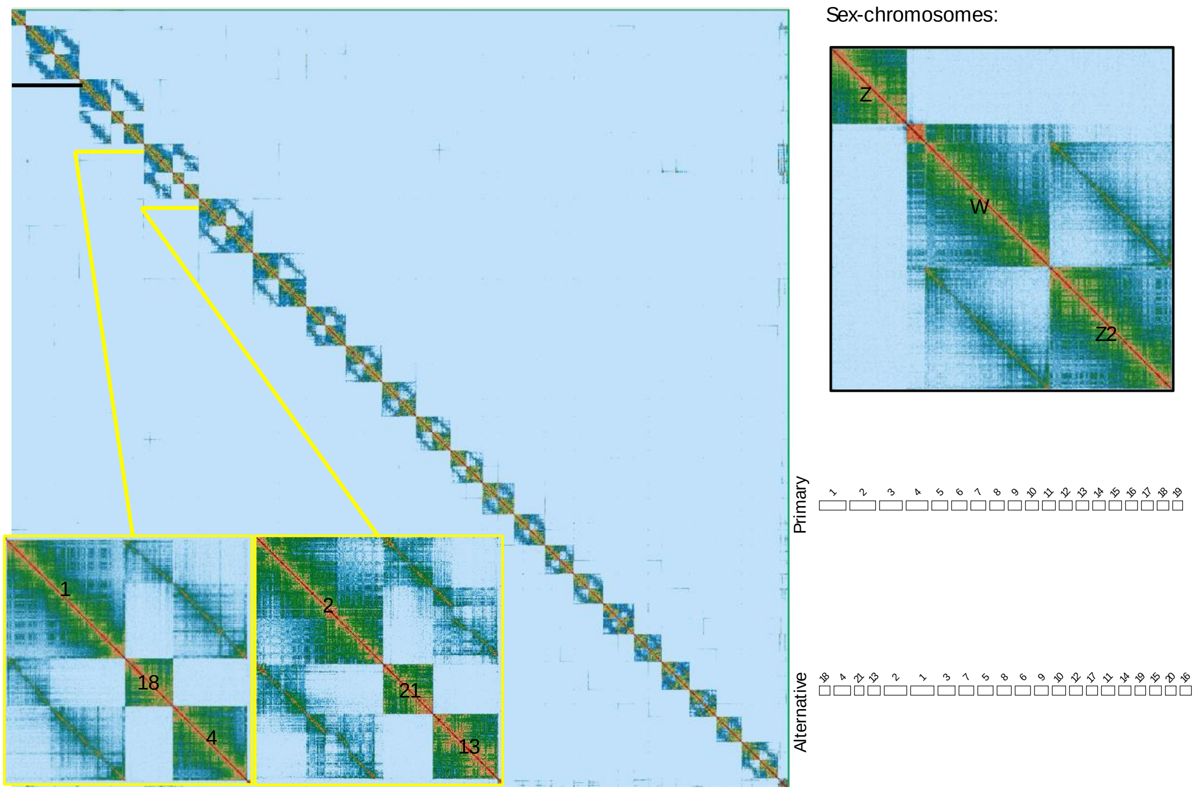

*Mel. marsaeus*

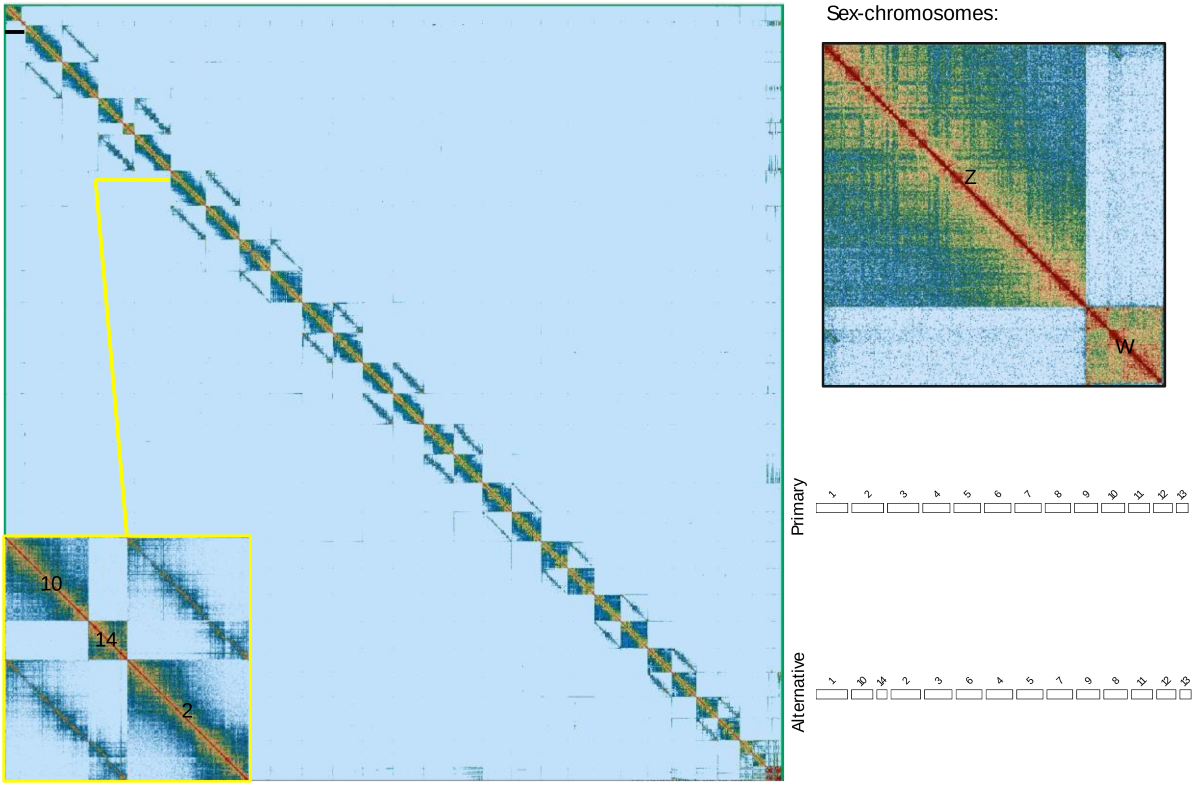


**Fig. S16b. Haplotype mapping**

*HiC contact maps of reads mapped simultaneously to the two haplotype phased assemblies in each* Melinaea *genome, with sex-chromosomes (black line) and heterozygous rearrangements (yellow line) highlighted. Insets: Examples of autosomal heterozygous rearrangements. Right panel: Zoom in of sex-chromosomes. Bottom panel: Whole genome alignment between the two phased assemblies, excluding hemizygous chromosomes.*

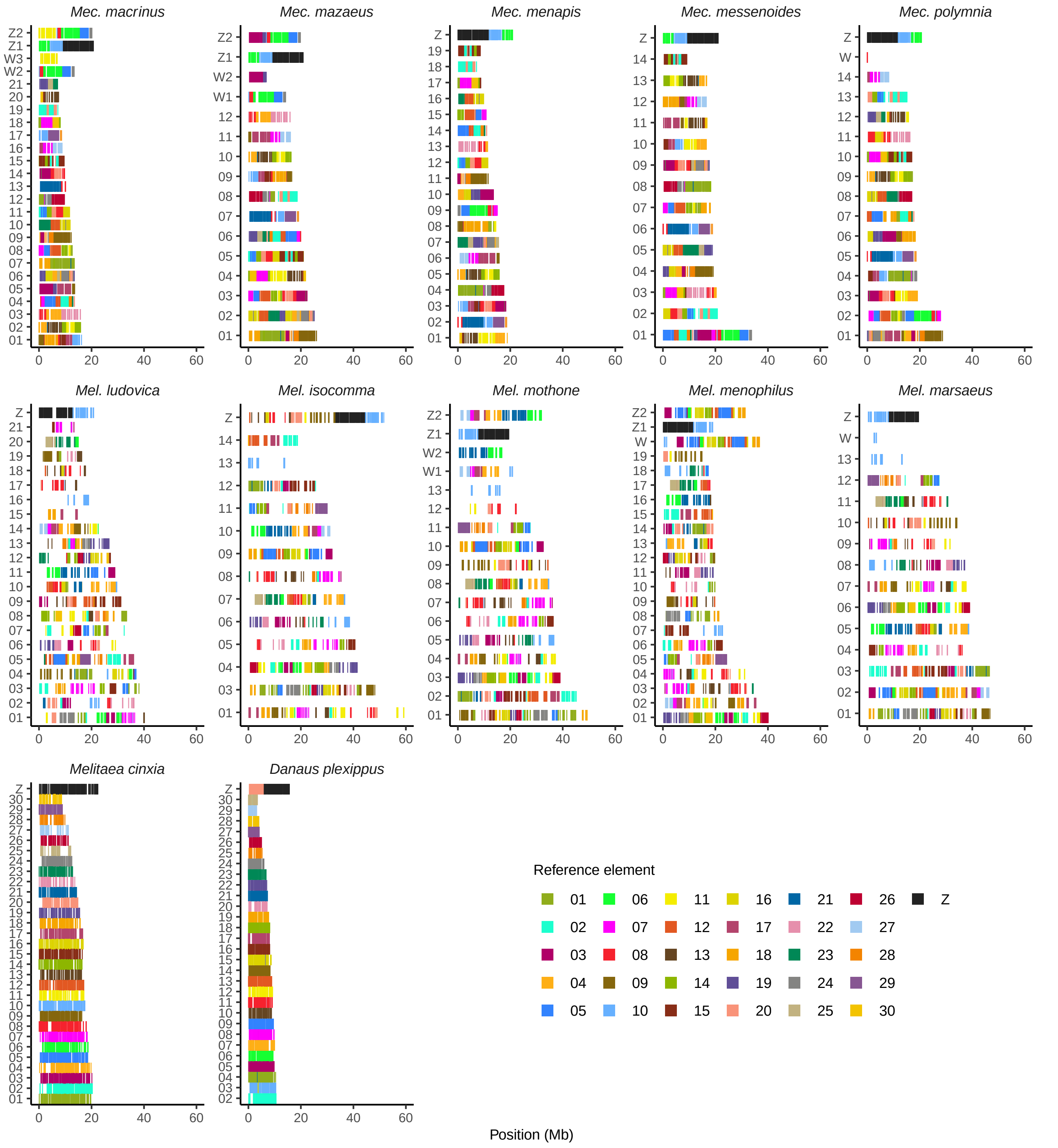


**Fig. S17. Chromosome BUSCO painting**

*Complete and duplicated BUSCO-genes in* Melinaea *and* Mechanitis *genomes coloured coloured by chromosome position in* Melitaea cinxia*. Outgroups* M. cinxia *and* Danaus plexippus *included for comparison. The chromosome number is on the y-axis and the start position of the BUSCO genes in megabases on the x-axis (Position(Mb)).*

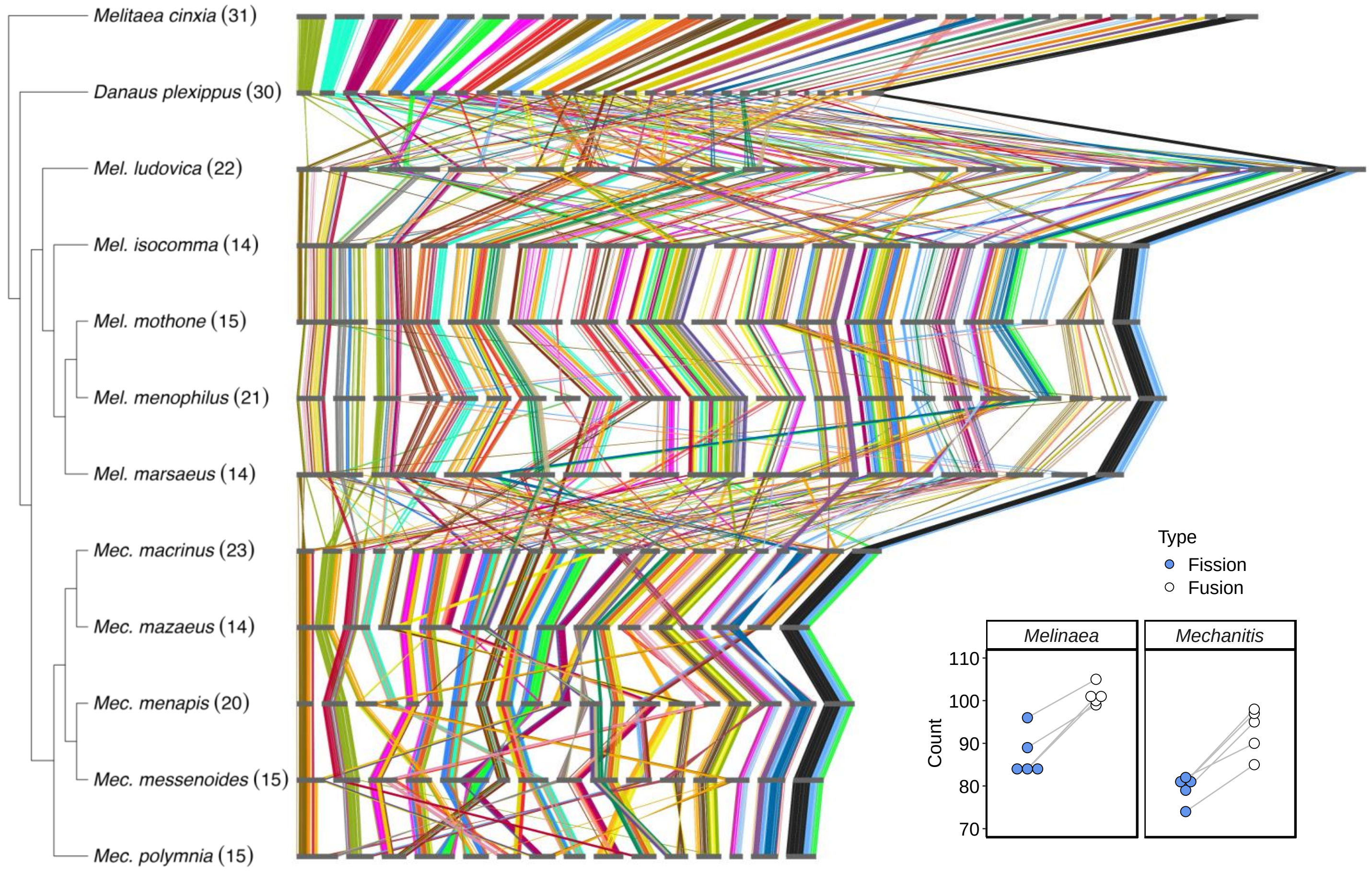


**Fig. S18. Synteny between *Melinaea* and *Mechanitis***

*Synteny between* Melinaea *(yellow) and* Mechanitis *(blue) genomes based on BUSCO gene positions using* Melitaea cinxia *and* Danaus plexippus *as outgroups (grey). Horizontal bars represent individual chromosomes and vertical links represent BUSCO genes coloured by their ancestral linkage group assignment performed with the reference species* Melitaea cinxia*. The cladogram is based on Fig. 2 and shows haploid chromosome numbers in parentheses. Inset: Count of minimum number of chromosomal fission (blue) and fusion (white) events in* Melinaea *and* Mechanitis *relative the reference species* Melitaea cinxia*.*

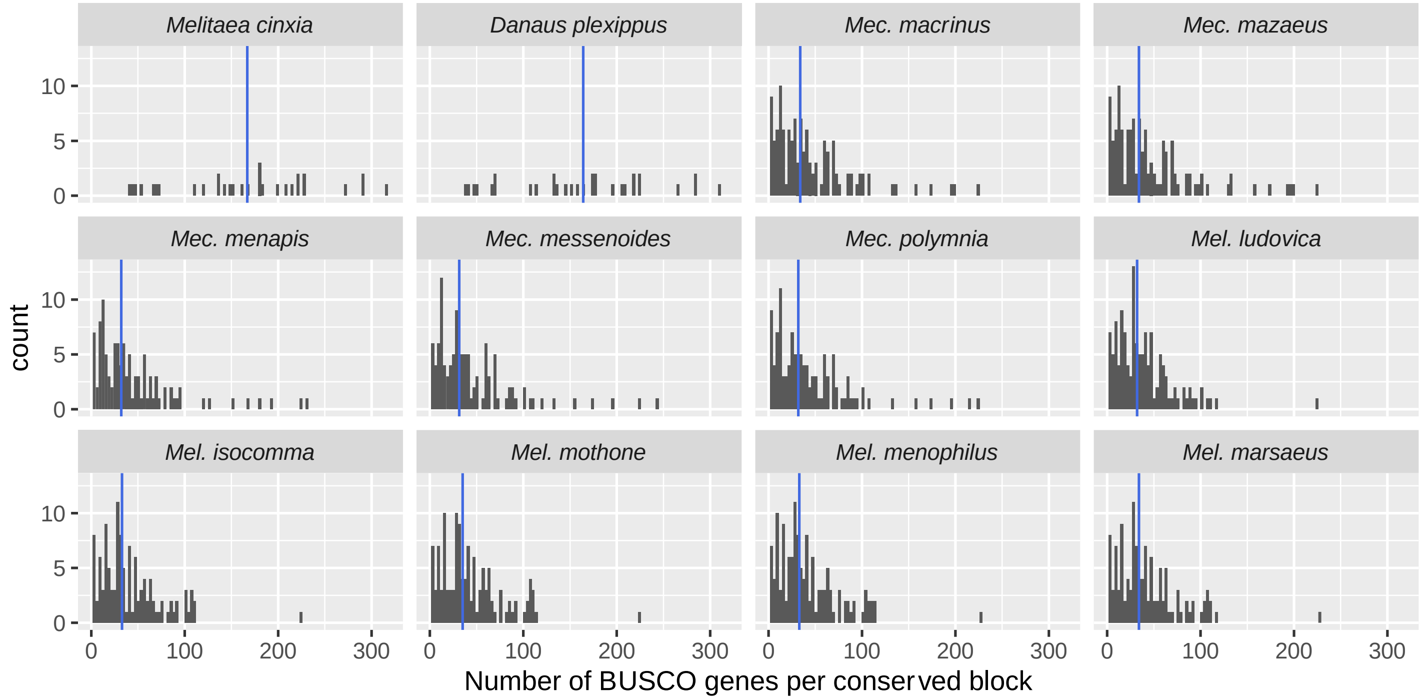


**Fig. S19. Conserved syntenic blocks**

Distribution of length of conserved syntenic blocks in number of BUSCO genes per block compared to the ancestral butterfly karyotype represented by *Melitaea cinxia*. Median number of genes per block (blue vertical line).

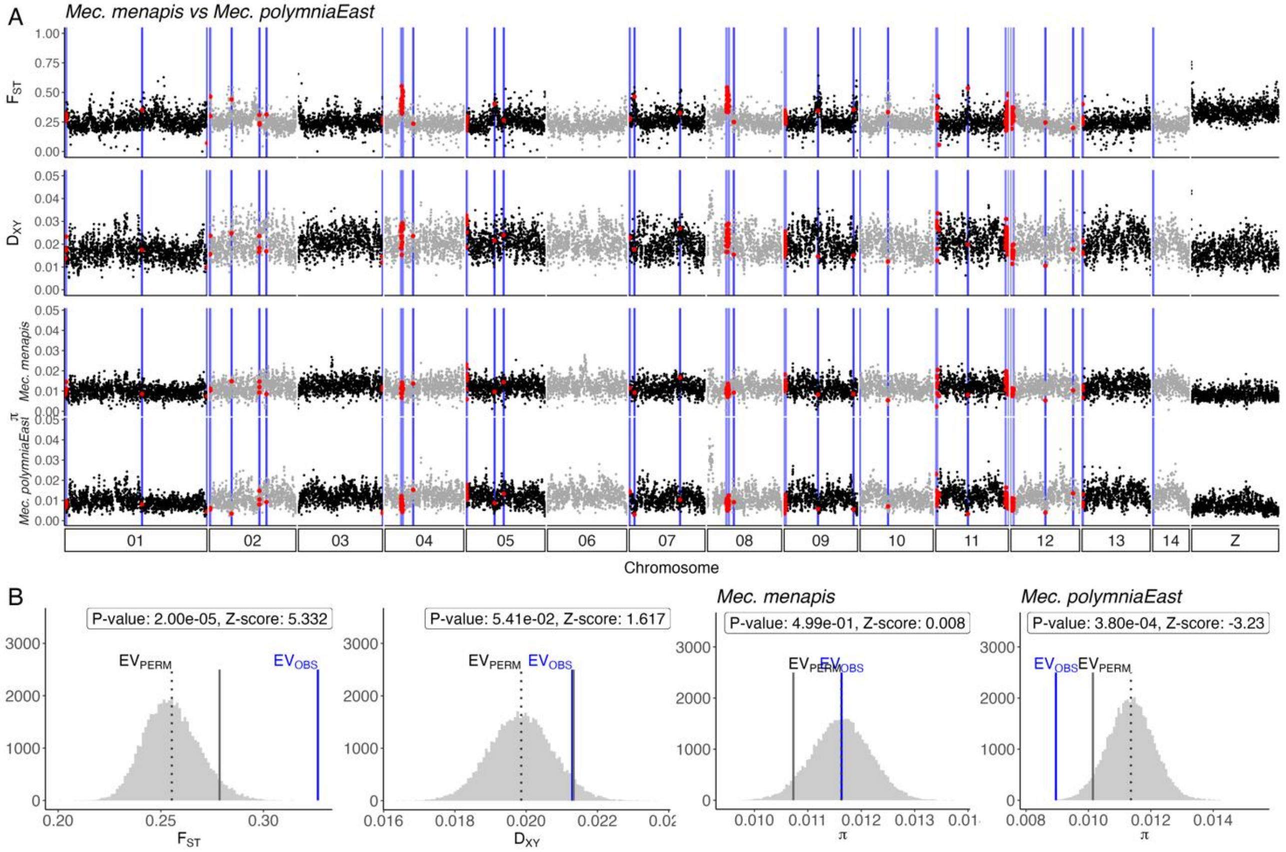
A
Mec. menapis vs Mec. polymniaEast
F_ST
D_XY
π
Mec. menapis
Mec. polymniaEast
01
02
03
04
05
06
07
08
09
10
11
12
13
14
Z
Chromosome
B
P-value: 2.00e-05, Z-score: 5.332
P-value: 5.41e-02, Z-score: 1.617
Mec. menapis
P-value: 4.99e-01, Z-score: 0.008
Mec. polymniaEast
P-value: 3.80e-04, Z-score: -3.23
EV_PERM
EV_OBS
F_ST
D_XY
π

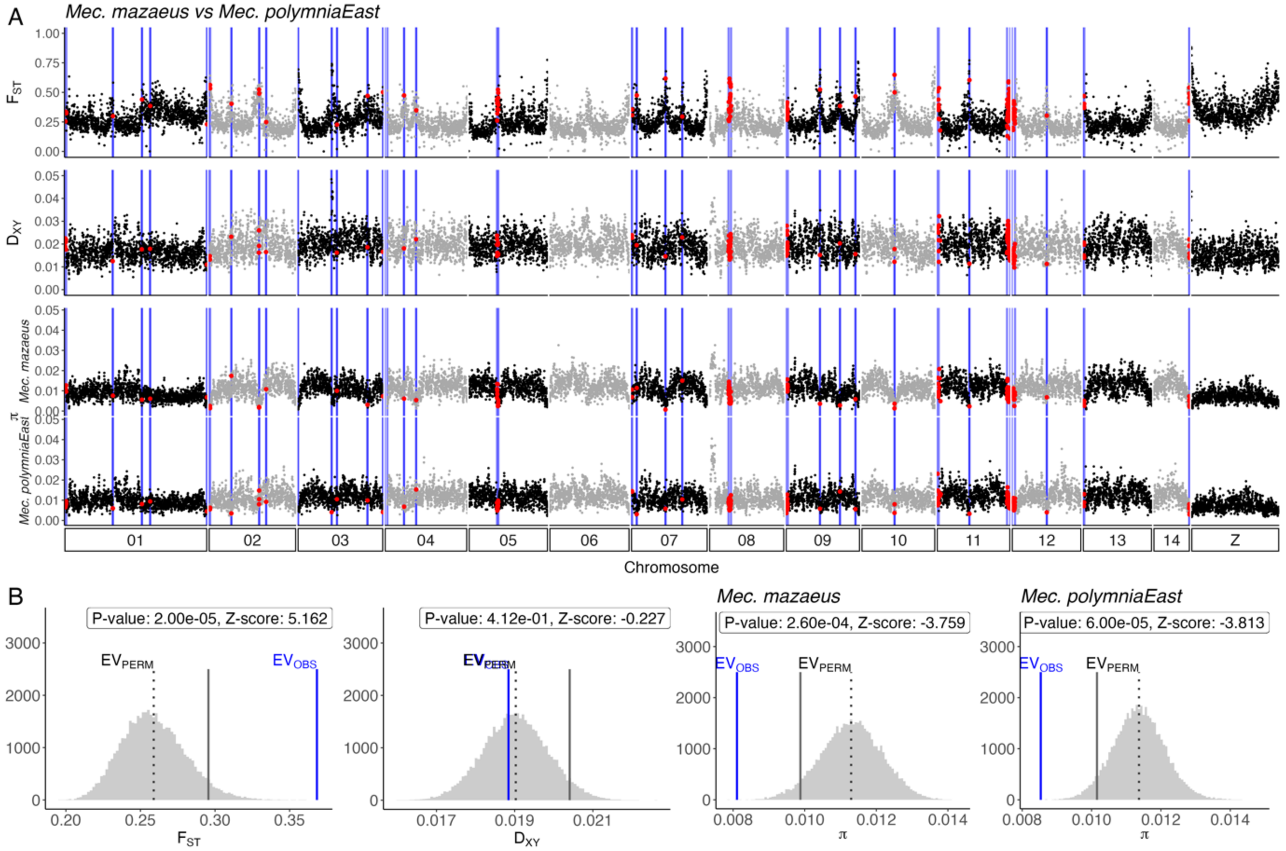
A
Mec. mazaeus vs Mec. polymniaEast
F_ST
D_XY
π
Mec. mazaeus
Mec. polymniaEast
01
02
03
04
05
06
07
08
09
10
11
12
13
14
Z
Chromosome
B
P-value: 2.00e-05, Z-score: 5.162
P-value: 4.12e-01, Z-score: -0.227
Mec. mazaeus
P-value: 2.60e-04, Z-score: -3.759
Mec. polymniaEast
P-value: 6.00e-05, Z-score: -3.813
EV_PERM
EV_OBS
F_ST
D_XY
π

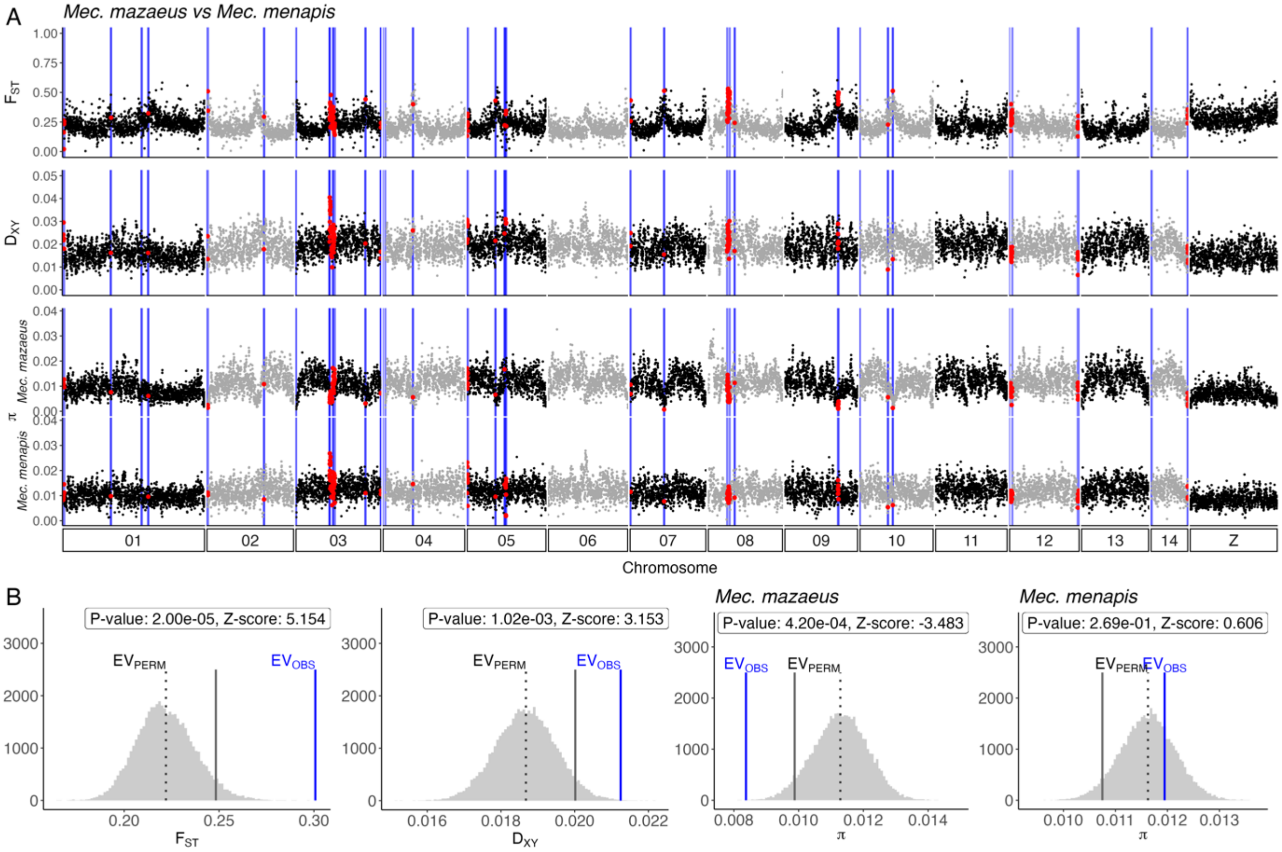
A
Mec. mazaeus vs Mec. menapis
$F_{ST}$
$D_{XY}$
π
Mec. mazaeus
Mec. menapis
01 02 03 04 05 06 07 08 09 10 11 12 13 14 Z
Chromosome
B
P-value: 2.00e-05, Z-score: 5.154
P-value: 1.02e-03, Z-score: 3.153
Mec. mazaeus
P-value: 4.20e-04, Z-score: -3.483
Mec. menapis
P-value: 2.69e-01, Z-score: 0.606
$EV_{PERM}$
$EV_{OBS}$
$F_{ST}$
$D_{XY}$
π

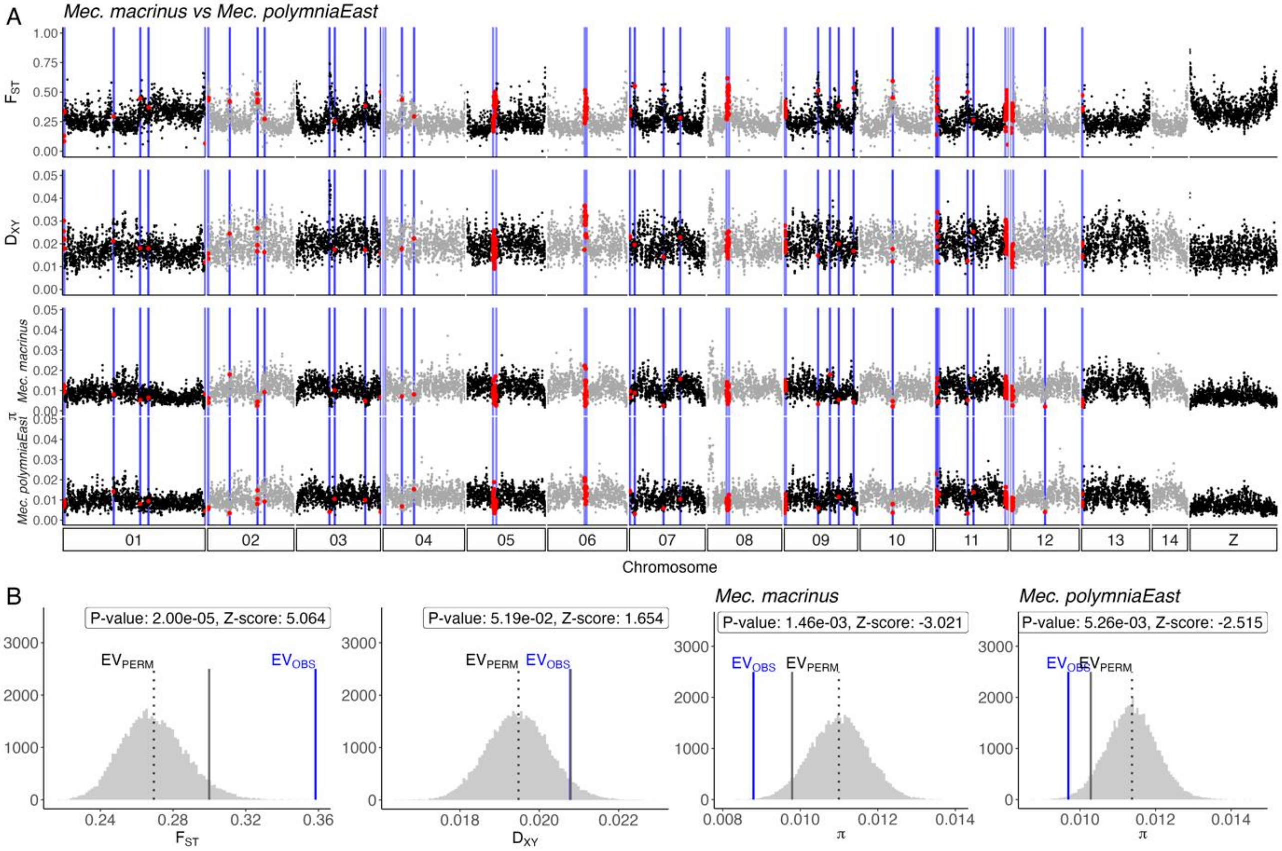
A
Mec. macrinus vs Mec. polymniaEast
$F_{ST}$
$D_{XY}$
π
Mec. macrinus
Mec. polymniaEast
01 02 03 04 05 06 07 08 09 10 11 12 13 14 Z
Chromosome
B
P-value: 2.00e-05, Z-score: 5.064
P-value: 5.19e-02, Z-score: 1.654
Mec. macrinus
P-value: 1.46e-03, Z-score: -3.021
Mec. polymniaEast
P-value: 5.26e-03, Z-score: -2.515
$EV_{PERM}$
$EV_{OBS}$
$F_{ST}$
$D_{XY}$
π

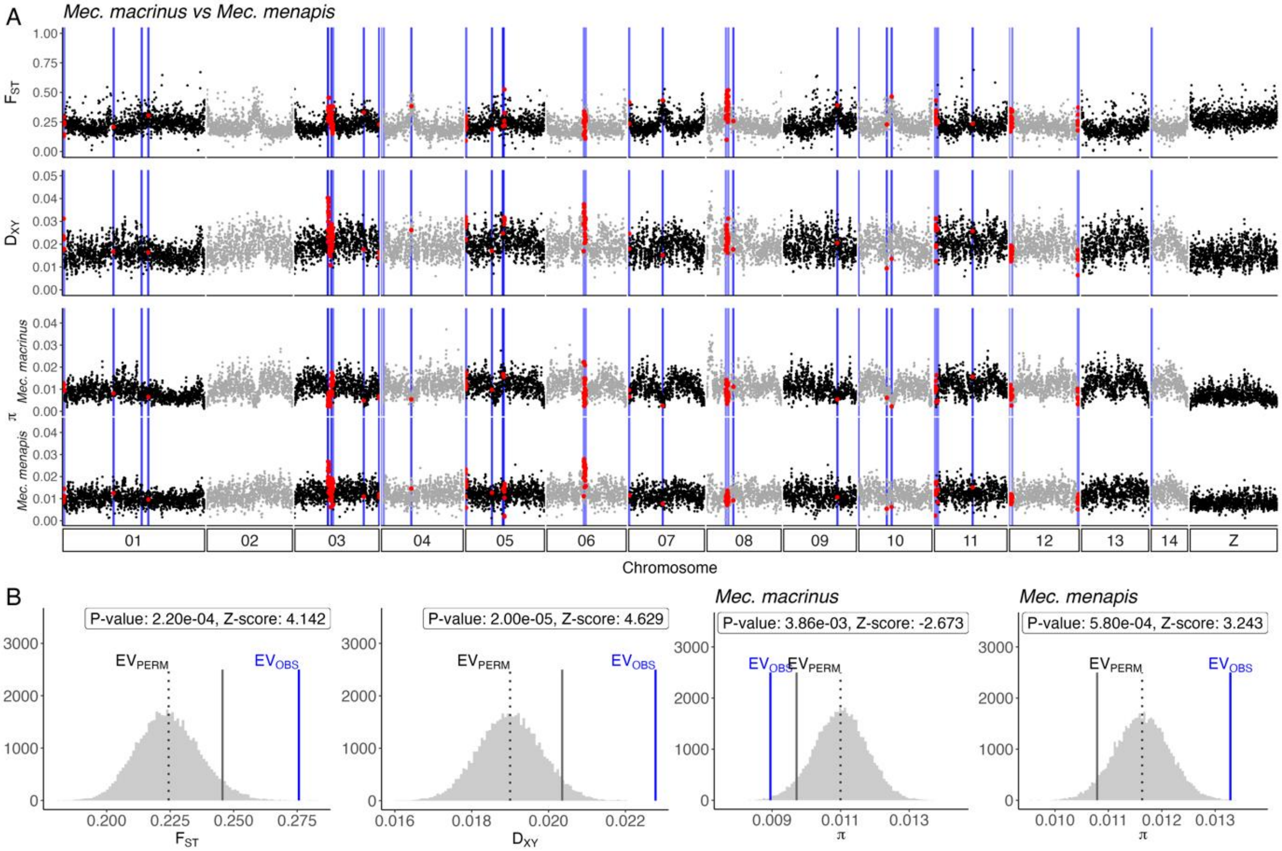
A
Mec. macrinus vs Mec. menapis
FST
DXY
Mec. macrinus
Mec. menapis
π
01
02
03
04
05
06
07
08
09
10
11
12
13
14
Z
Chromosome
B
P-value: 2.20e-04, Z-score: 4.142
P-value: 2.00e-05, Z-score: 4.629
P-value: 3.86e-03, Z-score: -2.673
P-value: 5.80e-04, Z-score: 3.243
EVPERM
EVOBS
FST
DXY
π

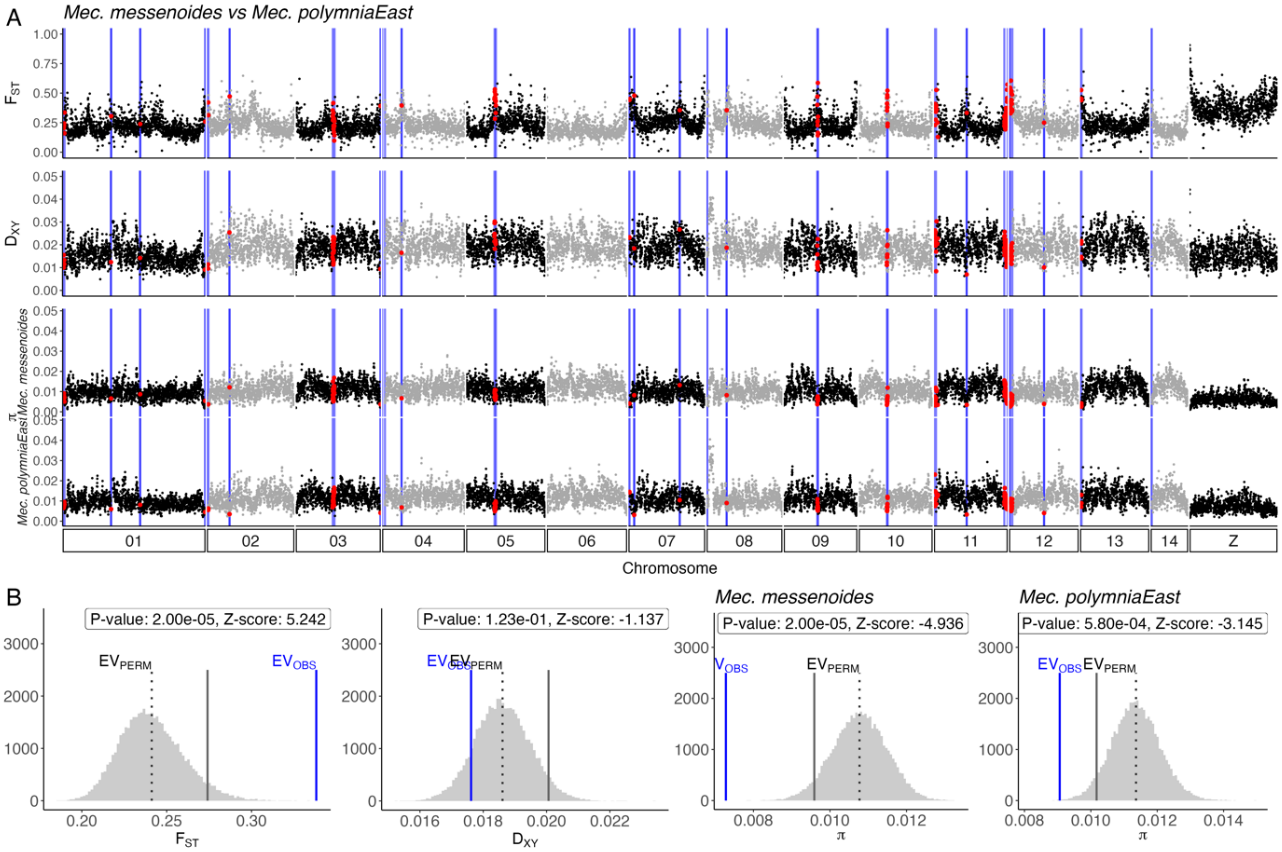
A
Mec. messenoides vs Mec. polymniaEast
FST
DXY
Mec. messenoides
Mec. polymniaEast
π
01
02
03
04
05
06
07
08
09
10
11
12
13
14
Z
Chromosome
B
Mec. messenoides
Mec. polymniaEast
P-value: 2.00e-05, Z-score: 5.242
P-value: 1.23e-01, Z-score: -1.137
P-value: 2.00e-05, Z-score: -4.936
P-value: 5.80e-04, Z-score: -3.145
EVPERM
EVOBS
FST
DXY
π

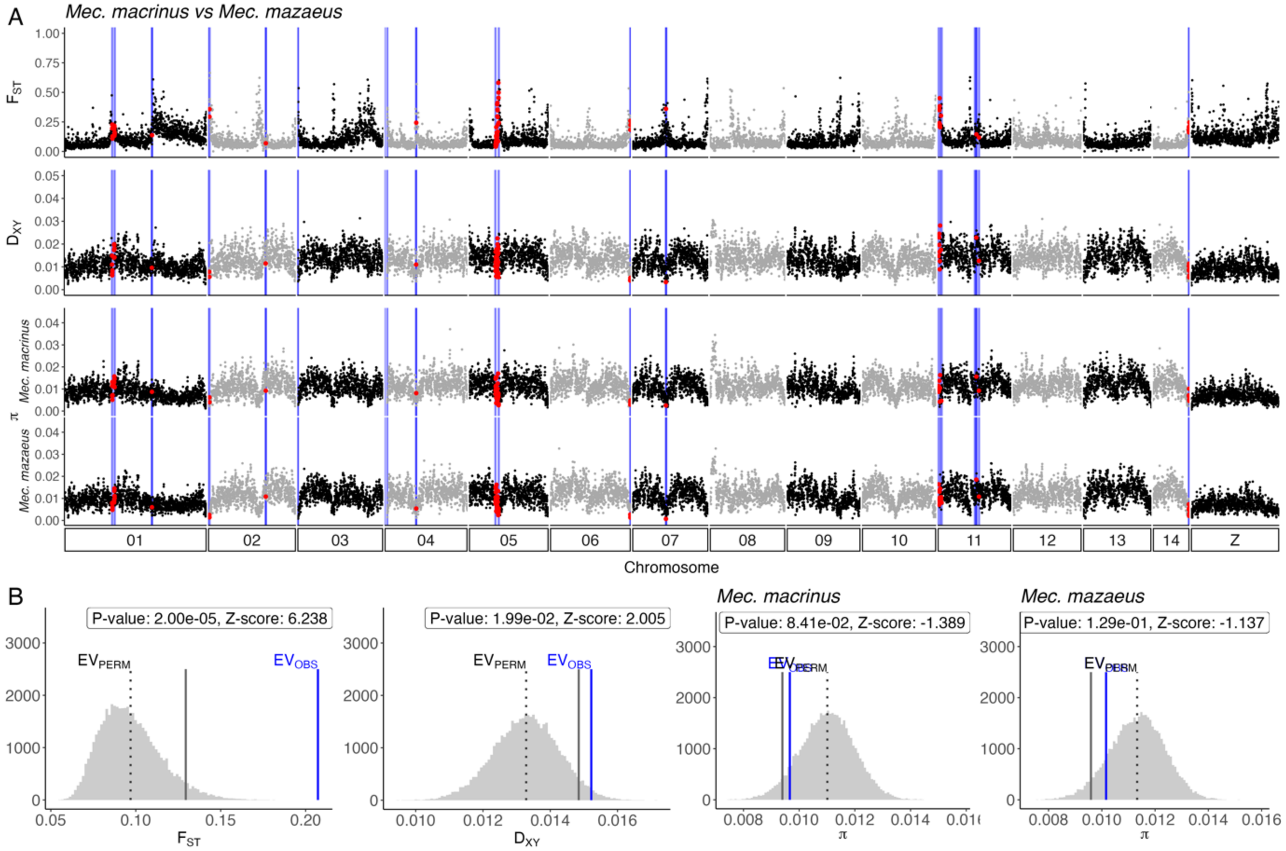
A
Mec. macrinus vs Mec. mazaeus
FST
DXY
π
Mec. macrinus
Mec. mazaeus
01 02 03 04 05 06 07 08 09 10 11 12 13 14 Z
Chromosome
B
P-value: 2.00e-05, Z-score: 6.238
P-value: 1.99e-02, Z-score: 2.005
Mec. macrinus
P-value: 8.41e-02, Z-score: -1.389
Mec. mazaeus
P-value: 1.29e-01, Z-score: -1.137
EVPERM
EVOBS
FST
DXY
π

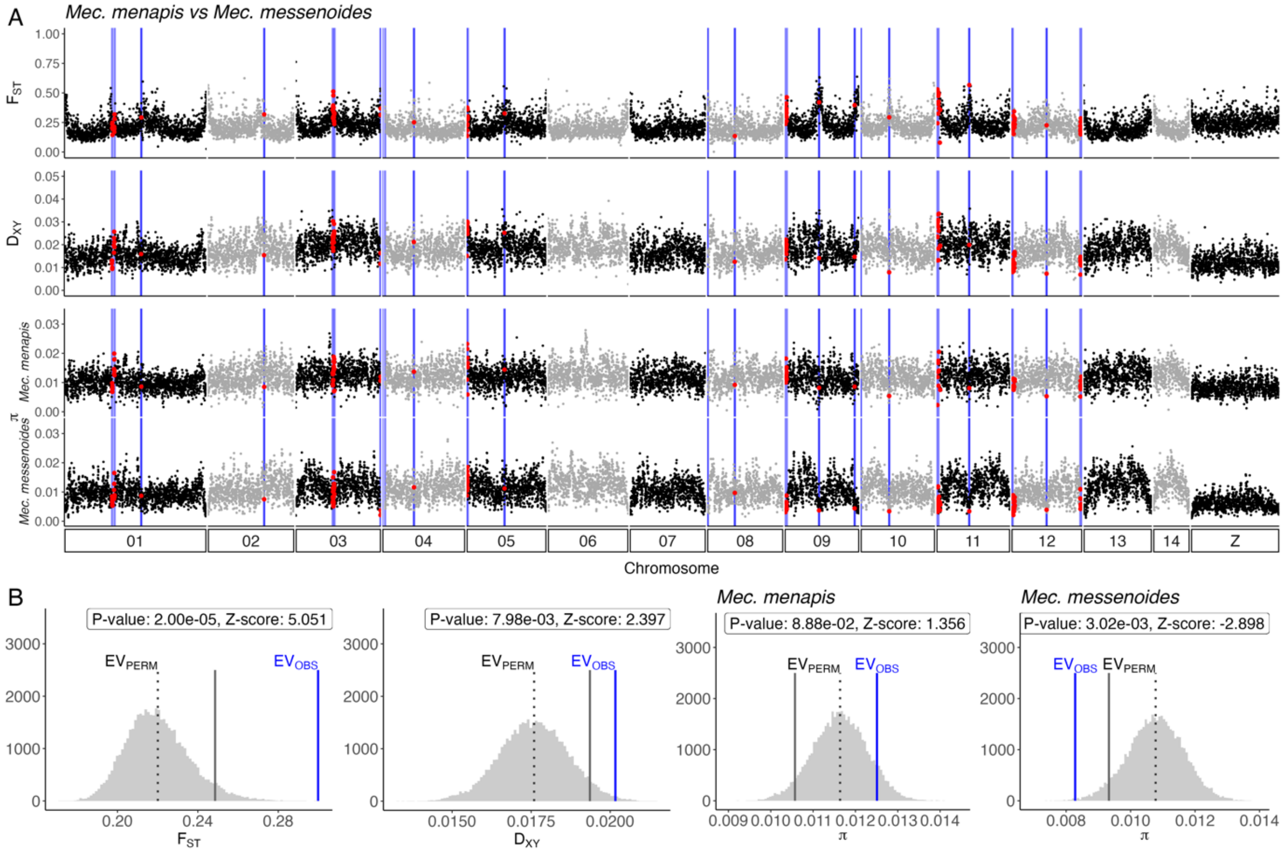
A
Mec. menapis vs Mec. messenoides
FST
DXY
π
Mec. menapis
Mec. messenoides
01 02 03 04 05 06 07 08 09 10 11 12 13 14 Z
Chromosome
B
P-value: 2.00e-05, Z-score: 5.051
P-value: 7.98e-03, Z-score: 2.397
Mec. menapis
P-value: 8.88e-02, Z-score: 1.356
Mec. messenoides
P-value: 3.02e-03, Z-score: -2.898
EVPERM
EVOBS
FST
DXY
π

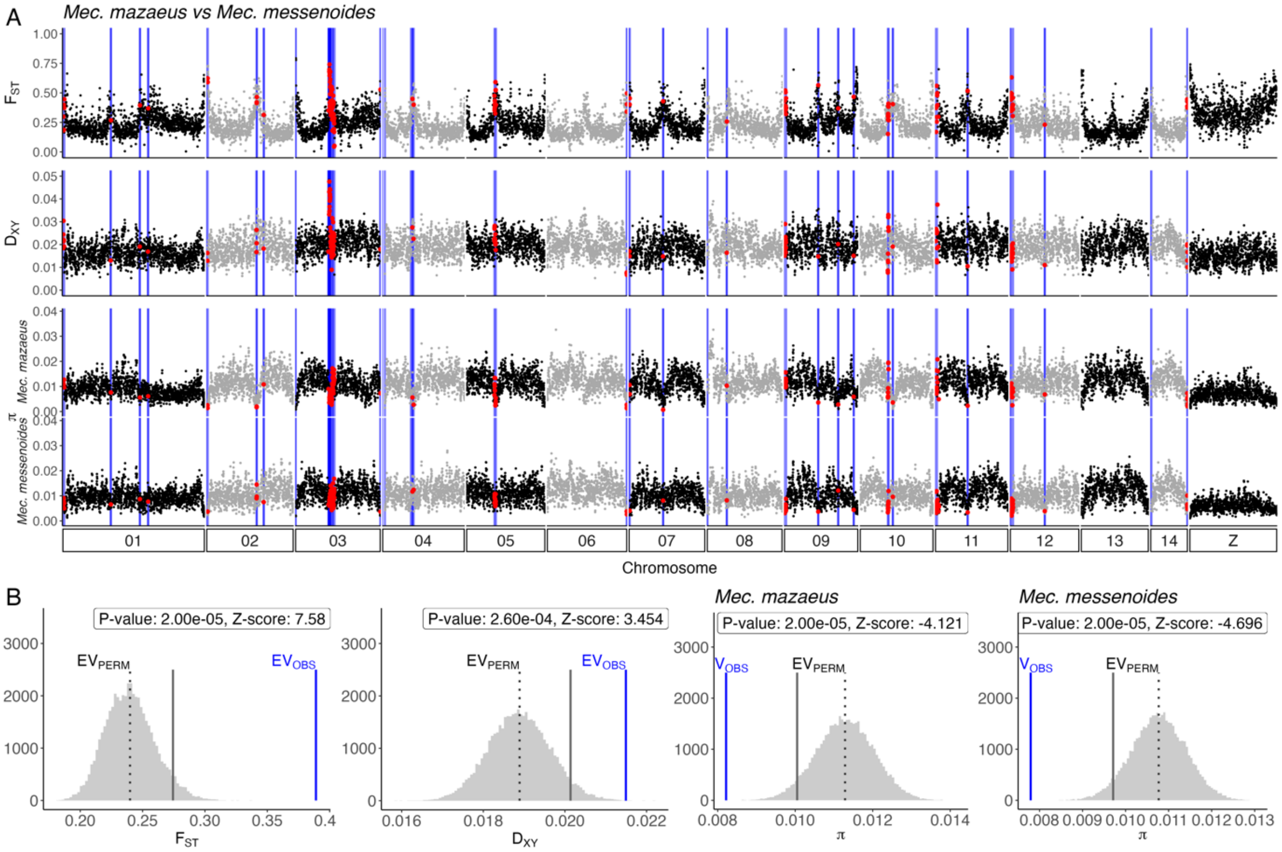
A
Mec. mazaeus vs Mec. messenoides
FST
DXY
π
Mec. mazaeus
Mec. messenoides
01
02
03
04
05
06
07
08
09
10
11
12
13
14
Z
Chromosome
B
P-value: 2.00e-05, Z-score: 7.58
P-value: 2.60e-04, Z-score: 3.454
Mec. mazaeus
P-value: 2.00e-05, Z-score: -4.121
Mec. messenoides
P-value: 2.00e-05, Z-score: -4.696
EVPERM
EVOBS
VOBS
FST
DXY
π

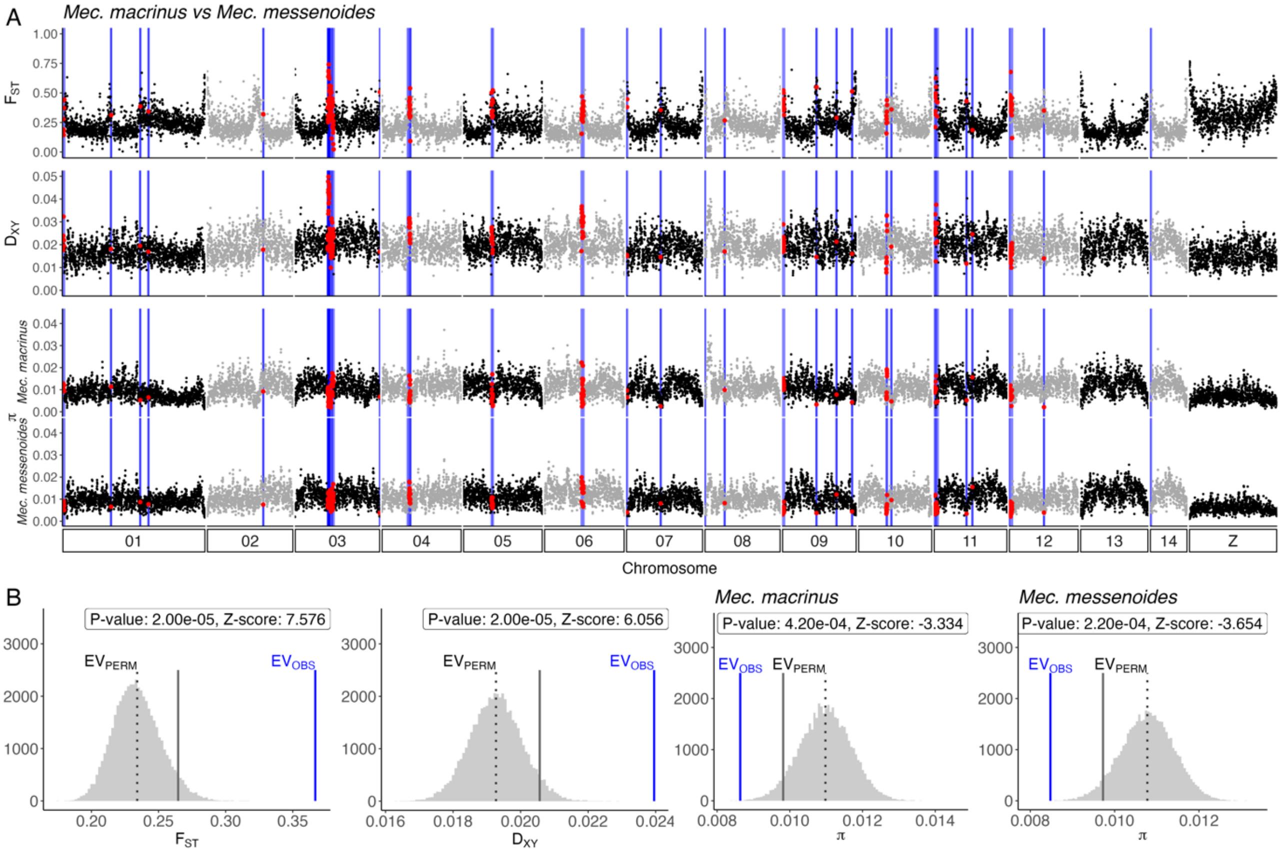
A
Mec. macrinus vs Mec. messenoides
FST
DXY
π
Mec. macrinus
Mec. messenoides
01
02
03
04
05
06
07
08
09
10
11
12
13
14
Z
Chromosome
B
P-value: 2.00e-05, Z-score: 7.576
P-value: 2.00e-05, Z-score: 6.056
Mec. macrinus
P-value: 4.20e-04, Z-score: -3.334
Mec. messenoides
P-value: 2.20e-04, Z-score: -3.654
EVPERM
EVOBS
FST
DXY
π

**Fig. S20a. Genome scans with breakpoints**

*A) Population genomics statistics per 20kb window across the genome for $F_{ST}$, $D_{XY}$ for each species comparison and $\pi$ for each species within* Mechanitis*. Grey and black dots are coloured by chromosome and red dots represent windows located in breakpoints.*

*B) Randomisation test for each statistic across the genome with P-value at 5% significance level (lesser than or greater than randomised sample, grey solid line represents the threshold for the primary alternative hypothesis). Z-score in label. Mean observed population statistic within breakpoint regions (blue, $EV_{OBS}$) and permutation mean (dotted grey line, $EV_{PERM}$) for $F_{ST}$, $D_{XY}$ and $\pi$.*

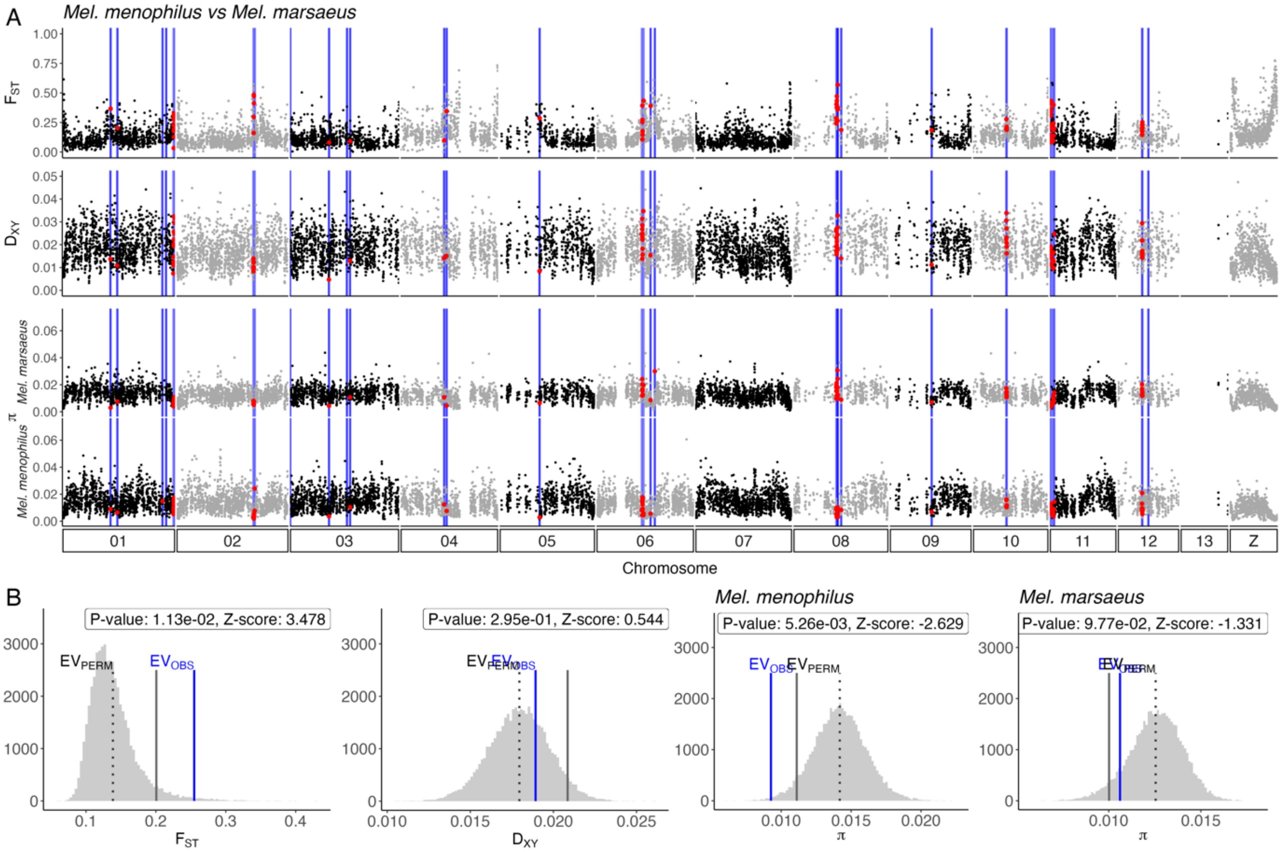
A
Mel. menophilus vs Mel. marsaeus
FST
DXY
π
Mel. marsaeus
Mel. menophilus
01
02
03
04
05
06
07
08
09
10
11
12
13
Z
Chromosome
B
P-value: 1.13e-02, Z-score: 3.478
EVPERM
EVOBS
FST
P-value: 2.95e-01, Z-score: 0.544
DXY
Mel. menophilus
P-value: 5.26e-03, Z-score: -2.629
π
Mel. marsaeus
P-value: 9.77e-02, Z-score: -1.331
π

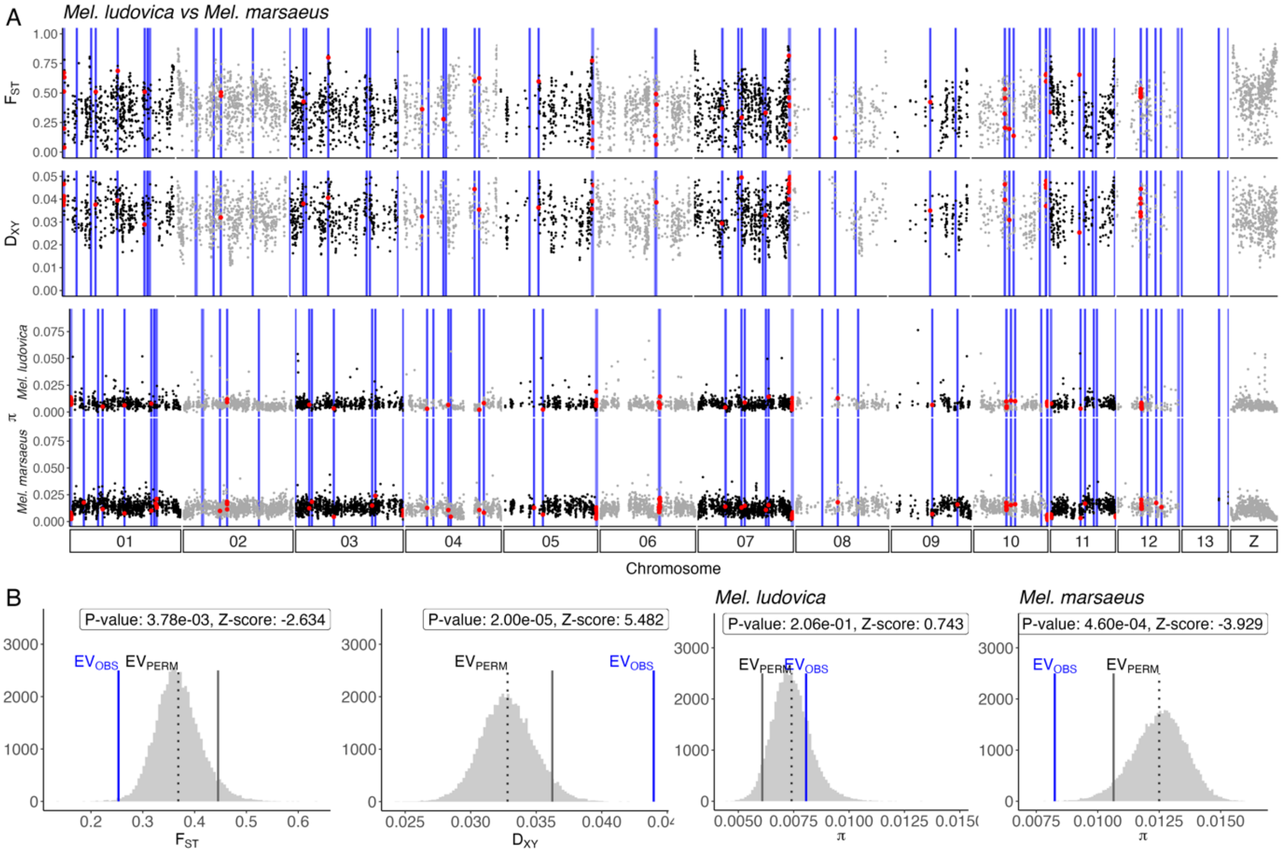
A
Mel. ludovica vs Mel. marsaeus
FST
DXY
π
Mel. ludovica
Mel. marsaeus
01
02
03
04
05
06
07
08
09
10
11
12
13
Z
Chromosome
B
P-value: 3.78e-03, Z-score: -2.634
EVOBS
EVPERM
FST
P-value: 2.00e-05, Z-score: 5.482
DXY
Mel. ludovica
P-value: 2.06e-01, Z-score: 0.743
π
Mel. marsaeus
P-value: 4.60e-04, Z-score: -3.929
π

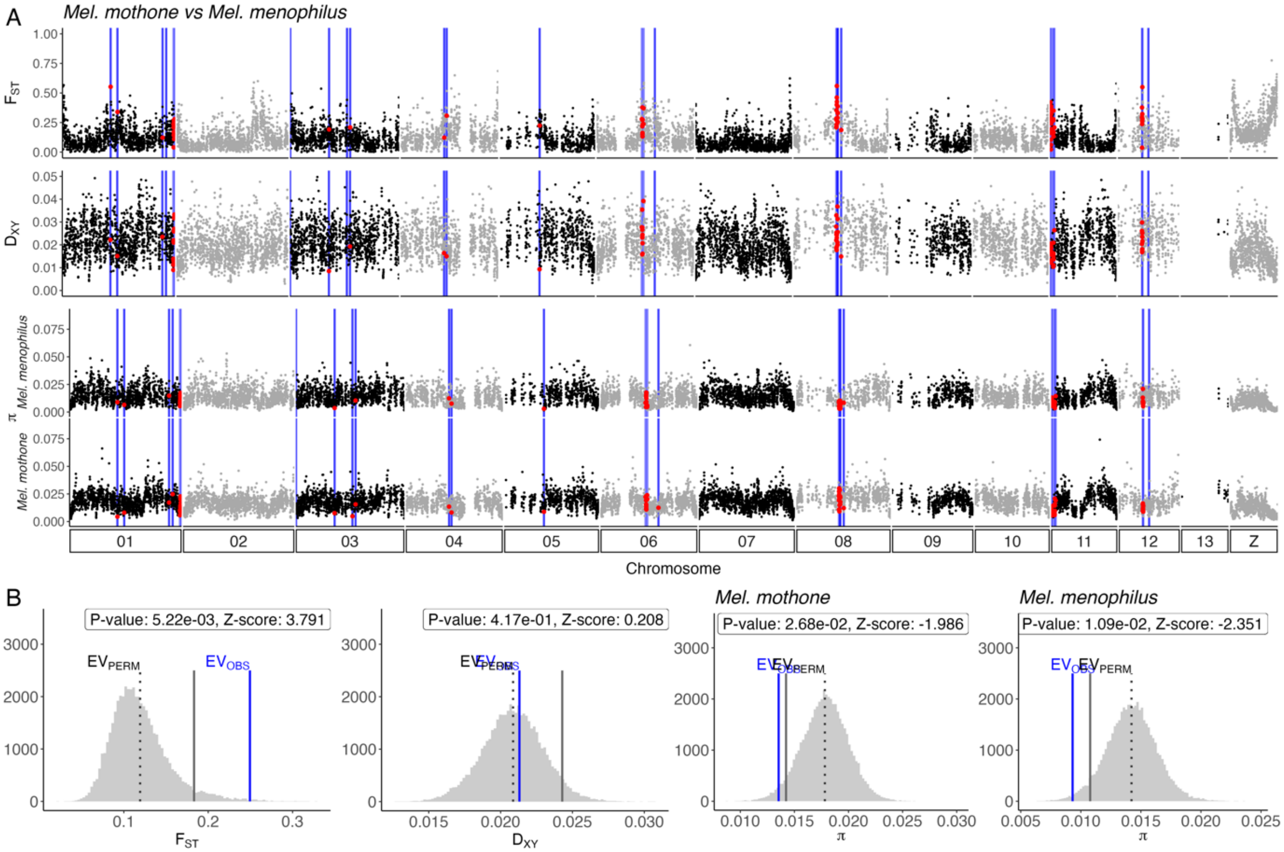
A
Mel. mothone vs Mel. menophilus
FST
DXY
π
Mel. menophilus
Mel. mothone
01
02
03
04
05
06
07
08
09
10
11
12
13
Z
Chromosome
B
P-value: 5.22e-03, Z-score: 3.791
P-value: 4.17e-01, Z-score: 0.208
Mel. mothone
P-value: 2.68e-02, Z-score: -1.986
Mel. menophilus
P-value: 1.09e-02, Z-score: -2.351
EVPERM
EVOBS
FST
DXY
π

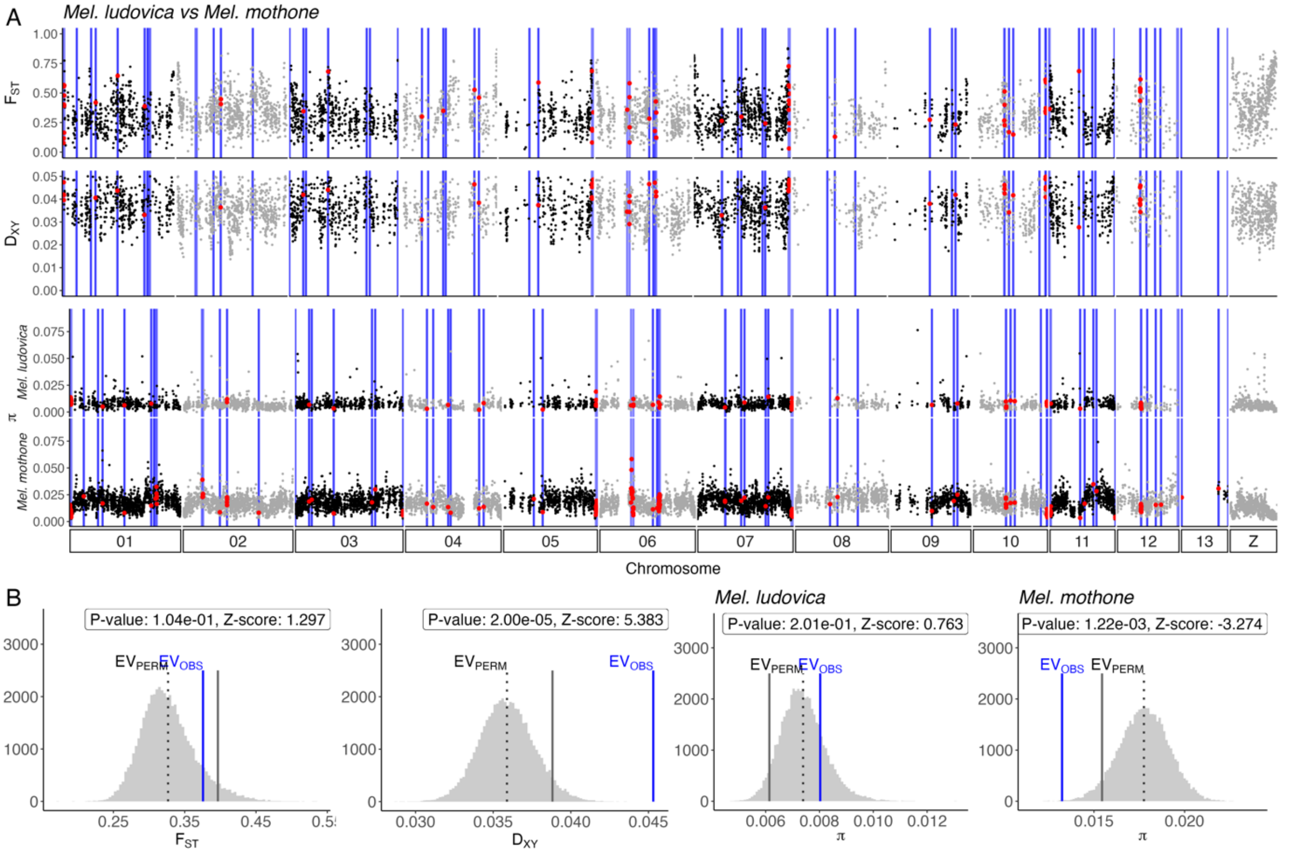
A
Mel. ludovica vs Mel. mothone
FST
DXY
π
Mel. ludovica
Mel. mothone
01
02
03
04
05
06
07
08
09
10
11
12
13
Z
Chromosome
B
P-value: 1.04e-01, Z-score: 1.297
P-value: 2.00e-05, Z-score: 5.383
Mel. ludovica
P-value: 2.01e-01, Z-score: 0.763
Mel. mothone
P-value: 1.22e-03, Z-score: -3.274
EVPERM
EVOBS
FST
DXY
π

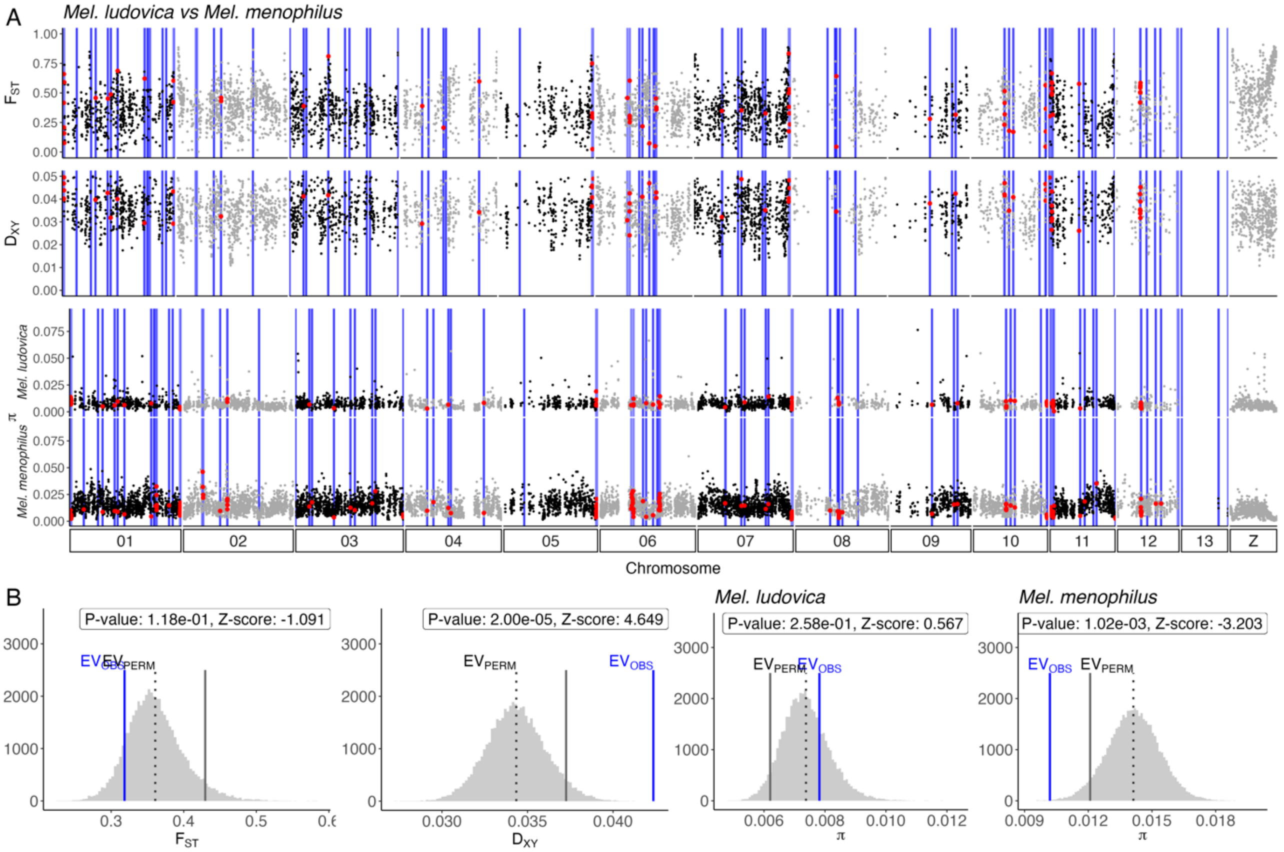
A
Mel. ludovica vs Mel. menophilus
F_ST
D_XY
Mel. ludovica
π
Mel. menophilus
01
02
03
04
05
06
07
08
09
10
11
12
13
Z
Chromosome
B
P-value: 1.18e-01, Z-score: -1.091
P-value: 2.00e-05, Z-score: 4.649
Mel. ludovica
P-value: 2.58e-01, Z-score: 0.567
Mel. menophilus
P-value: 1.02e-03, Z-score: -3.203
EV_OBS
EV_PERM
F_ST
D_XY
π

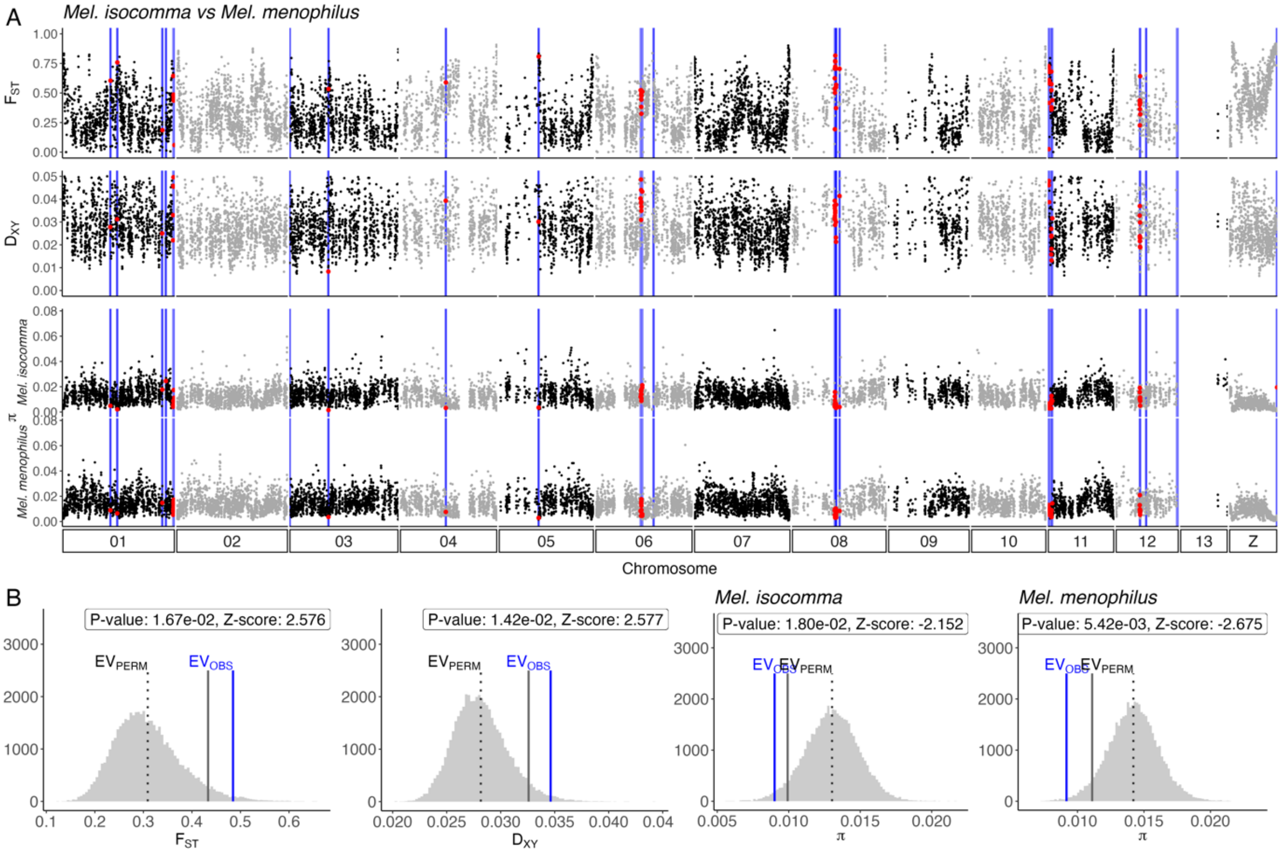
A
Mel. isocomma vs Mel. menophilus
F_ST
D_XY
Mel. isocomma
π
Mel. menophilus
01
02
03
04
05
06
07
08
09
10
11
12
13
Z
Chromosome
B
P-value: 1.67e-02, Z-score: 2.576
P-value: 1.42e-02, Z-score: 2.577
Mel. isocomma
P-value: 1.80e-02, Z-score: -2.152
Mel. menophilus
P-value: 5.42e-03, Z-score: -2.675
EV_PERM
EV_OBS
F_ST
D_XY
π

*Mel. ludovica vs Mel. isocomma*

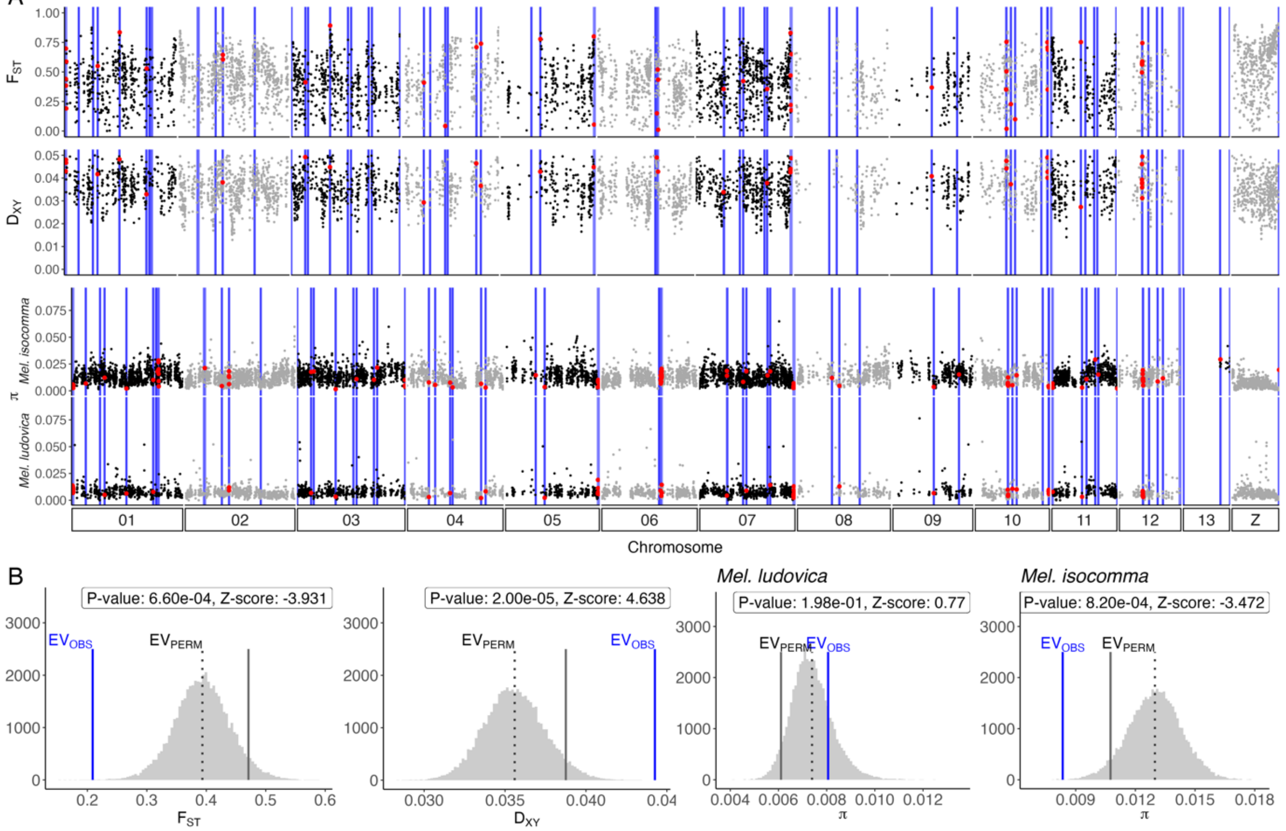


*Mel. isocomma vs Mel. marsaeus*

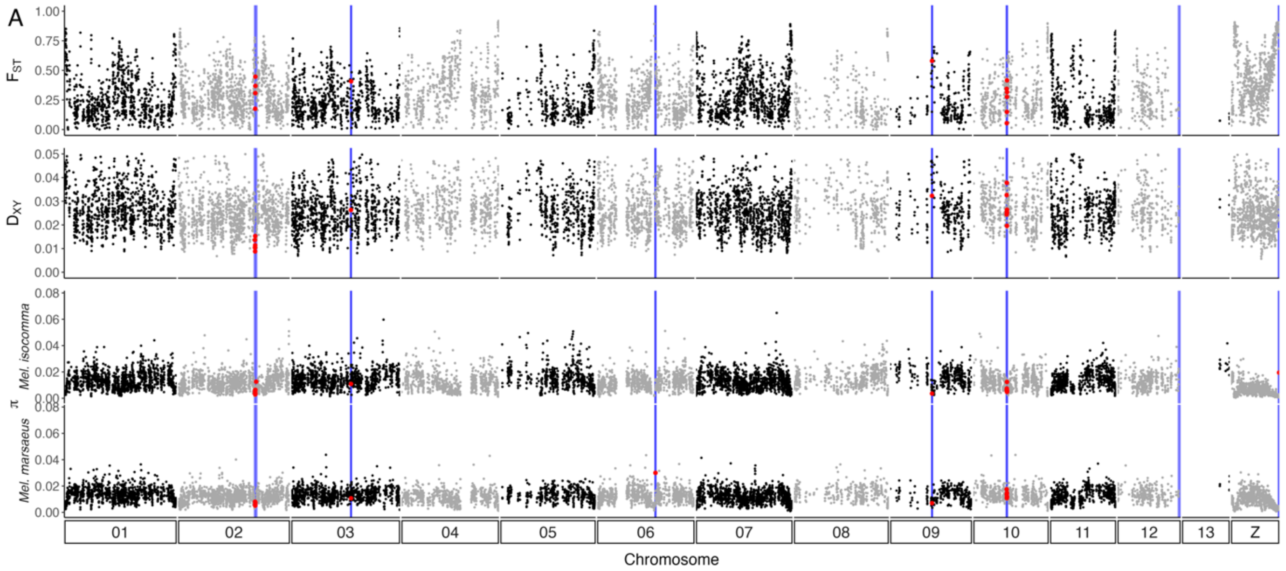

*Mel. isocomma vs Mel. mothone*

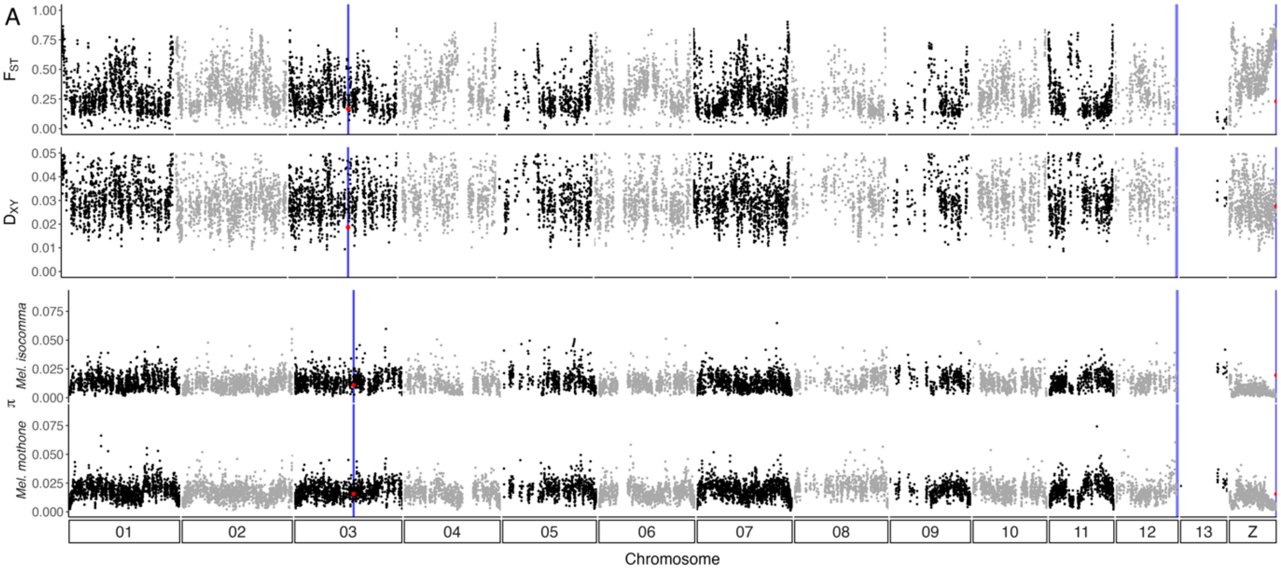


*Mel. Mothone vs Mel. marsaeus*

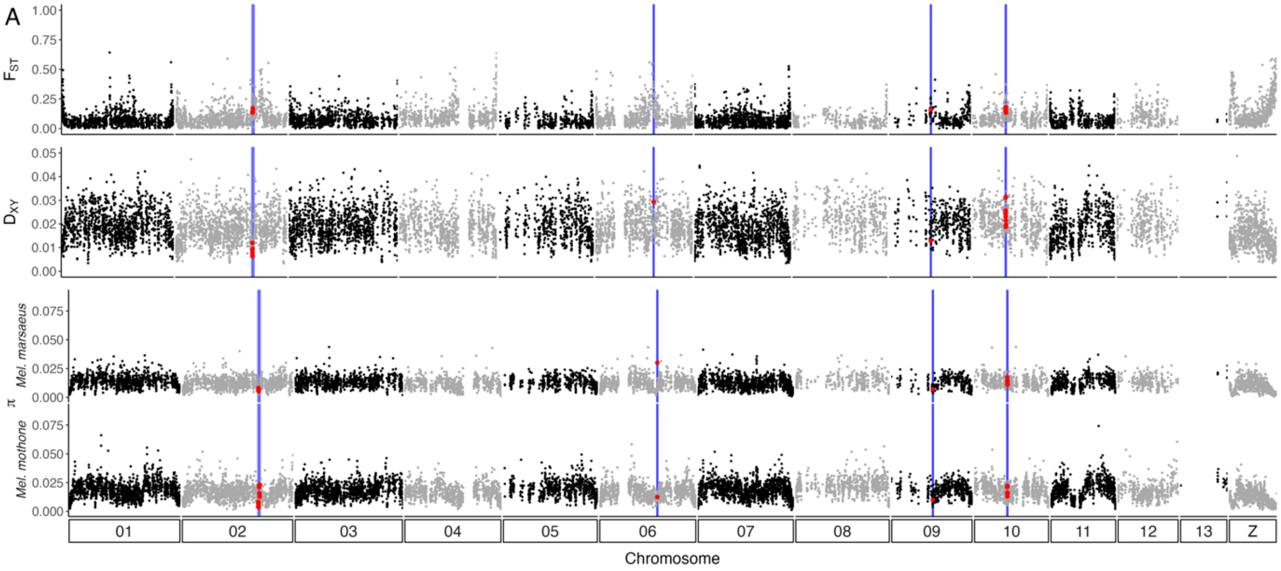


**Fig. S20b. Genome scans with breakpoints**

*A) Population genomics statistics per 20kb window across the genome for $F_{ST}$, $D_{XY}$ for each species comparison and $\pi$ for each species within* Melinaea*. Grey and black dots are coloured by chromosome and red dots represent windows located in breakpoints.*
*B) Randomisation test for each statistic across the genome with P-value at 5% significance level (lesser than or greater than randomised sample, grey solid line represents the threshold for the primary alternative hypothesis). Z-score in label. Mean observed population statistic within breakpoint regions (blue, $EV_{OBS}$) and permutation mean (dotted grey line, $EV_{PERM}$) for $F_{ST}$, $D_{XY}$ and $\pi$.*

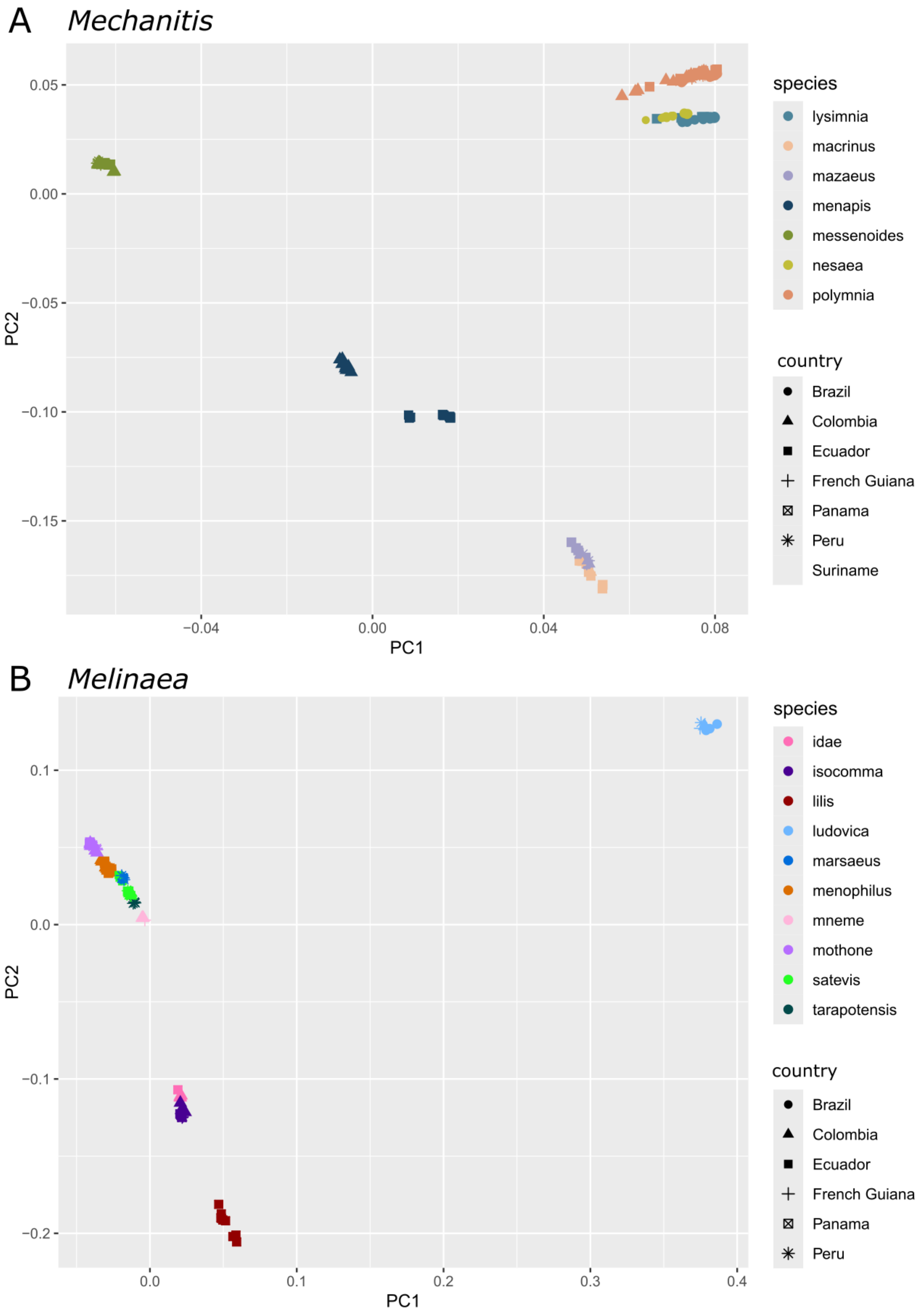


**Fig. S21. PCA for *Mechanitis* and *Melinaea***

*PCA from all the individuals for* Mechanitis *(A) and* Melinaea *(B). The colours indicate species, and the shapes indicate countries.*

**Supplementary Table 1: Sequenced individuals. Information on all individuals resequenced with Illumina.**

sex without asterisk: based on photos of hindwings (with bristles = male; without = female)

sex with asterisk: based on W and Z chromosome depth compared to autosome depth (for Mechanitis: W close to 0 and Z equal to autosome = male; W > 0 (usually half of depth Z) and Z half of autosome = female; for Melinaea: W < autosome and Z > autosome = male; W > autosome and Z < autosome = female. NOTE: take with some caution, there is a lot of variety)

| ID (sequencing) | Original ID | full name (ID before reclassification) | full name (after reclassification) | Dsuite/TWISST | Phylogeny | Country | Year | Depth | latitude | longitude | Sex | Location |
|---|---|---|---|---|---|---|---|---|---|---|---|---|
| CAM072004 | CAM072004 | *Eutresis hypereia* | *Eut. hypereia* | Outgroup (Mel.) | include | Ecuador | NA | 4.5897 | -0.730204 | -77.773665 | male | Narupa Birding |
| CAM028090 | CAM028090 | *Forbestra equicola equicoloides* | *For. equicola equiloides* | Outgroup (Mec.) | Include | Ecuador | 2019 | 8.39578 | -1.1155667 | -77.77833 | female* | Apuya km2.7, Napo |
| CAM028271 | CAM028271 | *Forbestra equicola equicoloides* | *For. equicola equiloides* | Outgroup (Mec.) | Include | Ecuador | 2019 | 8.54284 | -0.9484307 | -77.59437 | male* | Rio Pusuno, Napo |
| CAM028327 | CAM028327 | *Forbestra equicola equicoloides* | *For. equicola equiloides* | Outgroup (Mec.) | Include | Ecuador | 2019 | 8.28167 | NA | NA | male* | Yutzupino 6.3 km, Napo |
| CAM028193 | CAM028193 | *Forbestra olivencia juntana* | *For. olivencia juntana* | Outgroup (Mec.) | Include | Ecuador | 2019 | 8.40125 | -0.9484307 | -77.59437 | male* | Rio Pusuno, Napo |
| CAM028312 | CAM028312 | *Forbestra olivencia juntana* | *For. olivencia juntana* | Outgroup (Mec.) | Include | Ecuador | 2019 | 8.32118 | -0.9484307 | -77.59437 | female* | Rio Pusuno, Napo |
| CAM070582 | CAM070582 | *Forbestra olivencia juntana* | *For. olivencia juntana* | Outgroup (Mec.) | Include | Ecuador | 2022 | 24.5779 | -1.1155667 | -77.77833 | male | Apuya km2.7, Napo |
| CAM070479 | CAM070479 | *Forbestra proceris* | *For. proceris* | Outgroup (Mec.) | Include | Ecuador | 2022 | 21.0066 | -1.09759 | -77.58389 | female | San Pedro de Arajuno |
| CAM070480 | CAM070480 | *Forbestra proceris* | *For. proceris* | Outgroup (Mec.) | Include | Ecuador | 2022 | 18.6076 | -1.09759 | -77.58389 | female | San Pedro de Arajuno |
| CAM070483 | CAM070483 | *Forbestra proceris* | *For. proceris* | Outgroup (Mec.) | Include | Ecuador | 2022 | 18.8403 | -1.09759 | -77.58389 | male | San Pedro de Arajuno |
| CAM036283 | CAM036283 | *Mechanitis lysimnia lysimnia* | *Mec. lysimnia lysimnia* | lysimniaBraz | exclude | Brazil | 2020 | 9.41379 | -15.406724 | -39.54846 | female | Serra Bonita |
| CAM036340 | CAM036340 | *Mechanitis lysimnia lysimnia* | *Mec. lysimnia lysimnia* | lysimniaBraz | exclude | Brazil | 2020 | 11.7506 | -15.406724 | -39.54846 | male | Serra Bonita |
| CAM036460 | CAM036460 | *Mechanitis lysimnia lysimnia* | *Mec. lysimnia lysimnia* | lysimniaBraz | exclude | Brazil | 2020 | 10.7128 | -22.437925 | -44.61271 | female | Itatiaia |
| CAM036461 | CAM036461 | *Mechanitis lysimnia lysimnia* | *Mec. lysimnia lysimnia* | lysimniaBraz | exclude | Brazil | 2020 | 12.1488 | -22.437925 | -44.61271 | female | Itatiaia |
| CAM036544 | CAM036544 | *Mechanitis lysimnia lysimnia* | *Mec. lysimnia lysimnia* | lysimniaBraz | exclude | Brazil | 2020 | 9.71835 | -23.230809 | -46.92723 | male | Japi |
| CAM036548 | CAM036548 | *Mechanitis lysimnia lysimnia* | *Mec. lysimnia lysimnia* | lysimniaBraz | exclude | Brazil | 2020 | 8.97937 | -23.230809 | -46.92723 | male | Japi |
| LBR0211 | LBR0211 | *Mechanitis lysimnia lysimnia* | *Mec. lysimnia lysimnia* | lysimniaBraz | exclude | Brazil | 2011 | 5.83743 | -22.656793 | -52.85771 | male* | Parana, Diamante do Norte |
| MCN001 | MCN001 | *Mechanitis lysimnia lysimnia* | *Mec. lysimnia lysimnia* | lysimniaBraz | exclude | Brazil | 2023 | 13.8941 | NA | NA | female* | Bacchus, RJ |
| MCN002 | MCN002 | *Mechanitis lysimnia lysimnia* | *Mec. lysimnia lysimnia* | lysimniaBraz | exclude | Brazil | 2023 | 14.5659 | NA | NA | female* | Bacchus, RJ |
| MCN003 | MCN003 | *Mechanitis lysimnia lysimnia* | *Mec. lysimnia lysimnia* | lysimniaBraz | include | Brazil | 2023 | 15.7072 | NA | NA | female* | Bacchus, RJ |
| MCN004 | MCN004 | *Mechanitis lysimnia lysimnia* | *Mec. lysimnia lysimnia* | lysimniaBraz | exclude | Brazil | 2023 | 14.1557 | NA | NA | female* | Bacchus, RJ |
| MCN005 | MCN005 | *Mechanitis lysimnia lysimnia* | *Mec. lysimnia lysimnia* | lysimniaBraz | exclude | Brazil | 2023 | 13.1734 | NA | NA | male* | Bacchus, RJ |
| MCN006 | MCN006 | *Mechanitis lysimnia lysimnia* | *Mec. lysimnia lysimnia* | lysimniaBraz | exclude | Brazil | 2023 | 14.9662 | NA | NA | female* | Bacchus, RJ |
| MCN007 | MCN007 | *Mechanitis lysimnia lysimnia* | *Mec. lysimnia lysimnia* | lysimniaBraz | exclude | Brazil | NA | 13.7085 | NA | NA | female* | Camcan, BA |
| MCN008 | MCN008 | *Mechanitis lysimnia lysimnia* | *Mec. lysimnia lysimnia* | lysimniaBraz | include | Brazil | 2015 | 15.3574 | NA | NA | female* | Camcan, BA |
| MCN009 | MCN009 | *Mechanitis lysimnia lysimnia* | *Mec. lysimnia lysimnia* | lysimniaBraz | include | Brazil | 2021 | 17.1437 | -15.361908 | -39.57668 | female* | Serra Bonita |
| MCN010 | MCN010 | *Mechanitis lysimnia lysimnia* | *Mec. lysimnia lysimnia* | lysimniaBraz | include | Brazil | 2021 | 17.4601 | -15.361908 | -39.57668 | male* | Serra Bonita |
| MCN012 | MCN012 | *Mechanitis lysimnia lysimnia* | *Mec. lysimnia lysimnia* | lysimniaBraz | include | Brazil | 2019 | 18.361 | -24.319406 | -46.99659 | male* | Peruibe, SP |
| MCN014 | MCN014 | *Mechanitis lysimnia lysimnia* | *Mec. lysimnia lysimnia* | lysimniaBraz | include | Brazil | 2019 | 15.6518 | -24.319406 | -46.99659 | female* | Peruibe, SP |
| CAM028192 | CAM028192 | *Mechanitis lysimnia roqueensis* | *Mec. lysimnia roqueensis* | lysimniaEcu | include | Ecuador | 2019 | 9.14504 | -0.9484307 | -77.59437 | male* | Rio Pusuno, Napo |
| CAM028197 | CAM028197 | *Mechanitis lysimnia roqueensis* | *Mec. lysimnia roqueensis* | lysimniaEcu | include | Ecuador | 2019 | 8.15159 | -0.9484307 | -77.59437 | male* | Rio Pusuno, Napo |
| CAM028198 | CAM028198 | *Mechanitis lysimnia roqueensis* | *Mec. lysimnia roqueensis* | lysimniaEcu | include | Ecuador | 2019 | 6.75433 | -0.9484307 | -77.59437 | male* | Rio Pusuno, Napo |
| Eli_20338 | 20338 | *Mechanitis lysimnia roqueensis* | *Mec. lysimnia roqueensis* | lysimniaEcu | include | Ecuador | 2005 | 9.38569 | -0.524863 | -76.38418 | male | Anangu, Orellana |
| Eli_20367 | 20367 | *Mechanitis lysimnia roqueensis* | *Mec. lysimnia roqueensis* | lysimniaEcu | include | Ecuador | 2005 | 14.4172 | -0.524863 | -76.38418 | female | Anangu, Orellana |
| Eli_20547 | 20547 | *Mechanitis lysimnia roqueensis* | *Mec. lysimnia roqueensis* | lysimniaEcu | include | Ecuador | 2005 | 14.4104 | -0.524863 | -76.38418 | female | Anangu, Orellana |
| CAM008057 | CAM008057 | *Mechanitis lysimnia utemaia* | *Mec. macrinus utemaia* | macrinus | include | Panama | 2000 | 11.0399 | 8.9569032 | -82.14786 | female | Chiriqui Grande - Almirante 9.5km, Bocas del Toro |
| CAM008092 | CAM008092 | *Mechanitis lysimnia utemaia* | *Mec. macrinus utemaia* | macrinus | include | Panama | 2001 | 11.2882 | 8.9569738 | -82.14413 | female | Bocas del Toro, Chiriquí |
| CAM008432 | CAM008432 | *Mechanitis lysimnia utemaia* | *Mec. macrinus utemaia* | macrinus | include | Panama | 2001 | 9.23806 | 8.7443078 | -82.24991 | male | Quebrada Hornito Trail Fortuna, Chiriquí |
| CAM008476 | CAM008476 | *Mechanitis lysimnia macrinus* | *Mec. macrinus macrinus* | macrinus | include | Panama | 2001 | 8.1879 | 8.713114 | -79.90659 | female | Cerro Campana |
| CAM008822 | CAM008822 | *Mechanitis lysimnia macrinus* | *Mec. macrinus macrinus* | macrinus | include | Panama | 2002 | 8.63882 | 7.8000592 | -77.46355 | female | Cana, around runway area, Darién |
| CAM008975 | CAM008975 | *Mechanitis lysimnia macrinus* | *Mec. macrinus macrinus* | macrinus | include | Panama | 2003 | 9.18807 | 7.6429589 | -78.19232 | female | Rio Piñas campsite, Darién |
| Sal_1986 | M1986 | *Mechanitis lysimnia macrinus* | *Mec. macrinus macrinus* | macrinus | include | Colombia | 2006 | 10.5562 | 4.9645722 | -74.43481 | female | Sasaima, Cundinamarca |
| Sal_4176 | M4176 | *Mechanitis lysimnia macrinus* | *Mec. macrinus macrinus* | macrinus | include | Colombia | 2016 | 12.7209 | 5.6585829 | -74.18148 | male | El Carmen1-Otanche, Boyaca |
| SAN25001776 | LEP-08641 | *Mechanitis lysimnia macrinus* | *Mec. macrinus macrinus* | macrinus | include | Ecuador | NA | 30.1365 | 1.13 | -78.658 | male* | Esmeraldas: Ridge N La Ceiba |
| SAN25001777 | LEP-08646 | *Mechanitis lysimnia macrinus* | *Mec. macrinus macrinus* | macrinus | include | Ecuador | NA | 35.3045 | 1.13 | -78.658 | male* | Esmeraldas: Ridge N La Ceiba |
| CAM042050 | CAM042050 | *Mechanitis mazaeus fallax* | *Mec. mazaeus fallax* | mazaeus | include | Ecuador | 2019 | 9.29747 | -0.993443 | -77.81284 | male | Napo |
| CAM042128 | CAM042128 | *Mechanitis mazaeus fallax* | *Mec. mazaeus fallax* | mazaeus | include | Ecuador | 2019 | 8.63007 | -0.993443 | -77.81284 | female | Napo |
| Eli_mm14_0014 | mm14-14 | *Mechanitis mazaeus fallax* | *Mec. mazaeus fallax* | mazaeus | include | Peru | 2014 | 10.9364 | -6.4805835 | -76.34799 | male | PB km 7.5 |
| Eli_mm15_0887 | mm15-887 | *Mechanitis mazaeus fallax* | *Mec. mazaeus fallax* | mazaeus | include | Peru | 2015 | 12.0112 | -6.4805835 | -76.34799 | female | PB km 7.5 |
| Eli_mm15_0888 | mm15-888 | *Mechanitis mazaeus fallax* | *Mec. mazaeus fallax* | mazaeus | include | Peru | 2015 | 12.2197 | -6.4805835 | -76.34799 | female | PB km 7.5 |
| Mon_2012_172 | Yas_2012_StMo_172 | *Mechanitis mazaeus fallax* | *Mec. mazaeus fallax* | mazaeus | include | Ecuador | 2012 | 16.0838 | -0.5680873 | -75.7582 | male | Yasuni, Napo |
| Mon_2012_173 | Yas_2012_StMo_173 | *Mechanitis mazaeus fallax* | *Mec. mazaeus fallax* | mazaeus | include | Ecuador | 2012 | 11.7799 | -0.5680873 | -75.7582 | female | Yasuni, Napo |
| Mon_2012_197 | Yas_2012_StMo_197 | *Mechanitis mazaeus fallax* | *Mec. mazaeus fallax* | mazaeus | include | Ecuador | 2012 | 13.8087 | -0.5680873 | -75.7582 | male | Yasuni, Napo |
| Sal_1056 | M1056 | *Mechanitis mazaeus fallax* | *Mec. mazaeus fallax* | mazaeus | include | Colombia | 2004 | 12.1138 | 4.1659898 | -73.68238 | female | Buenavista, Meta |
| Sal_1298 | M1298 | *Mechanitis mazaeus fallax* | *Mec. mazaeus fallax* | mazaeus | include | Colombia | 2005 | 13.2132 | 1.9097464 | -75.15582 | female | Puerto Rico, Caquetá |
| Sal_5319 | M5319 | *Mechanitis mazaeus fallax* | *Mec. mazaeus fallax* | mazaeus | include | Colombia | 2018 | 11.0242 | -1.3220609 | -69.57811 | female | Vereda Angostura - La Pedrera, Amazonas |
| Sal_5427 | M5427 | *Mechanitis mazaeus fallax* | *Mec. mazaeus fallax* | mazaeus | include | Colombia | 2018 | 12.2512 | -1.3220609 | -69.57811 | female | Sendero Bacurí - La Pedrera, Amazonas |
| Sal_2064 | M2064 | *Mechanitis menapis caribensis* | *Mec. menapis caribensis* | menapis | include | Colombia | 2006 | 11.0798 | 6.097464 | -73.32032 | female | Tolota, Santander |
| Sal_2066 | M2066 | *Mechanitis menapis caribensis* | *Mec. menapis caribensis* | menapis | include | Colombia | 2006 | 12.0029 | 6.097464 | -73.32032 | female | Tolota, Santander |
| Sal_5127 | M5127 | *Mechanitis menapis caribensis* | *Mec. menapis caribensis* | menapis | exclude | Colombia | NA | 10.263 | 4.481846 | -75.76605 | male | La paila, Cucuta |
| Sal_5640 | M5640 | *Mechanitis menapis caribensis* | *Mec. menapis caribensis* | menapis | include | Colombia | 2020 | 12.8262 | 5.6803162 | -73.52136 | female | La Rosita -casa familia Valero, Boyaca |
| Sal_5641 | M5641 | *Mechanitis menapis caribensis* | *Mec. menapis caribensis* | menapis | include | Colombia | 2020 | 12.2877 | 5.6803162 | -73.52136 | female | La Rosita -casa familia Valero, Boyaca |
| Sal_1938 | M1938 | *Mechanitis menapis dariensis* | *Mec. menapis dariensis* | menapis | include | Colombia | 2006 | 10.3853 | 3.6105042 | -76.59797 | female | Bitaco, Valle del Cauca |
| Sal_3342 | M3342 | *Mechanitis menapis dariensis* | *Mec. menapis dariensis* | menapis | include | Colombia | 2011 | 15.2219 | 6.5593804 | -75.8256 | female | Santafé de Antioquia, Antioquia |
| CAM070759 | CAM070759 | *Mechanitis menapis mantineus* | *Mec. menapis mantineus* | menapis | include | Ecuador | 2022 | 27.525 | 0.1659866 | -78.87828 | male | Mashpi Lodge, Azuay |
| SAN25001770 | LEP-55153 | *Mechanitis menapis mantineus* | *Mec. menapis mantineus* | menapis | include | Ecuador | NA | 29.9997 | 0.739 | -78.242 | female* | Imbabura: Bosque de Paz |
| SAN25001771 | LEP-55155 | *Mechanitis menapis mantineus* | *Mec. menapis mantineus* | menapis | include | Ecuador | NA | 31.3096 | 0.888 | -78.438 | female* | Carchi: Lita, Ridge east of Rio Baboso |
| SAN25001772 | LEP-58365 | *Mechanitis menapis mantineus* | *Mec. menapis mantineus* | menapis | include | Ecuador | NA | 24.2407 | 1.13 | -78.658 | female* | Esmeraldas: Ridge N La Ceiba |
| SAN25001773 | LEP-58366 | *Mechanitis menapis mantineus* | *Mec. menapis mantineus* | menapis | include | Ecuador | NA | 26.5063 | 1.13 | -78.658 | female* | Esmeraldas: Ridge N La Ceiba |
| Sal_5638 | M5638 | *Mechanitis menapis menapis* | *Mec. menapis menapis* | menapis | include | Colombia | 2020 | 11.6516 | 5.6803162 | -73.52136 | female | La Rosita -casa familia Valero, Boyaca |
| Sal_4723 | M4723 | *Mechanitis menapis occasiva* | *Mec. menapis occasiva* | menapis | include | Colombia | 2017 | 15.8544 | 3.3230839 | -76.62916 | female | Pance - Topacio, Valle del Cauca |
| Sal_5681 | M5681 | *Mechanitis menapis occasiva* | *Mec. menapis occasiva* | menapis | include | Colombia | 2020 | 9.7703 | 3.6855664 | -76.52802 | female | Montañitas, Valle del Cauca |
| Sal_5683 | M5683 | *Mechanitis menapis occasiva* | *Mec. menapis occasiva* | menapis | include | Colombia | 2020 | 12.5592 | 3.6855664 | -76.52802 | male | Montañitas, Valle del Cauca |
| CAM008026 | CAM008026 | *Mechanitis menapis saturata* | *Mec. menapis saturata* | menapis | include | Panama | 2000 | 10.1695 | 8.8340568 | -82.71632 | female | Cafe Duran Finca near Santa Clara, Chiriquí |
| CAM008324 | CAM008324 | *Mechanitis menapis saturata* | *Mec. menapis saturata* | menapis | include | Panama | 2001 | 12.5072 | 8.7443078 | -82.24991 | male | Quebrada Hornito Trail Fortuna, Chiriquí |
| CAM008325 | CAM008325 | *Mechanitis menapis saturata* | *Mec. menapis saturata* | menapis | include | Panama | 2001 | 13.4405 | 8.7443078 | -82.24991 | male | Quebrada Hornito Trail Fortuna, Chiriquí |
| CAM008423 | CAM008423 | *Mechanitis menapis saturata* | *Mec. menapis saturata* | menapis | include | Panama | 2001 | 11.8244 | 8.7829144 | -82.63632 | female | Cafe Vista Hermosa Fortuna, Chiriquí |
| CAM077079 | M6152 | *Mechanitis messenoides deceptus* | *Mec. messenoides deceptus* | messenoides | include | Colombia | 2022 | 32.2813 | 1.147492 | -76.64801 | female* | Mocoa, Putumayo |
| CAM077080 | M6153 | *Mechanitis messenoides deceptus* | *Mec. messenoides deceptus* | messenoides | include | Colombia | 2022 | 43.3148 | 1.147492 | -76.64801 | male* | Mocoa, Putumayo |
| Mon_2011_124 | Yas_2011_StMo_124 | *Mechanitis messenoides deceptus* | *Mec. messenoides deceptus* | messenoides | include | Ecuador | 2011 | 15.133 | -0.5680873 | -75.7582 | female | Yasuni, Napo |
| Mon_2011_195 | Yas_2011_StMo_195 | *Mechanitis messenoides deceptus* | *Mec. messenoides deceptus* | messenoides | include | Ecuador | 2011 | 14.7891 | -0.5680873 | -75.7582 | male | Yasuni, Napo |
| Mon_2012_146 | Yas_2012_StMo_146 | *Mechanitis messenoides deceptus* | *Mec. messenoides deceptus* | messenoides | include | Ecuador | 2012 | 19.084 | -0.5680873 | -75.7582 | male | Yasuni, Napo |
| Mon_2012_198 | Yas_2012_StMo_198 | *Mechanitis messenoides deceptus* | *Mec. messenoides deceptus* | messenoides | include | Ecuador | 2012 | 17.194 | -0.5680873 | -75.7582 | female | Yasuni, Napo |
| Mon_2012_243 | Yas_2012_StMo_243 | *Mechanitis messenoides deceptus* | *Mec. messenoides deceptus* | messenoides | include | Ecuador | 2012 | 16.2453 | -0.5680873 | -75.7582 | male | Yasuni, Napo |
| Mon_2012_88 | Yas_2012_StMo_88 | *Mechanitis messenoides deceptus* | *Mec. messenoides deceptus* | messenoides | include | Ecuador | 2012 | 15.2826 | -0.5680873 | -75.7582 | female* | Yasuni, Napo |
| Sal_1360 | M1360 | *Mechanitis messenoides deceptus* | *Mec. messenoides deceptus* | messenoides | include | Colombia | 2005 | 12.7963 | 4.1659898 | -73.68238 | female | Buenavista, Meta |
| Sal_1532 | M1532 | *Mechanitis messenoides deceptus* | *Mec. messenoides deceptus* | messenoides | include | Colombia | 2005 | 12.0195 | 4.1659898 | -73.68238 | female | Buenavista, Meta |
| Mon_2012_219 | Yas_2012_StMo_219 | *Mechanitis messenoides mes/dec* | *Mec. messenoides mes/dec* | messenoides | include | Ecuador | 2012 | 14.8432 | -0.5680873 | -75.7582 | male | Yasuni, Napo |
| CAM077045 | M6102 | *Mechanitis messenoides messenoides* | *Mec. messenoides messenoides* | messenoides | include | Colombia | 2022 | 33.8994 | 1.147492 | -76.64801 | female* | Mocoa, Putumayo |
| CAM077081 | M6154 | *Mechanitis messenoides messenoides* | *Mec. messenoides messenoides* | messenoides | include | Colombia | 2022 | 35.5861 | 1.147492 | -76.64801 | male* | Mocoa, Putumayo |
| CAM077083 | M6156 | *Mechanitis messenoides messenoides* | *Mec. messenoides messenoides* | messenoides | include | Colombia | 2022 | 34.4739 | 1.147492 | -76.64801 | male* | Mocoa, Putumayo |
| CAM077084 | M6157 | *Mechanitis messenoides messenoides* | *Mec. messenoides messenoides* | messenoides | include | Colombia | 2022 | 36.8726 | 1.147492 | -76.64801 | male* | Mocoa, Putumayo |
| CAM077091 | M6158 | *Mechanitis messenoides messenoides* | *Mec. messenoides messenoides* | messenoides | include | Colombia | 2022 | 42.212 | 1.147492 | -76.64801 | male* | Mocoa, Putumayo |
| CAM077092 | M6165 | *Mechanitis messenoides messenoides* | *Mec. messenoides messenoides* | messenoides | include | Colombia | 2022 | 37.6697 | 1.147492 | -76.64801 | male* | Mocoa, Putumayo |
| Mon_2012_220 | Yas_2012_StMo_220 | *Mechanitis messenoides messenoides* | *Mec. messenoides messenoides* | messenoides | include | Ecuador | 2012 | 15.2761 | -0.5680873 | -75.7582 | male | Yasuni, Napo |
| Mon_2012_242 | Yas_2012_StMo_242 | *Mechanitis messenoides messenoides* | *Mec. messenoides messenoides* | messenoides | include | Ecuador | 2012 | 14.9384 | -0.5680873 | -75.7582 | male | Yasuni, Napo |
| Mon_2012_244 | Yas_2012_StMo_244 | *Mechanitis messenoides messenoides* | *Mec. messenoides messenoides* | messenoides | include | Ecuador | 2012 | 14.7535 | -0.5680873 | -75.7582 | male | Yasuni, Napo |
| Mon_2012_90 | Yas_2012_StMo_90 | *Mechanitis messenoides messenoides* | *Mec. messenoides messenoides* | messenoides | include | Ecuador | 2012 | 14.2619 | -0.5680873 | -75.7582 | male | Yasuni, Napo |
| Mon_2012_99 | Yas_2012_StMo_99 | *Mechanitis messenoides messenoides* | *Mec. messenoides messenoides* | messenoides | include | Ecuador | 2012 | 15.419 | -0.5680873 | -75.7582 | female* | Yasuni, Napo |
| 50-02-348 | 50-02-348 | *Mechanitis messenoides phasianita* | *Mec. messenoides phasianita* | messenoides | include | Peru | 2002 | 30.6697 | -6.3353 | -76.2542 | female | Pongo de Caynarachi |
| 55-02-1284 | 55-02-1284 | *Mechanitis messenoides phasianita* | *Mec. messenoides phasianita* | messenoides | include | Peru | 2002 | 28.5735 | -6.2868 | -76.2317 | female | Km-7.2, Pongo-Barranquita |
| 60-02-1290 | 60-02-1290 | *Mechanitis messenoides phasianita* | *Mec. messenoides phasianita* | messenoides | include | Peru | 2002 | 34.3459 | -6.2868 | -76.2317 | female | Km-7.2, Pongo-Barranquita |
| 61-02-1896 | 61-02-1896 | *Mechanitis messenoides phasianita* | *Mec. messenoides phasianita* | messenoides | include | Peru | 2002 | 27.4076 | -6.6247 | -76.4564 | female | Km-30, Tarapoto-Yurimaguas |
| 65-02-477 | 65-02-477 | *Mechanitis messenoides phasianita* | *Mec. messenoides phasianita* | messenoides | include | Peru | 2002 | 30.6185 | -6.4083 | -76.325 | female | Km-28, Tarapoto-Yurimaguas |
| 68-02-677 | 68-02-677 | *Mechanitis messenoides phasianita* | *Mec. messenoides phasianita* | messenoides | include | Peru | 2002 | 31.2007 | -6.4618 | -76.2919 | male | Biodiversidad USM, Km-19, Tarapoto-Yurimaguas |
| CAM036038 | CAM036038 | *Mechanitis lysimnia nesaea* | *Mec. nesaea nesaea* | nesaea | exclude | Brazil | 2020 | 9.94508 | -8.725661 | -35.17961 | female | Saltinho |
| CAM036042 | CAM036042 | *Mechanitis lysimnia nesaea* | *Mec. nesaea nesaea* | nesaea | exclude | Brazil | 2020 | 9.85616 | -8.725661 | -35.17961 | female | Saltinho |
| CAM036060 | CAM036060 | *Mechanitis lysimnia nesaea* | *Mec. nesaea nesaea* | nesaea | exclude | Brazil | 2020 | 10.6923 | -8.725661 | -35.17961 | female | Saltinho |
| CAM036081 | CAM036081 | *Mechanitis lysimnia nesaea* | *Mec. nesaea nesaea* | nesaea | exclude | Brazil | 2020 | 10.131 | -7.925017 | -34.93332 | male | Caetes |
| CAM036084 | CAM036084 | *Mechanitis lysimnia nesaea* | *Mec. nesaea nesaea* | nesaea | include | Brazil | 2020 | 12.0614 | -7.925017 | -34.93332 | male | Caetes |
| CAM036146 | CAM036146 | *Mechanitis lysimnia nesaea* | *Mec. nesaea nesaea* | nesaea | exclude | Brazil | 2020 | 10.224 | -6.730095 | -35.1427 | male | Guaribas |
| CAM036153 | CAM036153 | *Mechanitis lysimnia nesaea* | *Mec. nesaea nesaea* | nesaea | exclude | Brazil | 2020 | 10.5213 | -6.724533 | -35.14482 | female | Guaribas |
| CAM036207 | CAM036207 | *Mechanitis lysimnia nesaea* | *Mec. nesaea nesaea* | nesaea | include | Brazil | 2020 | 12.0259 | -6.724533 | -35.14482 | male | Guaribas |
| CAM036278 | CAM036278 | *Mechanitis lysimnia nesaea* | *Mec. nesaea nesaea* | nesaea | exclude | Brazil | 2020 | 8.94742 | -15.405748 | -39.54844 | male | Serra Bonita |
| CAM036280 | CAM036280 | *Mechanitis lysimnia nesaea* | *Mec. nesaea nesaea* | nesaea | include | Brazil | 2020 | 12.021 | -15.405748 | -39.54844 | male | Serra Bonita |
| CAM036281 | CAM036281 | *Mechanitis lysimnia nesaea* | *Mec. nesaea nesaea* | nesaea | exclude | Brazil | 2020 | 9.48588 | -15.403946 | -39.54858 | female | Serra Bonita |
| CAM036337 | CAM036337 | *Mechanitis lysimnia nesaea* | *Mec. nesaea nesaea* | nesaea | exclude | Brazil | 2020 | 11.0083 | -15.406724 | -39.54846 | male | Serra Bonita |
| MCN011 | MCN011 | *Mechanitis lysimnia nesaea* | *Mec. nesaea nesaea* | nesaea | include | Brazil | 2021 | 17.6524 | -15.361908 | -39.57668 | male* | Serra Bonita |
| MCN015 | MCN015 | *Mechanitis lysimnia nesaea* | *Mec. nesaea nesaea* | nesaea | include | Brazil | 2022 | 13.1977 | -10.685365 | -37.43606 | female* | PNSerra Itabaiana, SE |
| MCN016 | MCN016 | *Mechanitis lysimnia nesaea* | *Mec. nesaea nesaea* | nesaea | include | Brazil | NA | 16.6144 | NA | NA | female* | Areia Branca, SE |
| CAM036418 | CAM036418 | *Mechanitis polymnia casabranca* | *Mec. polymnia casabranca* | polymniaBraz | exclude | Brazil | 2020 | 11.9244 | -22.428577 | -44.61974 | male | Itatiaia |
| CAM036419 | CAM036419 | *Mechanitis polymnia casabranca* | *Mec. polymnia casabranca* | polymniaBraz | exclude | Brazil | 2020 | 10.2508 | -22.437925 | -44.61271 | male | Itatiaia |
| LBR0644 | LBR0644 | *Mechanitis polymnia casabranca* | *Mec. polymnia casabranca* | polymniaBraz | include | Brazil | 2018 | 16.6629 | -18.355049 | -39.84209 | male* | ES, Conceição da Barra, Flona do Rio Preto, Trilha da Lagoa Seca |
| LBR0693 | LBR0693 | *Mechanitis polymnia casabranca* | *Mec. polymnia casabranca* | polymniaBraz | exclude | Brazil | 2018 | 14.5857 | -18.355049 | -39.84209 | male* | ES, Conceição da Barra, Flona do Rio Preto, Trilha da Lagoa Seca |
| LBR0703 | LBR0703 | *Mechanitis polymnia casabranca* | *Mec. polymnia casabranca* | polymniaBraz | include | Brazil | 2018 | 15.0083 | -18.582374 | -39.72888 | male* | Conceicao da Barra, ES |

| | | | | | | | | | | | | |
|---|---|---|---|---|---|---|---|---|---|---|---|---|
| LBR0716 | LBR0716 | *Mechanitis polymnia casabranca* | *Mec. polymnia casabranca* | polymniaBraz | include | Brazil | 2018 | 16.0719 | -22.317409 | -46.32517 | male* | Incofidentes, MG |
| LBR2110 | LBR2110 | *Mechanitis polymnia casabranca* | *Mec. polymnia casabranca* | polymniaBraz | include | Brazil | 2019 | 17.5549 | -9.2423133 | -36.41815 | female* | Alagoas, Ponto 3, REBIO de Pedra Talhada |
| MCN013 | MCN013 | *Mechanitis polymnia casabranca* | *Mec. polymnia casabranca* | polymniaBraz | include | Brazil | 2018 | 17.4261 | -24.319406 | -46.99659 | female* | Peruibe, SP |
| MCN017 | MCN017 | *Mechanitis polymnia casabranca* | *Mec. polymnia casabranca* | polymniaBraz | include | Brazil | 2017 | 16.8255 | -20.284035 | -50.24386 | female* | Fernandopolis, SP |
| Sal_2038 | M2038 | *Mechanitis polymnia caucaensis* | *Mec. polymnia caucaensis* | polymniaWest | include | Colombia | 2008 | 11.4886 | 3.6604101 | -76.92442 | male | Ladrilleros, Valle del Cauca |
| Sal_2113 | M2113 | *Mechanitis polymnia caucaensis* | *Mec. polymnia caucaensis* | polymniaWest | include | Colombia | 2008 | 11.0647 | 3.6604101 | -76.92442 | male | Ladrilleros, Valle del Cauca |
| Sal_2115 | M2115 | *Mechanitis polymnia caucaensis* | *Mec. polymnia caucaensis* | polymniaWest | include | Colombia | 2008 | 12.3321 | 3.6604101 | -76.92442 | female | Ladrilleros, Valle del Cauca |
| Sal_2193 | M2193 | *Mechanitis polymnia caucaensis* | *Mec. polymnia caucaensis* | polymniaWest | include | Colombia | 2008 | 14.9999 | 3.6604101 | -76.92442 | male | Ladrilleros, Valle del Cauca |
| Sal_2195 | M2195 | *Mechanitis polymnia caucaensis* | *Mec. polymnia caucaensis* | polymniaWest | include | Colombia | 2008 | 14.4424 | 3.6604101 | -76.92442 | female | Ladrilleros, Valle del Cauca |
| Sal_4604 | M4604 | *Mechanitis polymnia caucaensis* | *Mec. polymnia caucaensis* | polymniaWest | include | Colombia | 2017 | 4.41947 | 3.6604101 | -76.92442 | female | Ladrilleros- Buenaventura, Valle del cauca |
| CAM071976 | CAM071976 | *Mechanitis polymnia chimborazona* | *Mec. polymnia chimborazona* | polymniaWest | include | Ecuador | 2023 | 30.7778 | 1.182986 | -78.75295 | male | Tundaloma Lodge, Esmeraldas |
| CAM071977 | CAM071977 | *Mechanitis polymnia chimborazona* | *Mec. polymnia chimborazona* | polymniaWest | include | Ecuador | 2023 | 31.4595 | 1.182986 | -78.75295 | male | Tundaloma Lodge, Esmeraldas |
| CAM071978 | CAM071978 | *Mechanitis polymnia chimborazona* | *Mec. polymnia chimborazona* | polymniaWest | include | Ecuador | 2023 | 32.8102 | 1.182986 | -78.75295 | male | Tundaloma Lodge, Esmeraldas |
| CAM071979 | CAM071979 | *Mechanitis polymnia chimborazona* | *Mec. polymnia chimborazona* | polymniaWest | include | Ecuador | 2023 | 32.495 | 1.182986 | -78.75295 | male | Tundaloma Lodge, Esmeraldas |
| Eli_20339 | 20339 | *Mechanitis polymnia dorissides* | *Mec. polymnia dorissides* | polymniaEast | include | Ecuador | 2005 | 13.0535 | -0.524863 | -76.38418 | male | Anangu, Orellana |
| Eli_20340 | 20340 | *Mechanitis polymnia dorissides* | *Mec. polymnia dorissides* | polymniaEast | include | Ecuador | 2005 | 15.4496 | -0.524863 | -76.38418 | male | Anangu, Orellana |
| Eli_20728 | 20728 | *Mechanitis polymnia dorissides* | *Mec. polymnia dorissides* | polymniaEast | include | Ecuador | 2005 | 11.4839 | -0.524863 | -76.38418 | male | Anangu, Orellana |
| Eli_21673 | 21673 | *Mechanitis polymnia dorissides* | *Mec. polymnia dorissides* | polymniaEast | include | Ecuador | 2007 | 15.3154 | -0.524863 | -76.38418 | male | Anangu, Orellana |
| Eli_21677 | 21677 | *Mechanitis polymnia dorissides* | *Mec. polymnia dorissides* | polymniaEast | exclude | Ecuador | 2007 | 9.50495 | -0.4617211 | -76.9936 | female | Coca, Orellana |
| Sal_1313 | M1313 | *Mechanitis polymnia dorissides* | *Mec. polymnia dorissides* | polymniaEast | include | Colombia | 2005 | 9.40955 | 1.9097464 | -75.15582 | female | Puerto Rico, Caquetá |
| Sal_1374 | M1374 | *Mechanitis polymnia dorissides* | *Mec. polymnia dorissides* | polymniaEast | include | Colombia | 2005 | 8.32645 | 1.7473478 | -75.68861 | female | Paraiso, Caquetá |
| Sal_1413 | M1413 | *Mechanitis polymnia dorissides* | *Mec. polymnia dorissides* | polymniaEast | include | Colombia | 2005 | 7.30326 | 1.5256897 | -75.47937 | male | Itarca, Caquetá |
| Sal_1416 | M1416 | *Mechanitis polymnia dorissides* | *Mec. polymnia dorissides* | polymniaEast | include | Colombia | 2005 | 7.03627 | 1.5256897 | -75.47937 | female | Itarca, Caquetá |
| Sal_4944 | M4944 | *Mechanitis polymnia dorissides* | *Mec. polymnia dorissides* | exclude | include | Colombia | 2017 | 15.7139 | 2.5824902 | -72.68893 | male | Playa Güio, Guaviare |
| CAM028186 | CAM028186 | *Mechanitis polymnia dorissides* | *Mec. polymnia dorissides* | polymniaEast | exclude | Ecuador | 2019 | 11.271 | -0.9484307 | -77.59437 | male* | Rio Pusuno, Napo |
| CAM028191 | CAM028191 | *Mechanitis polymnia dorissides* | *Mec. polymnia dorissides* | polymniaEast | exclude | Ecuador | 2019 | 10.8787 | -0.9484307 | -77.59437 | male* | Rio Pusuno, Napo |
| CAM028219 | CAM028219 | *Mechanitis polymnia dorissides* | *Mec. polymnia dorissides* | polymniaEast | exclude | Ecuador | 2019 | 5.06811 | -0.9484307 | -77.59437 | male* | Rio Pusuno, Napo |
| CAM028274 | CAM028274 | *Mechanitis polymnia dorissides* | *Mec. polymnia dorissides* | polymniaEast | exclude | Ecuador | 2019 | 11.2457 | -0.9484307 | -77.59437 | male* | Rio Pusuno, Napo |
| CAM028276 | CAM028276 | *Mechanitis polymnia dorissides* | *Mec. polymnia dorissides* | polymniaEast | include | Ecuador | 2019 | 11.5295 | -0.9484307 | -77.59437 | male* | Rio Pusuno, Napo |
| CAM028331 | CAM028331 | *Mechanitis polymnia dorissides* | *Mec. polymnia dorissides* | polymniaEast | include | Ecuador | 2019 | 13.4645 | -0.9501798 | -77.86562 | female* | Via Tena-Nuevo Angosta Km2.7, Napo |
| CAM002881 | CAM002881 | *Mechanitis polymnia isthmia* | *Mec. polymnia isthmia* | polymniaWest | include | Panama | 2000 | 7.87924 | 8.7663202 | -78.17282 | female | El Tirao, Darién |
| CAM002894 | CAM002894 | *Mechanitis polymnia isthmia* | *Mec. polymnia isthmia* | polymniaWest | include | Panama | 2000 | 9.20144 | 8.7663202 | -78.17282 | female | El Tirao, Darién |
| CAM008058 | CAM008058 | *Mechanitis polymnia isthmia* | *Mec. polymnia isthmia* | polymniaWest | include | Panama | 2000 | 12.2865 | 8.9569032 | -82.14786 | male | Chiriqui Grande - Almirante 9.5km, Bocas del Toro |
| CAM008287 | CAM008287 | *Mechanitis polymnia isthmia* | *Mec. polymnia isthmia* | polymniaWest | include | Panama | 2001 | 13.5779 | 8.713114 | -79.90659 | male | Cerro Campana |
| CAM008288 | CAM008288 | *Mechanitis polymnia isthmia* | *Mec. polymnia isthmia* | polymniaWest | include | Panama | 2001 | 9.89353 | 8.713114 | -79.90659 | male | Cerro Campana |
| Sal_5107 | M5107 | *Mechanitis polymnia isthmia* | *Mec. polymnia isthmia* | polymniaWest | include | Colombia | 2018 | 13.8266 | 5.7121727 | -75.31006 | female | Municipio de Sonson - Experiencia viva Hotel campestre, Antioquia |
| CAM009318 | CAM009318 | *Mechanitis polymnia polymnia* | *Mec. polymnia polymnia* | polymniaGui | include | Fr Guiana | NA | 9.222 | 4.5493027 | -52.49203 | male | Cacao - Cayenne, 9.9km, Cayenne |
| CAM009319 | CAM009319 | *Mechanitis polymnia polymnia* | *Mec. polymnia polymnia* | polymniaGui | include | Fr Guiana | NA | 3.99727 | 4.9399653 | -52.33921 | male | Devez Road, Cayenne |
| CAM021051 | CAM021051 | *Mechanitis polymnia polymnia* | *Mec. polymnia polymnia* | polymniaGui | include | Suriname | 2014 | 10.4614 | 5.0255233 | -54.99384 | female | Brokopondo Road East, Brokopondo |
| Eli_mm15_0350 | mm15-350 | *Mechanitis polymnia proceriformis* | *Mec. polymnia proceriformis* | polymniaPeru | include | Peru | 2015 | 12.5199 | -6.025899 | -75.87202 | female | shucushyacu |
| Eli_mm15_0528 | mm15-528 | *Mechanitis polymnia proceriformis* | *Mec. polymnia proceriformis* | polymniaPeru | include | Peru | 2015 | 12.7524 | -6.0421934 | -76.96754 | male | Moyo, Rumipata |
| Eli_mm15_0530 | mm15-530 | *Mechanitis polymnia proceriformis* | *Mec. polymnia proceriformis* | polymniaPeru | include | Peru | 2015 | 13.2699 | -6.0421934 | -76.96754 | male | Moyo, Rumipata |
| CAM008101 | CAM008101 | *Melinaea idae idae* | *Mel. idae idae* | idea | include | Panama | 2001 | 3.82078 | 9.2261912 | -80.01518731 | male* | Achiote Road, Colón |
| Sal_4172 | M4172 | *Melinaea idae idae* | *Mel. idae idae* | idea | include | Colombia | 2016 | 4.86217 | 5.6585829 | -74.18148119 | male | El Carmen1-Otanche, Boyaca |
| Sal_4174 | M4174 | *Melinaea idae idae* | *Mel. idae idae* | idea | include | Colombia | 2016 | 6.19511 | 5.6585829 | -74.18148119 | male | El Carmen1-Otanche, Boyaca |
| Sal_5620 | M5620 | *Melinaea idae idae* | *Mel. idae idae* | idea | include | Colombia | 2020 | 5.45716 | 5.6585668 | -74.18118948 | male | Potrero doña Blanca-Otanche, Boyaca |
| Sal_4286 | M4286 | *Melinaea isocomma isocomma* | *Mel. isocomma isocomma* | isocomma | include | Colombia | 2017 | 5.28586 | 1.2040722 | -76.70956725 | male | Campucana, Putumayo |
| SAN25001766 | NHMUK015104092 | *Melinaea isocomma isocomma* | *Mel. isocomma isocomma* | isocomma | include | Colombia | NA | 3.80833 | NA | NA | male | NA |
| SAN25001769 | NHMUK015200312 | *Melinaea isocomma isocomma* | *Mel. isocomma isocomma* | isocomma | include | Colombia | NA | 2.42611 | NA | NA | male | NA |
| CAM070923 | CAM070923 | *Melinaea isocomma simulator* | *Mel. isocomma simulator* | isocomma | include | Ecuador | 2023 | 13.4451 | -0.725164 | -77.767354 | male | Finca Narupa, Bridge, Napo |
| CAM070930 | CAM070930 | *Melinaea isocomma simulator* | *Mel. isocomma simulator* | isocomma | include | Ecuador | 2022 | 12.0203 | -0.725164 | -77.767354 | male | Finca Narupa, Bridge, Napo |
| CAM070960 | CAM070960 | *Melinaea isocomma simulator* | *Mel. isocomma simulator* | isocomma | include | Ecuador | 2022 | 13.6209 | -0.725164 | -77.767354 | male | Finca Narupa, Bridge, Napo |
| CAM071351 | CAM071351 | *Melinaea isocomma simulator* | *Mel. isocomma simulator* | isocomma | include | Ecuador | 2022 | 12.7849 | -0.6746389 | -77.60886111 | female | Wildsumaco |
| CAM071523 | CAM071523 | *Melinaea isocomma simulator* | *Mel. isocomma simulator* | isocomma | include | Ecuador | 2023 | 12.399 | -0.730204 | -77.773665 | female | Narupa Birding |
| CAM072002 | CAM072002 | *Melinaea isocomma simulator* | *Mel. isocomma simulator* | isocomma | include | Ecuador | 2023 | 12.1011 | -0.730204 | -77.773665 | female | Narupa Birding |
| FS50864630 | 2002-0688 | *Melinaea isocomma simulator* | *Mel. isocomma simulator* | isocomma | include | Peru | 2002 | 6.48787 | -5.6754 | -77.6746 | female* | Puente Serranoyacu, San Martin |
| FS50864631 | 2002-0689 | *Melinaea isocomma simulator* | *Mel. isocomma simulator* | isocomma | include | Peru | 2002 | 8.66875 | -5.6754 | -77.6746 | female | Puente Serranoyacu, San Martin |
| FS50864632 | 2002-0690 | *Melinaea isocomma simulator* | *Mel. isocomma simulator* | isocomma | include | Peru | 2002 | 5.43407 | -5.6754 | -77.6746 | female | Puente Serranoyacu, San Martin |
| McD_I01 | McD_I01 | *Melinaea lilis imitata* | *Mel. lilis imitata* | lilis | include | Panama | 2020 | 3.10555 | 9.1604389 | -82.51808014 | NA | Palo Seco Forest Reserve, Bocas del Toro |
| McD_I02 | McD_I02 | *Melinaea lilis imitata* | *Mel. lilis imitata* | lilis | include | Panama | 2020 | 3.22922 | 9.1604389 | -82.51808014 | NA | Palo Seco Forest Reserve, Bocas del Toro |
| McD_I03 | McD_I03 | *Melinaea lilis imitata* | *Mel. lilis imitata* | lilis | include | Panama | 2020 | 3.17569 | 9.1604389 | -82.51808014 | NA | Palo Seco Forest Reserve, Bocas del Toro |
| McD_I04 | McD_I04 | *Melinaea lilis imitata* | *Mel. lilis imitata* | lilis | include | Panama | 2020 | 3.58474 | 9.1604389 | -82.51808014 | NA | Palo Seco Forest Reserve, Bocas del Toro |
| CAM027255 | CAM027255 | *Melinaea lilis parallelis* | *Mel. lilis parallelis* | lilis | include | Panama | 2018 | 2.64285 | 8.6254434 | -80.0512129 | female | Laguna San Carlo |
| McD_P01 | McD_P01 | *Melinaea lilis parallelis* | *Mel. lilis parallelis* | lilis | include | Panama | 2020 | 3.80605 | 9.1604389 | -82.51808014 | NA | Palo Seco Forest Reserve, Bocas del Toro |
| McD_P02 | McD_P02 | *Melinaea lilis parallelis* | *Mel. lilis parallelis* | lilis | include | Panama | 2020 | 3.58577 | 9.1604389 | -82.51808014 | NA | Palo Seco Forest Reserve, Bocas del Toro |
| McD_P03 | McD_P03 | *Melinaea lilis parallelis* | *Mel. lilis parallelis* | lilis | include | Panama | 2020 | 3.53204 | 9.1604389 | -82.51808014 | NA | Palo Seco Forest Reserve, Bocas del Toro |
| McD_P04 | McD_P04 | *Melinaea lilis parallelis* | *Mel. lilis parallelis* | lilis | include | Panama | 2020 | 3.5841 | 9.1604389 | -82.51808014 | NA | Palo Seco Forest Reserve, Bocas del Toro |
| CAM009348 | CAM009348 | *Melinaea ludovica* | *Mel. ludovica* | ludovica | include | Fr Guiana | NA | 3.78025 | 3.6437425 | -53.99537461 | male | N. Wakapou, Maripausala, Saint-Laurent-du-Maroni |
| Eli_mm16_007 | mm17-007 | *Melinaea ludovica ludovica* | *Mel. ludovica ludovica* | ludovica | include | Peru | 2017 | 2.45813 | -6.025899 | -75.87202157 | male | Shucushyacu |
| Sal_5553 | M5553 | *Melinaea ludovica ludovica* | *Mel. ludovica ludovica* | ludovica | include | Colombia | NA | 4.04485 | -3.79 | -70.36 | male | Puerto Nariño- Quebrada Aguas Rojas, Amazonas |
| CAM036282 | CAM036282 | *Melinaea ludovica parayia* | *Mel. ludovica parayia* | ludovica | include | Brazil | 2020 | 7.5135 | -15.403946 | -39.54857899 | male | Serra Bonita |
| CAM036354 | CAM036354 | *Melinaea ludovica parayia* | *Mel. ludovica parayia* | ludovica | include | Brazil | 2020 | 4.07876 | -21.684803 | -48.09023939 | male | Santa Lucia |
| CAM036594 | CAM036594 | *Melinaea ludovica parayia* | *Mel. ludovica parayia* | ludovica | include | Brazil | 2020 | 5.13687 | NA | NA | male | Jureia |
| FS50864637 | 2002-3562 | *Melinaea marsaeus clara* | *Mel. marsaeus clara* | marsaeus | include | Peru | 2002 | 7.60517 | -13.2333 | -70.75 | male | Quincemil, Cuzco |
| Eli_21708 | 21708 | *Melinaea marsaeus macaria* | *Mel. marsaeus macaria* | marsaeus | include | Ecuador | 2007 | 3.85919 | NA | NA | male | Luis Garcia, Panayaku |
| Eli_21726 | 21726 | *Melinaea marsaeus macaria* | *Mel. marsaeus macaria* | marsaeus | include | Ecuador | 2007 | 3.98785 | -0.4235457 | -76.10119703 | female | Ramiro Viteri, Panacocha |
| Eli_mm16_0292 | mm16-292 | *Melinaea marsaeus phasiana* | *Mel. marsaeus phasiana* | marsaeus | include | Peru | 2016 | 4.87241 | -6.573466 | -76.1375285 | male | Chazuta |
| Eli_mm16_0327 | mm16-327 | *Melinaea marsaeus phasiana* | *Mel. marsaeus phasiana* | marsaeus | include | Peru | 2016 | 4.0949 | -6.573466 | -76.1375285 | female | Chazuta |
| Eli_20811 | 20811 | *Melinaea marsaeus rileyi* | *Mel. marsaeus rileyi* | marsaeus | include | Ecuador | 2005 | 3.4349 | -0.524863 | -76.384183 | male | Anangu, Napo |
| Eli_mm16_0227 | mm16-227 | *Melinaea marsaeus rileyi* | *Mel. marsaeus rileyi* | marsaeus | include | Peru | 2016 | 3.92723 | -6.025899 | -75.87202157 | female | Shucushyacu |
| Eli_mm16_0326 | mm16-326 | *Melinaea marsaeus rileyi* | *Mel. marsaeus rileyi* | marsaeus | include | Peru | 2016 | 3.6942 | -6.025899 | -75.87202157 | male | Shucushyacu |
| CAM077050 | M6120 | *Melinaea menophilus ernestoi* | *Mel. menophilus ernestoi* | menophilus | include | Colombia | 2022 | 14.628 | 1.147492 | -76.64800808 | male | Mocoa, Putumayo |
| CAM077076 | M6149 | *Melinaea menophilus ernestoi* | *Mel. menophilus ernestoi* | menophilus | include | Colombia | 2022 | 15.3842 | 1.147492 | -76.64800808 | male | Mocoa, Putumayo |
| Sal_1135 | M1135 | *Melinaea menophilus ernestoi* | *Mel. menophilus ernestoi* | menophilus | include | Colombia | 2005 | 3.93483 | 1.786823 | -75.65298657 | female | Sucre, Caquetá |
| Sal_2983 | M2983 | *Melinaea menophilus ernestoi* | *Mel. menophilus ernestoi* | menophilus | include | Colombia | 2010 | 4.41367 | 0.8794321 | -73.86996313 | female | Q. Doraditas 1, Caquetá |
| Sal_408 | M408 | *Melinaea menophilus ernestoi* | *Mel. menophilus ernestoi* | menophilus | include | Colombia | 2003 | 4.97289 | 1.786823 | -75.65298657 | male | Sucre, Caquetá |
| Sal_4284 | M4284 | *Melinaea menophilus ernestoi* | *Mel. menophilus ernestoi* | menophilus | include | Colombia | 2017 | 4.74927 | 1.2040722 | -76.70956725 | female | Campucana, Putumayo |
| Eli_mm16_0141 | mm16-141 | *Melinaea menophilus hicetas* | *Mel. menophilus hicetas* | menophilus | include | Peru | 2016 | 4.00594 | -6.025899 | -75.87202157 | male | Shucushyacu |
| MJ09_4179 | MJ09_4179 | *Melinaea menophilus mediatrix* | *Mel. menophilus mediatrix* | menophilus | include | Fr Guiana | 2019 | 13.8982 | 4.01106 | -52.64197 | NA | Les Nouragues-Saut Pararé |
| MJ09_4208 | MJ09_4208 | *Melinaea menophilus mediatrix* | *Mel. menophilus mediatrix* | menophilus | include | Fr Guiana | 2019 | 10.1738 | 4.01106 | -52.64197 | NA | Les Nouragues-Inselberg |
| CAM070948 | CAM070948 | *Melinaea menophilus menophilus* | *Mel. menophilus menophilus* | menophilus | include | Ecuador | 2022 | 12.1792 | -0.49673 | -76.839035 | female | Pozo Anaconda |
| CAM071272 | CAM071272 | *Melinaea menophilus menophilus* | *Mel. menophilus menophilus* | menophilus | include | Ecuador | 2022 | 13.0653 | 0.0101333 | -76.259317 | male | Pozo Cuyabeno |
| Eli_20234 | 20234 | *Melinaea menophilus menophilus* | *Mel. menophilus menophilus* | menophilus | include | Ecuador | 2005 | 3.60695 | -0.524863 | -76.384183 | male | Anangu, Napo |
| Eli_20252 | 20252 | *Melinaea menophilus menophilus* | *Mel. menophilus menophilus* | menophilus | include | Ecuador | 2005 | 3.55453 | -0.524863 | -76.384183 | male | Anangu, Napo |
| Eli_20302 | 20302 | *Melinaea menophilus menophilus* | *Mel. menophilus menophilus* | menophilus | include | Ecuador | 2005 | 3.61136 | -0.524863 | -76.384183 | male | Anangu, Napo |
| Eli_20336 | 20336 | *Melinaea menophilus menophilus* | *Mel. menophilus menophilus* | menophilus | include | Ecuador | 2005 | 4.03646 | -0.524863 | -76.384183 | female | Anangu, Napo |
| Eli_20730 | 20730 | *Melinaea menophilus menophilus* | *Mel. menophilus menophilus* | menophilus | include | Ecuador | 2005 | 3.9815 | -0.524863 | -76.384183 | female | Anangu, Napo |
| Sal_447 | M447 | *Melinaea menophilus menophilus* | *Mel. menophilus menophilus* | menophilus | include | Colombia | 2003 | 6.1394 | 1.786823 | -75.65298657 | male | Sucre, Caquetá |
| Eli_mm16_0313 | mm16-313 | *Melinaea menophilus ssp1* | *Mel. menophilus ssp1* | menophilus | include | Peru | 2016 | 4.2959 | -6.4969024 | -76.35857168 | male | rio shilcayo |
| Eli_mm16_0314 | mm16-314 | *Melinaea menophilus ssp1* | *Mel. menophilus ssp1* | menophilus | include | Peru | 2016 | 4.39352 | -6.4969024 | -76.35857168 | male | rio shilcayo |
| Eli_mm16_0317 | mm16-317 | *Melinaea menophilus ssp1* | *Mel. menophilus ssp1* | menophilus | include | Peru | 2016 | 4.62917 | -6.4969024 | -76.35857168 | female | rio shilcayo |
| Eli_mm16_0324 | mm16-324 | *Melinaea menophilus ssp1* | *Mel. menophilus ssp1* | menophilus | include | Peru | 2016 | 4.00648 | -6.0421934 | -76.96753632 | female | Moyo, Rumipata |
| CAM008526 | CAM008526 | *Melinaea menophilus zaneka* | *Mel. menophilus zaneka* | menophilus | include | Ecuador | 2002 | 2.96625 | -1.4074569 | -78.19110354 | female* | El Topo, Baños - Puyo road, Tungurahua |
| CAM008527 | CAM008527 | *Melinaea menophilus zaneka* | *Mel. menophilus zaneka* | menophilus | include | Ecuador | 2002 | 4.54474 | -1.4074569 | -78.19110354 | female* | El Topo, Baños - Puyo road, Tungurahua |
| CAM009136 | CAM009136 | *Melinaea menophilus zaneka* | *Mel. menophilus zaneka* | menophilus | include | Ecuador | 2006 | 4.23755 | -2.2627388 | -78.20150396 | NA | Rio Abanico, Macas, Morona-Santiago |
| CAM028162 | CAM028162 | *Melinaea menophilus zaneka* | *Mel. menophilus zaneka* | menophilus | include | Ecuador | 2019 | 4.8315 | -0.725164 | -77.767354 | female* | Finca Narupa, Bridge, Napo |
| CAM028163 | CAM028163 | *Melinaea menophilus zaneka* | *Mel. menophilus zaneka* | menophilus | include | Ecuador | 2019 | 3.41743 | -0.725164 | -77.767354 | female* | Finca Narupa, Bridge, Napo |
| CAM028398 | CAM028398 | *Melinaea menophilus zaneka* | *Mel. menophilus zaneka* | menophilus | include | Ecuador | 2019 | 3.83182 | -0.7200278 | -77.74708333 | female* | Mina Negra, Napo |
| CAM028402 | CAM028402 | *Melinaea menophilus zaneka* | *Mel. menophilus zaneka* | menophilus | include | Ecuador | 2019 | 4.10872 | -0.725164 | -77.767354 | NA | Finca Narupa, Bridge, Napo |
| CAM028403 | CAM028403 | *Melinaea menophilus zaneka* | *Mel. menophilus zaneka* | menophilus | include | Ecuador | 2019 | 3.91533 | -0.725164 | -77.767354 | female* | Finca Narupa, Bridge, Napo |
| CAM009349 | CAM009349 | *Melinaea mneme* | *Mel. mneme* | mneme | include | Fr Guiana | NA | 4.53344 | 3.6437425 | -53.99537461 | male | N. Wakapou, Maripausala, Saint-Laurent-du-Maroni |
| MJ09_4176 | MJ09_4176 | *Melinaea mneme* | *Mel. mneme* | mneme | include | Fr Guiana | 2019 | 9.8136 | 4.01106 | -52.64197 | NA | Les Nouragues-Saut Pararé |
| MJ09_4199 | MJ09_4199 | *Melinaea mneme* | *Mel. mneme* | mneme | include | Fr Guiana | 2019 | 11.5887 | 4.01106 | -52.64197 | NA | Les Nouragues-Inselberg |
| Sal_5371 | M5371 | *Melinaea mneme* | *Mel. mneme* | mneme | include | Colombia | 2018 | 5.40237 | -1.3220609 | -69.57810912 | male | Vereda Madroño - La Pedrera, Amazonas |
| Sal_5393 | M5393 | *Melinaea mneme* | *Mel. mneme* | mneme | include | Colombia | 2018 | 5.79792 | -1.3220609 | -69.57810912 | female | Sendero Bacurí - La Pedrera, Amazonas |
| CAM077033 | M6090 | *Melinaea mothone messenina* | *Mel. mothone messenina* | mothone | include | Colombia | 2023 | 12.6359 | 1.147492 | -76.64800808 | female | Mocoa, Putumayo |
| CAM077035 | M6092 | *Melinaea mothone messenina* | *Mel. mothone messenina* | mothone | include | Colombia | 2022 | 16.3272 | 1.147492 | -76.64800808 | female | Mocoa, Putumayo |
| Sal_1069 | M1069 | *Melinaea mothone messenina* | *Mel. mothone messenina* | mothone | include | Colombia | 2005 | 5.68367 | 1.786823 | -75.65298657 | male | Sucre, Caquetá |
| Sal_1136 | M1136 | *Melinaea mothone messenina* | *Mel. mothone messenina* | mothone | include | Colombia | 2005 | 5.18706 | 1.786823 | -75.65298657 | female | Sucre, Caquetá |
| CAM077000 | M6039 | *Melinaea mothone mot/mess* | *Mel. mothone mot/mess* | mothone | include | Colombia | 2022 | 15.2665 | 1.147492 | -76.64800808 | female | Mocoa, Putumayo |
| CAM070362 | CAM070362 | *Melinaea mothone mothone* | *Mel. mothone mothone* | mothone | include | Ecuador | 2023 | 15.8951 | -0.725164 | -77.767354 | male* | Finca Narupa, Bridge, Napo |
| CAM070921 | CAM070921 | *Melinaea mothone mothone* | *Mel. mothone mothone* | mothone | include | Ecuador | 2023 | 14.5282 | -0.725164 | -77.767354 | male* | Finca Narupa, Bridge, Napo |

| | | | | | | | | | | | | |
|---|---|---|---|---|---|---|---|---|---|---|---|---|
| CAM070922 | CAM070922 | *Melinaea mothone mothone* | *Mel. mothone mothone* | mothone | include | Ecuador | 2023 | 14.6653 | -0.725164 | -77.767354 | male* | Finca Narupa, Bridge, Napo |
| CAM070924 | CAM070924 | *Melinaea mothone mothone* | *Mel. mothone mothone* | mothone | include | Ecuador | 2023 | 16.0467 | -0.725164 | -77.767354 | male* | Finca Narupa, Bridge, Napo |
| CAM070925 | CAM070925 | *Melinaea mothone mothone* | *Mel. mothone mothone* | mothone | include | Ecuador | 2023 | 15.0785 | -0.725164 | -77.767354 | male* | Finca Narupa, Bridge, Napo |
| CAM070926 | CAM070926 | *Melinaea mothone mothone* | *Mel. mothone mothone* | mothone | include | Ecuador | 2023 | 16.5245 | -0.725164 | -77.767354 | male* | Finca Narupa, Bridge, Napo |
| CAM077032 | M6089 | *Melinaea mothone mothone* | *Mel. mothone mothone* | mothone | include | Colombia | 2023 | 13.6359 | 1.147492 | -76.64800808 | female | Mocoa, Putumayo |
| FS50864628 | 2002-0378 | *Melinaea mothone mothone* | *Mel. mothone mothone* | mothone | include | Peru | 2023 | 8.77395 | -6.455 | -76.2983 | female | La Antena, Km-16, Tarapoto-Yurimaguas; San Martin |
| FS50864629 | 2002-0675 | *Melinaea mothone mothone* | *Mel. mothone mothone* | mothone | include | Peru | 2023 | 8.20904 | -6.46183 | -76.2919 | male | Biodiversidad USM, Km-19, Tarapoto-Yurimaguas; San Martin |
| FS50864633 | 2002-0982 | *Melinaea mothone mothone* | *Mel. mothone mothone* | mothone | include | Peru | 2023 | 8.06032 | -6.4619 | -76.3231 | male | Km-10, Tarapoto-Yurimaguas, San Martin |
| FS50864635 | 2002-3558 | *Melinaea mothone mothone* | *Mel. mothone mothone* | mothone | include | Peru | 2023 | 9.31932 | -13.2167 | -70.75 | male | Quincemil, Cuzco |
| FS50864636 | 2002-3560 | *Melinaea mothone mothone* | *Mel. mothone mothone* | mothone | include | Peru | 2023 | 7.51254 | -13.2167 | -70.75 | male | Quincemil, Cuzco |
| FS50864642 | 2005-1342 | *Melinaea mothone mothone* | *Mel. mothone mothone* | mothone | include | Peru | 2023 | 9.24518 | -6.4519 | -76.3461 | male* | Río Shilcayo, Tarapoto, San Martin |
| Eli_mm16_0205 | mm16-205 | *Melinaea satevis cydon* | *Mel. satevis cydon* | satevis | include | Peru | 2016 | 3.53011 | -6.4805835 | -76.34799034 | male | PB Km 7.5 |
| Eli_mm16_0206 | mm16-206 | *Melinaea satevis cydon* | *Mel. satevis cydon* | satevis | include | Peru | 2016 | 4.11701 | -6.4805835 | -76.34799034 | female | PB Km 7.5 |
| Sal_5517 | M5517 | *Melinaea satevis cydon* | *Mel. satevis cydon* | satevis | include | Colombia | NA | 5.11485 | -3.7886904 | -70.35546689 | male | Puerto Nariño- Quebrada Aguas Rojas, Amazonas |
| Sal_5570 | M5570 | *Melinaea satevis cydon* | *Mel. satevis cydon* | satevis | include | Colombia | NA | 4.7983 | -3.79 | -70.36 | male | Puerto Nariño- Quebrada Aguas Rojas, Amazonas |
| SAN25001774 | LEP-56663 | *Melinaea satevis cydon* | *Mel. satevis cydon* | satevis | include | Ecuador | NA | 14.1981 | -2.902 | -77.742 | female* | 2.5 km N Puerto Morona |
| SAN25001775 | LEP-57574 | *Melinaea satevis cydon* | *Mel. satevis cydon* | satevis | include | Ecuador | NA | 15.5379 | -2.912 | -77.707 | female* | km 3 Puerto Morona-San José de Morona |
| FS50864634 | 2002-3262 | *Melinaea satevis lamasi* | *Mel. satevis lamasi* | satevis | include | Peru | 2002 | 8.72197 | -12.9 | -71.4 | female | Pilcopata, Cuzco |
| Sal_5372 | M5372 | *Melinaea satevis maelus* | *Mel. satevis maelus* | satevis | include | Colombia | 2018 | 4.21181 | -1.3220609 | -69.57810912 | female | Vereda Madroño - La Pedrera, Amazonas |
| CAM028332 | CAM028332 | *Melinaea satevis maeonis* | *Mel. maeonis* | maeonis | include | Ecuador | 2019 | 5.0276 | -0.9501798 | -77.8656204 | female* | Via Tena-Nuevo Angosta Km2.7, Napo |
| CAM071086 | CAM071086 | *Melinaea satevis maeonis* | *Mel. maeonis* | maeonis | include | Ecuador | 2022 | 8.67416 | 0.1995667 | -77.471 | male | Puerto Libre Linea 2 |
| Eli_20301 | 20301 | *Melinaea satevis maeonis* | *Mel. maeonis* | maeonis | include | Ecuador | 2005 | 4.12592 | -0.524863 | -76.384183 | male | Anangu, Napo |
| Eli_20549 | 20549 | *Melinaea satevis maeonis* | *Mel. maeonis* | maeonis | include | Ecuador | 2005 | 3.90169 | -0.524863 | -76.384183 | female | Anangu, Napo |
| Eli_20594 | 20594 | *Melinaea satevis maeonis* | *Mel. maeonis* | maeonis | include | Ecuador | 2005 | 4.45302 | -0.524863 | -76.384183 | female* | Anangu, Napo |
| FS50864626 | E450 | *Melinaea satevis maeonis* | *Mel. maeonis* | maeonis | include | Ecuador | 2002 | 9.60045 | NA | NA | female | Jatun Sacha, Napo |
| Eli_mm16_0161 | mm16-161 | *Melinaea tarapotensis* | *Mel. tarapotensis* | tarapotensis | include | Peru | 2016 | 4.12229 | -6.4837118 | -76.3706028 | male | Tarapoto |
| Eli_mm16_0173 | mm16-173 | *Melinaea tarapotensis* | *Mel. tarapotensis* | tarapotensis | include | Peru | 2016 | 3.81347 | -6.4837118 | -76.3706028 | female | Tarapoto |
| Eli_mm16_0197 | mm16-197 | *Melinaea tarapotensis* | *Mel. tarapotensis* | tarapotensis | include | Peru | 2016 | 4.38264 | -6.4837118 | -76.3706028 | male | Tarapoto |
| Eli_mm16_0296 | mm16-296 | *Melinaea tarapotensis* | *Mel. tarapotensis* | tarapotensis | include | Peru | 2016 | 4.34819 | -6.4969024 | -76.35857168 | female | rio shilcayo |
| FS50864856 | CAM071929 | *Olyras insignis* | *Oly. insignis* | Outgroup (Mel.) | include | Ecuador | 2023 | 2.64989 | 0.885174 | -78.436529 | female | Lita Ridge |

**Supplementary Table 2: MSCi results.**

Raw (A) and scaled (B) results of the MSCi runs for each set of species. Scaling factors are shown at the top. For each replicate and parameter a median and confidence interval are shown.

| | | | |
|---|---|---|---|
| **u** | 2.90E-09 mutation rate subs/site/generation | **Ne** | Effective population size (x10^6) |
| **g** | 4 generations per year | **Tdiv** | Time in years (x10^6) |

A

Raw

| Set | Run | | theta_1A | theta_2B | theta_3C | theta_4R | theta_5S | theta_6T | theta_7H | theta_8H | tau_4R | tau_5S | tau_6T | tau_7H | phi_H<-T | lnL |
|---|---|---|---|---|---|---|---|---|---|---|---|---|---|---|---|---|
| lil,ida,mar | rep1 | median | 0.005709 | 0.023926 | 0.006146 | 0.007284 | 0.00491 | 0.002574 | 0.014967 | 0.0048085 | 0.004639 | 0.003083 | 0.002973 | 0.002958 | 0.226106 | -4104416 |
| | | 2.5%HPD | 0.00538 | 0.020673 | 0.00565 | 0.006949 | 0.004449 | 0.002282 | 0.003142 | 0.001193 | 0.004512 | 0.00298 | 0.002752 | 0.002729 | 0.180464 | -4104588 |
| | | 97.5%HPD | 0.006042 | 0.027756 | 0.006813 | 0.007588 | 0.005364 | 0.002898 | 0.056334 | 0.03309 | 0.004755 | 0.003191 | 0.003104 | 0.00309 | 0.267851 | -4104243 |
| | rep2 | median | 0.007895 | 0.01357 | 0.006961 | 0.007334 | 0.002109 | 0.004215 | 0.001902 | 0.015137 | 0.004556 | 0.00244 | 0.003629 | 0.002429 | 0.738719 | -4104328 |
| | | 2.5%HPD | 0.007261 | 0.01251 | 0.006462 | 0.007024 | 0.001913 | 0.003598 | 0.001543 | 0.003384 | 0.004449 | 0.002331 | 0.003512 | 0.00232 | 0.712043 | -4104498 |
| | | 97.5%HPD | 0.008573 | 0.014657 | 0.00746 | 0.00765 | 0.002302 | 0.004853 | 0.002266 | 0.054689 | 0.004657 | 0.002541 | 0.003743 | 0.002532 | 0.764557 | -4104155 |
| | rep3 | median | 0.007894 | 0.013566 | 0.006964 | 0.007334 | 0.002113 | 0.004216 | 0.001902 | 0.0147135 | 0.004556 | 0.002439 | 0.00363 | 0.002428 | 0.738495 | -4104336 |
| | | 2.5%HPD | 0.007248 | 0.012524 | 0.006479 | 0.007023 | 0.001915 | 0.003569 | 0.001546 | 0.003658 | 0.004449 | 0.002334 | 0.003508 | 0.002318 | 0.711632 | -4104507 |
| | | 97.5%HPD | 0.008558 | 0.014676 | 0.007477 | 0.007645 | 0.002306 | 0.004846 | 0.002266 | 0.047249 | 0.004657 | 0.002547 | 0.003743 | 0.002533 | 0.76465 | -4104165 |
| | rep4 | median | 0.005794 | 0.023113 | 0.005998 | 0.007191 | 0.005005 | 0.002651 | 0.03758 | 0.035415 | 0.004717 | 0.003025 | 0.003016 | 0.003004 | 0.247922 | -4104416 |
| | | 2.5%HPD | 0.005459 | 0.02046 | 0.005671 | 0.006881 | 0.004582 | 0.002399 | 0.008969 | 0.008048 | 0.004622 | 0.002952 | 0.002938 | 0.002929 | 0.218803 | -4104588 |
| | | 97.5%HPD | 0.006127 | 0.02597 | 0.006343 | 0.007493 | 0.00546 | 0.002907 | 0.121584 | 0.118853 | 0.004813 | 0.003097 | 0.003088 | 0.003077 | 0.276993 | -4104246 |
| | rep5 | median | 0.0058 | 0.023127 | 0.005994 | 0.007193 | 0.005017 | 0.002655 | 0.0373215 | 0.035993 | 0.004717 | 0.003023 | 0.003014 | 0.003002 | 0.247677 | -4104423 |
| | | 2.5%HPD | 0.005464 | 0.020469 | 0.005648 | 0.006891 | 0.004588 | 0.002405 | 0.008984 | 0.007798 | 0.004619 | 0.002947 | 0.002939 | 0.002925 | 0.219389 | -4104593 |
| | | 97.5%HPD | 0.00613 | 0.025987 | 0.00633 | 0.007495 | 0.00546 | 0.002913 | 0.118262 | 0.126347 | 0.004808 | 0.003094 | 0.00309 | 0.003074 | 0.27737 | -4104250 |
| | rep6 | median | 0.005793 | 0.023151 | 0.005998 | 0.007194 | 0.00502 | 0.00265 | 0.037641 | 0.0362615 | 0.004716 | 0.003023 | 0.003014 | 0.003002 | 0.247292 | -4104430 |
| | | 2.5%HPD | 0.005459 | 0.02043 | 0.005658 | 0.006895 | 0.004571 | 0.002393 | 0.00856 | 0.00829 | 0.004616 | 0.002947 | 0.002934 | 0.002924 | 0.218636 | -4104600 |
| | | 97.5%HPD | 0.006131 | 0.025998 | 0.006331 | 0.007499 | 0.005473 | 0.002911 | 0.128795 | 0.125983 | 0.004808 | 0.003097 | 0.00309 | 0.003077 | 0.276438 | -4104255 |

| Set | Run | | theta_1A | theta_2B | theta_3C | theta_4R | theta_5S | theta_6T | theta_7H | theta_8H | tau_4R | tau_5S | tau_6T | tau_7H | phi_H<-T | lnL |
|---|---|---|---|---|---|---|---|---|---|---|---|---|---|---|---|---|
| men,mes,pol | rep1 | median | 0.002439 | 0.020932 | 0.022299 | 0.008954 | 0.018812 | 0.007279 | 0.0142365 | 0.001115 | 0.008334 | 0.005175 | 0.0049 | 0.004878 | 0.092437 | -19212611 |
| | | 2.5%HPD | 0.00238 | 0.020105 | 0.02128 | 0.008736 | 0.017825 | 0.006922 | 0.003166 | 0.000892 | 0.008262 | 0.005101 | 0.004815 | 0.004797 | 0.083397 | -19212942 |
| | | 97.5%HPD | 0.002498 | 0.021781 | 0.023362 | 0.009164 | 0.019851 | 0.00764 | 0.044824 | 0.001359 | 0.008405 | 0.005245 | 0.004983 | 0.004957 | 0.10177 | -19212288 |
| | rep2 | median | 0.002426 | 0.01634 | 0.022952 | 0.009017 | 0.007364 | 0.011995 | 0.002296 | 0.013993 | 0.008346 | 0.004629 | 0.006775 | 0.004624 | 0.7325595 | -19212861 |
| | | 2.5%HPD | 0.002364 | 0.015831 | 0.021902 | 0.008784 | 0.006695 | 0.011003 | 0.001965 | 0.003448 | 0.008265 | 0.004516 | 0.006666 | 0.004514 | 0.712119 | -19213192 |
| | | 97.5%HPD | 0.002485 | 0.016849 | 0.024044 | 0.009231 | 0.008036 | 0.013005 | 0.002633 | 0.041549 | 0.008421 | 0.004729 | 0.006879 | 0.004727 | 0.752037 | -19212533 |
| | rep3 | median | 0.002426 | 0.016341 | 0.022938 | 0.00902 | 0.007367 | 0.011996 | 0.002285 | 0.014879 | 0.008345 | 0.004634 | 0.006773 | 0.004629 | 0.732153 | -19212854 |
| | | 2.5%HPD | 0.002363 | 0.015824 | 0.02186 | 0.008793 | 0.006714 | 0.010977 | 0.001937 | 0.00325 | 0.008266 | 0.004512 | 0.006661 | 0.004504 | 0.71147 | -19213180 |
| | | 97.5%HPD | 0.002483 | 0.016838 | 0.02402 | 0.009236 | 0.008045 | 0.012987 | 0.002632 | 0.048943 | 0.008421 | 0.004742 | 0.006875 | 0.004735 | 0.752086 | -19212524 |
| | rep4 | median | 0.00243 | 0.016313 | 0.023064 | 0.009038 | 0.007238 | 0.012048 | 0.002547 | 0.0356325 | 0.008335 | 0.004572 | 0.006801 | 0.004566 | 0.7427165 | -19212828 |
| | | 2.5%HPD | 0.002367 | 0.015817 | 0.021985 | 0.008812 | 0.006589 | 0.011024 | 0.002231 | 0.009034 | 0.008256 | 0.004467 | 0.006694 | 0.004463 | 0.724138 | -19213156 |
| | | 97.5%HPD | 0.002486 | 0.016831 | 0.024161 | 0.009258 | 0.007911 | 0.013052 | 0.002866 | 0.106088 | 0.008412 | 0.004677 | 0.006901 | 0.004674 | 0.761274 | -19212502 |
| | rep5 | median | 0.00243 | 0.016312 | 0.02307 | 0.009036 | 0.00724 | 0.012046 | 0.002556 | 0.0407865 | 0.008335 | 0.004568 | 0.006802 | 0.004562 | 0.742936 | -19212823 |
| | | 2.5%HPD | 0.002369 | 0.015803 | 0.022021 | 0.008809 | 0.006589 | 0.011013 | 0.002219 | 0.009558 | 0.008257 | 0.004462 | 0.006695 | 0.004457 | 0.723981 | -19213156 |
| | | 97.5%HPD | 0.002489 | 0.01681 | 0.024178 | 0.00926 | 0.007906 | 0.013063 | 0.002881 | 0.147218 | 0.008414 | 0.004676 | 0.006902 | 0.004671 | 0.761847 | -19212497 |
| | rep6 | median | 0.00243 | 0.016315 | 0.023064 | 0.009035 | 0.007249 | 0.012054 | 0.002546 | 0.039615 | 0.008336 | 0.004571 | 0.006799 | 0.004565 | 0.742421 | -19212816 |
| | | 2.5%HPD | 0.002373 | 0.015814 | 0.021982 | 0.008807 | 0.00661 | 0.011063 | 0.002229 | 0.00894 | 0.008255 | 0.004458 | 0.006694 | 0.00446 | 0.723496 | -19213153 |
| | | 97.5%HPD | 0.002492 | 0.016828 | 0.024154 | 0.009255 | 0.007928 | 0.013083 | 0.002878 | 0.114832 | 0.008412 | 0.004673 | 0.0069 | 0.004676 | 0.761143 | -19212494 |

| Set | Run | | theta_1A | theta_2B | theta_3C | theta_4R | theta_5S | theta_6T | theta_7H | theta_8H | tau_4R | tau_5S | tau_6T | tau_7H | phi_H<-T | lnL |
|---|---|---|---|---|---|---|---|---|---|---|---|---|---|---|---|---|
| pol,nes,lys | rep1 | median | 0.040143 | 0.017117 | 0.007383 | 0.006955 | 0.003729 | 0.00978 | 0.001525 | 0.014472 | 0.006226 | 0.004193 | 0.005037 | 0.004185 | 0.92675 | -19413288.66 |
| | | 2.5%HPD | 0.037547 | 0.016539 | 0.007168 | 0.0068 | 0.003518 | 0.009032 | 0.001319 | 0.003476 | 0.006178 | 0.004113 | 0.004962 | 0.004107 | 0.917633 | -19413635.29 |
| | | 97.5%HPD | 0.042721 | 0.017712 | 0.007598 | 0.007107 | 0.003931 | 0.010544 | 0.001727 | 0.046269 | 0.006277 | 0.004267 | 0.005108 | 0.004262 | 0.935551 | -19412935.88 |
| | rep2 | median | 0.040165 | 0.017115 | 0.007383 | 0.006955 | 0.003731 | 0.009783 | 0.001528 | 0.014533 | 0.006226 | 0.004191 | 0.005038 | 0.004183 | 0.926784 | -19413297.32 |
| | | 2.5%HPD | 0.037595 | 0.016546 | 0.007167 | 0.006798 | 0.003514 | 0.009022 | 0.001337 | 0.003526 | 0.006178 | 0.004115 | 0.004963 | 0.004107 | 0.91768 | -19413648.24 |
| | | 97.5%HPD | 0.042764 | 0.017718 | 0.007597 | 0.007108 | 0.003934 | 0.010528 | 0.001731 | 0.041393 | 0.006277 | 0.004271 | 0.005108 | 0.004265 | 0.935586 | -19412948.16 |
| | rep3 | median | 0.040185 | 0.017115 | 0.007382 | 0.006953 | 0.003735 | 0.009774 | 0.001531 | 0.014828 | 0.006227 | 0.004191 | 0.005038 | 0.004183 | 0.926875 | -19413294.37 |
| | | 2.5%HPD | 0.037622 | 0.016518 | 0.007171 | 0.006799 | 0.00353 | 0.009046 | 0.001329 | 0.003759 | 0.006178 | 0.004109 | 0.004966 | 0.004098 | 0.917832 | -19413641.94 |
| | | 97.5%HPD | 0.042817 | 0.01769 | 0.007598 | 0.007111 | 0.003944 | 0.010525 | 0.001741 | 0.045361 | 0.006277 | 0.004268 | 0.00511 | 0.004259 | 0.935698 | -19412938.77 |

B

Scaled

| Set | Run | Stat | Ne_1A | Ne_2B | Ne_3C | Ne_4R | Ne_5S | Ne_6T | Ne_7H | Ne_8H | Tdiv_4R | Tdiv_5S | Tdiv_6T | Tdiv_7H | phi_H<-T | lnL |
|---|---|---|---|---|---|---|---|---|---|---|---|---|---|---|---|---|
| lil,ida,mar | rep1 | median | 0.49 | 2.06 | 0.53 | 0.63 | 0.42 | 0.22 | 1.29 | 0.41 | 0.400 | 0.266 | 0.256 | 0.255 | 22.6% | |
| | | 2.5%HPD | 0.46 | 1.78 | 0.49 | 0.60 | 0.38 | 0.20 | 0.27 | 0.10 | 0.389 | 0.257 | 0.237 | 0.235 | 18.0% | -4,104,243 |
| | | 97.5%HPD | 0.52 | 2.39 | 0.59 | 0.65 | 0.46 | 0.25 | 4.86 | 2.85 | 0.410 | 0.275 | 0.268 | 0.266 | 26.8% | |
| | rep2 | median | 0.68 | 1.17 | 0.60 | 0.63 | 0.18 | 0.36 | 0.16 | 1.30 | 0.393 | 0.210 | 0.313 | 0.209 | 73.9% | |
| | | 2.5%HPD | 0.63 | 1.08 | 0.56 | 0.61 | 0.16 | 0.31 | 0.13 | 0.29 | 0.384 | 0.201 | 0.303 | 0.200 | 71.2% | -4,104,155 |
| | | 97.5%HPD | 0.74 | 1.26 | 0.64 | 0.66 | 0.20 | 0.42 | 0.20 | 4.71 | 0.401 | 0.219 | 0.323 | 0.218 | 76.5% | |
| | rep3 | median | 0.68 | 1.17 | 0.60 | 0.63 | 0.18 | 0.36 | 0.16 | 1.27 | 0.39 | 0.21 | 0.31 | 0.21 | 73.8% | |
| | | 2.5%HPD | 0.62 | 1.08 | 0.56 | 0.61 | 0.17 | 0.31 | 0.13 | 0.32 | 0.38 | 0.20 | 0.30 | 0.20 | 71.2% | |
| | | 97.5%HPD | 0.74 | 1.27 | 0.64 | 0.66 | 0.20 | 0.42 | 0.20 | 4.07 | 0.40 | 0.22 | 0.32 | 0.22 | 76.5% | -4,104,165 |
| | rep4 | median | 0.50 | 1.99 | 0.52 | 0.62 | 0.43 | 0.23 | 3.24 | 3.05 | 0.41 | 0.26 | 0.26 | 0.26 | 24.8% | |
| | | 2.5%HPD | 0.47 | 1.76 | 0.49 | 0.59 | 0.40 | 0.21 | 0.77 | 0.69 | 0.40 | 0.25 | 0.25 | 0.25 | 21.9% | |
| | | 97.5%HPD | 0.53 | 2.24 | 0.55 | 0.65 | 0.47 | 0.25 | 10.48 | 10.25 | 0.41 | 0.27 | 0.27 | 0.27 | 27.7% | -4,104,246 |
| | rep5 | median | 0.50 | 1.99 | 0.52 | 0.62 | 0.43 | 0.23 | 3.22 | 3.10 | 0.41 | 0.26 | 0.26 | 0.26 | 24.8% | |
| | | 2.5%HPD | 0.47 | 1.76 | 0.49 | 0.59 | 0.40 | 0.21 | 0.77 | 0.67 | 0.40 | 0.25 | 0.25 | 0.25 | 21.9% | |
| | | 97.5%HPD | 0.53 | 2.24 | 0.55 | 0.65 | 0.47 | 0.25 | 10.20 | 10.89 | 0.41 | 0.27 | 0.27 | 0.27 | 27.7% | -4,104,250 |
| | rep6 | median | 0.50 | 2.00 | 0.52 | 0.62 | 0.43 | 0.23 | 3.24 | 3.13 | 0.41 | 0.26 | 0.26 | 0.26 | 24.7% | |
| | | 2.5%HPD | 0.47 | 1.76 | 0.49 | 0.59 | 0.39 | 0.21 | 0.74 | 0.71 | 0.40 | 0.25 | 0.25 | 0.25 | 21.9% | |
| | | 97.5%HPD | 0.53 | 2.24 | 0.55 | 0.65 | 0.47 | 0.25 | 11.10 | 10.86 | 0.41 | 0.27 | 0.27 | 0.27 | 27.6% | -4,104,255 |

| Set | Run | Stat | Ne_1A | Ne_2B | Ne_3C | Ne_4R | Ne_5S | Ne_6T | Ne_7H | Ne_8H | Tdiv_4R | Tdiv_5S | Tdiv_6T | Tdiv_7H | phi_H<-T | lnL |
|---|---|---|---|---|---|---|---|---|---|---|---|---|---|---|---|---|
| men,mes,pol | rep1 | median | 0.21 | 1.80 | 1.92 | 0.77 | 1.62 | 0.63 | 1.23 | 0.10 | 0.718 | 0.446 | 0.422 | 0.421 | 9.2% | |
| | | 2.5%HPD | 0.21 | 1.73 | 1.83 | 0.75 | 1.54 | 0.60 | 0.27 | 0.08 | 0.712 | 0.440 | 0.415 | 0.414 | 8.3% | -19,212,611 |
| | | 97.5%HPD | 0.22 | 1.88 | 2.01 | 0.79 | 1.71 | 0.66 | 3.86 | 0.12 | 0.725 | 0.452 | 0.430 | 0.427 | 10.2% | |
| | rep2 | median | 0.21 | 1.41 | 1.98 | 0.78 | 0.63 | 1.03 | 0.20 | 1.21 | 0.719 | 0.399 | 0.584 | 0.399 | 73.3% | |
| | | 2.5%HPD | 0.20 | 1.36 | 1.89 | 0.76 | 0.58 | 0.95 | 0.17 | 0.30 | 0.713 | 0.389 | 0.575 | 0.389 | 71.2% | -19,212,861 |
| | | 97.5%HPD | 0.21 | 1.45 | 2.07 | 0.80 | 0.69 | 1.12 | 0.23 | 3.58 | 0.726 | 0.408 | 0.593 | 0.408 | 75.2% | |
| | rep3 | median | 0.21 | 1.41 | 1.98 | 0.78 | 0.64 | 1.03 | 0.20 | 1.28 | 0.719 | 0.399 | 0.584 | 0.399 | 73.2% | |
| | | 2.5%HPD | 0.20 | 1.36 | 1.88 | 0.76 | 0.58 | 0.95 | 0.17 | 0.28 | 0.713 | 0.389 | 0.574 | 0.388 | 71.1% | -19,212,854 |
| | | 97.5%HPD | 0.21 | 1.45 | 2.07 | 0.80 | 0.69 | 1.12 | 0.23 | 4.22 | 0.726 | 0.409 | 0.593 | 0.408 | 75.2% | |
| | rep4 | median | 0.21 | 1.41 | 1.99 | 0.78 | 0.62 | 1.04 | 0.22 | 3.07 | 0.719 | 0.394 | 0.586 | 0.394 | 74.3% | |
| | | 2.5%HPD | 0.20 | 1.36 | 1.90 | 0.76 | 0.57 | 0.95 | 0.19 | 0.78 | 0.712 | 0.385 | 0.577 | 0.385 | 72.4% | -19,212,828 |
| | | 97.5%HPD | 0.21 | 1.45 | 2.08 | 0.80 | 0.68 | 1.13 | 0.25 | 9.15 | 0.725 | 0.403 | 0.595 | 0.403 | 76.1% | |
| | rep5 | median | 0.21 | 1.41 | 1.99 | 0.78 | 0.62 | 1.04 | 0.22 | 3.52 | 0.719 | 0.394 | 0.586 | 0.393 | 74.3% | |
| | | 2.5%HPD | 0.20 | 1.36 | 1.90 | 0.76 | 0.57 | 0.95 | 0.19 | 0.82 | 0.712 | 0.385 | 0.577 | 0.384 | 72.4% | -19,212,823 |
| | | 97.5%HPD | 0.21 | 1.45 | 2.08 | 0.80 | 0.68 | 1.13 | 0.25 | 12.69 | 0.725 | 0.403 | 0.595 | 0.403 | 76.2% | |
| | rep6 | median | 0.21 | 1.41 | 1.99 | 0.78 | 0.62 | 1.04 | 0.22 | 3.42 | 0.719 | 0.394 | 0.586 | 0.394 | 74.2% | |
| | | 2.5%HPD | 0.20 | 1.36 | 1.90 | 0.76 | 0.57 | 0.95 | 0.19 | 0.77 | 0.712 | 0.384 | 0.577 | 0.384 | 72.3% | -19,212,816 |
| | | 97.5%HPD | 0.21 | 1.45 | 2.08 | 0.80 | 0.68 | 1.13 | 0.25 | 9.90 | 0.725 | 0.403 | 0.595 | 0.403 | 76.1% | |

| Set | Run | Stat | Ne_1A | Ne_2B | Ne_3C | Ne_4R | Ne_5S | Ne_6T | Ne_7H | Ne_8H | Tdiv_4R | Tdiv_5S | Tdiv_6T | Tdiv_7H | phi_H<-T | lnL |
|---|---|---|---|---|---|---|---|---|---|---|---|---|---|---|---|---|
| pol,nes,lys | rep1 | median | 3.46 | 1.48 | 0.64 | 0.60 | 0.32 | 0.84 | 0.13 | 1.25 | 0.537 | 0.361 | 0.434 | 0.361 | 92.7% | |
| | | 2.5%HPD | 3.24 | 1.43 | 0.62 | 0.59 | 0.30 | 0.78 | 0.11 | 0.30 | 0.533 | 0.355 | 0.428 | 0.354 | 91.8% | -19,413,635 |
| | | 97.5%HPD | 3.68 | 1.53 | 0.66 | 0.61 | 0.34 | 0.91 | 0.15 | 3.99 | 0.541 | 0.368 | 0.440 | 0.367 | 0.936 | |
| | rep2 | median | 3.46 | 1.48 | 0.64 | 0.60 | 0.32 | 0.84 | 0.13 | 1.25 | 0.537 | 0.361 | 0.434 | 0.361 | 92.7% | |
| | | 2.5%HPD | 3.24 | 1.43 | 0.62 | 0.59 | 0.30 | 0.78 | 0.12 | 0.30 | 0.533 | 0.355 | 0.428 | 0.354 | 91.8% | -19413648.24 |
| | | 97.5%HPD | 3.69 | 1.53 | 0.65 | 0.61 | 0.34 | 0.91 | 0.15 | 3.57 | 0.541 | 0.368 | 0.440 | 0.368 | 0.936 | |
| | rep3 | median | 3.46 | 1.48 | 0.64 | 0.60 | 0.32 | 0.84 | 0.13 | 1.28 | 0.537 | 0.361 | 0.434 | 0.361 | 92.7% | |
| | | 2.5%HPD | 3.24 | 1.42 | 0.62 | 0.59 | 0.30 | 0.78 | 0.11 | 0.32 | 0.533 | 0.354 | 0.428 | 0.353 | 91.8% | -19413641.94 |
| | | 97.5%HPD | 3.69 | 1.53 | 0.66 | 0.61 | 0.34 | 0.91 | 0.15 | 3.91 | 0.541 | 0.368 | 0.441 | 0.367 | 0.936 | |

**Supplementary Table 3: Details of the androconial chemicals**

| ID | Endemism Centre | Municipality / State | Sampling Area | Sampling altitude (m) | 1150 | 1252 | 1264 | 1318 | 1365 | 1481 | 1543 | 1551 | 1589 | 1599 | 1619 | 1673 | 1749 | 1792 | 1804 | 1843 | 1852 | 1950 | 2145 | 2162 | 2172 | 2220 | 2572 | 2675 | 2832 | 2873 |
|---|---|---|---|---|---|---|---|---|---|---|---|---|---|---|---|---|---|---|---|---|---|---|---|---|---|---|---|---|---|---|
| M.polymnia662 | Serra do Mar | Tremembé / SP | Urban Park | 500-550 | 0 | 0 | 0 | 1.81439521 | 0 | 83.6593395 | 0 | 0 | 10.4149599 | 0 | 0 | 0.17222797 | 0 | 0 | 0 | 0 | 0 | 0.79566418 | 0 | 0 | 0 | 1.15038913 | 0 | 0 | 1.99302409 | 0 |
| M.polymnia663 | Serra do Mar | Tremembé / SP | Urban Park | 500-550 | 0 | 0 | 0 | 1.94328697 | 0 | 63.7618607 | 0 | 0 | 13.3624124 | 0 | 0 | 0 | 0 | 0 | 0 | 0 | 0 | 1.24065814 | 0 | 0 | 8.8986205 | 5.53252409 | 0 | 0 | 5.26063714 | 0 |
| M.polymnia664 | Serra do Mar | Tremembé / SP | Urban Park | 500-550 | 0 | 0 | 0 | 2.9964548 | 0 | 77.2110151 | 0 | 0 | 13.1212435 | 0 | 0 | 0.35608499 | 0 | 0 | 0 | 0 | 0 | 1.846734 | 0 | 0 | 0 | 2.3477962 | 0 | 0 | 2.12067133 | 0 |
| M.polymnia665 | Serra do Mar | Tremembé / SP | Urban Park | 500-550 | 0 | 0 | 0 | 5.67371665 | 0 | 73.217209 | 0 | 0 | 11.8430092 | 0 | 0 | 0 | 0 | 0 | 0 | 0 | 0 | 1.80468429 | 0 | 0 | 3.2520484 | 2.55198207 | 0 | 0 | 1.65735041 | 0 |
| M.polymnia667 | Serra do Mar | Tremembé / SP | Urban Park | 500-550 | 0 | 0 | 0 | 2.19323793 | 0 | 76.7622863 | 0 | 0 | 9.31193319 | 0 | 0 | 0 | 0 | 0 | 0 | 0 | 0 | 1.06136489 | 0 | 0 | 0 | 2.2846597 | 0 | 0 | 8.38651798 | 0 |
| M.polymnia833 | Bahia | Camacã / BA | Serra Bonita Private Reserve | 900-1200 | 0 | 0 | 0 | 9.36687735 | 0 | 64.0583884 | 0 | 0 | 18.3922813 | 0 | 0 | 0.73056347 | 0 | 0 | 0 | 0 | 0 | 0 | 0 | 0 | 0 | 5.06884983 | 0 | 0 | 2.38303962 | 0 |
| M.polymnia854 | Bahia | Camacã / BA | Serra Bonita Private Reserve | 900-1200 | 0 | 0 | 0 | 0 | 0 | 72.4732005 | 0 | 0 | 14.7582432 | 0 | 0 | 0.4780874 | 0 | 0 | 0 | 0 | 0 | 0 | 0 | 0 | 3.83463556 | 3.28049463 | 0 | 0 | 5.17533874 | 0 |
| M.polymnia900 | Bahia | Camacã / BA | Serra Bonita Private Reserve | 900-1200 | 0 | 0 | 0 | 1.41084366 | 0 | 86.0571083 | 0 | 0 | 9.54388731 | 0 | 0 | 0.47157343 | 0 | 0 | 0 | 0 | 0 | 0.90067489 | 0 | 0 | 0 | 1.03601031 | 0 | 0 | 0.57990212 | 0 |
| M.nesaea020 | Pernambuco | Recife / PE | Dois Irmãos State Park | 0-90 | 1.52572254 | 0 | 0 | 5.24432672 | 0 | 44.5626369 | 0 | 0 | 10.1267351 | 5.58784774 | 1.59271975 | 0 | 31.3600112 | 0 | 0 | 0 | 0 | 0 | 0 | 0 | 0 | 0 | 0 | 0 | 0 | 0 |
| M.nesaea035 | Pernambuco | Recife / PE | Dois Irmãos State Park | 0-90 | 0.33821466 | 0.13306722 | 0.08442876 | 1.54591636 | 0 | 48.668333 | 0 | 0 | 26.1962521 | 0.97397827 | 0 | 0 | 10.8023542 | 5.58187185 | 0 | 0 | 0 | 0 | 3.26903446 | 2.40654909 | 0 | 0 | 0 | 0 | 0 | 0 |
| M.nesaea036 | Pernambuco | Recife / PE | Dois Irmãos State Park | 0-90 | 0.31349946 | 0.14552659 | 0.11681349 | 1.4956552 | 0 | 72.1367891 | 0 | 0 | 12.6474082 | 0.71529957 | 0 | 0 | 4.43678864 | 4.06444378 | 0 | 0 | 0 | 0 | 1.97862165 | 1.94915426 | 0 | 0 | 0 | 0 | 0 | 0 |
| M.nesaea038 | Pernambuco | Recife / PE | Dois Irmãos State Park | 0-90 | 0.15338311 | 0.18760495 | 0.13020199 | 2.59710595 | 0 | 15.5581534 | 0 | 0 | 45.0832855 | 1.90221249 | 0 | 0 | 10.6471401 | 15.6697098 | 0 | 0 | 0 | 0 | 4.68990615 | 3.3812965 | 0 | 0 | 0 | 0 | 0 | 0 |
| M.nesaea039 | Pernambuco | Recife / PE | Dois Irmãos State Park | 0-90 | 0.10870219 | 0.03930754 | 0.0291563 | 0.7355774 | 0 | 79.9825674 | 0 | 0 | 10.8642061 | 0.79703566 | 0.16757587 | 0 | 2.47709684 | 1.26979241 | 0 | 0 | 0 | 0 | 1.75651129 | 1.77247102 | 0 | 0 | 0 | 0 | 0 | 0 |
| M.nesaea040 | Pernambuco | Recife / PE | Dois Irmãos State Park | 0-90 | 0.20016814 | 0.14663471 | 0.06540147 | 2.03564319 | 0 | 50.3294291 | 0 | 0 | 18.2613387 | 0.88415731 | 0.5889274 | 0 | 9.28768588 | 6.70928124 | 0 | 0 | 0 | 0 | 7.78531065 | 3.70602225 | 0 | 0 | 0 | 0 | 0 | 0 |
| M.nesaea043 | Pernambuco | Recife / PE | Dois Irmãos State Park | 0-90 | 0.5902113 | 0.18628157 | 0.05506498 | 2.41954446 | 0 | 31.3911179 | 0 | 0 | 32.0312501 | 1.0186932 | 0.80223424 | 0 | 7.74394019 | 10.2370252 | 0 | 0 | 0 | 0 | 8.65055345 | 4.87408334 | 0 | 0 | 0 | 0 | 0 | 0 |
| M.nesaea044 | Pernambuco | Recife / PE | Dois Irmãos State Park | 0-90 | 0.55445468 | 0.1612009 | 0.47202586 | 1.75752794 | 0 | 25.0544892 | 0 | 0 | 39.7925166 | 0.96280315 | 0.62234884 | 0 | 9.15891223 | 7.75568651 | 0 | 0 | 0 | 0 | 10.2535338 | 3.45450034 | 0 | 0 | 0 | 0 | 0 | 0 |
| M.nesaea045 | Pernambuco | Recife / PE | Dois Irmãos State Park | 0-90 | 0.23157719 | 0.10939794 | 0.07978331 | 1.24165094 | 0 | 74.4950508 | 0 | 0 | 14.5900225 | 0.6038832 | 0.33231768 | 0 | 3.88687802 | 2.38655513 | 0 | 0 | 0 | 0 | 1.28170406 | 0.76117922 | 0 | 0 | 0 | 0 | 0 | 0 |
| M.nesaea046 | Pernambuco | Recife / PE | Dois Irmãos State Park | 0-90 | 0.18724146 | 0.08390034 | 0 | 1.75335352 | 0 | 42.5794399 | 0 | 0 | 30.8738894 | 1.06610547 | 0.39054728 | 0 | 5.63316346 | 5.15455461 | 0 | 0 | 0 | 0 | 8.34567795 | 3.93212669 | 0 | 0 | 0 | 0 | 0 | 0 |
| M.nesaea087 | Pernambuco | Jaqueira / PE | Frei Caneca Private Reserve | 65-750 | 0 | 0 | 0 | 0 | 0 | 46.5063357 | 0 | 0 | 25.429765 | 2.59915127 | 0 | 0 | 4.97726985 | 9.41545418 | 0 | 0 | 0 | 0 | 3.99685721 | 7.07516681 | 0 | 0 | 0 | 0 | 0 | 0 |
| M.nesaea102 | Pernambuco | Jaqueira / PE | Frei Caneca Private Reserve | 65-750 | 0 | 0.14464449 | 0 | 0.91308019 | 0 | 50.6636059 | 0 | 0 | 26.7843712 | 0 | 0 | 0 | 10.9851887 | 7.9535418 | 1.62188134 | 0.93368629 | 0 | 0 | 0 | 0 | 0 | 0 | 0 | 0 | 0 | 0 |
| M.nesaea118 | Pernambuco | Jaqueira / PE | Frei Caneca Private Reserve | 65-750 | 0 | 0 | 0 | 0 | 0 | 81.1633652 | 0 | 0 | 17.1518647 | 0 | 0 | 0 | 0 | 0.41583589 | 0 | 1.26893418 | 0 | 0 | 0 | 0 | 0 | 0 | 0 | 0 | 0 | 0 |
| M.nesaea202 | Pernambuco | Jaqueira / PE | Frei Caneca Private Reserve | 65-750 | 0 | 0 | 0 | 0 | 0 | 92.2113636 | 0 | 0 | 0 | 0 | 5.8348718 | 0 | 0 | 0 | 0 | 0 | 0 | 0 | 0 | 1.95376459 | 0 | 0 | 0 | 0 | 0 | 0 |
| M.nesaea203 | Pernambuco | Jaqueira / PE | Frei Caneca Private Reserve | 65-750 | 0 | 0 | 0 | 0 | 0 | 53.8744099 | 0 | 0 | 33.3416348 | 0 | 0 | 0 | 8.18048358 | 3.53078494 | 0 | 1.07268675 | 0 | 0 | 0 | 0 | 0 | 0 | 0 | 0 | 0 | 0 |
| M.nesaea221 | Pernambuco | Jaqueira / PE | Frei Caneca Private Reserve | 65-750 | 0 | 0 | 0 | 0.27993289 | 0 | 93.6045275 | 0 | 0 | 0 | 0 | 0 | 0 | 1.67288721 | 0 | 0 | 0 | 3.94576115 | 0 | 0 | 0.49689124 | 0 | 0 | 0 | 0 | 0 | 0 |
| M.nesaea222 | Pernambuco | Jaqueira / PE | Frei Caneca Private Reserve | 65-750 | 0 | 0 | 0 | 0.18136872 | 0 | 70.4832876 | 0 | 0 | 19.2036469 | 0 | 0 | 0 | 3.86227704 | 4.50174246 | 0 | 0.97324084 | 0 | 0 | 0.34168013 | 0.45275629 | 0 | 0 | 0 | 0 | 0 | 0 |
| M.nesaea234 | Pernambuco | Jaqueira / PE | Frei Caneca Private Reserve | 65-750 | 0 | 0 | 0 | 4.9714E-08 | 0 | 59.6803655 | 0 | 0 | 26.6567765 | 0 | 0 | 0 | 7.51637575 | 4.13183701 | 0 | 0.18914715 | 0 | 0 | 1.42623378 | 0.39926428 | 0 | 0 | 0 | 0 | 0 | 0 |
| M.nesaea235 | Pernambuco | Jaqueira / PE | Frei Caneca Private Reserve | 65-750 | 0 | 0 | 0 | 0.95068788 | 0 | 60.6470534 | 0 | 0 | 11.367888 | 0 | 0 | 0 | 11.3809504 | 12.1622495 | 2.10827498 | 1.38289585 | 0 | 0 | 0 | 0 | 0 | 0 | 0 | 0 | 0 | 0 |
| M.nesaea236 | Pernambuco | Jaqueira / PE | Frei Caneca Private Reserve | 65-750 | 0 | 0.36932291 | 0 | 2.53410742 | 0 | 68.9102389 | 0 | 0 | 7.8613312 | 0 | 0 | 0 | 8.37015876 | 10.9336561 | 0 | 1.02118472 | 0 | 0 | 0 | 0 | 0 | 0 | 0 | 0 | 0 | 0 |
| M.nesaea533 | Pernambuco | São Lourenço da Ma | Tapacurá Ecological Station | 90-200 | 0.17159864 | 0 | 0 | 0.53690802 | 0 | 54.9852364 | 0 | 0 | 19.8689227 | 0 | 0 | 0 | 3.15558407 | 6.57707692 | 1.87827533 | 0.69560409 | 0 | 0 | 10.0700481 | 2.06074575 | 0 | 0 | 0 | 0 | 0 | 0 |
| M.nesaea536 | Pernambuco | São Lourenço da Ma | Tapacurá Ecological Station | 90-200 | 0 | 0 | 0 | 0.15454465 | 0 | 80.3805376 | 0 | 0.4203331 | 10.3003392 | 0 | 0.25796396 | 0 | 2.04237404 | 1.39763109 | 0.24334424 | 0.26952226 | 0 | 0 | 3.61897201 | 0.9144378 | 0 | 0 | 0 | 0 | 0 | 0 |
| M.nesaea537 | Pernambuco | São Lourenço da Ma | Tapacurá Ecological Station | 90-200 | 0.17758948 | 0 | 0 | 0.51496119 | 0 | 82.2010896 | 0 | 0.29852666 | 7.95694948 | 0 | 0.14295581 | 0 | 2.54203843 | 1.43715413 | 0 | 0 | 0 | 0 | 3.55634453 | 1.17239069 | 0 | 0 | 0 | 0 | 0 | 0 |
| M.nesaea538 | Pernambuco | São Lourenço da Ma | Tapacurá Ecological Station | 90-200 | 0 | 0 | 0 | 0.24257932 | 0 | 72.3208027 | 0 | 0.54985388 | 11.3750309 | 0 | 0.23403757 | 0 | 4.05225666 | 3.26565574 | 0 | 0.46672734 | 0 | 0 | 5.79222339 | 1.70083249 | 0 | 0 | 0 | 0 | 0 | 0 |
| M.nesaea539 | Pernambuco | São Lourenço da Ma | Tapacurá Ecological Station | 90-200 | 0 | 0 | 0 | 0.54563051 | 0 | 66.5919142 | 0 | 1.26983502 | 17.448529 | 0 | 0 | 0 | 4.31080425 | 5.47475637 | 1.40535676 | 0 | 0 | 0 | 1.89082447 | 1.06234948 | 0 | 0 | 0 | 0 | 0 | 0 |
| M.nesaea540 | Pernambuco | São Lourenço da Ma | Tapacurá Ecological Station | 90-200 | 0 | 0 | 0 | 0.27128333 | 0 | 76.9589273 | 0 | 0 | 10.1985806 | 0 | 0.38672858 | 0 | 1.88102054 | 3.5339065 | 0.2546119 | 0.62638047 | 0 | 0 | 5.72648277 | 0.16207804 | 0 | 0 | 0 | 0 | 0 | 0 |
| M.nesaea541 | Pernambuco | São Lourenço da Ma | Tapacurá Ecological Station | 90-200 | 0 | 0 | 0 | 0.18157334 | 0 | 57.7408749 | 0 | 0 | 20.5707512 | 0 | 0.47767806 | 0 | 2.53068235 | 4.17170471 | 2.5712897 | 0 | 0 | 0 | 9.88407036 | 1.87137534 | 0 | 0 | 0 | 0 | 0 | 0 |
| M.nesaea542 | Pernambuco | São Lourenço da Ma | Tapacurá Ecological Station | 90-200 | 0 | 0 | 0 | 1.11709317 | 0.32362685 | 68.9265684 | 0 | 0 | 11.9678201 | 0 | 0.4064465 | 0 | 4.66185672 | 6.95795162 | 0.71054373 | 0.57704831 | 0 | 0 | 2.77277846 | 1.57826609 | 0 | 0 | 0 | 0 | 0 | 0 |
| M.nesaea543 | Pernambuco | São Lourenço da Ma | Tapacurá Ecological Station | 90-200 | 0.12423537 | 0 | 0 | 0.21131605 | 0.09539051 | 61.5862993 | 0 | 0.38092025 | 12.9809013 | 0 | 0.36693071 | 0 | 2.28850126 | 1.85429498 | 1.00629475 | 0.28209576 | 0 | 0 | 15.9485028 | 2.87431691 | 0 | 0 | 0 | 0 | 0 | 0 |
| M.nesaea544 | Pernambuco | São Lourenço da Ma | Tapacurá Ecological Station | 90-200 | 0 | 0 | 0 | 0.26835783 | 0.16284207 | 76.3000568 | 0 | 0.29209251 | 7.83703832 | 0 | 0.2030904 | 0 | 2.46034377 | 4.65597284 | 0.3618209 | 0.36424184 | 0 | 0 | 5.01040837 | 2.08373435 | 0 | 0 | 0 | 0 | 0 | 0 |
| M.nesaea545 | Pernambuco | São Lourenço da Ma | Tapacurá Ecological Station | 90-200 | 0 | 0 | 0 | 0.75262587 | 0.17674278 | 74.01797 | 0 | 0.49784931 | 11.8039213 | 0 | 0.3675869 | 0 | 2.81624867 | 7.47172875 | 0.18717271 | 0.67701446 | 0 | 0 | 0.85010156 | 0.38103776 | 0 | 0 | 0 | 0 | 0 | 0 |
| M.nesaea277 | Pernambuco | Igarassu / PE | Usina São José | 45-150 | 0 | 0 | 0 | 0.05634414 | 0 | 77.7230607 | 0 | 1.07195327 | 13.9293281 | 0 | 0 | 0 | 2.78586211 | 3.85382121 | 0 | 0.57963049 | 0 | 0 | 0 | 0 | 0 | 0 | 0 | 0 | 0 | 0 |
| M.nesaea320 | Pernambuco | Igarassu / PE | Usina São José | 45-150 | 0 | 0 | 0 | 0.28043298 | 0 | 55.9353938 | 0 | 0.35165242 | 27.7710039 | 0 | 0 | 0 | 5.35201552 | 6.56824979 | 2.24475303 | 1.49649861 | 0 | 0 | 0 | 0 | 0 | 0 | 0 | 0 | 0 | 0 |
| M.nesaea496 | Pernambuco | Igarassu / PE | Usina São José | 45-150 | 0 | 0 | 0 | 0.70203145 | 0 | 60.4550456 | 0 | 1.46176474 | 23.2237326 | 0 | 0 | 0 | 5.95989483 | 3.57466127 | 0 | 2.20209271 | 0 | 0 | 1.1694511 | 1.25132571 | 0 | 0 | 0 | 0 | 0 | 0 |
| M.nesaea500 | Pernambuco | Igarassu / PE | Usina São José | 45-150 | 0 | 0 | 0 | 0.89808957 | 0 | 66.9952625 | 0 | 0.52823674 | 15.4414509 | 0 | 0 | 0 | 6.83414799 | 4.84191571 | 0 | 2.57491844 | 0 | 0 | 1.30796172 | 0.57801641 | 0 | 0 | 0 | 0 | 0 | 0 |
| M.nesaea505 | Pernambuco | Igarassu / PE | Usina São José | 45-150 | 0 | 0 | 0 | 0.45685696 | 0 | 70.701603 | 0 | 0.65172337 | 15.7417586 | 0 | 0 | 0 | 4.19645369 | 3.44652309 | 0.47547359 | 1.70939902 | 0 | 0 | 1.56804338 | 1.05216535 | 0 | 0 | 0 | 0 | 0 | 0 |
| M.nesaea507 | Pernambuco | Igarassu / PE | Usina São José | 45-150 | 0 | 0 | 0 | 0.90966346 | 0 | 63.8239869 | 0 | 0.40268144 | 11.8497254 | 0 | 0 | 0 | 4.33811877 | 5.72234753 | 0.45652846 | 1.170253 | 0 | 0 | 6.74923171 | 4.5774633 | 0 | 0 | 0 | 0 | 0 | 0 |
| M.nesaea508 | Pernambuco | Igarassu / PE | Usina São José | 45-150 | 0 | 0 | 0 | 1.32613139 | 0 | 59.1077337 | 0 | 0.54179944 | 27.9673951 | 0 | 0 | 0 | 3.27765349 | 1.58565487 | 0 | 3.17303925 | 0 | 0 | 1.99220816 | 1.02838466 | 0 | 0 | 0 | 0 | 0 | 0 |
| M.nesaea509 | Pernambuco | Igarassu / PE | Usina São José | 45-150 | 0 | 0 | 0 | 1.09334491 | 0 | 62.594211 | 0 | 0.54933791 | 21.0720504 | 0 | 0 | 0 | 2.8466889 | 2.24611469 | 0 | 2.49399752 | 0 | 0 | 4.44371797 | 2.66053665 | 0 | 0 | 0 | 0 | 0 | 0 |
| M.nesaea510 | Pernambuco | Igarassu / PE | Usina São José | 45-150 | 0 | 0 | 0 | 0.94581047 | 0 | 63.452466 | 0 | 0.28478783 | 14.0836797 | 0 | 0 | 0 | 5.2454887 | 4.99973604 | 0.8713368 | 2.69204388 | 0 | 0 | 4.13124903 | 3.29340156 | 0 | 0 | 0 | 0 | 0 | 0 |
| M.nesaea552 | Pernambuco | Igarassu / PE | Usina São José | 45-150 | 0 | 0 | 0 | 2.00585217 | 8.65E-07 | 37.4260056 | 0 | 8.65E-07 | 23.9752062 | 0.84601945 | 0 | 0 | 4.42277181 | 22.0483151 | 0 | 0 | 0 | 0 | 3.96708827 | 5.30873977 | 0 | 0 | 0 | 0 | 0 | 0 |
| M.nesaea551 | Pernambuco | Igarassu / PE | Usina São José | 45-150 | 0 | 0 | 0 | 0.20365179 | 0.69331635 | 44.463291 | 0 | 3.6827E-07 | 24.0967616 | 0.69772237 | 0 | 0 | 7.78218469 | 7.44895376 | 0 | 0 | 0 | 0 | 7.86883865 | 6.74527935 | 0 | 0 | 0 | 0 | 0 | 0 |
| M.nesaea388 | Brejos Nordestinos | Caruara / PE | Serro dos Cavalos Municipal Park | 780-950 | 0 | 0 | 0 | 0 | 0 | 16.1119837 | 0 | 3.72837702 | 22.1410509 | 0 | 0 | 0 | 12.1463365 | 4.67309381 | 0 | 3.54555642 | 0 | 0 | 18.1891243 | 19.4644773 | 0 | 0 | 0 | 0 | 0 | 0 |
| M.nesaea390 | Brejos Nordestinos | Caruara / PE | Serro dos Cavalos Municipal Park | 780-950 | 0 | 0 | 0 | 0 | 0 | 96.8852822 | 0 | 1.81226163 | 0 | 0 | 0 | 0 | 0 | 0 | 0 | 0 | 0 | 0 | 0 | 1.30245618 | 0 | 0 | 0 | 0 | 0 | 0 |
| M.nesaea394 | Brejos Nordestinos | Caruara / PE | Serro dos Cavalos Municipal Park | 780-950 | 0 | 0 | 0 | 0.18225358 | 0 | 62.8233766 | 0 | 1.37105538 | 27.628429 | 0 | 0 | 0 | 2.48969723 | 3.71747642 | 0.16286055 | 1.62485123 | 0 | 0 | 0 | 0 | 0 | 0 | 0 | 0 | 0 | 0 |
| M.nesaea395 | Brejos Nordestinos | Caruara / PE | Serro dos Cavalos Municipal Park | 780-950 | 0 | 0 | 0 | 0.49473308 | 0 | 64.9160115 | 0 | 1.92919768 | 23.4516103 | 0 | 0 | 0 | 2.9861818 | 4.39505189 | 0 | 1.82721384 | 0 | 0 | 0 | 0 | 0 | 0 | 0 | 0 | 0 | 0 |
| M.nesaea396 | Brejos Nordestinos | Caruara / PE | Serro dos Cavalos Municipal Park | 780-950 | 0 | 0 | 0 | 0.25437375 | 0 | 98.9859345 | 0 | 0.7596918 | 0 | 0 | 0 | 0 | 0 | 0 | 0 | 0 | 0 | 0 | 0 | 0 | 0 | 0 | 0 | 0 | 0 | 0 |
| M.nesaea397 | Brejos Nordestinos | Caruara / PE | Serro dos Cavalos Municipal Park | 780-950 | 0 | 0 | 0 | 0.65521053 | 0 | 82.3308724 | 0 | 0.63471038 | 8.64324002 | 0 | 0 | 0 | 2.99146168 | 3.75125446 | 0 | 0.99325054 | 0 | 0 | 0 | 0 | 0 | 0 | 0 | 0 | 0 | 0 |
| M.nesaea398 | Brejos Nordestinos | Caruara / PE | Serro dos Cavalos Municipal Park | 780-950 | 0 | 0 | 0 | 0.21029195 | 0 | 77.6793295 | 0 | 1.64913193 | 13.7271934 | 0 | 0 | 0 | 2.61140116 | 3.62859151 | 2.0052E-07 | 0.49406037 | 0 | 0 | 0 | 0 | 0 | 0 | 0 | 0 | 0 | 0 |
| M.nesaea399 | Brejos Nordestinos | Caruara / PE | Serro dos Cavalos Municipal Park | 780-950 | 0 | 0 | 0 | 0 | 0 | 82.2137399 | 0 | 1.25944171 | 10.4731659 | 0 | 0 | 0 | 3.00554347 | 2.5148674 | 0 | 0.53324166 | 0 | 0 | 0 | 0 | 0 | 0 | 0 | 0 | 0 | 0 |
| M.nesaea405 | Brejos Nordestinos | Caruara / PE | Serro dos Cavalos Municipal Park | 780-950 | 0 | 0 | 0 | 0.67301442 | 0 | 67.3947729 | 0 | 1.58340446 | 18.8843408 | 0 | 0 | 0 | 7.05015909 | 3.37927139 | 0 | 1.03503695 | 0 | 0 | 0 | 0 | 0 | 0 | 0 | 0 | 0 | 0 |
| M.nesaea406 | Brejos Nordestinos | Caruara / PE | Serro dos Cavalos Municipal Park | 780-950 | 0 | 0 | 0 | 0 | 0 | 72.8401964 | 0 | 0.78541377 | 20.4889964 | 0 | 0 | 0 | 2.49715257 | 1.56941933 | 0 | 1.17777363 | 0 | 0 | 0.42963048 | 0.21141749 | 0 | 0 | 0 | 0 | 0 | 0 |
| M.nesaea_BA839 | Bahia | Camacã / BA | Serra Bonita Private Reserve | 900-1200 | 8.0992228 | 0 | 0 | 5.30904006 | 0 | 63.7847457 | 0 | 0 | 11.9499554 | 0 | 0 | 0 | 0 | 0 | 0 | 0 | 0 | 0 | 0 | 10.857036 | 0 | 0 | 0 | 0 | 0 | 0 |
| M.nesaea_BA859 | Bahia | Camacã / BA | Serra Bonita Private Reserve | 900-1200 | 0 | 0 | 0 | 0.81582799 | 1.62021591 | 45.2250879 | 0 | 3.39706368 | 24.5555012 | 0 | 0 | 0 | 7.6459486 | 6.30211393 | 0 | 2.10563671 | 0 | 0 | 0 | 7.92558821 | 0 | 0 | 0 | 0.4070159 | 0 | 0 |
| M.nesaea_BA860 | Bahia | Camacã / BA | Serra Bonita Private Reserve | 900-1200 | 0 | 0 | 0 | 0.25579374 | 0 | 45.5849012 | 0 | 1.61585511 | 23.1059642 | 0 | 0 | 0 | 3.5116344 | 5.18359937 | 0 | 0.89076369 | 0 | 0 | 10.7065303 | 8.75543942 | 0 | 0 | 0 | 0.38951844 | 0 | 0 |
| M.nesaea_BA867 | Bahia | Camacã / BA | Serra Bonita Private Reserve | 900-1200 | 1.30278721 | 0 | 0 | 0.76998378 | 0 | 63.8785459 | 0 | 0.7692554 | 10.3407428 | 0 | 0 | 0 | 5.7143533 | 4.93774402 | 0 | 1.29358238 | 0 | 0 | 5.9372679 | 5.05573724 | 0 | 0 | 0 | 0 | 0 | 0 |
| M.nesaea_BA870 | Bahia | Camacã / BA | Serra Bonita Private Reserve | 900-1200 | 0 | 0 | 0 | 0 | 0 | 53.9770754 | 2.3678E-07 | 3.98959814 | 25.5440064 | 0 | 0 | 0 | 4.14023545 | 4.06639219 | 0 | 0 | 0 | 0 | 2.89081409 | 5.39187807 | 0 | 0 | 0 | 0 | 0 | 0 |
| M.nesaea_BA896 | Bahia | Camacã / BA | Serra Bonita Private Reserve | 900-1200 | 0 | 0.28507915 | 0 | 1.59769373 | 0 | 54.9955068 | 1.0769E-07 | 7.55753194 | 13.8430969 | 0 | 0 | 0 | 4.17280083 | 1.8375579 | 0 | 1.23024519 | 0 | 0 | 6.41772642 | 8.0627611 | 0 | 0 | 0 | 0 | 0 | 0 |
| M.nesaea_BA901 | Bahia | Camacã / BA | Serra Bonita Private Reserve | 900-1200 | 0 | 0 | 0 | 0.62798312 | 0 | 67.9284588 | 1.9509E-07 | 1.9509E-07 | 20.6426119 | 0 | 0 | 0 | 5.32497448 | 3.36597346 | 0 | 0.79823777 | 0 | 0 | 1.9509E-07 | 1.3117599 | 0 | 0 | 0 | 0 | 0 | 0 |
| M.nesaea_BA902 | Bahia | Camacã / BA | Serra Bonita Private Reserve | 900-1200 | 0 | 0 | 0 | 0.32148326 | 0 | 57.5366063 | 7.8267E-08 | 3.53543235 | 25.8334977 | 0 | 0 | 0 | 3.91160995 | 2.06339339 | 0 | 1.18335986 | 0 | 0 | 2.89581243 | 2.71880463 | 0 | 0 | 0 | 0 | 0 | 0 |
| M.nesaea_BA903 | Bahia | Camacã / BA | Serra Bonita Private Reserve | 900-1200 | 0 | 0 | 0 | 0.62511315 | 7.8446E-08 | 54.5306714 | 7.8446E-08 | 1.72716156 | 16.5301704 | 0 | 0 | 0 | 4.28834424 | 2.37328454 | 0 | 1.08924402 | 0 | 0 | 9.14067746 | 9.69533304 | 0 | 0 | 0 | 0 | 0 | 0 |
| M.nesaea_BA904 | Bahia | Camacã / BA | Serra Bonita Private Reserve | 900-1200 | 0 | 0 | 0 | 0.47785162 | 0 | 47.7019442 | 3.529E-07 | 4.22305855 | 20.7101864 | 0 | 0 | 0 | 14.9425637 | 8.36740363 | 0 | 0 | 0 | 0 | 0 | 3.57699156 | 0 | 0 | 0 | 0 | 0 | 0 |
| M.nesaea_BA905 | Bahia | Camacã / BA | Serra Bonita Private Reserve | 900-1200 | 0 | 0 | 0 | 1.58043559 | 0 | 64.7703745 | 4.805E-07 | 4.99377903 | 9.88951589 | 0 | 0 | 0 | 9.77176831 | 5.31952502 | 0 | 0 | 0 | 0 | 0 | 3.67460121 | 0 | 0 | 0 | 0 | 0 | 0 |
| M.nesaea_BA907 | Bahia | Camacã / BA | Serra Bonita Private Reserve | 900-1200 | 0 | 0 | 0 | 0.08818658 | 0 | 61.3384521 | 2.3176E-07 | 7.59782526 | 13.5358067 | 0 | 0 | 0 | 7.10769804 | 3.64528046 | 0 | 0 | 0 | 0 | 2.65229621 | 4.03445442 | 0 | 0 | 0 | 0 | 0 | 0 |
| M.nesaea_BA908 | Bahia | Camacã / BA | Serra Bonita Private Reserve | 900-1200 | 0 | 0 | 0 | 0.19041305 | 0 | 83.2554207 | 1.0259E-07 | 1.66668529 | 11.1654831 | 0 | 0 | 0 | 1.85920208 | 1.86279558 | 0 | 0 | 0 | 0 | 0 | 0 | 0 | 0 | 0 | 0 | 0 | 0 |
| M.lysimnia589 | Serra do Mar | Jundiaí / SP | Serra do Japi Biological Reserve | 900-1200 | 0 | 0 | 0 | 0.14052936 | 0 | 5.31147495 | 0.43210003 | 0.35093199 | 0 | 0 | 0 | 0 | 0.27990906 | 0 | 0 | 0 | 0.79653893 | 0 | 0 | 0 | 0 | 0 | 0 | 88.4268811 | 0 | 4.26163461 |
| M.lysimnia591 | Serra do Mar | Jundiaí / SP | Serra do Japi Biological Reserve | 900-1200 | 0 | 0 | 0 | 0.23464036 | 0 | 0 | 0.81789675 | 1.03102831 | 0 | 0 | 0 | 0 | 0 | 0 | 0 | 0 | 6.81580343 | 0 | 0 | 0 | 0 | 0 | 0 | 88.9122651 | 0 | 2.18836602 |
| M.lysimnia596 | Serra do Mar | Jundiaí / SP | Serra do Japi Biological Reserve | 900-1200 | 0 | 0 | 0 | 0.03915117 | 0 | 6.72422316 | 0.07998742 | 0.06936629 | 0 | 0 | 0 | 0 | 0.09175904 | 0 | 0 | 0 | 0.85733598 | 0 | 0 | 0 | 0 | 0 | 2.28933707 | 86.9884168 | 0 | 2.86042307 |
| M.lysimnia597 | Serra do Mar | Jundiaí / SP | Serra do Japi Biological Reserve | 900-1200 | 0 | 0 | 0 | 0.00645283 | 0 | 2.07448828 | 0.34946033 | 0.16651877 | 0 | 0 | 0 | 0 | 0.25697011 | 0 | 0 | 0 | 2.29751036 | 0 | 0 | 0 | 0 | 0 | 1.71138566 | 90.5609475 | 0 | 2.57626616 |
| M.lysimnia636 | Serra do Mar | Jundiaí / SP | Serra do Japi Biological Reserve | 900-1200 | 0 | 0 | 0 | 0.10986487 | 0 | 6.59742104 | 0.04274752 | 0.07053895 | 0 | 0 | 0 | 0 | 0.1117124 | 0 | 0 | 0 | 0.77332049 | 0 | 0 | 0 | 0 | 0 | 0 | 89.9841975 | 0 | 2.31019722 |
| M.lysimnia650 | Serra do Mar | Jundiaí / SP | Serra do Japi Biological Reserve | 900-1200 | 0 | 0 | 0 | 0 | 0 | 1.3992E-06 | 1.12946198 | 1.3992E-06 | 0 | 0 | 0 | 0 | 0.41978898 | 0 | 0 | 0 | 2.14239862 | 0 | 0 | 0 | 0 | 0 | 0 | 93.3740169 | 0 | 2.93433072 |
| M.lysimnia656 | Serra do Mar | Jundiaí / SP | Serra do Japi Biological Reserve | 900-1200 | 0 | 0 | 0 | 0.16094917 | 0 | 14.045796 | 0.2052649 | 0.24658731 | 0 | 0 | 0 | 0 | 0.24358273 | 0 | 0 | 0 | 2.3324385 | 0 | 0 | 0 | 0 | 0 | 2.42759821 | 78.3918119 | 0 | 1.94597129 |
| M.lysimnia_BA827 | Bahia | Camacã / BA | Serra Bonita Private Reserve | 900-1200 | 0 | 0 | 0 | 0 | 0 | 1.58067177 | 0.47791712 | 0.27413119 | 0 | 0 | 0 | 0 | 0 | 0 | 0 | 0 | 0.71741309 | 0 | 0 | 0.07924479 | 0 | 0 | 0.98682932 | 92.9024271 | 0 | 2.98136562 |
| M.lysimnia_BA845 | Bahia | Camacã / BA | Serra Bonita Private Reserve | 900-1200 | 0 | 0 | 0 | 0 | 0 | 17.1109466 | 0 | 0.15548734 | 0.27432519 | 0 | 0 | 0 | 0.30716948 | 0 | 0 | 0 | 0.35723725 | 0 | 0 | 0.05328654 | 0 | 0 | 1.18812441 | 77.7457301 | 0 | 2.80769311 |
| M.lysimnia_BA855 | Bahia | Camacã / BA | Serra Bonita Private Reserve | 900-1200 | 0 | 0 | 0 | 0 | 0 | 0.21924994 | 0 | 0.01371651 | 0 | 0 | 0 | 0 | 0 | 0 | 0 | 0 | 0.19125711 | 0 | 0 | 0.60763643 | 0 | 0 | 0.36583366 | 97.8774511 | 0 | 0.72485522 |
| M.lysimnia_BA858 | Bahia | Camacã / BA | Serra Bonita Private Reserve | 900-1200 | 0 | 0 | 0 | 0 | 0 | 0.12550766 | 0 | 0 | 0 | 0 | 0 | 0 | 0 | 0 | 0 | 0 | 0.03738595 | 0 | 0 | 0 | 0 | 0 | 0 | 99.703839 | 0 | 0.13326741 |
| M.lysimnia_BA877 | Bahia | Camacã / BA | Serra Bonita Private Reserve | 900-1200 | 0 | 0 | 0 | 0 | 0 | 0.34411777 | 1.19130692 | 0 | 0 | 0 | 0 | 0 | 0 | 0 | 0 | 0 | 1.26151335 | 0 | 0 | 0 | 0 | 0 | 0 | 96.5346245 | 0 | 0.66843741 |
| M.lysimnia_BA878 | Bahia | Camacã / BA | Serra Bonita Private Reserve | 900-1200 | 0 | 0 | 0 | 0 | 0 | 1.95105027 | 0 | 0 | 0 | 0 | 0 | 0 | 0 | 0 | 0 | 0 | 0.13159952 | 0 | 0 | 0 | 0 | 0 | 0 | 97.6140244 | 0 | 0.30332581 |
| M.lysimnia_BA879 | Bahia | Camacã / BA | Serra Bonita Private Reserve | 900-1200 | 0 | 0 | 0 | 0 | 0 | 2.17434067 | 0 | 0 | 0 | 0 | 0 | 0 | 0 | 0 | 0 | 0 | 0 | 0 | 0 | 0 | 0 | 0 | 0 | 97.402953 | 0 | 0.42270629 |
| M.lysimnia_BA880 | Bahia | Camacã / BA | Serra Bonita Private Reserve | 900-1200 | 0 | 0 | 0 | 0 | 0 | 1.2737009 | 0.02613134 | 0 | 0 | 0 | 0 | 0 | 0 | 0 | 0 | 0 | 0.1816224 | 0 | 0 | 0 | 0 | 0 | 0 | 97.9720837 | 0 | 0.54646164 |
| M.lysimnia_BA892 | Bahia | Camacã / BA | Serra Bonita Private Reserve | 900-1200 | 0 | 0 | 0 | 0 | 0 | 0.98577966 | 0 | 0 | 0 | 0 | 0 | 0 | 0 | 0 | 0 | 0 | 0 | 0 | 0 | 0 | 0 | 0 | 0 | 98.7128486 | 0 | 0.30137174 |
| M.lysimnia_BA898 | Bahia | Camacã / BA | Serra Bonita Private Reserve | 900-1200 | 0 | 0 | 0 | 0 | 0 | 1.14553891 | 0.07527714 | 0.03599215 | 0 | 0 | 0 | 0 | 0.04596619 | 0 | 0 | 0 | 0 | 0 | 0 | 0 | 0 | 0 | 0 | 98.3244313 | 0 | 0.37279429 |
| M.lysimnia_BA906 | Bahia | Camacã / BA | Serra Bonita Private Reserve | 900-1200 | 0 | 0 | 0 | 0 | 0 | 0.22879488 | 0 | 0 | 0 | 0 | 0 | 0 | 0.01341094 | 0 | 0 | 0 | 0.03162097 | 0 | 0 | 0 | 0 | 0 | 0 | 99.0613974 | 0 | 0.66477584 |
| M.lysimnia_BA909 | Bahia | Camacã / BA | Serra Bonita Private Reserve | 900-1200 | 0 | 0 | 0 | 0 | 0 | 2.43979649 | 0 | 0 | 0 | 0 | 0 | 0 | 0 | 0 | 0 | 0 | 0 | 0 | 0 | 0 | 0 | 0 | 0 | 96.4758354 | 0 | 1.08436806 |

**Table S4. Pheromone results summary**

Median relative concentration values of chemical compounds identified in androconial extracts of *Mechanitis* individuals from the Brazilian Atlantic forest. Values between brackets indicate the number of analysed samples in which the compound was present, per investigated species. *RI*: Retention Index.

| ***RI*** | **Compound identity or mass spectra** | ***M. nesaea* N = 65** | ***M. l. lysimnia* N = 19** | ***M. polymnia casabranca* N = 8** |
|---|---|---|---|---|
| **1150** | 2,6,6-Trimethyl-2-cyclohexene-1,4-dione | 0.18 [15] | - | - |
| **1252** | *m/z:* 98, 41, 43, 69, 39 | 0.12 [12] | - | - |
| **1264** | *m/z:* 43, 41, 57, 55, 39 | 0.06 [8] | - | - |
| **1318** | 4-Hydroxy-3,5,5-trimethylcyclohex-2-enone | 0.45 [56] | 0.12 [6] | 2.19 [7] |
| **1365** | Eugenol | 0.11 [8] | - | - |
| **1486** | Hydroxydanaidal | 44.68 [65] | 1.76 [18] | 74.99 [8] |
| **1543** | *m/z:* 98, 43, 71, 41, 57 | - | 0.35 [11] | - |
| **1551** | Dihydroactinidiolide | 0.67 [40] | 0.15 [11] | - |
| **1588** | Methyl hydroxydanaidoate | 11.81 [61] | 0.27 [1] | 12.48 [8] |
| **1599** | Hexadecane | 0.77 [13] | - | - |
| **1619** | *m/z:* 86, 43, 71, 41, 84 | 0.27 [16] | - | - |
| **1673** | *m/z* 120, 43, 65, 44, 55 | - | - | 0.47 [5] |
| **1749** | Methyl farnesoate isomer | 3.09 [61] | 0.23 [9] | - |
| **1792** | Methyl (*E,E*)-farnesoate | 2.97 [60] | - | - |
| **1804** | 4-Hydroxy-3,5,5-trimethyl-4-[3-oxo-1-butenyl]-2-cyclohexen-1-one | 0.41 [18] | - | - |
| **1843** | Methyl oxodanaidoate | 0.77 [39] | - | - |
| **1852** | *m/z:* 57, 45, 55, 43, 71 | 2.95 [1] | 0.76 [15] | - |
| **1950** | *m/z* 57, 43, 55, 56, 85 | - | - | 1.15 [6] |
| **2145** | *m/z* 79, 67, 55, 41, 95 | 2.69 [41] | - | - |
| **2160** | Octadecanoic acid | 1.66 [48] | 0.08 [3] | - |
| **2172** | Ethyl linolenate | - | - | 3.83 [3] |
| **2220** | Phytol acetate <(*E*)-> | - | - | 2.45 [8] |
| **2572** | Hexacosene | 4.26 [1] | 1.44 [6] | - |
| **2675** | Heptacosene | 0.27 [2] | 96.27 [19] | - |
| **2832** | Squalene | - | - | 2.25 [8] |
| **2873** | Nonacosene | - | 1.08 [19] | - |

**Table S5. Reference genome statistics**

Assembled chromosomes and completeness of the genomes generated in this study, with primary and alternative haplotype for each individual. Individuals with a different alternative autosome number are heterozygous for chromosomal rearrangements. All sex-chromosomes are included in the primary assembly. Estimated k-mer coverage included for PacBio, HiC-data and, where applicable, 10X-data.

| Species | NCBI accession | Autos. | Sex-chr | Sex | Genome size (Mb) | In chr (Mb) | % | Scaffold N50 (Mb) | Contigs N50 (Mb) | Gaps | Scaffolds | PacBio | HiC | 10X |
|---|---|---|---|---|---|---|---|---|---|---|---|---|---|---|
| *Mec. polymnia* | In progress | 14 | W and Z | F | 295.9 | 293.1 | 99 | 18.4 | 1.4 | 0.03% | 122 | 29.03 | 136.9 | |
| | In progress | 15 | | | 264.4 | 261.7 | 99 | 17.5 | 1.2 | 0.03% | 104 | | | |
| *Mec. menapis* | In progress | 19 | W and Z | F | 305.3 | 301.8 | 99 | 15.7 | 0.5 | 0.08% | 213 | 13.61 | 54.46 | |
| | In progress | 19 | | | 237.4 | 231.5 | 98 | 13.4 | 0.5 | 0.07% | 140 | | | |
| *Mec. messenoides* | In progress | 14 | W and Z | F | 311.1 | 307.5 | 99 | 19.1 | 8.6 | 0.01% | 102 | 52.83 | 164.6 | |
| | In progress | 15 | | | 269.0 | 266.2 | 99 | 18.7 | 5.9 | 0.01% | 53 | | | |
| *Mec. macrinus* | In progress | 21 | W1, W2, W3, Z1 and Z2 | F | 320.1 | 319.4 | 100 | 13.6 | 6.6 | 0.01% | 53 | 12.2 | 73.91 | |
| | In progress | 21 | | | 241.6 | 241.0 | 100 | 12.5 | 6.6 | 0.00% | 45 | | | |
| *Mec. mazaeus* | In progress | 12 | W1, W2, Z1, Z2 | F | 318.5 | 315.7 | 99 | 20.9 | 9.3 | 0.01% | 88 | 45.32 | 196.9 | |
| | In progress | 12 | | | 244.5 | 242.9 | 99 | 20.6 | 15.9 | 0.00% | 46 | | | |
| *Mel. ludovica* | In progress | 21 | Z\|Z | M | 661.4 | 627.9 | 95 | 33.1 | 4.6 | 0.01% | 176 | 57.19 | 60.47 | |
| | In progress | 23 | | | 647.8 | 625.1 | 97 | 30.3 | 3.8 | 0.01% | 282 | | | |
| *Mel. isocomma* | In progress | 13 | Z\|Z1 and Z2 | M | 527.6 | 519.4 | 98 | 39.4 | 4.9 | 0.01% | 78 | 21.4 | 33.3 | |
| | In progress | 14 | | | 504.8 | 498.2 | 99 | 34.9 | 4.1 | 0.01% | 94 | | | |
| *Mel. mothone* | In progress | 13 | W1, W2, Z1 and Z2 | F | 559.0 | 552.3 | 99 | 37.0 | 5.0 | 0.01% | 185 | 20.74 | 87.54 | |
| | In progress | 14 | | | 461.6 | 457.7 | 99 | 35.6 | 4.4 | 0.01% | 92 | | | |
| *Mel. marsaeus* | In progress | 13 | W and Z | F | 516.2 | 510.6 | 99 | 39.5 | 17.2 | 0.00% | 143 | 25.7 (CLR) | 19.45 | 15.11 |
| | In progress | 14 | | | 490.1 | 483.7 | 99 | 38.8 | 22.0 | 0.00% | 141 | | | |
| *Mel. menophilus* | In progress | 19 | W, Z1, Z2 | F | 541.2 | 536.4 | 99 | 22.9 | 14.5 | 0.00% | 130 | 24.86 (CLR) | 20.19 | 11.03 |
| | In progress | 21 | | | 453.2 | 449.8 | 99 | 21.2 | 10.7 | 0.00% | 81 | | | |

**Table S6. BUSCO results**

Result from BUSCO analysis of the genomes included in the analyses. All results are in percent of the total number of genes (5286 genes) in the BUSCO *lepidoptera_odb10* database.

| Species | | Complete | Single | Duplicated | Fragmented | Missing | Internal stop codon (%) |
|---|---|---|---|---|---|---|---|
| *Mec. polymnia* | Hap1 | 99 | 98.1 | 0.9 | 0.5 | 0.5 | 2.7 |
| | Hap2 | 91.8 | 91.3 | 0.5 | 0.5 | 7.7 | 2.9 |
| *Mec. menapis* | Hap1 | 96.6 | 96.1 | 0.5 | 1.2 | 2.2 | 3.4 |
| | Hap2 | 75.4 | 74.9 | 0.5 | 1.3 | 23.3 | 3.5 |
| *Mec. messenoides* | Hap1 | 99.2 | 98.7 | 0.5 | 0.5 | 0.3 | 2.8 |
| | Hap2 | 91.9 | 91.4 | 0.5 | 0.5 | 7.6 | 2.9 |
| *Mec. macrinus* | Hap1 | 99.1 | 91.6 | 7.5 | 0.5 | 0.4 | 2.8 |
| | Hap2 | 79.3 | 78.9 | 0.4 | 0.4 | 20.3 | 2.7 |
| *Mec. mazaeus* | Hap1 | 99.2 | 89.3 | 9.9 | 0.5 | 0.3 | 2.6 |
| | Hap2 | 82.5 | 82.2 | 0.3 | 0.4 | 17.1 | 2.7 |
| *Mel. ludovica* | Hap1 | 98.8 | 97.2 | 1.6 | 0.7 | 0.5 | 2.6 |
| | Hap2 | 98.4 | 97.4 | 1 | 0.8 | 0.8 | 2.6 |
| *Mel. isocomma* | Hap1 | 98.1 | 96.3 | 1.8 | 0.6 | 1.3 | 2.7 |
| | Hap2 | 97.9 | 96.6 | 1.3 | 0.7 | 1.4 | 2.7 |
| *Mel. mothone* | Hap1 | 98.9 | 90.6 | 8.3 | 0.6 | 0.5 | 2.6 |
| | Hap2 | 85.5 | 84.8 | 0.7 | 0.5 | 14 | 2.5 |
| *Mel. marsaeus* | Hap1 | 99 | 98.1 | 0.9 | 0.6 | 0.4 | 2.8 |
| | Hap2 | 93.1 | 92.4 | 0.7 | 0.5 | 6.4 | 2.6 |
| *Mel. menophilus* | Hap1 | 99 | 87.3 | 11.7 | 0.6 | 0.4 | 2.7 |
| | Hap2 | 82.6 | 81.8 | 0.8 | 0.5 | 16.9 | 2.7 |

**Table S7. Gene count per conserved syntenic block**

Total number of BUSCO-genes included per taxa, mean, standard deviation, median and maximum number of genes per ancestral syntenic block per taxa.

| Species | Total nr genes | Mean | SD | Median | Max |
|---|---|---|---|---|---|
| *Melitaea cinxia* | 5076.00 | 163.74 | 77.02 | 167 | 315 |
| *Danaus plexippus* | 4963.00 | 160.10 | 75.98 | 164 | 311 |
| *Mec. macrinus* | 5390.00 | 45.29 | 42.93 | 34 | 226 |
| *Mec. mazaeus* | 5506.00 | 46.27 | 45.08 | 34 | 226 |
| *Mec. menapis* | 4754.00 | 45.28 | 45.29 | 32 | 230 |
| *Mec. messenoides* | 5014.00 | 45.58 | 44.78 | 31.5 | 244 |
| *Mec. polymnia* | 4998.00 | 44.23 | 42.97 | 32 | 226 |
| *Mel. ludovica* | 4956.00 | 39.02 | 31.10 | 32 | 225 |
| *Mel. isocomma* | 4899.00 | 42.60 | 33.68 | 33 | 226 |
| *Mel. mothone* | 5383.00 | 44.12 | 34.14 | 35 | 226 |
| *Mel. menophilus* | 5532.00 | 42.23 | 33.42 | 33 | 227 |
| *Mel. marsaeus* | 4998.00 | 42.72 | 33.88 | 34 | 227 |

**Supplementary Table 8. Sex-chromosomes and autosomal polymorphisms.** Constituent parts of the sex-chromosomes (Chr) in the ten *Mechanitis* and *Melinaea* genome assemblies based on BUSCO-genes and whole genome alignment (W). Fusions of autosomes or autosomal parts (Ancestral element) with the canonical Z-chromosome (Z*) are indicated with "+" signs and the number of the ancestral autosome. The "w*" in the W-ancestral column is a common W-element in all *Mechanitis* assemblies with high sequence homology but no BUSCO-genes. Autosomal polymorphic fusion-fissions with homologous chromosomes separated with "=" and the split chromosomes in one haplotype are separated by "|". Ancestral elements in parenthesis ( ) are only found in the split chromosomes, elements in square brackets [ ] are only found in the fused chromosome and "|" represent the breakpoint between the split chromosomes.

| **Species** | **Z** | **Ancestral elements** | **W** | **Ancestral elements** | **Autosomal polymorphism** | **Ancestral elements** |
|---|---|---|---|---|---|---|
| *Melinaea Ludovica (male)* | Z | Z*+10 | NA | NA | 2 = 13 \| 17 | 21+20+2+3+10 \| 8+22 |
| | | | | | 7 = 20 \| 18 | 11+2+26+5 \| 14+17 |
| | | | | | 10 = 23 \| 14 | 12+8+16+1+21 \| 21+4+10 |
| | | | | | 1,18,14,16 = 1,10,12 | Complex rearrangement |
| *Melinaea isocomma (male)* | Z = Z2 \| Z1 | 12+13+16+[24]+ 8+20+11+9 \| Z*+10 | NA | NA | 1 = 3 \|13 | 17+18+9+11+7+27+13+2 \| 8+4+20 +(11) |
| *Melinaea mothone* | Z1 \| Z2 | Z*+10 \| 27+7+17+4+21+6 | W1 \| W2 | 27+7+17+4+10 \| 21+6 | 2 = 11 \| 14 | 1+21+20+2+3+15 \| 18+12+17+(2) |
| *Melinaea menophilus* | Z1 \| Z2 | Z*+10 \| 3+5+6+16+14+12+18 | W | 10+3+5+6+16+14+12+18 | 1 = 18 \| 4 | 19+24+4+14 \|14+30+6+3+11+2+26 |
| | | | | | 2 = 21 \| 13 | 27+7+17+[18]+4+(3) \| 11+9+[Z]+(18+17) |
| *Melinaea marsaeus* | Z | Z*+10 | W | 10 | 2 = 10 \| 14 | 3+5+6+16+14+12+18 \| 4+17+7+27 |
| *Mechanitis macrinus* | Z1 \| Z2 | Z*+10+6 \| 11+10+8+6+5+24 | W1 \| W2 \| W3 | w* \| 10+8+6+5+24 \| 11 | Not found | |
| *Mechanitis mazaeus* | Z1 \| Z2 | Z*+10+6 \| 3+19+10+8+6+5+24 | W1 \| W2 | w*+10+8+6+5+24 \| 3+19 | Not found | |
| *Mechanitis menapis* | Z | Z*+10+6 | W | w* | Not found | |
| *Mechanitis messenoides* | Z | Z*+10+6 | W | w* | 1 = 6 \| 14 | 5+12+2+3+28+23+20+19 \| 7+8+10+6+(5)+24 |
| *Mechanitis polymnia* | Z | Z*+10+6 | W | w*+8 | 1 = 14 \| 13 | 19+24+20+4+25+17 \| (4)+9+3+28 |

**Table S9. Result table for randomization test**

Mean values across *Mechanitis* and *Melinaea* genomes and within breakpoint regions (break) for a) $F_{ST}$, $D_{XY}$ and b) $\pi$. Result of a randomisation test of observed population statistic within breakpoint regions and permutated regions across the genome. Z-score and P-value at 5% significance level (lesser than or greater than randomised sample). Bold represent significant results after Bonferroni correction. The randomisation test was not performed in three of the comparisons due to limited number of informative sites.

A)

| Comparison Mec | $F_{ST}$ | SD | $F_{ST}$ break | SD | P-value | Z-score | $D_{XY}$ | SD | $D_{XY}$ break | SD | P-value | Z.score |
|---|---|---|---|---|---|---|---|---|---|---|---|---|
| *Mec. macrinus vs Mec. mazaeus* | 9.84E-02 | 7.32E-02 | 2.06E-01 | 1.18E-01 | **2.00E-05** | **6.238** | 1.32E-02 | 4.46E-03 | 1.26E-02 | 5.60E-03 | 1.99E-02 | 2.01E+00 |
| *Mec. macrinus vs Mec. menapis* | 2.25E-01 | 9.61E-02 | 2.80E-01 | 1.18E-01 | **2.20E-04** | **4.142** | 1.89E-02 | 5.06E-03 | 2.32E-02 | 6.93E-03 | **2.00E-05** | **4.63E+00** |
| *Mec. macrinus vs Mec. messenoides* | 2.35E-01 | 1.01E-01 | 3.63E-01 | 1.37E-01 | **2.00E-05** | **7.576** | 1.92E-02 | 5.13E-03 | 2.39E-02 | 8.02E-03 | **2.00E-05** | **6.06E+00** |
| *Mec. macrinus vs Mec. polymna* | 2.71E-01 | 9.51E-02 | 3.56E-01 | 1.30E-01 | **2.00E-05** | **5.064** | 1.94E-02 | 5.33E-03 | 2.05E-02 | 5.55E-03 | 5.19E-02 | 1.65E+00 |
| *Mec. mazaeus vs Mec. menapis* | 2.22E-01 | 9.27E-02 | 3.09E-01 | 1.21E-01 | **2.00E-05** | **5.154** | 1.86E-02 | 4.88E-03 | 2.14E-02 | 6.39E-03 | **1.02E-03** | **3.15E+00** |
| *Mec. mazaeus vs Mec. messenoides* | 2.40E-01 | 1.07E-01 | 3.94E-01 | 1.32E-01 | **2.00E-05** | **7.58** | 1.88E-02 | 4.95E-03 | 2.13E-02 | 8.01E-03 | **2.60E-04** | **3.45E+00** |
| *Mec. mazaeus vs Mec. polymnia* | 2.60E-01 | 1.04E-01 | 3.74E-01 | 1.32E-01 | **2.00E-05** | **5.162** | 1.90E-02 | 5.06E-03 | 1.90E-02 | 4.39E-03 | 4.12E-01 | -2.27E-01 |
| *Mec. menapis vs Mec. messenoides* | 2.21E-01 | 7.63E-02 | 2.92E-01 | 1.06E-01 | **2.00E-05** | **5.051** | 1.75E-02 | 5.01E-03 | 1.85E-02 | 6.69E-03 | 7.98E-03 | 2.40E+00 |
| *Mec. menapis vs Mec. polymnia* | 2.55E-01 | 8.33E-02 | 3.25E-01 | 1.23E-01 | **2.00E-05** | **5.332** | 1.98E-02 | 5.29E-03 | 2.15E-02 | 4.95E-03 | 5.41E-02 | 1.62E+00 |
| *Mec. messenoides vs Mec. polymnia* | 2.42E-01 | 9.23E-02 | 3.34E-01 | 1.16E-01 | **2.00E-05** | **5.242** | 1.86E-02 | 5.21E-03 | 1.79E-02 | 4.82E-03 | 1.23E-01 | -1.14E+00 |

| Comparison Mel | $F_{ST}$ | SD | $F_{ST}$ break | SD | P-value | Z-score | $D_{XY}$ | SD | $D_{XY}$ break | SD | P-value | Z.score |
|---|---|---|---|---|---|---|---|---|---|---|---|---|
| *Mel. isocomma vs Mel. marsaeus* | 2.73E-01 | 2.13E-01 | 2.69E-01 | 1.86E-01 | NA | NA | 2.57E-02 | 8.48E-03 | 2.16E-02 | 9.00E-03 | NA | NA |
| *Mel. isocomma vs Mel. menophilus* | 3.11E-01 | 2.02E-01 | 4.94E-01 | 1.92E-01 | 1.67E-02 | 2.576 | 2.79E-02 | 9.37E-03 | 3.42E-02 | 1.11E-02 | 1.42E-02 | 2.58E+00 |
| *Mel. isocomma vs Mel. mothone* | 3.13E-01 | 1.82E-01 | 1.98E-01 | 4.55E-02 | NA | NA | 3.07E-02 | 8.52E-03 | 2.31E-02 | 6.22E-03 | NA | NA |
| *Mel. ludovica vs Mel. isocomma* | 4.00E-01 | 2.57E-01 | 2.53E-01 | 4.95E-01 | **6.60E-04** | **-3.931** | 3.54E-02 | 7.64E-03 | 4.36E-02 | 7.23E-03 | **2.00E-05** | **4.64E+00** |
| *Mel. ludovica vs Mel. marsaeus* | 3.75E-01 | 2.06E-01 | 2.98E-01 | 3.39E-01 | **3.78E-03** | **-2.634** | 3.26E-02 | 8.03E-03 | 4.27E-02 | 8.53E-03 | **2.00E-05** | **5.48E+00** |
| *Mel. ludovica vs Mel. menophilus* | 3.65E-01 | 1.95E-01 | 3.41E-01 | 2.61E-01 | 1.18E-01 | -1.091 | 3.42E-02 | 8.26E-03 | 4.13E-02 | 8.25E-03 | **2.00E-05** | **4.65E+00** |
| *Mel. ludovica vs Mel. mothone* | 3.30E-01 | 1.61E-01 | 3.79E-01 | 1.82E-01 | 1.04E-01 | 1.297 | 3.57E-02 | 7.66E-03 | 4.46E-02 | 7.74E-03 | **2.00E-05** | **5.38E+00** |
| *Mel. menophilus vs Mel. marsaeus* | 1.39E-01 | 1.00E-01 | 2.52E-01 | 1.30E-01 | 1.13E-02 | 3.478 | 1.77E-02 | 7.02E-03 | 1.85E-02 | 6.61E-03 | 2.90E-01 | 5.44E-01 |
| *Mel. mothone vs Mel. marsaeus* | 8.79E-02 | 8.07E-02 | 1.19E-01 | 1.31E-01 | NA | NA | 1.85E-02 | 6.61E-03 | 1.74E-02 | 9.14E-03 | NA | NA |
| *Mel. mothone vs Mel. menophilus* | 1.20E-01 | 1.03E-01 | 2.52E-01 | 1.15E-01 | 5.22E-03 | 3.791 | 2.06E-02 | 7.57E-03 | 2.09E-02 | 6.58E-03 | 4.17E-01 | 2.08E-01 |

B)

| **Comparison Mec** | **Species** | **π** | **SD** | **π break** | **SD** | **P-value** | **Z-score** |
|---|---|---|---|---|---|---|---|
| *Mec. macrinus vs Mec. mazaeus* | *Mec. macrinus* | 1.09E-02 | 4.28E-03 | 8.61E-03 | 3.71E-03 | 8.41E-02 | -1.389 |
| *Mec. macrinus vs Mec. mazaeus* | *Mec. mazaeus* | 1.12E-02 | 4.35E-03 | 8.13E-03 | 4.13E-03 | 1.29E-01 | -1.137 |
| *Mec. macrinus vs Mec. menapis* | *Mec. macrinus* | 1.09E-02 | 4.27E-03 | 9.02E-03 | 4.15E-03 | **3.86E-03** | **-2.673** |
| *Mec. macrinus vs Mec. menapis* | *Mec. menapis* | 1.16E-02 | 3.44E-03 | 1.34E-02 | 5.41E-03 | **5.80E-04** | **3.243** |
| *Mec. macrinus vs Mec. messenoides* | *Mec. macrinus* | 1.09E-02 | 4.27E-03 | 8.94E-03 | 4.20E-03 | **4.20E-04** | **-3.334** |
| *Mec. macrinus vs Mec. messenoides* | *Mec. messenoides* | 1.07E-02 | 3.98E-03 | 8.58E-03 | 3.74E-03 | **2.20E-04** | **-3.654** |
| *Mec. macrinus vs Mec. polymna* | *Mec. macrinus* | 1.09E-02 | 4.28E-03 | 9.01E-03 | 3.86E-03 | **1.46E-03** | **-3.021** |
| *Mec. macrinus vs Mec. polymna* | *Mec. polymniaEast* | 1.13E-02 | 4.10E-03 | 9.37E-03 | 3.60E-03 | 5.26E-03 | -2.515 |
| *Mec. mazaeus vs Mec. menapis* | *Mec. mazaeus* | 1.13E-02 | 4.34E-03 | 7.91E-03 | 4.00E-03 | **4.20E-04** | **-3.483** |
| *Mec. mazaeus vs Mec. menapis* | *Mec. menapis* | 1.16E-02 | 3.47E-03 | 1.19E-02 | 4.18E-03 | 2.69E-01 | 0.606 |
| *Mec. mazaeus vs Mec. messenoides* | *Mec. mazaeus* | 1.13E-02 | 4.34E-03 | 8.38E-03 | 4.33E-03 | **2.00E-05** | **-4.121** |
| *Mec. mazaeus vs Mec. messenoides* | *Mec. messenoides* | 1.08E-02 | 3.98E-03 | 7.33E-03 | 2.86E-03 | **2.00E-05** | **-4.696** |
| *Mec. mazaeus vs Mec. polymnia* | *Mec. mazaeus* | 1.13E-02 | 4.35E-03 | 8.15E-03 | 3.81E-03 | **2.60E-04** | **-3.759** |
| *Mec. mazaeus vs Mec. polymnia* | *Mec. polymniaEast* | 1.13E-02 | 4.10E-03 | 8.55E-03 | 2.97E-03 | **6.00E-05** | **-3.813** |
| *Mec. menapis vs Mec. messenoides* | *Mec. menapis* | 1.16E-02 | 3.48E-03 | 1.19E-02 | 4.00E-03 | 8.88E-02 | 1.356 |
| *Mec. menapis vs Mec. messenoides* | *Mec. messenoides* | 1.07E-02 | 3.98E-03 | 7.51E-03 | 3.63E-03 | **3.02E-03** | **-2.898** |
| *Mec. menapis vs Mec. polymnia* | *Mec. menapis* | 1.16E-02 | 3.49E-03 | 1.17E-02 | 3.26E-03 | 4.99E-01 | 8.00E-03 |
| *Mec. menapis vs Mec. polymnia* | *Mec. polymniaEast* | 1.13E-02 | 4.10E-03 | 8.97E-03 | 3.17E-03 | **3.80E-04** | **-3.23** |
| *Mec. messenoides vs Mec. polymnia* | *Mec. messenoides* | 1.07E-02 | 3.98E-03 | 7.63E-03 | 3.12E-03 | **2.00E-05** | **-4.936** |
| *Mec. messenoides vs Mec. polymnia* | *Mec. polymniaEast* | 1.13E-02 | 4.10E-03 | 9.04E-03 | 3.14E-03 | **5.80E-04** | **-3.145** |

| **Comparison Mel** | **Species** | **π** | **SD** | **π break** | **SD** | **P-value** | **Z-score** |
|---|---|---|---|---|---|---|---|
| *Mel. isocomma vs Mel. marsaeus* | *Mel. isocomma* | 1.29E-02 | 6.67E-03 | 6.81E-03 | 4.11E-03 | NA | NA |
| *Mel. isocomma vs Mel. marsaeus* | *Mel. marsaeus* | 1.24E-02 | 5.19E-03 | 1.10E-02 | 6.47E-03 | NA | NA |
| *Mel. isocomma vs Mel. menophilus* | *Mel. isocomma* | 1.29E-02 | 6.67E-03 | 8.92E-03 | 5.43E-03 | 1.80E-02 | -2.152 |
| *Mel. isocomma vs Mel. menophilus* | *Mel. menophilus* | 1.40E-02 | 7.14E-03 | 9.12E-03 | 3.86E-03 | 5.42E-03 | -2.675 |
| *Mel. isocomma vs Mel. mothone* | *Mel. isocomma* | 1.29E-02 | 6.67E-03 | 1.52E-02 | 6.08E-03 | NA | NA |
| *Mel. isocomma vs Mel. mothone* | *Mel. mothone* | 1.76E-02 | 7.05E-03 | 1.56E-02 | 7.07E-05 | NA | NA |
| *Mel. ludovica vs Mel. isocomma* | *Mel. isocomma* | 1.29E-02 | 6.66E-03 | 8.95E-03 | 6.19E-03 | 1.98E-01 | 0.77 |
| *Mel. ludovica vs Mel. isocomma* | *Mel. ludovica* | 7.32E-03 | 4.99E-03 | 7.54E-03 | 3.45E-03 | **8.20E-04** | **-3.472** |
| *Mel. ludovica vs Mel. marsaeus* | *Mel. ludovica* | 7.32E-03 | 4.99E-03 | 7.54E-03 | 3.45E-03 | 2.06E-01 | 0.743 |
| *Mel. ludovica vs Mel. marsaeus* | *Mel. marsaeus* | 1.24E-02 | 5.17E-03 | 8.86E-03 | 5.12E-03 | **4.60E-04** | **-3.929** |
| *Mel. ludovica vs Mel. menophilus* | *Mel. ludovica* | 7.32E-03 | 5.00E-03 | 7.43E-03 | 3.19E-03 | 2.58E-01 | 0.567 |
| *Mel. ludovica vs Mel. menophilus* | *Mel. menophilus* | 1.41E-02 | 7.12E-03 | 1.06E-02 | 6.67E-03 | **1.02E-03** | **-3.203** |
| *Mel. ludovica vs Mel. mothone* | *Mel. ludovica* | 7.32E-03 | 4.99E-03 | 7.52E-03 | 3.34E-03 | 2.01E-01 | 0.763 |
| *Mel. ludovica vs Mel. mothone* | *Mel. mothone* | 1.77E-02 | 7.01E-03 | 1.38E-02 | 7.90E-03 | **1.22E-03** | **-3.274** |
| *Mel. menophilus vs Mel. marsaeus* | *Mel. marsaeus* | 1.24E-02 | 5.18E-03 | 1.06E-02 | 5.68E-03 | 5.26E-03 | -2.63E+00 |
| *Mel. menophilus vs Mel. marsaeus* | *Mel. menophilus* | 1.40E-02 | 7.14E-03 | 9.13E-03 | 4.30E-03 | 9.77E-02 | -1.331 |
| *Mel. mothone vs Mel. marsaeus* | *Mel. marsaeus* | 1.24E-02 | 5.19E-03 | 1.11E-02 | 6.71E-03 | NA | NA |
| *Mel. mothone vs Mel. marsaeus* | *Mel. mothone* | 1.76E-02 | 7.05E-03 | 1.26E-02 | 6.31E-03 | NA | NA |
| *Mel. mothone vs Mel. menophilus* | *Mel. menophilus* | 1.40E-02 | 7.14E-03 | 9.17E-03 | 3.83E-03 | 2.68E-02 | -1.986 |
| *Mel. mothone vs Mel. menophilus* | *Mel. mothone* | 1.76E-02 | 7.05E-03 | 1.33E-02 | 5.99E-03 | 1.09E-02 | -2.351 |
